# The self-peptide repertoire plays a critical role in transplant tolerance induction

**DOI:** 10.1101/2020.11.09.359968

**Authors:** Eric T. Son, Pouya Faridi, Moumita Paul-Heng, Mario Leong, Kieran English, Sri H. Ramarathinam, Asolina Braun, Nadine L. Dudek, Ian E. Alexander, Leszek Lisowski, Patrick Bertolino, David G. Bowen, Anthony W. Purcell, Nicole A. Mifsud, Alexandra F. Sharland

**Affiliations:** Transplantation Immunobiology Group, University of Sydney Central Clinical School, Charles Perkins Centre, Faculty of Medicine and Health, Sydney, NSW, Australia; Infection and Immunity Program, Department of Biochemistry and Molecular Biology, Biomedicine Discovery Institute, Monash University, Clayton, Victoria, Australia; Liver Immunology Group and AW Morrow Gastroenterology and Liver Centre, The University of Sydney and Royal Prince Alfred Hospital, Sydney, NSW, Australia; Gene Therapy Research Unit, Children’s Medical Research Institute, The University of Sydney, Faculty of Medicine and Health and Sydney Children’s Hospitals Network, Westmead, NSW, Australia; The University of Sydney, Sydney Medical School, Discipline of Child and Adolescent Health, Westmead, NSW, Australia; Translational Vectorology Research Unit, Children’s Medical Research Institute, Faculty of Medicine and Health, The University of Sydney, Westmead, Australia; Vector and Genome Engineering Facility, Children’s Medical Research Institute, Faculty of Medicine and Health, The University of Sydney, Westmead, Australia; Military Institute of Medicine, Laboratory of Molecular Oncology and Innovative Therapies, 04-141 Warsaw, Poland

## Abstract

While direct allorecognition underpins both solid organ allograft rejection and tolerance induction, the specific molecular targets of most directly-alloreactive CD8^+^ T cells have not been defined. In this study, we used a combination of genetically-engineered MHC class I (MHC I) constructs, mice with a hepatocyte-specific mutation in the class I antigen-presentation pathway and immunopeptidomic analysis to provide definitive evidence for the contribution of the peptide cargo of allogeneic MHC I molecules to transplant tolerance induction. We established a systematic approach for the discovery of directly-recognised pMHC epitopes, and identified 17 strongly immunogenic H-2K^b^-associated peptides recognised by CD8^+^ T cells from B10.BR (H-2^k^) mice, 13 of which were also recognised by BALB/c (H-2^d^) mice. As few as five different tetramers used together were able to identify a high proportion of alloreactive T cells within a polyclonal population, suggesting that there are immunodominant allogeneic MHC-peptide complexes that can account for a large component of the alloresponse.

## Introduction

Allorecognition may result in either graft rejection, graft versus host disease or transplantation tolerance, depending upon the context in which the recipient immune system encounters the allogeneic major histocompatibility complex (MHC). When donor MHC class I (MHC I) molecules are expressed in the hepatocytes of recipient mice, subsequent skin or pancreatic islet grafts bearing the same donor allomorph are accepted indefinitely(1–3). Insights from this model can inform our understanding of allorecognition and transplant tolerance induction more broadly. MHC I molecules are ubiquitously expressed, and display a range of endogenous peptides (the class I immunopeptidome) reflecting protein turnover and normal cellular processes(4). While tolerance induction depends upon recognition of intact donor MHC I molecules by recipient CD8^+^ T cells(3), the contribution of the self-peptide cargo of these molecules to tolerance induction in this setting is unknown. Many alloreactive T cell clones recognise epitopes comprising allogeneic MHC I molecules complexed with self-peptides(5–12). Conversely, peptide-independent direct allorecognition is also described(7, 13, 14), and the ability of peptide-independent cytotoxic T lymphocyte (CTL) clones to bring about rapid destruction of allogeneic skin grafts has been demonstrated(15). At the level of a polyclonal alloresponse in vivo, there is limited information about the role of the donor’s tissue-specific class I immunopeptidome in allorecognition and consequent immune responses that impact on graft survival.

Here, we examined the role of the liver immunopeptidome in transplantation tolerance induction using two different approaches. Firstly, we engineered liver-specific adeno-associated viral (AAV) vectors encoding H-2K^b^ or H-2K^d^ as a single chain trimer (SCT) comprising the polymorphic heavy chain (HC), *β*2 microglobulin (β2m) light chain and a defined covalently-bound peptide species(16, 17), thus excluding presentation of endogenous peptides by the allogeneic MHC I when these constructs were expressed in recipient hepatocytes (Figure 1A). In parallel, we introduced a global shift in the repertoire of peptides bound to allogeneic H-2K^d^. To accomplish this, we generated recipient mice in which the transporter associated with antigen processing (TAP1) protein was deficient only in hepatocytes (*Tap1*KOHep, H-2^b^), and designed a construct expressing the H-2K^d^ HC with a modification (YCAC) which stabilises the molecule and permits occupancy by lower affinity peptides(18). Induction of tolerance to subsequent donor skin grafts was blocked by these manipulations, establishing that the endogenous peptide repertoire of hepatocytes makes an essential contribution to transplant tolerance induction.

**Figure 1.**
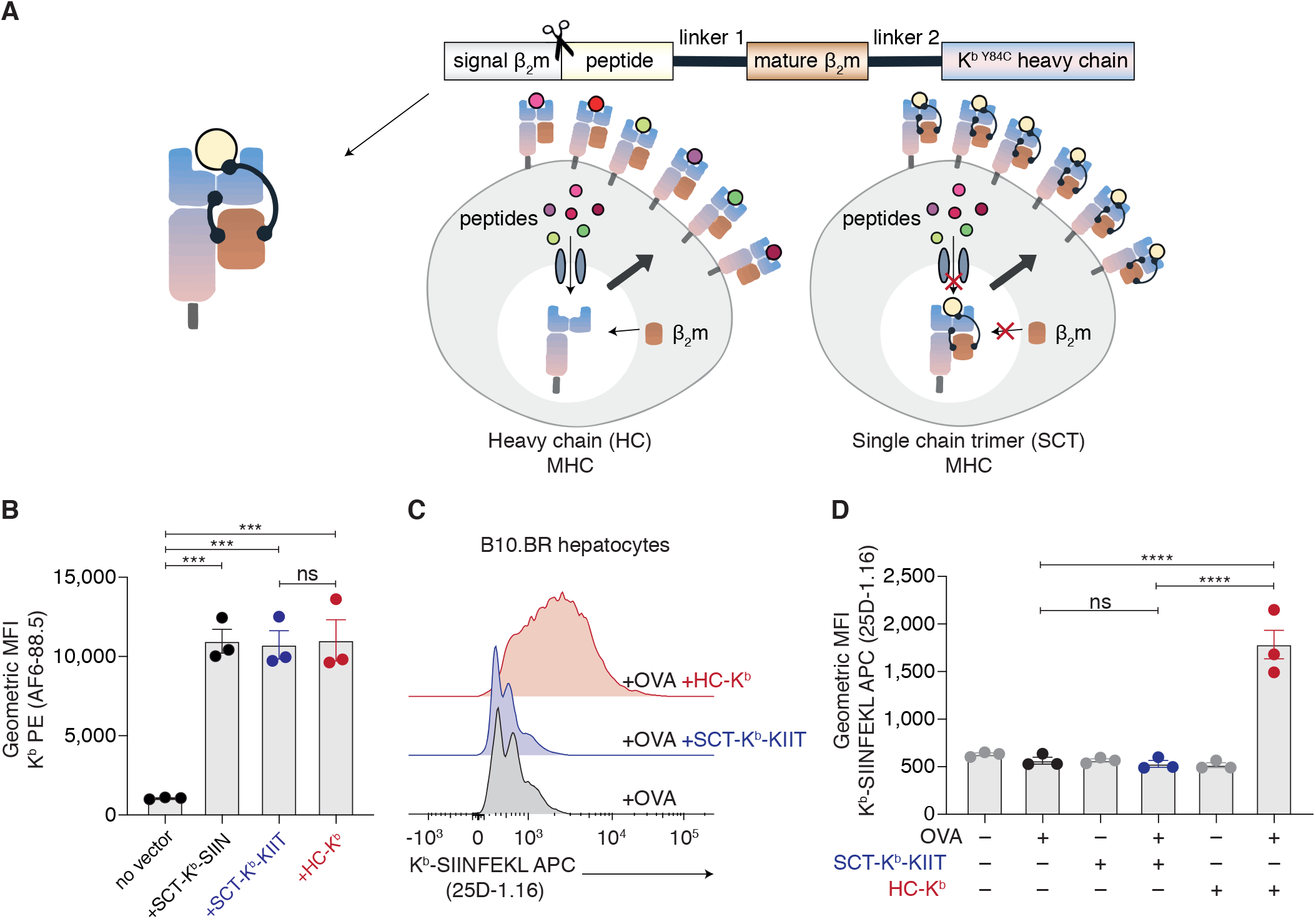
SCT constructs express a single pMHC epitope at the cell surface. **(A)** Schematic diagram of SCT pMHC constructs. MHC I HC expressed in hepatocytes can bind endogenous peptides and associate with native β2m to form a wide repertoire of pMHC complexes. Conversely, SCT constructs exclude binding of endogenous peptides and present a single type of pMHC epitope on the cell surface. **(B-D)** B10.BR mice were transduced with AAV-SCT-K^b^-SIIN, AAV-SCT-K^b^-KIIT or AAV-HC-K^b^ with or without AAV-OVA. Flow cytometric analysis of isolated hepatocytes (d7) is shown. **(B)** Equivalent levels of H-2K^b^ cell surface expression were demonstrated using the mAb AF6-88.5. **(C,D)** Presentation of SIINFEKL (OVA_257-264_) peptide bound to H-2K^b^ was detected using an anti-K^b^-SIINFEKL mAb (25D-1.16). Co-transduction with AAV-OVA permitted presentation of SIINFEKL by hepatocytes transduced with AAV-HC-K^b^. Conversely, the SCT-K^b^-KIIT construct excluded presentation of endogenously-processed SIINFEKL peptide. For **(B)** and **(D)** n = 3 biological replicates per group, mean ± SEM are shown, Statistical analysis comprised one-way ANOVA with Sidak’s multiple comparison test: ns, not significant, *** p < 0.001, **** p < 0.0001. Panel **(C)** depicts one representative experiment (from n = 3).

Differential responses by alloreactive T cells to MHC I allomorphs expressed by various target tissues is cited as support for peptide-specific allorecognition(6, 19, 20). Given that expression of allogeneic donor MHC I HC in recipient hepatocytes attenuates responses against both donor skin grafts and splenocyte stimulators(1, 3), we hypothesised that peptides critical for tolerance induction would be found within a subset shared by these tissues. We identified these peptides and assessed their recognition by activated alloreactive B10.BR T cells. Of 100 peptides selected for screening, 17 were bound by more than 5% of this T cell population, and 13 of these peptide-MHC (pMHC) epitopes were also recognised by BALB/c mice. As few as five different pMHC tetramers used together were able to identify around 40% of alloreactive T cells within a polyclonal population, suggesting that there are immunodominant allogeneic MHC-peptide complexes that can account for a large proportion of the alloresponse. Accordingly, this panel enabled quantitation and phenotyping of alloreactive T cells in a model of secondary skin graft tolerance or rejection. The findings of this study represent a significant advance in our understanding of the role of endogenous peptides in direct T cell alloreactivity, and a springboard for further knowledge gain and technological development.

## Results

### Single chain trimer constructs exclude presentation of endogenous peptides

In preceding studies(1, 3), we have used AAV vectors encoding the donor MHC I HC. Within transduced hepatocytes, allogeneic HC associates with native β2m and the resulting heterodimers are loaded with a repertoire of endogenous peptides (Figure 1A). To express allogeneic MHC I at high levels on recipient hepatocytes while excluding the presentation of naturally processed peptides, we engineered SCT constructs, each encoding the HC of H-2K^b^, β2m and a single, defined H-2K^b^-restricted peptide [SIINFEKL (SIIN) or KIITYRNL (KIIT), Figure 1A], and packaged them in hepatocyte-specific AAV2/8 vectors. Sequences are shown in Supplementary Figure 1. Transgene expression in hepatocytes was close to maximal by d7 following intravenous (iv) inoculation, and persisted through to at least d100, no significant increases in serum aspartate aminotransferase (AST) or alanine aminotransferase (ALT) levels were observed, and minimal cellular infiltration was detected by histology (Supplementary Figure 2). SCT molecules were expressed on transduced hepatocytes at equivalent levels to the heterotrimer formed by transgenic H-2K^b^ HC with native β2m and peptide (Figure 1B). To demonstrate exclusion of naturally processed peptides, we co-transduced B10.BR (H-2^k^) hepatocytes with AAV vectors encoding full-length chicken ovalbumin (OVA) and either HC-K^b^ or SCT-K^b^-KIIT, and stained them with a monoclonal antibody, 25D-1.16, which is specific for the OVA peptide SIINFEKL complexed with K^b^. K^b^-SIINFEKL was only detected at the surface of cells co-transduced with HC-K^b^ and not those expressing SCT-K^b^-KIIT (Figure 1C-D). We extended this analysis to the broader endogenous peptide repertoire of K^b^-transduced hepatocytes using immunoaffinity purification with the H-2K^b^-specific antibody K9-178, followed by reverse phase high performance liquid chromatography (RP-HPLC) to collect peptide-containing fractions that were then analysed using liquid chromatography with tandem mass spectrometry (LC-MS/MS) to identify bound peptides (Figure 2A). The K^b^ binding motif of peptides eluted from HC-K^b^-transduced hepatocytes mirrored that obtained from C57BL/6 (H-2^b^) hepatocytes (Figure 2B). While there was greater diversity among the unique peptides isolated from HC-K^b^-transduced hepatocytes (Figure 2C), there was substantial overlap between the peptide repertoires, with >90% of the peptides from C57BL/6 hepatocytes also identified in HC-K^b^-transduced B10.BR (Figure 2D). In contrast, almost no K^b^-bound peptides could be identified in association with SCT-K^b^-KIIT, confirming exclusion of the endogenous peptide repertoire from presentation by this molecule (Figure 2C). Hepatocytes transduced with SCT-K^b^-KIIT did not manifest any generalised defect in antigen processing, with both the numbers and repertoire of peptides isolated from the native allomorph H-2K^k^ being similar across all groups (Figures 2E-F). The full peptide dataset can be found in Supplementary Data 1.

**Figure 2.**
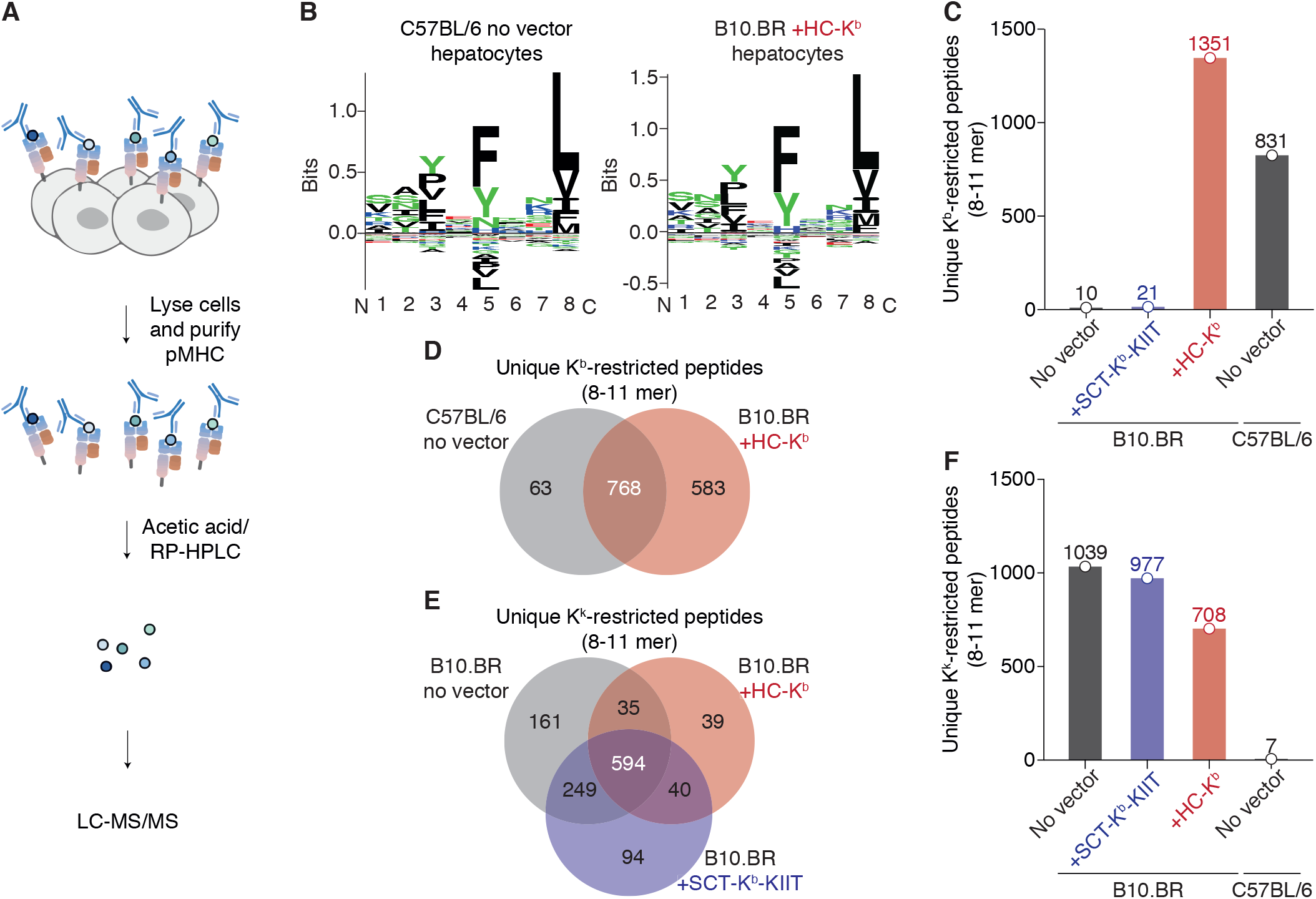
SCT-K^b^-KIIT constructs exclude presentation of endogenous peptides. **(A)** Control B10.BR hepatocytes, B10.BR hepatocytes transduced with either AAV-HC-K^b^ or AAV-SCT-K^b^-KIIT or control C57BL/6 hepatocytes underwent immunoaffinity purification in order to identify unique H-2K^b^ and H-2K^k^ peptides. A false discovery rate (FDR) of 5% was employed. Two replicate experiments were performed. Samples from 4 mice were pooled/group for each replicate, and combined lists of peptides present in either or both replicates were generated. **(B)** H-2K^b^-binding motifs were generated from a non-redundant list of 8-11-mer peptides using the GibbsCluster-2.0 algorithm. The binding motif obtained from B10.BR hepatocytes transduced with AAV-HC-K^b^ closely reflected that from C57BL/6 (H-2^b^) hepatocytes. **(C,D)** Abundant K^b^-binding peptides were isolated from both C57BL/6 and AAV-HC-K^b^-transduced B10.BR hepatocytes, with significant overlap between repertoires, while very few K^b^-binding peptides could be eluted from B10.BR hepatocytes transduced with AAV-SCT-K^b^-KIIT. **(E,F)** Comparable numbers of K^k^-binding peptides were isolated from all B10.BR hepatocyte samples, but not from C57BL/6 mice which lack this allomorph. **(C,F)** Bars represent the cumulative number of unique peptides obtained from replicate experiments.

### SCT-K^b^-KIIT induces tolerance in alloreactive T cells expressing the cognate T cell receptor but not in a polyclonal alloreactive population

We first verified that SCT molecules were recognised by cells bearing their cognate TCRs in a manner analogous to the native epitope. Pulsing RMA-S cells with synthetic peptides KIITYRNL, SIINFEKL and AAAAFAAL (minimum binding requirement for H-2K^b^) stabilised K^b^ expression on the cell surface (Supplementary Figure 3A-B). Similar levels of surface expression were achieved by transient transfection of RMA-S cells with the corresponding SCT constructs (Supplementary Figure 3C). Using interferon-gamma (IFN-*γ*) ELISPOT, we found a strong, specific response to SCT-K^b^-KIIT by Des-RAG TCR-transgenic T cells which express the cognate TCR; SCT-K^b^-SIIN was similarly recognised by OT-I-RAG T cells (Supplementary Figure 3D). Following adoptive transfer into B10.BR mice transduced with SCT-K^b^-KIIT, Des-RAG T cells proliferated vigorously, expanding more than those transferred to the positive control 178.3 (H-2^k^ + K^b^) mice after 2 days (p<0.0001, Supplementary Figure 3E-F). The polyclonal responder population of liver leukocytes from B10.BR mice also contained CD8^+^ T cells which were activated in response to transduction with SCT-K^b^-KIIT. Naïve or primed B10.BR mice were inoculated with SCT-K^b^-KIIT, and liver leukocytes isolated 7 days later (Supplementary Figure 4A). Activated CD8^+^ T cells, defined as CD44^+^PD-1^hi^, increased upon priming or transduction with SCT-K^b^-KIIT, with a further augmentation when primed mice received SCT-K^b^-KIIT vector (Supplementary Figure 4B). Activated CD8^+^ T cells from transduced mice recognised K^b^-KIIT dextramers but not dextramers of the self-pMHC complex K^k^-EEEPVKKI, while PD-1^-^ bystander CD8^+^ T cells did not bind either dextramer (Supplementary Figure 4C-D).

Next, we determined that Des-RAG T cells alone were capable of rejecting K^b^-bearing 178.3 skin grafts upon adoptive transfer to B10.BR-RAG recipients (Figure 3A-B). Graft survival was inversely related to the T cell dose; 50,000 transferred cells yielded a median graft survival comparable to that in B10.BR mice with a polyclonal K^b^-reactive T cell repertoire and this dose was used in subsequent experiments. All grafts to reconstituted B10.BR-RAG mice treated with SCT-K^b^-KIIT survived indefinitely, whereas no graft survival prolongation was observed in reconstituted mice receiving the control vector SCT-K^b^-SIIN (Figure 3C). Surviving grafts appeared normal macroscopically (Figure 3D) and upon histology (Figure 3E) with continued expression of H-2K^b^. In contrast, transduction of wild-type B10.BR mice with either SCT-K^b^-KIIT or SCT-K^b^-SIIN only briefly delayed graft rejection (Figure 3F).

**Figure 3.**
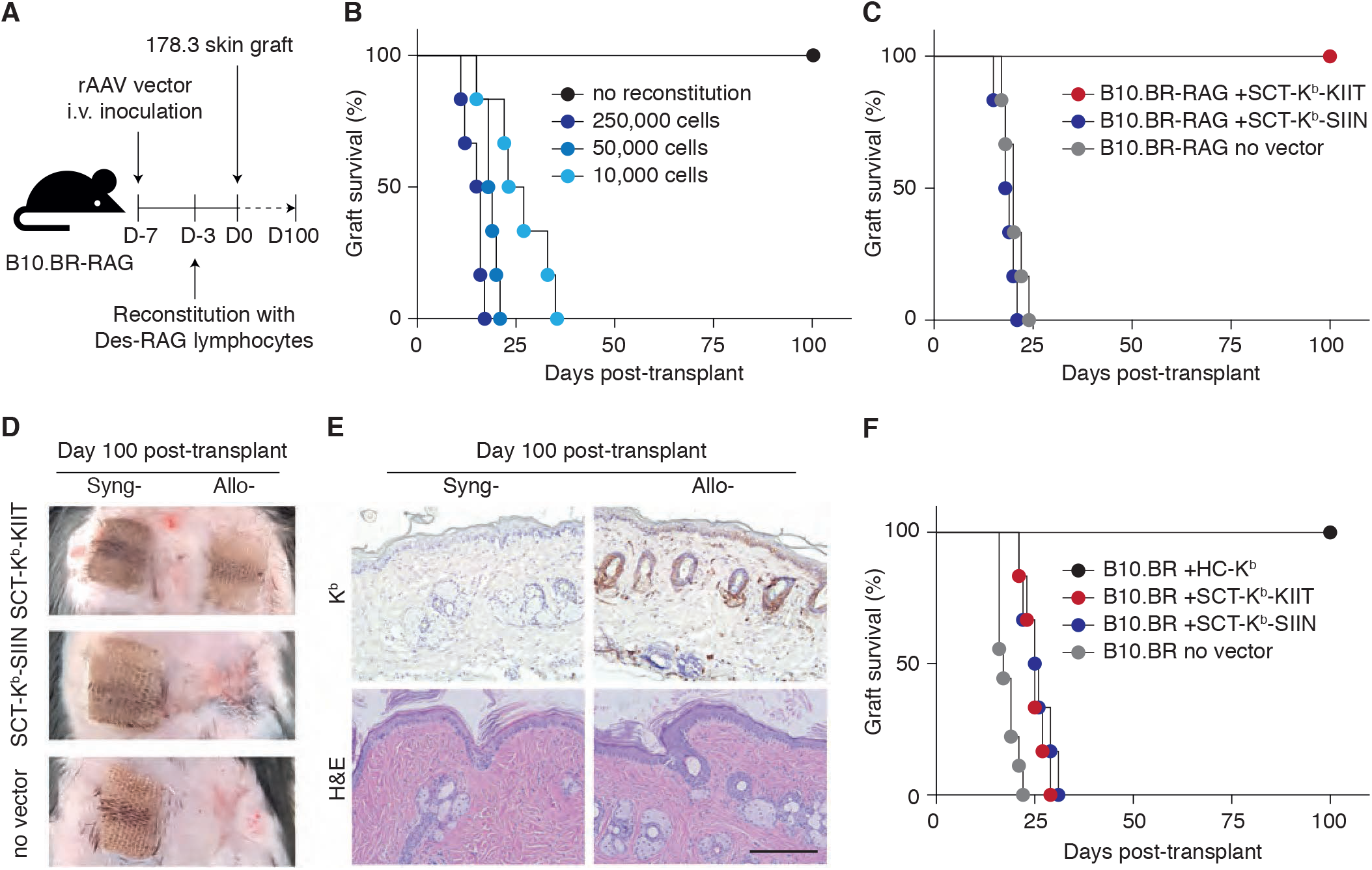
Expression of the SCT-K^b^-KIIT construct in recipient hepatocytes prevents skin graft rejection mediated by alloreactive CD8^+^ T cells expressing the cognate TCR but does not induce tolerance in a polyclonal alloreactive population. **(A)** B10.BR-RAG mice were reconstituted with 1.0×10^4^ -2.5×10^5^ Des-RAG lymphocytes, 3 days prior to receiving a K^b^-bearing 178.3 skin graft. Some mice were also inoculated with AAV vectors. **(B)** 178.3 skin grafts survived indefinitely on immunodeficient B10.BR-RAG hosts. Progressive shortening in survival accompanied adoptive transfer of increasing cell numbers (Mantel-Cox log-rank test for trend: p < 0.0001, n = 6 per group). A dose of 50,000 cells was used subsequently. **(C)** Inoculation of reconstituted B10.BR-RAG recipients with AAV-SCT-K^b^-KIIT resulted in indefinite graft survival, whereas treatment with AAV-SCT-K^b^-SIIN offered no survival prolongation compared with that in untransduced recipients; median survival time (MST) of 18.5 days versus 20 days (p = 0.2, p = 0.0005 between mice receiving AAV-SCT-K^b^-KIIT or AAV-SCT-K^b^-SIIN, all groups n = 6). **(D)** Representative macroscopic images (from n = 3) demonstrate continued allograft survival in reconstituted B10.BR-RAG mice treated with AAV-SCT-K^b^-KIIT but not other groups at 100 days after transplantation. **(E)** Representative IHC and H&E (n = 6) images showing syngeneic (B10.BR-RAG) and allogeneic (178.3) skin grafts 100 days post-transplant (scale bar: 100 µm). Skin transplants are morphologically normal, with persistent expression of H-2K^b^ in 178.3 grafts. **(F)** Immunosufficient B10.BR mice were inoculated with either AAV-HC-K^b^, AAV-SCT-K^b^-KIIT or AAV-SCT-K^b^-SIIN (n = 6 per group) or were not transduced (n = 9). 7 days post-inoculation, the mice received 178.3 skin grafts. In these mice, expression of either SCT-K^b^-KIIT or SCT-K^b^-SIIN led to a modest increase in graft survival (MST of 25 days versus 17 days in no vector controls, p = 0.0007). However, only expression of H-2K^b^ loaded with the endogenous peptide repertoire was able to induce tolerance to 178.3 skin grafts. p = 0.0005 between B10.BR inoculated with AAV-HC-K^b^ and either AAV-SCT-K^b^-KIIT or AAV-SCT-K^b^-SI-IN. **(C,F)** Mantel-Cox log-rank test.

### The YCAC mutation alters the repertoire of H-2K^d^-bound peptides in both *TAP1*KOHep and C57BL/6 mice

Mice with a conditional deletion of *Tap1* in hepatocytes (*Tap1*KOHep, H-2^b^) express trace amounts of the native MHC I allomorphs H-2K^b^ and H-2D^b^ at the hepatocyte surface (Figure 4A-D). Importantly, expression of these molecules on most other cell types within the liver, in other tissues (such as spleen, thymus and lymph node), and in *Tap1*^fl/fl^ control mice was normal (Figure 4A-D and Supplementary Figure 5). Transduction of *Tap1*KOHep mice with a vector encoding the H-2K^d^ HC (HC-K^d^) resulted in surface expression of H-2K^d^ which was clearly positive with respect to untransduced controls, yet was reduced in comparison to that in C57BL/6 or *Tap1*^fl/fl^ mice (Figure 4E-F). Increasing the vector dose did not yield an appreciable increase in surface expression (data not shown), most likely because of instability of suboptimally-loaded H-2K^b^ molecules and their rapid recycling from the cell surface(21–23). To counter this, we designed a construct where the point mutations Y84C and A139C (YCAC) in the H-2K^d^ HC result in the formation of a disulphide bridge, which stabilises the molecule when empty or loaded with lower affinity peptides(18). This modification does not interfere with TCR recognition of the bound peptide(18). The construct sequence is shown in Supplementary Figure 1. Comparable strong expression of H-2K^d^ on the surface of hepatocytes was achieved in *Tap1*KOHep mice treated with HC-K^d^-YCAC and in *Tap1*^fl/fl^ mice receiving either the HC-K^d^ or HC-K^d^-YCAC vectors (Figure 4E-G). Immunoaffinity purification and LC-MS/MS were used to characterise the bound self-peptide repertoire of transduced hepatocytes. 9570 unique peptides were identified from C57BL/6 hepatocytes transduced with HC-K^d^ (B6/HC), compared with 7690, 7776, and 6417 unique peptides respectively, from C57BL/6, *Tap1*^fl/fl^ and *Tap1*KOHep transduced with HC-K^d^-YCAC (*i.e.* B6/YCAC, *Tap1*^fl/fl^/YCAC and *Tap1*KO/YCAC). The full dataset can be found in Supplementary Data 1, while distribution of peptide lengths and spectral intensities is shown in Supplementary Figure 6, along with the gene ontology analysis of the subcellular location and function of the source proteins. Whilst the sequences of eluted 9-mer peptides from B6/HC, B6/YCAC and *Tap1*^fl/fl^/YCAC corresponded to the canonical motif for H-2K^d^, with a tyrosine (Y) residue predominant at position 2, and leucine (L) or isoleucine (I) most frequently found at position 9, this motif was not observed for the peptides eluted from *Tap1*KO/YCAC (Figure 5A-B). Compared to B6/HC, just under 45% of peptides were common to B6/YCAC or *Tap1*^fl/fl^/YCAC, while 22% were shared with *Tap1*KOHep/YCAC (Figure 5C). Similarity across peptide repertoires was increased when the comparison was weighted for peptide abundance (Figure 5C). The peptide SYFPEITHI (SYFP) was common to all K^d^ repertoires, and comparable proportions of CD8^+^ T cells recognising K^d^-SYFP could be detected among the liver leukocytes isolated from primed *Tap1*KO/YCAC as well as from primed B6/YCAC and B6/HC (Figure 5D) consistent with published reports that the YCAC mutation alters the peptide repertoire but does not interfere with TCR recognition of presented peptides(18, 24).

**Figure 4.**
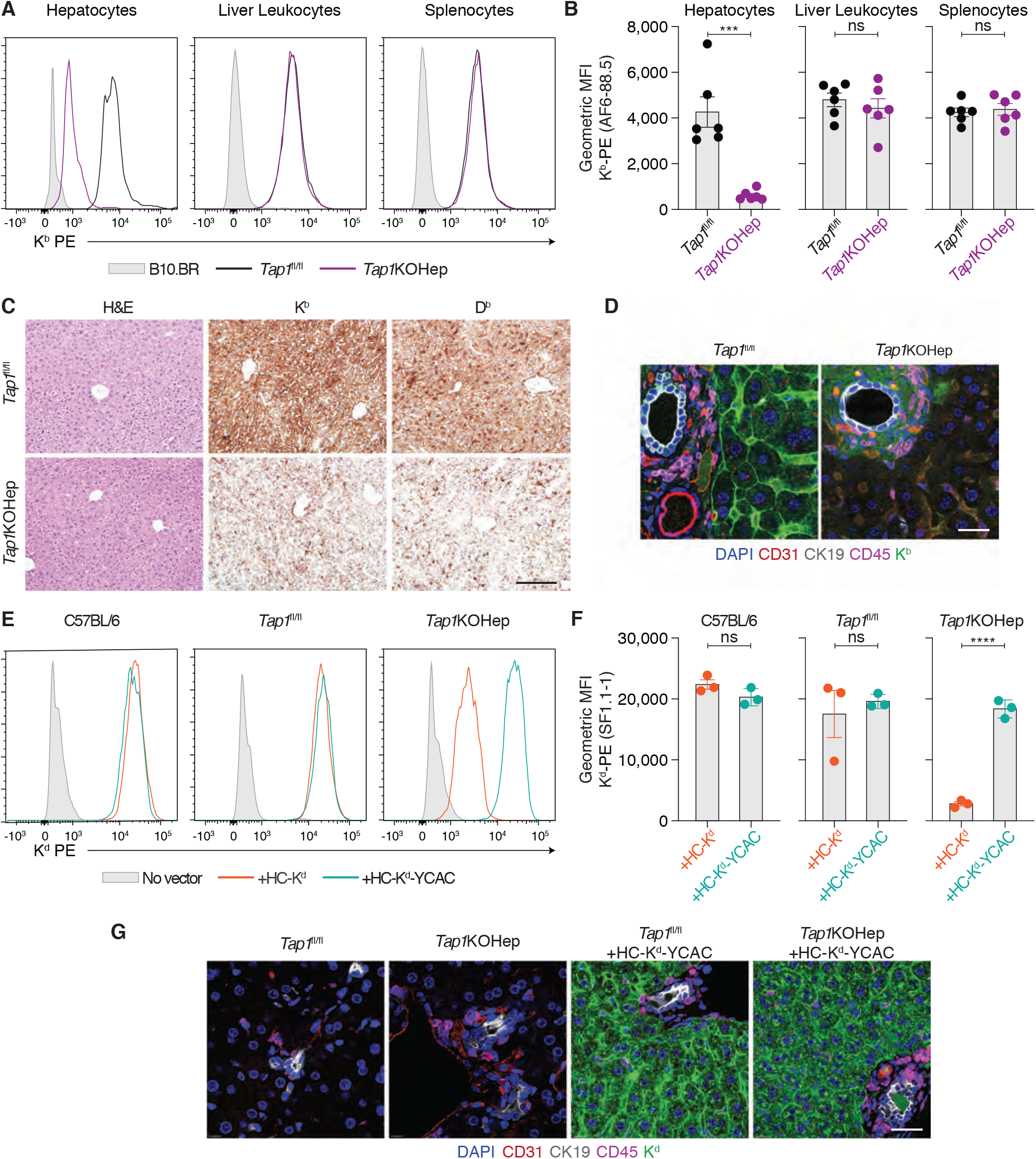
Hepatocytes from *Tap1K*O Hep mice express high levels of H-2K^d^ following transduction with AAV-HC-K^d^-YCAC. **(A-D)** Expression of K^b^ on liver leukocytes and splenocytes was equivalent between *Tap1*KOHep and *Tap1*^fl/fl^ mice, while much lower levels were detected on hepatocytes from *Tap1*KOHep. Representative flow plots from n = 6. **(C)** *Tap1*KOHep and *Tap1*^fl/fl^ livers are morphologically normal. IHC staining demonstrates expression of K^b^ and D^b^ in nonparenchymal cells from *Tap1*KOHep livers, while K^b^ and D^b^ are absent from hepatocytes in these mice (representative images, n = 3). **(D)** Thick sections (150 μm) from *Tap1*KOHep or *Tap1*^fl/fl^ livers, were stained with antibodies against H-2K^b^, CD31, CK19 and CD45. Confocal micrographs were obtained. K^b^ is ubiquitously expressed in *Tap1*^fl/fl^, but absent from hepatocytes in *Tap1*KOHep (representative images, n = 3). **(E,F)** While H-2K^d^ was clearly present on hepatocytes from *Tap1*KOHep transduced with AAV-HC-K^d^, expression did not reach that in TAP-sufficient mice transduced with the same vector. To achieve robust cell surface expression of K^d^, we designed a construct where point mutations (Y84C and A139C; YCAC) stabilise expression of K^d^ when empty or loaded with low-affinity peptides. Inoculation of *Tap1*KOHep mice with AAV-HC-K^d^-YCAC yielded comparable expression to that of TAP-sufficient mice transduced with AAV-HC-K^d^. Representative flow plots from n = 3. **(G)** Thick sections of *Tap1*KO/YCAC and *Tap1*^fl/fl^/YCAC livers were stained with antibodies against H-2K^d^, CD31, CK19 and CD45. Equivalent strong expression of K^d^ in hepatocytes, but not other cell types, was observed for both treatment groups (representative of n = 3). Scale bars **(C,E)** 100 μm, **(D,G)** 40 μm. **(B,F)** Mean ± SEM are shown. Statistical analysis comprised unpaired Student’s t-test: ns, not significant, *** p < 0.001, **** p < 0.0001.

**Figure 5.**
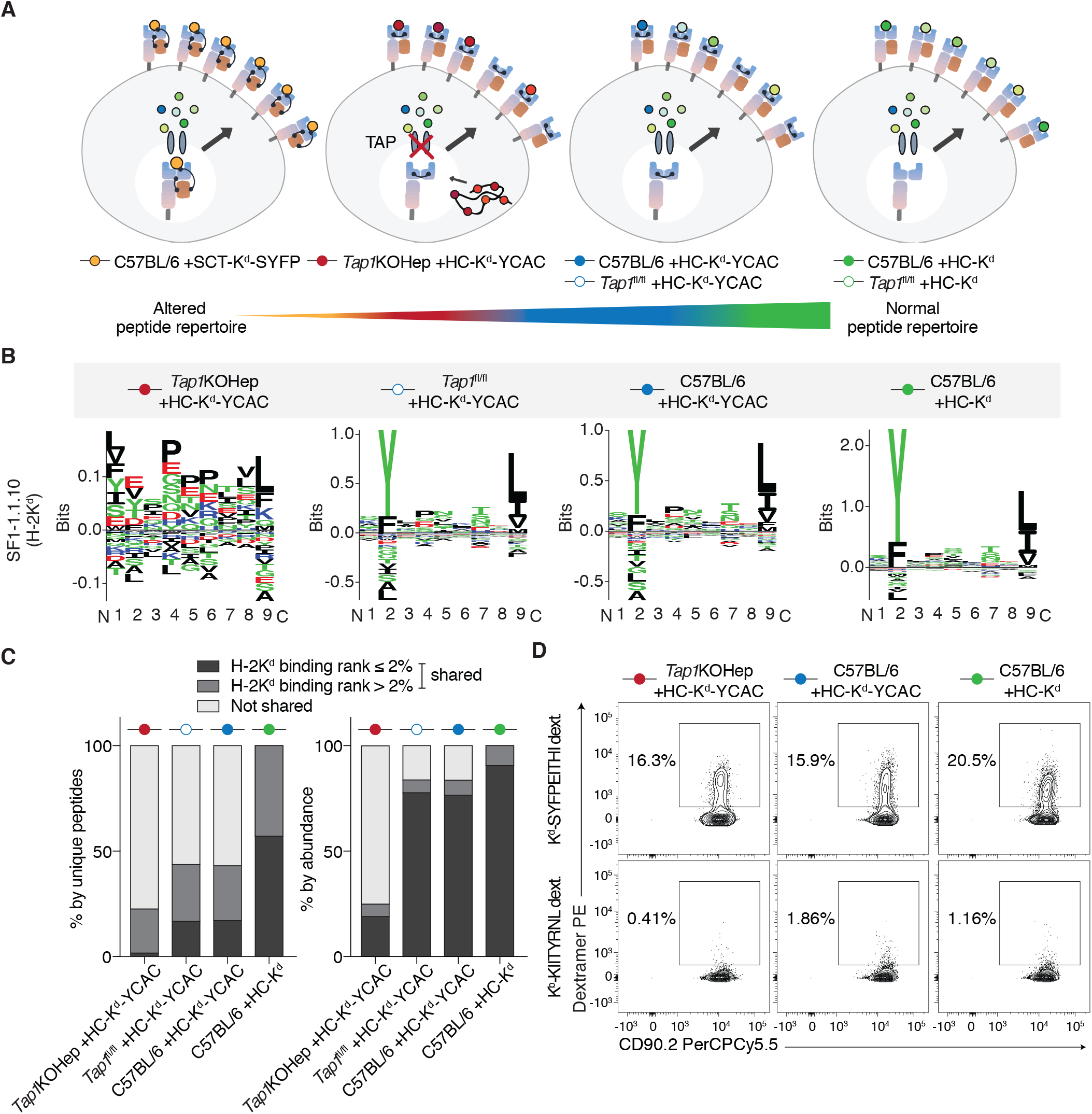
TAP-deficient hepatocytes transduced with AAV-K^d^-YCAC present an abnormal peptide repertoire. **(A)** Differences in peptide presentation between TAP-sufficient and deficient hepatocytes transduced with AAV vectors encoding one of three H-2K^d^ constructs. From left to right, diagrams show a single presented peptide (SYFPEITHI), a substantially-altered repertoire resulting from the combination of K^d^-YCAC expression and absence of TAP, a moderately altered repertoire when K^d^-YCAC is expressed in cells where TAP is active, and the repertoire normally associated with K^d^ in TAP-sufficient hepatocytes. Both empty and peptide-loaded K^d^-YCAC molecules are expected to be present at the cell surface. **(B)** H2-K^d^-binding motifs were generated from a non-redundant list of 9-mer peptides using GibbsCluster-2.0. Peptides eluted from B6/HC, B6/YCAC or *Tap1*^fl/fl^/YCAC, hepatocytes displayed the canonical K^d^ binding motif of tyrosine (Y) at position 2 and leucine (L) or isoleucine (I) at the C-terminus while no clear binding motif was discernible in the sequences for peptides eluted from *Tap1*KO/YCAC hepatocytes. **(C)** Just under 45% of peptides eluted from B6/YCAC or *Tap1*^fl/fl^/YCAC hepatocytes were common to the B6/HC repertoire, while only 22% of peptides from *Tap1*KO/YCAC were shared with B6/HC. Similarity across peptide repertoires was increased when the comparison was weighted for peptide abundance. The peptide SYFPEITHI was present in all repertoires. **(D)** Comparable proportions of activated CD8^+^ T cells able to recognise K^d^-SYFPEITHI were detected among the liver leukocytes of primed C57BL/6 mice inoculated with AAV-HC-K^d^ or AAV-HC-K^d^-YCAC and those of *Tap1*KOHep mice treated with AAV-HC-K^d^-YCAC. Representative flow plots from n = 3 are shown.

### Increasing perturbation of the hepatocyte H-2K^d^ peptide repertoire correlates with progressive reduction of K^d^-bearing skin graft survival in transduced mice

Mice were inoculated with AAV vectors encoding HC-K^d^, HC-K^d^-YCAC, or SCT-K^d^ with the peptide SYFPEITHI (SCT-K^d^-SYFP), seven days prior to transplantation with a B6.Kd skin graft (Figure 6A). The construct sequence and expression data for SCT-K^d^-SYFP are shown in Supplementary Figures 1 and 7, respectively. The majority of C57BL/6 or *Tap1*^fl/fl^ mice transduced with HC-K^d^ accepted K^d^-bearing B6.Kd skin grafts indefinitely, whereas median survival of B6.Kd skin grafts to C57BL/6 or *Tap1*^fl/fl^ mice treated with HC-K^d^-YCAC was reduced to 62.5±5.3 days and 52.5±9.2 days respectively (Figure 6B). Graft survival in *Tap1*KOHep mice inoculated with HC-K^d^-YCAC was further reduced to 20±1.9 days, similar to that in C57BL/6 mice which received a vector encoding SCT-K^d^-SYFP (17±1.2 days). Median graft survival in no vector control mice was 12.5±0.6 days for C57BL/6 and 10±0.6 days for *Tap1*KOHep (p<0.0001 for overall survival trend, Figure 6B). Failure to induce tolerance was not associated with loss of transgene expression in the transduced livers (Supplementary Figure 7). Instead, progressive loss of ability of H-2K^d^ gene transfer to prolong B6.Kd skin graft survival correlated with increasing disturbance of the hepatocyte K^d^ peptide repertoire in the various experimental groups. These findings, along with the inability of SCT-K^b^ vectors to induce tolerance in polyclonal alloreactive T cell populations demonstrate the importance of the hepatocyte immunopeptidome in transplantation tolerance induction following donor MHC I gene transfer to the liver.

**Figure 6.**
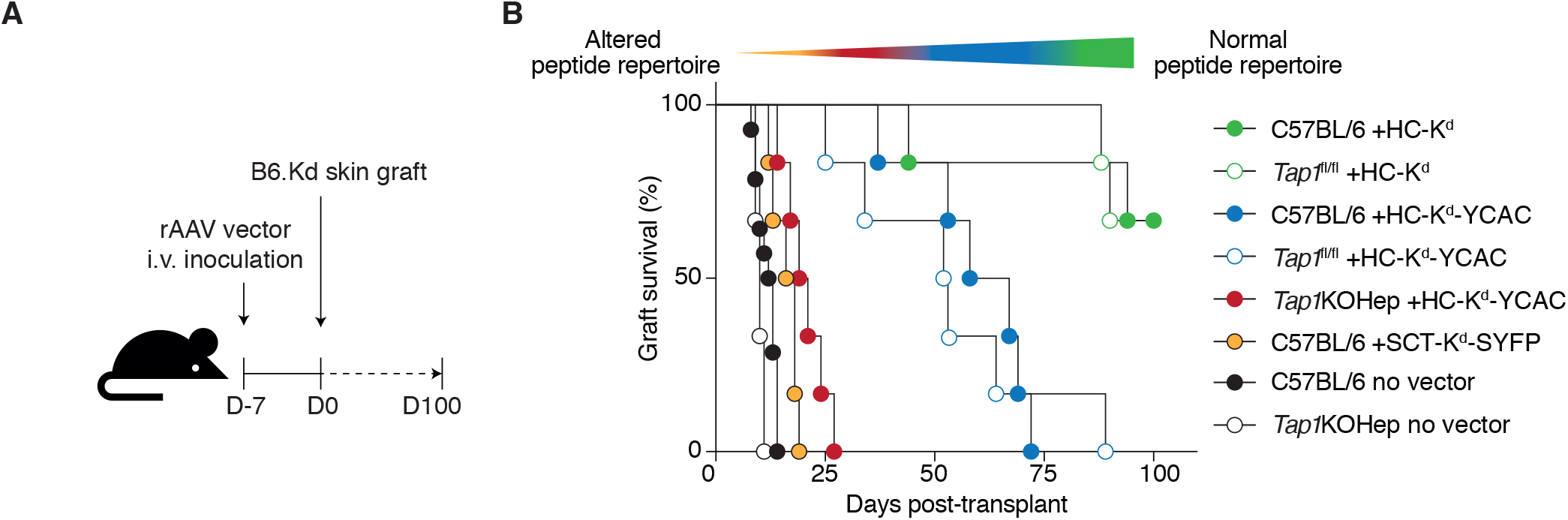
Alteration in the endogenous peptide repertoire impedes transplantation tolerance induction. **(A)** *Tap1*KOHep, *Tap1*^fl/fl^ or C57BL6 mice were inoculated with AAV-HC-K^d^-YCAC or were not transduced. Another group of C57BL/6 were transduced with AAV-SCT-K^d^-SYFP, while C57BL6 and *Tap1*^fl/fl^ controls received AAV-HC-K^d^. 7 days post-inoculation, the mice were challenged with B6.Kd skin grafts. **(B)** Expression of HC-K^d^ loaded with the endogenous peptide repertoire was able to induce tolerance to B6.Kd skin grafts in the majority of C57BL/6 and *Tap1*^fl/fl^ recipients. Increasing perturbation of the H-2K^d^-bound peptide repertoire progressively shortened graft survival of H-2K^d^-bearing skin grafts [C57BL/6 inoculated with AAV-HC-K^d^ (MST: indefinite), C57BL/6 inoculated with AAV-HC-K^d^-YCAC (MST: 62.5 days), *Tap1*^fl/fl^ inoculated with AAV-HC-K^d^-YCAC (MST: 52.5 days), *Tap1*KOHep inoculated with AAV-HC-K^d^-YCAC (MST: 20 days), Mantel-Cox log-rank test for trend p < 0.0001, all groups n = 6].

**Figure 7:**
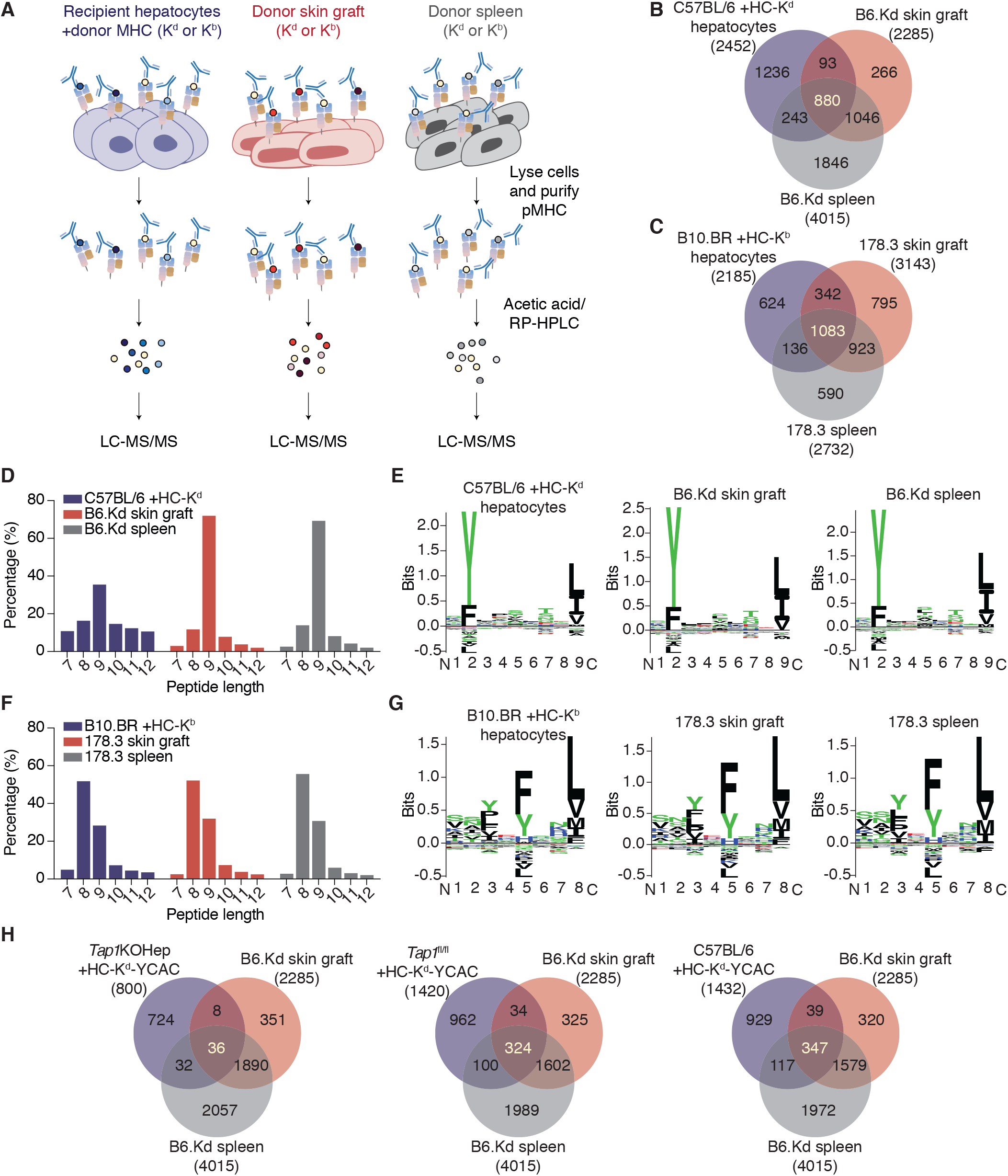
Profiling of the H-2K^d^ and H-2K^b^-associated peptide repertoires of transduced hepatocytes, skin and spleen. **(A)** A schematic diagram illustrating the immunoaffinity purification workflow. **(B,C)** Unique peptides were identified from transduced hepatocytes, skin and spleen. For H-2K^d^, the peptide repertoires of C57BL/6 hepatocytes transduced with AAV-HC-K^d^, B6.Kd spleen and B6.Kd skin grafts collected 7 days after transplantation were determined, while for H-2K^b^, the corresponding tissues were B10.BR hepatocytes transduced with AAV-HC-K^b^, 178.3 spleen and 178.3 skin grafts sampled at 7 days post-transplant. Data from two independent experiments are shown. Within each replicate experiment, samples from 3-4 mice were pooled per condition. For K^d^, 880 9-mer peptides were found to be shared across all three tissue types, while 1083 K^b^ peptides (8-11 mer, IC < 500nM) were common to the three tissues. **(D)** Length distribution of filtered H-2K^d^ peptides from hepatocytes, spleen and skin graft tissue samples. The number of peptides of each length identified with a 5% FDR cut-off are shown. **(E)** Peptide-binding motifs for H2-K^d^ peptides generated from a non-redundant list of 9-mer peptides using GibbsCluster-2.0. **(F)** Length distribution of H-2K^b^ peptides from hepatocytes, spleen and skin graft tissue samples (as for **D**). Most eluted peptides are 8-mers, with 9-mers also relatively frequent. **(G)** Peptide-binding motifs for H2-K^b^ peptides generated from a list of 8-11-mer peptides using the GibbsCluster-2.0 algorithm as above. The canonical binding motifs were observed for all three tissues for both H2-K^d^ and H2-K^b^. **(H)** The extent of peptide sharing between the H2-K^d^ repertoires of hepatocytes, spleen and skin was substantially reduced when *Tap1*KOHep, *Tap1*^fl/fl^ or C57BL6 mice inoculated with AAV-HC-K^d^-YCAC were substituted for C57BL/6 transduced with AAV-HC-K^d^. This reduction was particularly striking for the combination of *Tap1*KOHep with AAV-HC-K^d^-YCAC.

### Profiling the tissue-specific immunopeptidomes of hepatocytes, skin and spleen

Given the ability of H-2K^b^ or K^d^ HC expressed in recipient hepatocytes to induce tolerance to allogeneic skin grafts and to downmodulate responses against donor splenocyte stimulators(1, 3), we postulated that the peptides critical for allorecognition and tolerance induction would be found within a subset common to these three tissue types. The self-peptide repertoires of transduced hepatocytes, grafted donor skin and donor spleen were determined using a combination of immunoaffinity purification and RP-HPLC to liberate and collect peptide-containing fractions from MHC I with LC-MS/MS for peptide identification, for both the 178.3 to B10.BR (K^b^ mismatch, H-2^k^ background) and B6.Kd to C57BL/6 (K^d^ mismatch, H-2^b^ background) strain combinations, as outlined in Figure 7A. The lists of common peptides are found in Supplementary Table 2 (H-2K^d^) and Supplementary Table 3 (H-2K^b^). For H-2K^d^, 880 common peptides were identified across the three tissue types (Figure 7B), whereas there were 1083 common K^b^-binding peptides (Figure 7C). The peptide length distributions (Figure 7D and F) and binding motifs (Figure 7E and G) were as anticipated for the respective allomorphs and were similar across tissue types. Of note, the common peptide pool was more limited when TAP-sufficient, or particularly TAP-deficient hepatocytes had been transduced with HC-K^d^-YCAC (324, 347 and 36 unique peptides respectively), compared to TAP-sufficient hepatocytes transduced with HC-K^d^ (880 unique peptides) (Figure 7H). Comparison of the different tissue immunopeptidomes showed that in the two settings where skin graft tolerance was achieved in wild-type recipient mice following expression of allogeneic donor MHC I in hepatocytes, the proportion of skin peptides common to hepatocytes was 43% and 45% for H-2K^d^ (Figure 7B) and H-2K^b^ (Figure 7C), respectively. Conversely, only 1.6% of skin peptides were also found in *Tap1*KOHep hepatocytes transduced with HC-K^d^-YCAC, while in the two groups with intermediate graft survival (*Tap1*^fl/fl^ or C57BL/6 inoculated with HC-K^d^-YCAC), the proportion of skin peptides present in hepatocytes was in the order of 15-17%. Gene ontology analysis of source proteins is shown in Supplementary Figure 6.

### H-2K^b^ peptides from the common peptide pool are recognised by activated alloreactive CD8^+^ T cells

A total of 100 peptides were selected for screening (listed in Supplementary Table 4). 96 peptides were drawn from the common peptide pool, and their identity was confirmed by a direct comparison between the mass spectra obtained from synthetic and eluted natural peptides (Supplementary Figure 8). A further four peptides had been previously identified as alloreactive CD8^+^ T cell epitopes in B10.BR mice(11, 25). Three of these four epitopes were detected within the common pool. Binding of pMHC tetramers was used to determine which peptides combined with H-2K^b^ to form immunogenic epitopes recognised by alloreactive B10.BR CD8^+^ T cells. B10.BR mice were first primed by placement of a K^b^-bearing 178.3 skin graft. Approximately 30 days after graft rejection, mice were inoculated with HC-K^b^, and after a further 7 days, liver leukocytes were isolated and stained by flow cytometry (Figure 8A). The gating strategy is shown in Figure 8B. Activated CD8^+^ T cells were defined as CD44^+^PD-1^hi^, whereas PD-1^-^ cells were considered to be an internal control population, which had been exposed in vivo to H-2K^b^ expressed on hepatocytes but were not activated. Peptides were deemed immunogenic when *≥*2% of CD44^+^PD-1^hi^ CD8^+^ T cells were bound by pMHC tetramer. Representative flow plots demonstrating T cell recognition of immunogenic and non-immunogenic peptides are shown in Figure 8C, while data summarising the results are shown in Figure 9A and Supplementary Table 4. Allorecognition of K^b^-bound peptides was then examined in recipient mice of a second background haplotype (BALB/c, H-2^d^) (Figures 9A-B and Supplementary Table 4).

**Figure 8.**
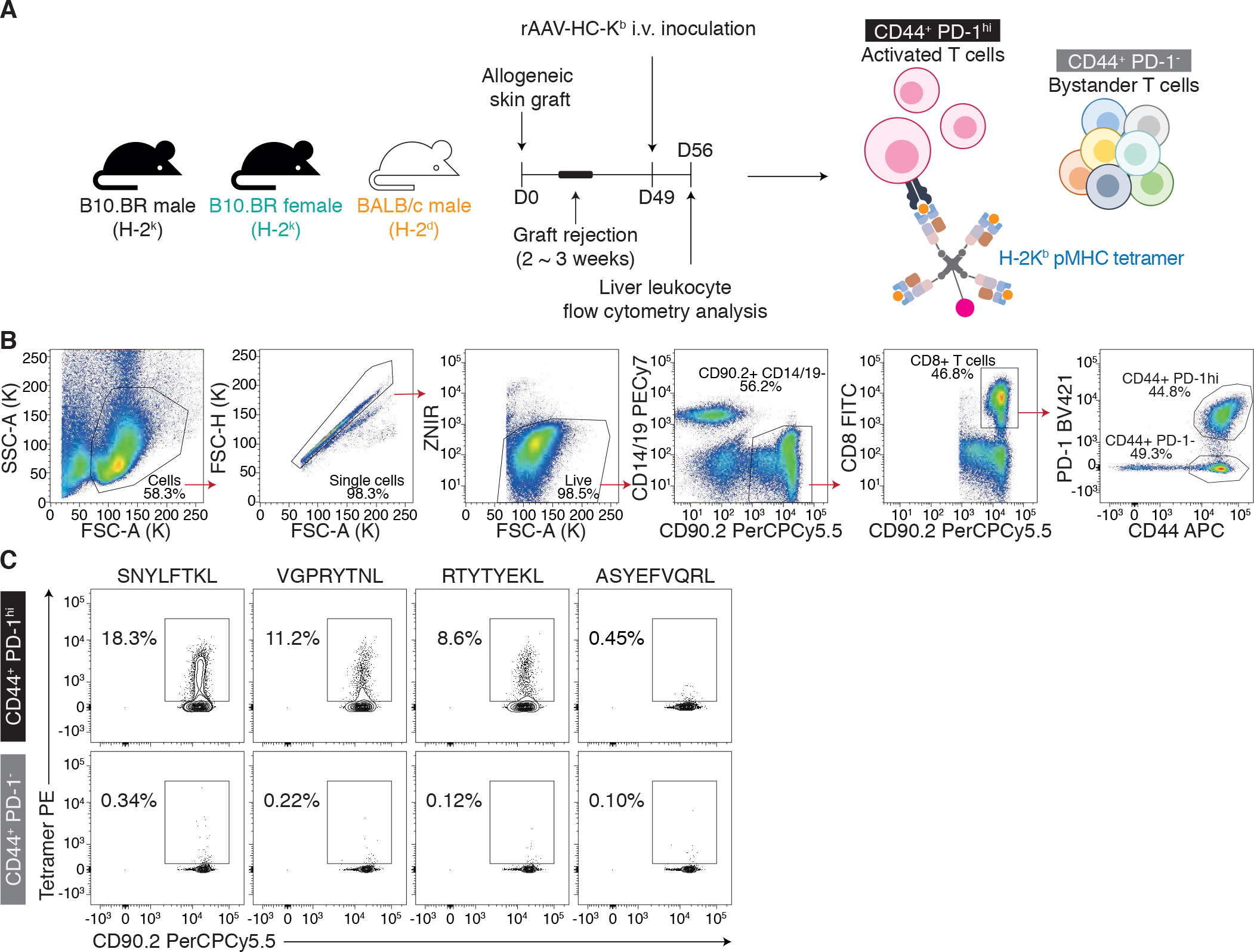
Identification of pMHC epitopes recognised by alloreactive CD8^+^ T cells. **(A)** Male B10.BR mice were primed with male 178.3 skin grafts and 30 days after rejection, mice were transduced with AAV-HC-K^b^. Liver leukocytes were isolated and pMHC tetramer binding was analysed using flow cytometry. **(B)** Shows the gating strategy that was employed to identify and stratify CD8^+^ T cells into two groups; CD44^+^PD-1^hi^ and CD44^+^PD-1^-^. CD44^+^PD-1^hi^ cells are activated alloreactive CD8^+^ T cells and CD44^+^PD-1^-^ cells are non-activated bystander CD8^+^ T cells which serve as internal controls. **(C)** Examples of pMHC tetramer binding by CD44^+^PD-1^hi^ and CD44^+^PD-1^-^ CD8^+^ T cells. Representative flow plots (from n = 3 biological replicates) for the selected pMHC epitopes are shown. K^b^-SNYLFTKL (Epas_387-394_), K^b^-VGPRYTNL (Mapk1_19-26_) and K^b^-RTYTYEKL (Ctnnb1_329-336_) were recognised by a large proportion of activated alloreactive T cells, whereas few T cells bound K^b^-ASYEFVQRL (Dync1h1_1379-1387_).

**Figure 9.**
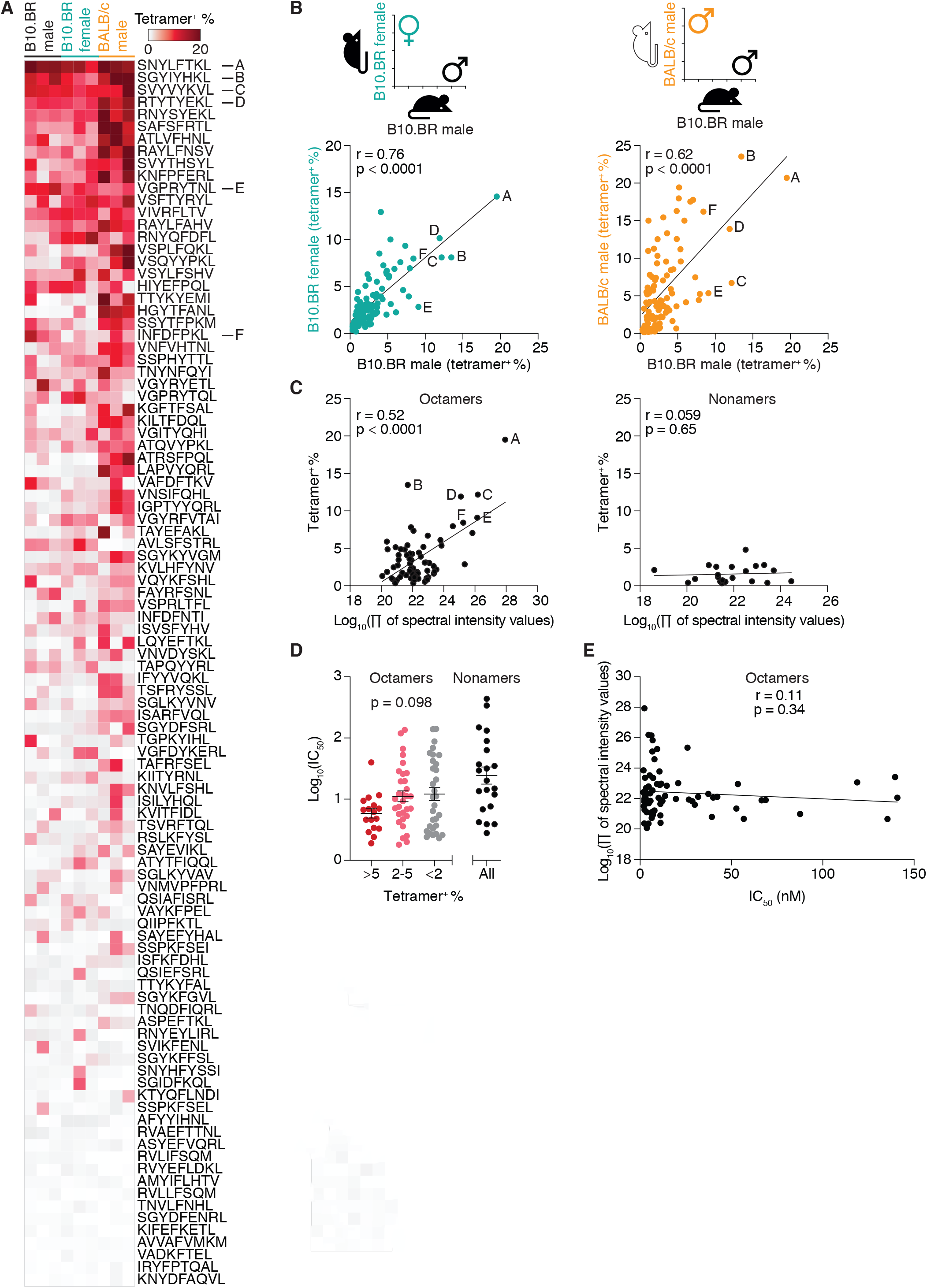
Features of immunogenic pMHC recognised by mice from different allogeneic backgrounds. **(A)** 100 peptides were screened for recognition by activated alloreactive T cells. T cells from female B10.BR mice primed with a female 178.3 skin graft and male BALB/c mice primed with a male C57BL/6 skin graft were used in parallel with T cells from primed male B10.BR. A heatmap was generated in order to compare recognition of pMHC epitopes across different sexes and strains. Epitopes are ordered from the top to the bottom by the average of pMHC tetramer binding across all samples. 13 pMHC were recognised by >5% of activated alloreactive T cells across all three groups. Full peptide screening data can be found in Supplementary Table 4. **(B)** Strong correlation of T cell binding to each pMHC was found between male and female B10.BR [Pearson correlation coefficient (r) = 0.76, p < 0.0001] and between male B10.BR and male BALB/c (r = 0.62, p < 0.0001). **(C)** For male B10.BR mice, the proportion of T cells binding 8-mer pMHC tetramers was proportional to log_10_ of the product of the spectral intensity values for each peptide across three different tissues (r = 0.52, p < 0.0001). Conversely, binding to 9-mer pMHC tetramers did not correlate with peptide spectral intensity (r = 0.059, p = 0.65). For each pMHC epitope, data was obtained from 1-2 independent experiments with a total of 3 biological replicates. **(D)** Predicted peptide binding affinity for H-2K^b^ (measured by IC_50_ ) did not differ significantly between strongly, moderately or non-immunogenic peptides (p = 0.098 by one-way ANOVA, mean ± SEM are shown), **(E)** nor was a correlation observed between IC_50_ and spectral intensity values (r = 0.11, p = 0.34).

Of 100 peptides screened, 17 peptides were recognised by >5% of activated recipient CD8^+^ T cells from male B10.BR mice (termed strongly immunogenic), and a further 39 were bound by 2-5% of cells (moderately immunogenic). These responses were mirrored in female B10.BR recipients (Figure 9A-B). A number of pMHC epitopes were recognised by BALB/c mice as well as B10.BR (Figure 9A-B). All peptides recognised by >5% of B10.BR responder cells and 42/43 of those binding >5% of BALB/c cells were 8-mers. For 8-mer peptides, there was a strong correlation between overall peptide abundance (as estimated by the product of the spectral intensity across the three tissue types) and the percentage of T cells with specificity for a given pMHC (r=0.52, p<0.0001, Figure 9C). No such relationship was observed for 9-mers. Predicted peptide binding affinity for H-2K^b^ (measured by IC_50_) did not differ significantly between strongly, moderately or non-immunogenic peptides (Figure 9D, p=0.098 by one-way ANOVA), nor was a correlation observed between IC_50_ and spectral abundance (Figure 9E). Simultaneous staining with two different pMHC tetramers was used to evaluate the proportion of T cells recognising more than one pMHC specificity (Figure 10A), with a total of six peptides being evaluated. A substantial proportion of T cells recognised two peptides (SGYIYHKL and/or SVYVYKVL) in addition to SNYLFTKL. 86.7±19.2% of T cells recognising SGYIYHKL-PE could recognise SNYLFTKL-APC and 66.8±9.0% of T cells recognising SVYVYKVL-PE could also recognise SNYLFKTKL-APC, whereas cross-reactivity between VGPRYTNL, INFDFPKL and RTYTYEKL was considerably lower (Figure 10B-C) When 5 of these 6 peptides (excluding SGYIYHKL), each binding between 7.2 and 15.2% of T cells, were used together as a panel, the proportion of alloreactive CD8^+^ T cells bound increased to 39.1±3.4% (Figure 10D-E, p<0.0001 compared with SNYLFTKL). This cumulative increase in binding is consistent with alloreactive T cell recognition of epitopes comprising both a self-peptide and allogeneic MHC I molecule.

**Figure 10.**
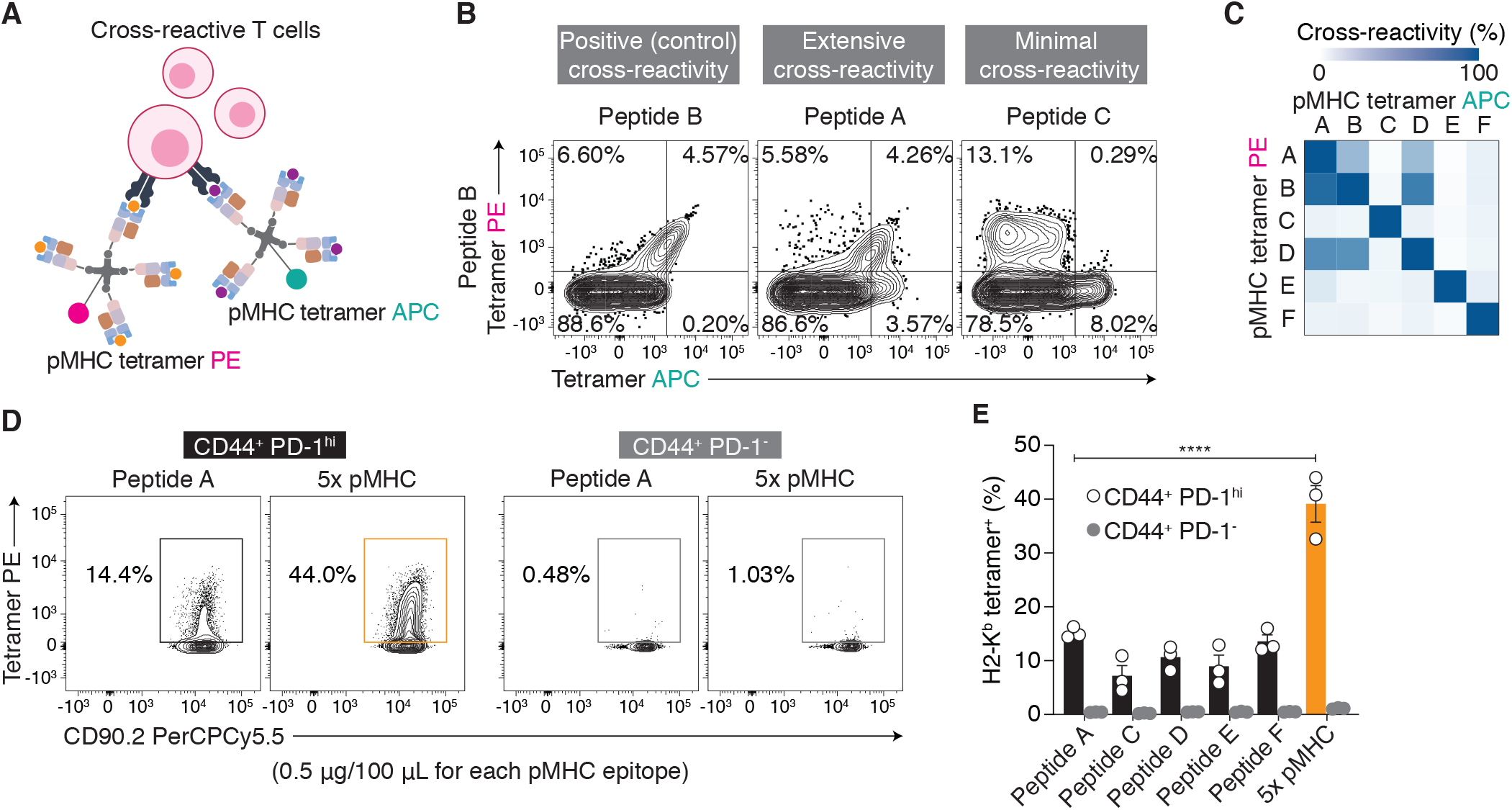
pMHC multimers can be combined to enhance detection of alloreactive T cells within a polyclonal population. **(A,B)** Staining with two different pMHC tetramers was used to evaluate the proportion of T cells recognising more than one pMHC specificity. Six strongly immunogenic peptides were tested. 86.7 ± 19.2 % of SNYLFTKL^+^ T cells recognised SGYIYHKL in addition to SNYLFTKL, while SVYVYKVL tetramers bound 66.8 ± 9.0 % of SNYLFTKL^+^ T cells and SGYIYHKL tetramers bound 75.8 ± 18.5 % of SVYVYKVL^+^ cells. Cross-reactivity between VGPRYTNL, INFDFPKL and RTYTYEKL was considerably lower (4.0 ± 1.1 and 8.4 ± 0.6 % of T cells recognised VGPRYTNL in addition to INFDFPKL and RTYTYEKL, respectively). **(B)** Representative flow plots from two independent experiments with 3-4 mice pooled/-experiment. **(C)** A heatmap was generated in order to compare cross-reactivity between different pMHC epitopes. **(D,E)** When 5 of these 6 peptides (excluding SGYIYHKL), each binding between 7.2 and 15.2% of T cells were used together as a panel, the proportion of alloreactive CD8^+^ T cells bound increased to 39.1 ± 3.4 % (p < 0.0001) compared with 15.2 ± 0.6 % for SNYLFTKL. Data were obtained from one experiment with n = 3 biological replicates/group. **(D)** Representative flow plots (from n = 3). **(E)** Data are shown as mean ± SEM, statistical analysis comprised one-way ANOVA in conjunction with Sidak’s multiple comparison test: **** p < 0.0001.

### A pMHC tetramer panel enables mechanistic insights into transplant immune responses

We used the 5-tetramer panel (above) to enumerate and phenotype alloreactive CD8^+^ T cells in a model of secondary skin graft tolerance or rejection (Figure 11A). Gating strategy is shown in Supplementary Figure 9A. Secondary skin grafts to control mice were promptly rejected, while grafts performed 7 days after inoculation of primed recipients with AAV-K^b^ survived indefinitely (Figure 11B). The number of tetramer-positive (tet^+^) CD8^+^ T cells in combined secondary lymphoid organs expanded three-fold following rejection of a secondary skin graft (12,000±1,500 versus 31,000±3,500 cells, p=0.0021), but had not increased significantly at a matching interval during graft acceptance (12,000±1,500 against 22,000±3,700, p=0.29) (Figure 11C). Induction of tolerance in primed mice by inoculation with AAV-K^b^ resulted in a sharp increase of tet^+^ cells within the liver (from 480±97 to 131,000±22,000 cells, p<0.0001), declining subsequently (Figure 11D). While the overall number of tet^+^ cells on protocol d14 was not significantly greater in rejecting skin grafts than in grafts destined to be accepted (640±190 versus 400±130 cells, p=0.78) (Figure 11E), rejecting transplants contained a tetramer-bright population (tet^hi^, 12.7±0.97% of CD8^+^ T cells) which was not detected in tolerated grafts (Figure 11 F-H). Few tet^+^ cells persisted long-term in accepted grafts (Figure 11E). Examining the tet^+^ cells revealed phenotypic changes which were partly obscured within the bulk CD8^+^ population. In naïve mice, the tet^+^ liver leukocytes included all CD8^+^ T cell subsets, whereas in mice having rejected a primary or secondary graft, tet^+^ cells were exclusively antigen-experienced, comprising both central memory and effector/resident memory-like cells. Following exposure to H-2K^b^ in the liver, tet^+^ cells with a central memory phenotype were no longer observed, and the majority of cells expressed markers of liver-homing or residency (CD69 and/or CXCR6, Figure 12A-B and Supplementary Figure 10). Seven days after inoculation with AAV-K^b^, tet^+^ cells in the liver have upregulated expression of PD-1, TIGIT, Tim-3 and LAG-3. While strong PD-1 expression persists through d84 after tolerance induction, expression of LAG-3, Tim-3 and to a lesser extent TIGIT in the tet^+^ population progressively declines in tandem with the decay of tet^+^ cell numbers, consistent with preferential deletion of cells expressing multiple exhaustion markers (Figure 12C and Supplementary Figure 11). All tet^+^ cells isolated from rejecting or tolerated grafts on protocol d14 were antigen experienced. Tet^hi^ cells from rejecting grafts were uniformly CD62L-negative and PD-1^hi^, likely denoting recent strong activation of these cells, while the majority of tet^+^ cells from tolerated grafts expressed CD62L and not PD-1.

**Figure 11.**
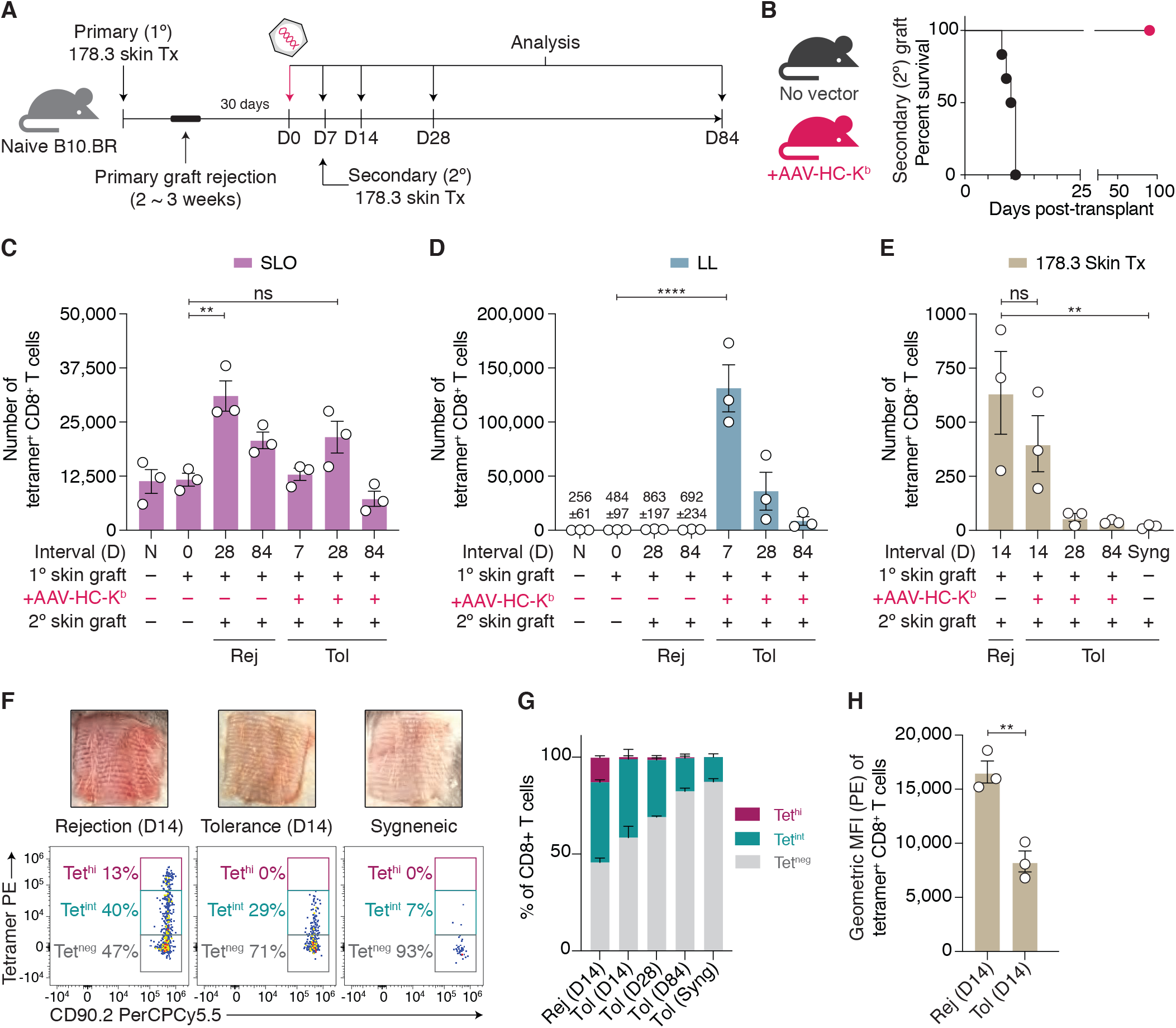
Tracking alloreactive T cells using a tetramer panel. **(A)** A 5-tetramer panel was used to enumerate and phenotype alloreactive CD8^+^ T cells in a model of secondary skin graft tolerance or rejection. **(B)** Secondary skin grafts to control mice were promptly rejected, while grafts performed 7 days after inoculation of primed recipients with AAV-K^b^ survived indefinitely (n = 6). **(C)** The number of tet^+^ cells in pooled SLO expanded three-fold following rejection of a secondary graft (12,000 ± 1,500 versus 31,000 ± 3,500 cells, p = 0.0021), but had not increased significantly after graft acceptance (12,000 ± 1,500 against 22,000 ± 3,700, p = 0.29). **(D)** Induction of tolerance in primed mice by inoculation with AAV-K^b^ resulted in a sharp increase of tet^+^ cells within the liver (from 480 ± 97 to 131,000 ± 22,000 cells, p < 0.0001), declining subsequently. **(E)** Numbers of tet^+^ cells in rejecting transplants were similar to those in tolerated grafts on d14 (640 ± 190 versus 400 ± 130 cells, p = 0.78), while few tetramer-positive cells persisted long-term in accepted grafts. **(F)** Rejecting skin grafts contained a population of CD8^+^ T cells which stained very strongly with the tetramer panel (representative flow plots from n = 3). **(F-H)** This tetramer-bright population was not detected in tolerated grafts. **(C,D,E,G,H)** Data are shown as mean ± SEM, n = 3/group, **(C,D,E)** one-way ANOVA in conjunction with Sidak’s multiple comparison test, **(H)** Student’s t-test: ns, not significant, ** p < 0.01, **** p < 0.0001.

**Figure 12.**
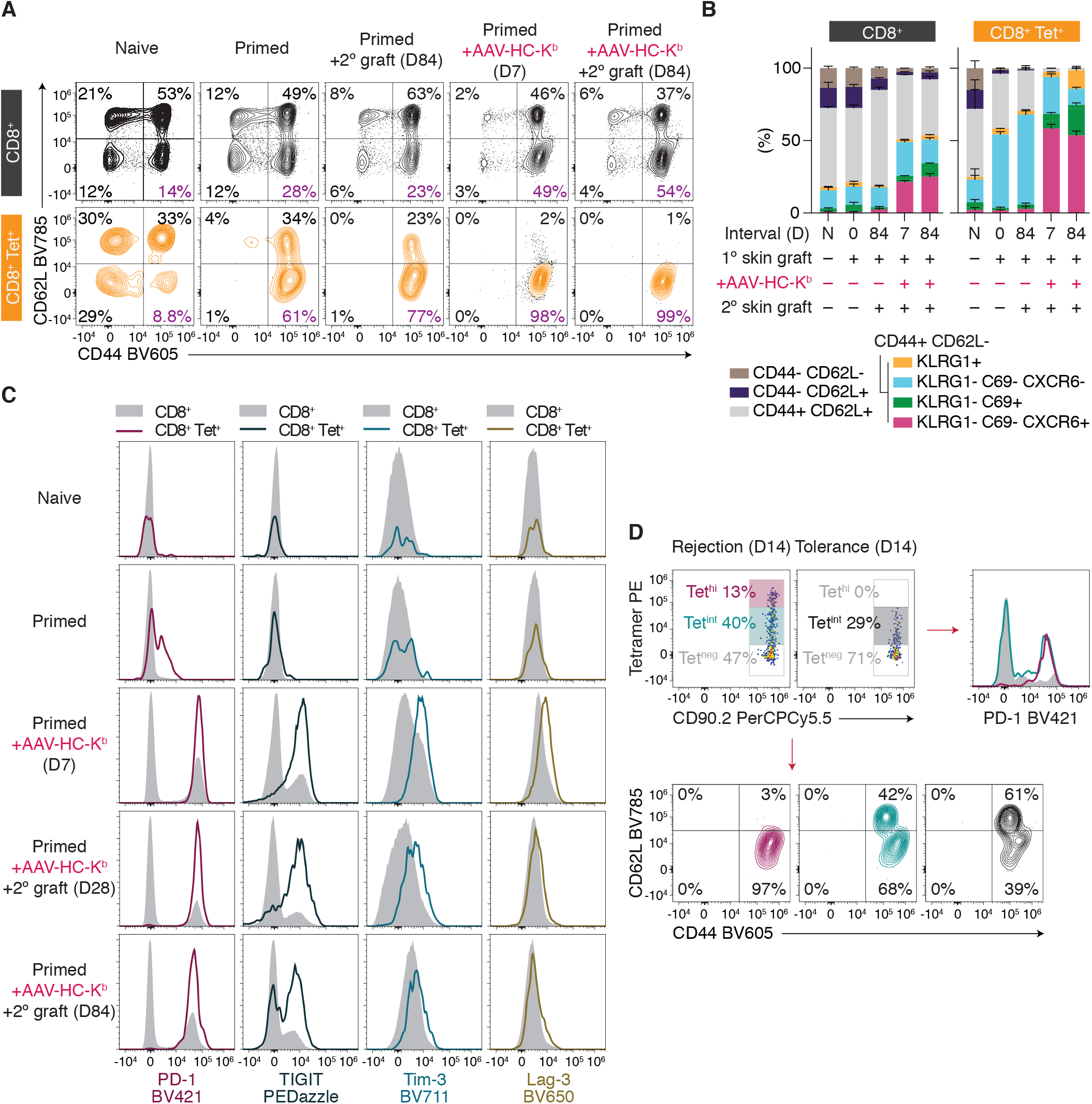
Tetramer-positive alloreactive cells are phenotypically distinct from bulk CD8^+^ T cells. Surface markers of naïve, effector and memory status were assessed in bulk and tet^+^ CD8^+^ T cells from naïve or transplanted mice. Relatively subtle phenotypic changes in the CD8^+^ T cell population were far more obvious when tet^+^ cells were examined. **(A)** In naïve mice, tet^+^ liver leukocytes contained all CD8^+^ T cell subsets, whereas in mice having rejected a primary or secondary graft, tet^+^ cells were exclusively antigen-experienced, comprising both central memory and effector/resident memory cells. **(B)** Following exposure to H-2K^b^ in the liver, CD62L^+^ tet^+^ cells were no longer observed, and the majority of cells expressed markers of liver-homing or residency (CD69 and/or CXCR6). **(C)** Seven days after inoculation with AAV-HC-K^b^, tet^+^ cells in the liver have upregulated PD-1, TIGIT, Tim-3 and LAG-3. While strong PD-1 expression persists through d84 after tolerance induction, expression of LAG-3, Tim-3 and to a lesser extent TIGIT in the tet^+^ population declines over this time. **(D)** All tet^+^ cells isolated from rejecting or tolerated grafts on protocol d14 were antigen-experienced. Tet^hi^ cells, only present in rejecting grafts, were uniformly CD62L-negative and PD-1^hi^. **(A,C,D)** Representative flow plots (from n = 3). **(B)** Data are shown as mean ± SEM (n = 3).

## Discussion

Direct allorecognition refers to the engagement of recipient TCRs by intact allogeneic MHC molecules on the surface of donor cells. In some settings, intact, unprocessed donor MHC is transferred to the surface of recipient APC, a variant known as the semi-direct pathway of allorecognition(26). While direct/semi-direct allorecognition is a critical driver of both solid organ allograft rejection(26) and tolerance induction(3), the specific molecular targets of most directly-alloreactive CD8^+^ T cells have not been defined, limiting our understanding of these phenomena. In this study, we provide definitive evidence for the contribution of the peptide cargo of allogeneic MHC I molecules to transplant tolerance induction. Moreover, we demonstrate the feasibility of a systematic approach incorporating mass spectrometry-based identification of alloantigen-presented peptides, alloreactive T cell enrichment and pMHC multimer screening for the discovery of pMHC epitopes that are directly recognised by alloreactive CD8^+^ T cells. To our knowledge, this represents the first example of such an approach in the context of direct allorecognition.

Studies over many years using T cell clones suggested that most directly alloreactive CD8^+^ clones recognise allogeneic MHC I molecules complexed with an endogenous peptide, while peptide-independent direct allorecognition was also reported. Our recent finding that direct recognition of hepatocyte-expressed donor MHC I by recipient CD8^+^ T cells was required for donor skin graft tolerance induction provided us with a unique opportunity to examine the role of the hepatocyte immunopeptidome in transplant tolerance, and by extension, in alloresponses more broadly. We used two complementary approaches to separate expression of donor MHC I molecules from presentation of their usual peptide cargo. In the first approach, we utilised an AAV vector encoding H-2K^b^ HC, β2m and the single peptide KIITYRNL to express high levels of K^b^ on the surface of recipient hepatocytes, while excluding presentation of naturally-processed endogenous peptides. Tolerance to K^b^-bearing skin grafts was able to be induced in reconstituted mice with a monoclonal population of T cells all recognising the K^b^-KIIT epitope, but not in wild-type mice possessing a full TCR repertoire. Secondly, we expressed a mutated version of the H-2K^d^ HC, which is permissive to loading with suboptimal peptides in a purpose-bred mouse strain (*Tap1*KOHep) with a hepatocyte-specific deletion of *Tap1*. This combination resulted in a significant alteration of the K^d^-bound peptide repertoire and prevented tolerance induction. Graft survival was shortened to a lesser extent in TAP1-sufficient mice transduced with HC-K^d^-YCAC, in keeping with the lesser perturbation of the K^d^ peptide repertoire observed in these mice. The point mutations Y84C and A139C in the K^d^ HC permit the creation of a disulphide bridge which stabilises the peptide binding groove when empty or populated by a low-affinity peptide(18). The presence of these mutations alters the composition of the K^d^-associated peptide repertoire but does not impede T cell responses to a given pMHC ligand(24). Accordingly, the ubiquitous peptide SYFPEITHI was present within the repertoires of all hepatocytes transduced with either HC-K^d^ or HC-K^d^-YCAC and comparable proportions of T cells isolated from the livers of these mice bound K^d^-SYFP dextramers.

Having demonstrated the importance of the peptide cargo of allogeneic MHC I molecules for transplant tolerance induction, we set about identifying the subset of peptides responsible for modulating the anti-graft alloresponse, using an immunoproteomic approach to identify and characterise the immunopeptidomes of relevant tissues. We hypothesised that hepatocyte expression of peptides that were also presented by cells in the grafted skin underpinned tolerance induction. Consistent with this hypothesis, in conditions where tolerance was achieved, 43% of K^d^-associated peptides and 45% of K^b^-bound peptides from transduced hepatocytes were also expressed in donor skin. When survival prolongation was minimal, only 1.6% of hepatocyte K^d^-bound peptides were common to donor skin, and this shared proportion increased to 15-17% in the groups with intermediate graft survival. Recognition of pMHC epitopes shared between hepatocytes and the graft could induce tolerance via deletion, sequestration within the liver or cell-intrinsic functional inactivation of specific CD8^+^ T cells, preventing them from attacking the skin. These mechanisms may operate in concert, while linked suppression of responses against epitopes expressed exclusively in the skin by T cells recognising shared epitopes may also contribute to the induction and maintenance of tolerance.

To identify immunogenic peptides from the shared subset, we generated an enriched population of alloreactive CD8^+^ T cells by priming mice with a skin graft expressing H-2K^b^, followed by boosting with a liver-specific AAV vector encoding HC-K^b^. Co-expression of CD44 with high levels of PD-1 allowed us to distinguish between activated cells and bystander cells which had been exposed to the alloantigens, but had not responded. Of the 100 K^b^-binding peptides selected for tetramer screening, 17 were recognised by >5% of the enriched alloreactive T cell population. INFDFNTI, previously reported to be recognised by a large proportion of the polyclonal alloreactive T cell population of B10.BR mice(25) was bound by an average of 3.7% of male B10.BR cells. To examine factors which might contribute to the immunogenicity of different pMHC, we used spectral intensity as an approximation for relative peptide abundance and IC_50_ to predict binding affinity of the peptide for H-2K^b^. An overall measure of abundance was derived by multiplying the spectral intensity recorded in each of the three tissues. For 8-mer peptides, a strong correlation was found between abundance and the proportion of CD8^+^ T cells recognising particular peptides. This is at variance with observations of CD8^+^ T cell recognition of K^b^-associated viral epitopes, where the frequency of T cells responding to a given epitope corresponds weakly at best to epitope abundance(27, 28). In contrast, while there was a tendency for the strongly immunogenic peptides to have high predicted affinity for K^b^, there was no significant difference in IC_50_ between the strongly-, moderately- and non-immunogenic groups, nor was there an association between IC_50_ and peptide abundance. While a prediction of immunogenicity based on peptide abundance is no substitute for direct measurement, these findings suggest that the likelihood of finding strongly immunogenic peptides is greater if the search is targeted towards the most highly-represented peptide species. The relationship between abundance and immunogenicity did not apply to 9-mer peptides.

One intriguing finding of this study was the broad overlap between strongly immunogenic K^b^-peptide epitopes recognised by BALB/c and those first identified in B10.BR mice (13/17 peptides). Further studies in mouse and human subjects will be needed to establish the prevalence of such “super-epitopes” recognised by allogeneic T cells across several genetic backgrounds. While extensive cross-reactivity is a feature of self MHC-restricted T cells, the range of possible cross-reactive epitopes is constrained by peptide length, with T cell clonotypes showing a strong preference for a single length peptide(29). Alloreactive T cell clones have undergone positive selection during thymic development based on self-MHC. The majority of peptides eluted from H-2K^k^ are 8-mers [(30) and Supplementary Data 1] which aligns with the result that in male B10.BR mice (H-2^k^) 17/79 8-mer peptides but 0/21 9-mers were strongly immunogenic. Most peptides associated with the H-2^d^ allomorphs D^d^, L^d^ and K^d^ are nonamers(30), yet 42/79 8-mers and only 1/21 9-mers were strongly immunogenic in BALB/c (H-2^d^) mice. Possible explanations for this asymmetry are that cross-reactivity involving an allogeneic MHC molecule is not subject to the same peptide length limitations as cross-reactivity between self MHC-restricted peptide epitopes, or that a small number of octamer-preferring CD8^+^ T cells from BALB/c mice expand preferentially in the presence of H-2K^b^ presenting mainly 8-mers. While the mean H-2K^b^ IC_50_ among the 9-mer peptides was greater than that of the 8-mers, a number of 9-mer peptides were in the same IC_50_ range as the strongly immunogenic 8-mers, so this is unlikely to fully explain the reduced frequency of recognition of 9-mer peptides.

A staining panel comprising only five strongly immunogenic pMHC epitopes identified nearly 40% of T cells in an enriched alloreactive population; we then used this panel to enumerate and phenotype alloreactive T cells in our model of secondary skin graft rejection or tolerance, thereby seeking insights into the mechanisms promoting tolerance. Tolerance induction upon AAV-K^b^ inoculation is accompanied by a rapid increase of tet^+^ T cells within the liver, decaying over subsequent weeks, similar to the kinetics of tolerance induction following administration of recombinant adenovirus expressing *β*-galactosidase(31). As tet^+^ T cell numbers decline, the proportion of the population expressing Tim-3 and LAG-3 decreases, while PD-1 and TIGIT expression persists. Cells expressing multiple co-inhibitory receptors may be preferentially deleted, while the remaining cells are functionally silenced(1, 31). While the number of tet^+^ cells was not significantly greater in rejecting skin grafts than in grafts destined to be accepted, a tetramer-bright population of CD8^+^ T cells was exclusively present in the rejecting grafts. Absence of these tet^hi^ cells from the tolerated grafts could reflect their deletion or sequestration in the liver following antigen encounter, or a difference in the avidity profiles of clones expanded after activation in different sites. Of note, reduced functional avidity of polyclonal alloreactive T cells was observed in mice accepting heart grafts after donor cell infusion and costimulatory blockade(32). There was minimal persistence of alloreactive CD8^+^ T cells within permanently accepted skin grafts. Tracking the evolution of the alloreactive TCR repertoire during rejection, emerging and established tolerance would provide definitive evidence for the expansion and/or deletion of particular clonotypes, and could be coupled with gene expression analysis and avidity measurements. Future experiments could address the contribution of linked suppression to the induction or maintenance of tolerance. Whether or not the performance of a tetramer-panel developed using a skin-graft model will be replicated in solid organ transplants or GvHD remains to be determined.

The findings of this study open a number of avenues for future research. Firstly, they suggest an approach for the systematic discovery of pMHC epitopes for directly alloreactive CD8^+^ T cells which may be adapted to additional MHC I allomorphs in mice, humans and other species. Identification of large numbers of allogeneic pMHC epitopes will enable the characterisation of alloreactive TCR repertoires, including biophysical and structural studies with many more receptor-ligand pairs than the handful currently known, which in turn will provide further insights into the fundamental basis of alloreactivity. Systematic allogeneic pMHC epitope discovery is complemented by the recent advent of methodologies such as oligonucleotide barcoding permitting the use of pMHC multimers for parallel tracking and enumeration of >1000 individual pMHC specificities, with options for coupling to single cell TCR sequence and gene expression analyses(33, 34). pMHC multimers conjugated with radionuclides are sensitive tools which could be applied to in vivo imaging of specific alloresponses(35). Beyond their potential as research reagents, allospecific pMHC multimers could enhance post-transplant immune monitoring and immunophenotyping of directly alloreactive T cells in a clinical setting. Finally, pMHC multimers are not only reagents for antigen-specific T cell detection, but potential vehicles for antigen-specific therapy, which may be employed for the depletion or immunomodulation of alloreactive T cells(36). In conclusion, the findings of this study represent a significant advance in our understanding of the role of endogenous peptides in direct T cell alloreactivity, enabling exploration of the alloreactive T cell repertoire and potential translation to clinically-applicable tools.

## Methods

### Peptides, antibodies and reagents

Peptides were synthesised with an average of 98% purity (GL Biochem Shanghai Ltd.). Lyophilised peptides were reconstituted in 10% DMSO and stored at -80°C. Dasatinib (Sigma-Aldrich, catalogue# CDS023389) was reconstituted in DMSO and 5mM stock was stored at -80°C. Primary and secondary antibodies used in this study are shown in Supplementary Table 5.

### Cell Lines

The *Tap2*-deficient T lymphoma cell line RMA-S (provided by Professor Klas Kärre, Karolinska Institutet) was cultured in RPMI-1640 medium supplemented with L-glutamine (Lonza, catalogue# 12-702F), penicillin-streptomycin (Invitrogen, catalogue# 15140) and 10% FCS (Sigma-Aldrich, catalogue# 13K179) at 37°C with 5% CO_2_.

### RMA-S peptide stabilisation and transfection

RMA-S cells were grown to confluence and passaged to a concentration of 3×10^5^ cells/mL then transferred to flat-bottomed 24 well culture plates. The cells were incubated at 27°C for 20 hours with 5% CO_2_, then pulsed with peptides at concentrations ranging from 0.0001–10μM. Following 1 hour incubation at 27°C, cells were returned to 37°C for 2 hours. Surface expression of stabilised H-2K^b^ was quantified using flow cytometry with the conformation-dependent mAb Y-3. For transient transfection of RMA-S cells, 2×10^6^ cells were nucleofected with 2μg of pcDNA3.1^+^ plasmid per reaction (Lonza-AMAXA program X-001, Nucleofector 2b). 24 hours after transfection, transgene expression was evaluated using flow cytometry.

### Mice

Information about all mouse strains used in this study can be found in the supplementary methods.

### AAV vectors

H-2K^b^ and H-2K^d^ were cloned as previously described(3). SCT constructs consist of a defined peptide sequence, β2m and MHC I HC joined together by flexible linkers(16). A tyrosine to cysteine substitution at HC position 84 and a cysteine at the second position of the peptide-β_2_m linker form a disulphide trap(17). SCT (H-2K^b^) constructs with peptide sequences KIITYRNL or SIINFEKL and a SCT (H-2K^d^) construct with peptide sequence SYFPEITHI (termed SCT-K^b^-KIIT, SCT-K^b^-SIIN and SCT-K^d^-SYFP, respectively) were codon-optimised and synthesised by GeneArt (Thermo Fisher Scientific). A further H2-K^d^ sequence incorporating the Y84C and A139C mutations (termed HC-K^d^-YCAC), was codon-optimised and synthesised by GenScript. All synthesised genes were delivered in pcDNA3.1^+^ plasmids. The full-length native chicken ovalbumin (OVA) gene inserted in a pcDNA3.1^+^ plasmid (clone ID: OGa28271) was purchased from GenScript. Gene inserts from pcDNA3.1^+^ plasmids were cloned into the pAM2AA backbone incorporating the liver-specific human *α*-1 antitrypsin promoter and human ApoE enhancer flanked by AAV2 inverted terminal repeats. Each gene was then packaged into an AAV2/8 vector, purified, and quantitated as previously described(37). AAV Vectors were either produced in-house or by the Vector and Genome Engineering Facility, Children’s Medical Research Institute, Westmead, Australia. Vector aliquots were stored at - 80°C. All vectors were used at a dose of 5×10^11^ vgc except SCT-K^b^-KIIT which was used at 2×10^12^ vgc. Vectors in 500 μL sterile PBS were administered by intravenous injection under general anaesthesia.

### Skin transplantation

Skin transplantation was performed as outlined previously(3), and in the supplementary methods. Grafts were deemed rejected when less than 20% of the viable skin graft remained. B10.BR and B10.BR-RAG mice (H-2^k^) received H-2K^b^ singly-mismatched allogeneic skin grafts from 178.3 strain donor mice. C57BL/6, *Tap1*^fl/fl^ and *Tap1*KOHep mice (H-2^b^) received H-2K^d^ singly-mismatched allogeneic skin grafts from B6.Kd donor mice. BALB/c mice (H-2^d^) received fully allogeneic skin grafts from C57BL/6 donor mice. Skin transplant donors and recipients were sex-matched. Isolation of infiltrating cells from skin grafts is described in the supplementary methods.

### Isolation of hepatocytes and leukocytes from liver, spleen and draining lymph nodes

These procedures were performed as described previously(3). Hepatocytes for immunoaffinity purification experiments underwent further washing with cold PBS before being stored at -80°C.

### Adoptive transfers

Lymphocytes from portal and mesenteric lymph nodes were collected and processed as above(3). Cells were resuspended in RPMI 1640 medium containing 10% FCS was labelled with 10μM CFSE dye (Thermo Fisher Scientific, catalogue# C34570) for 4 minutes at RT. The reaction was quenched by adding more RPMI 1640 medium containing 10% FCS. CFSE-labelled lymphocytes were washed with cold RMPI/FCS10 medium, filtered through 40μm nylon mesh and resuspended in 500μL cold sterile PBS. CFSE-labelled lymphocytes were administered via intravenous injection under general anaesthesia. CFSE-labelling and cell viability were assessed using flow cytometry.

### Histology and immunostaining

Immunohistochemical staining and histology were performed as described previously(3), and in the supplementary methods. Stained sections were examined by a blinded observer.

### Confocal imaging

The livers of freshly-sacrificed mice were perfused retrogradely via the IVC(3) with 3 ml of PBS and then 10 ml of 2% paraformaldehyde (Sigma, catalogue# 30525-89-4) in PBS. The gallbladder was removed and the liver was fixed in 2% paraformaldehyde in PBS for 8 hours. A section of the liver was embedded in 3% agarose (Fisher Biotec, catalogue# AGR-LM-50) and 150μm thick sections were cut using a Vibratome 1000 Plus Sectioning System (Harvard Apparatus, Holliston MA). Sections were blocked with 4% bovine serum albumin (Tocris bioscience, catalogue# 9048-46-8), 5% normal goat serum (Invitrogen, catalogue# 31873) and 0.3% Triton-X 100 (Sigma, catalogue# 9002-93-1) in PBS for 20 hours at 4°C. Sections were stained with primary antibodies for 20 hours at 4°C, washed, then incubated with secondary antibodies for 20 hours at 4°C, followed by staining with DAPI (Sigma, catalogue# 28718-90-3) for 1 hour at 4°C. Primary and secondary antibodies were diluted in blocking buffer. Washing buffer comprised 0.1% Triton-X 100 in PBS. Images were acquired using a Leica SP8 confocal microscope at 93x objective magnification with a numerical aperture of 1.35. The images were analysed using Imaris v9.5 (Oxford instruments).

### Flow cytometry

Cells resuspended in cold staining buffer (2% FCS in PBS) were blocked with mouse Fc Block (BD Biosciences, catalogue# 553141) for 10 minutes at 4°C and stained with a panel of antibodies (Supplementary Table 5). Cells were washed twice with PBS before staining with viability dyes Zombie NIR (BioLegend, catalogue# 423105) or LIVE/DEAD Fixable Blue (Thermo Fisher, catalogue# L23105) for 15 minutes at RT. Cells were then washed with staining. Sample data was acquired using Fortessa X-20, LSR-II (both BD Biosciences) or Cytek Aurora 5L (Cytek Biosciences) instruments and analysed using FlowJo v10 (BD).

### ELISpot

IFN-*γ* ELISpot assays were performed according to the manufacturer’s protocol (U-Cytech, catalogue# CT317-PR5). RMA-S cells either pulsed with peptides or transiently transfected with SCT plasmids, were irradiated with a dose of 3000 rad. Responders were either OT-I-RAG or Des-RAG splenocytes. For pre-stimulation, 1×10^6^ irradiated stimulator cells and 1×10^6^ responder splenocytes were suspended in 250 μL of RPMI/FCS10 medium with penicillin-streptomycin in each well of a 96-well U-bottom plate (Corning, catalogue# 3788). They were cultured at 37°C with 5% CO_2_ for 24 hours and then transferred into an antibody-coated polyvinylidene difluoride (PVDF) plate, serially diluted, and incubated for a further 16 hours. The plates were then developed, and the spots were counted using an AID iSpot plate reader as previously described(1).

### pMHC multimer staining

The pMHC multimer staining method was adapted from that of Dolton *et al.*(38). Cells were incubated with a protein kinase inhibitor, 50nM dasatinib, for 30 minutes at 37°C. PE- or APC-conjugated tetramers or dextramers were centrifuged at 16,000 *g* for 1 minute to remove aggregates. The cells were stained with 0.5μg of pMHC tetramer or pMHC dextramer at 6.4nM in 50μL (unless stated otherwise) for 30 minutes on ice or for one hour at room temperature. Following pMHC multimer staining, the cells were washed with cold staining buffer twice. Samples were incubated with mouse Fc Block (BD Biosciences, catalogue# 553141) for 10 minutes at 4°C and either or both mouse anti-PE and anti-APC were added at 0.5μg/100μL depending on the pMHC multimer conjugates used. The cells were washed and incubated with a cocktail of antibodies against surface markers for 30 minutes on ice. Viability staining, acquisition and analysis were performed as above. Dextramers were purchased from Immudex. QuickSwitch Custom Tetramer Kits (MBL International) were utilised to generate multiple tetramers with selected peptides in order to screen an array of pMHC epitopes. Quantitation of peptide exchange with selected peptides was performed according to the manufacturer’s protocol.

### Immunoaffinity purification

Two replicate samples were prepared for each tissue or experimental group. MHC complexes were isolated from the supernatant of ultracentrifuged tissue lysates by immunoaffinity purification using solid-phase-bound monoclonal antibodies SF1-1.1.10 (anti-H-2K^d^), K9-178 (anti-H-2K^b^), Y3 (anti-H-2K^b^/K^k^) and 28.14.8s (anti-H-2D^b^) as described previously(28, 39). Further information can be found in the supplementary methods.

### Mass Spectrometry

Reconstituted peptides were analysed by LC-MS/MS using an information-dependent acquisition strategy on a Q-Exactive Plus Hybrid Quadrupole Orbitrap (Thermo Fisher Scientific, Bremen, Germany) coupled to a Dionex UltiMate 3000 RSLCnano system (Thermo Fisher Scientific) as described previously(28). Further detail is available in the supplementary methods. The mass spectrometry data have been deposited to the ProteomeXchange Consortium via the PRIDE(40) partner repository with accession number PXD022695.

### Data Dependent Analysis (DDA)

For peptide identification, the acquired raw files were searched with PEAKS Studio X+ (Bioinformatics Solutions) against the *Mus musculus* (SwissProt) database. The parent mass error tolerance was set to 10 ppm and the fragment mass error tolerance to 0.02 Da. Oxidation of methionine (M) was set as variable modifications and a false-discovery rate (FDR) cut-off of 5% was applied (41).

### Data Independent Analysis (DIA)

PEAKS Studio X+ was used to generate a spectral library from all DDA data. DIA raw files were imported to Spectronaut 11 Pulsar (Biognosys). The parent mass error tolerance was set to 10 ppm and the fragment mass error tolerance to 0.02 Da. Oxidation of M was set as variable modifications and a peptide list was exported at Q-value = 1%.

### Immunopeptidomic analysis

Known contaminants were removed from the analysis. Unique peptides from the DDA (replicate 1) and DIA (replicate 2) datasets were combined to increase the coverage of the tissue immunopeptidomes. For analysis requiring spectral intensity values, only DDA datasets were used. Binding motifs of 8-mer peptides from samples that were immunoaffinity purified with K9-178 antibody (H-2K^b^ group) and 9-mer peptides from samples that had been immunoaffinity purified with SF1-1.1.10 antibody (H-2K^d^ group) were visualised using the GibbsCluster2.0 algorithm (NetMHC4.0)(42). For comparison of unique H-2K^b^ peptides between different tissues, 8-mer to 11-mer peptides with a predicted half-maximum inhibitory concentration (IC_50_) of binding to H-2K^b^ less than 500nM (NetMHC4.0 database) were selected. For comparison of unique H-2K^d^ peptides between tissues, all 9-mer peptides were included. The source proteins associated with the eluted peptides were analysed using the PANTHER Gene Ontology classification system(43). Function classification analysis and statistical over-representation tests were performed(44).

### Validation of peptide identification using retrospectively synthesised peptides

We validated the identity of a panel of peptides by comparing chromatographic retention and MS/MS spectra of synthesised peptides (GL Biochem, Shanghai) with those of the corresponding eluted peptides, as previously reported(45) and outlined in the supplementary methods.

### Statistical Analysis and Data Visualisation

Data are represented as mean±SEM unless otherwise stated. Unpaired Student’s t-tests were performed to calculate statistical differences in a single variable between the means of two groups and one-way analysis of variance (ANOVA) in conjunction with Sidak’s multiple comparison tests were used to calculate statistical differences between the means of three or more groups. For analysis of 2 variables, two-way ANOVA with Sidak’s multiple comparison tests were used. Graft survival curves were compared using Mantel Cox log-rank tests. Synthetic and corresponding eluted peptide spectra were compared using Pearson correlation tests. The relationship between overall peptide abundance and alloreactive T cell binding, and the impact of differences in sexes and strains on alloreactive T cell binding, were analysed using linear regression and Pearson correlation tests. P<0.05 was considered significant. Statistical tests were performed using GraphPad Prism version 8.01 (GraphPad Software, La Jolla CA). Heatmaps were generated using Morpheus software (https://software.broadinstitute.org/morpheus).

### Study Approvals

All animal procedures were approved by the University of Sydney Animal Ethics Committee (protocol 2017/1253) and carried out in accordance with the Australian code for the care and use of animals for scientific purposes.

### Author contributions

E.T.S., P.B., D.G.B., A.W.P., N.A.M. and A.F.S. designed experiments. E.T.S., P.F., M.P-H., M.L., K.E., S.H.R., A.B., N.L.D. and N.A.M. performed experiments. I.E.A., L.L., P.B., and A.W.P. provided reagents and/or samples. E.T.S., P.F., S.H.R., A.W.P., N.A.M. and A.F.S. analysed data. E.T.S., P.F., S.H.R., N.A.M. and A.F.S. wrote the manuscript. All authors read and approved the manuscript.

## Acknowledgements

Dr Nathan P. Croft (Monash University) for discussions of RMA-S peptide-pulsing experiments. Technical assistance from R. Ayala and I. Hanchapola (Monash University). The authors acknowledge the provision of instrumentation and technical support by the Monash Biomedical Proteomics Facility, and by Sydney Cytometry. Computational resources were supported by the R@CMon/Monash Node of the NeCTAR Research Cloud. We thank Laboratory Animal Services, University of Sydney, for superb mouse care, and the Vector and Genome Engineering Facility, CMRI for AAV vector production and advice. A number of funding bodies provided project and people support. These include the National Health and Medical Research Council of Australia (NHMRC); Principal Research Fellowship (1137739) to A.W.P., Ideas Grant (APP1183806) to A.F.S. and N.A.M, project grants (APP1108311, APP1156431 and APP1161583) to L.L. and (APP1086451 and APP1146677) to P.B. and D.G.B. M.L. and K.E. received Research Training Program Postgraduate Awards from the Department of Education and Training, Australian Federal Government, while M.P-H. received an Earl Owen Fellowship from the Sydney Medical School Foundation, and E.T.S. was supported by the Royal Prince Alfred Hospital Transplant Institute. LL held research grants from the Department of Science and Higher Education of Ministry of National Defense, Republic of Poland, (“Kościuszko” k/10/8047/DNiSW/T–WIHE/3) and the National Science Centre, Republic of Poland (OPUS13) (UMO-2017/25/B/NZ1/02790). P.F. was granted the Victorian Cancer Agency Fellowship. This project was also supported by Grants-in-aid from Sydney Medical School Foundation (to A.F.S and D.G.B.) and the Myee Codrington Medical Research Foundation (to A.F.S. and E.T.S).

## References

1. Cunningham EC, Tay SS, Wang C, Rtshiladze M, Wang ZZ, McGuffog C, et al. Gene therapy for tolerance: high-level expression of donor major histocompatibility complex in the liver overcomes naive and memory alloresponses to skin grafts. Transplantation. 2013;95(1):70–7.

2. Le Guen V, Judor JP, Boeffard F, Gauttier V, Ferry N, Soulillou JP, et al. Alloantigen gene transfer to hepatocytes promotes tolerance to pancreatic islet graft by inducing CD8(+) regulatory T cells. J Hepatol. 2017;66(4):765–77.

3. Paul-Heng M, Leong M, Cunningham E, Bunker DLJ, Bremner K, Wang Z, et al. Direct recognition of hepatocyte-expressed MHC class I alloantigens is required for tolerance induction. JCI Insight. 2018;3(15).

4. Caron E, Vincent K, Fortier MH, Laverdure JP, Bramoulle A, Hardy MP, et al. The MHC I immunopeptidome conveys to the cell surface an integrative view of cellular regulation. Mol Syst Biol. 2011;7:533.

5. Heath WR, Kane KP, Mescher MF, and Sherman LA. Alloreactive T cells discriminate among a diverse set of endogenous peptides. Proc Natl Acad Sci U S A. 1991;88(12):5101–5.

6. Heath WR, and Sherman LA. Cell-type-specific recognition of allogeneic cells by alloreactive cytotoxic T cells: a consequence of peptide-dependent allorecognition. Eur J Immunol. 1991;21(1):153–9.

7. Aosai F, Ohlen C, Ljunggren HG, Hoglund P, Franksson L, Ploegh H, et al. Different types of allospecific CTL clones identified by their ability to recognize peptide loading-defective target cells. Eur J Immunol. 1991;21(11):2767–74.

8. Udaka K, Tsomides TJ, and Eisen HN. A naturally occurring peptide recognized by alloreactive CD8+ cytotoxic T lymphocytes in association with a class I MHC protein. Cell. 1992;69(6):989–98.

9. Crumpacker DB, Alexander J, Cresswell P, and Engelhard VH. Role of endogenous peptides in murine allogenic cytotoxic T cell responses assessed using transfectants of the antigen-processing mutant 174xCEM.T2. Journal of immunology. 1992;148(10):3004–11.

10. Kuzushima K, Sun R, van Bleek GM, Vegh Z, and Nathenson SG. The role of self peptides in the allogeneic cross-reactivity of CTLs. Journal of immunology. 1995;155(2):594–601.

11. Guimezanes A, Barrett-Wilt GA, Gulden-Thompson P, Shabanowitz J, Engelhard VH, Hunt DF, et al. Identification of endogenous peptides recognized by in vivo or in vitro generated alloreactive cytotoxic T lymphocytes: distinct characteristics correlated with CD8 dependence. Eur J Immunol. 2001;31(2):421–32.

12. D’Orsogna LJ, Nguyen TH, Claas FH, Witt C, and Mifsud NA. Endogenous-peptide-dependent alloreactivity: new scientific insights and clinical implications. Tissue Antigens. 2013;81(6):399–407.

13. Mullbacher A, Hill AB, Blanden RV, Cowden WB, King NJ, and Hla RT. Alloreactive cytotoxic T cells recognize MHC class I antigen without peptide specificity. Journal of immunology. 1991;147(6):1765–72.

14. Jankovic V, Remus K, Molano A, and Nikolich-Zugich J. T Cell Recognition of an Engineered MHC Class I Molecule: Implications for Peptide-Independent Alloreactivity. The Journal of Immunology. 2002;169(4):1887–92.

15. Smith PA, Brunmark A, Jackson MR, and Potter TA. Peptide-independent recognition by alloreactive cytotoxic T lymphocytes (CTL). The Journal of experimental medicine. 1997;185(6):1023–33.

16. Yu YY, Netuschil N, Lybarger L, Connolly JM, and Hansen TH. Cutting edge: single-chain trimers of MHC class I molecules form stable structures that potently stimulate antigen-specific T cells and B cells. Journal of immunology. 2002;168(7):3145–9.

17. Truscott SM, Lybarger L, Martinko JM, Mitaksov VE, Kranz DM, Connolly JM, et al. Disulfide bond engineering to trap peptides in the MHC class I binding groove. Journal of immunology. 2007;178(10):6280–9.

18. Hein Z, Uchtenhagen H, Abualrous ET, Saini SK, Janssen L, Van Hateren A, et al. Peptide-independent stabilization of MHC class I molecules breaches cellular quality control. J Cell Sci. 2014;127(Pt 13):2885–97.

19. Jutte NH, Knoop CJ, Heijse P, Balk AH, Mochtar B, Claas FH, et al. Human heart endothelial-cell-restricted allorecognition. Transplantation. 1996;62(3):403–6.

20. Deckers JG, Boonstra JG, Van der Kooij SW, Daha MR, and Van der Woude FJ. Tissue-specific characteristics of cytotoxic graft-infiltrating T cells during renal allograft rejection. Transplantation. 1997;64(1):178–81.

21. Martin-Galiano AJ, and Lopez D. Computational characterization of the peptidome in transporter associated with antigen processing (TAP)-deficient cells. PLoS One. 2019;14(1):e0210583.

22. Boulanger DSM, Eccleston RC, Phillips A, Coveney PV, Elliott T, and Dalchau N. A Mechanistic Model for Predicting Cell Surface Presentation of Competing Peptides by MHC Class I Molecules. Front Immunol. 2018;9:1538.

23. Van Kaer L, Ashton-Rickardt PG, Ploegh HL, and Tonegawa S. TAP1 mutant mice are deficient in antigen presentation, surface class I molecules, and CD4-8+ T cells. Cell. 1992;71(7):1205–14.

24. Saini SK, Tamhane T, Anjanappa R, Saikia A, Ramskov S, Donia M, et al. Empty peptide-receptive MHC class I molecules for efficient detection of antigen-specific T cells. Sci Immunol. 2019;4(37).

25. Scifo C, Mekaelian L, Munyazesa E, Schmitt-Verhulst AM, and Guimezanes A. Selection of T-cell receptors with a recurrent CDR3beta peptide-contact motif within the repertoire of alloreactive CD8(+) T cells. Eur J Immunol. 2011;41(8):2414–23.

26. Siu JHY, Surendrakumar V, Richards JA, and Pettigrew GJ. T cell Allorecognition Pathways in Solid Organ Transplantation. Front Immunol. 2018;9:2548.

27. Croft NP, Smith SA, Wong YC, Tan CT, Dudek NL, Flesch IE, et al. Kinetics of antigen expression and epitope presentation during virus infection. PLoS Pathog. 2013;9(1):e1003129.

28. Croft NP, Smith SA, Pickering J, Sidney J, Peters B, Faridi P, et al. Most viral peptides displayed by class I MHC on infected cells are immunogenic. Proc Natl Acad Sci U S A. 2019;116(8):3112–7.

29. Ekeruche-Makinde J, Miles JJ, van den Berg HA, Skowera A, Cole DK, Dolton G, et al. Peptide length determines the outcome of TCR/peptide-MHCI engagement. Blood. 2013;121(7):1112–23.

30. DeVette CI, Andreatta M, Bardet W, Cate SJ, Jurtz VI, Jackson KW, et al. NetH2pan: A Computational Tool to Guide MHC Peptide Prediction on Murine Tumors. Cancer Immunol Res. 2018;6(6):636–44.

31. Krebs P, Scandella E, Odermatt B, and Ludewig B. Rapid functional exhaustion and deletion of CTL following immunization with recombinant adenovirus. J Immunol. 2005;174(8):4559–66.

32. Miller ML, McIntosh CM, Williams JB, Wang Y, Hollinger MK, Isaad NJ, et al. Distinct Graft-Specific TCR Avidity Profiles during Acute Rejection and Tolerance. Cell Rep. 2018;24(8):2112–26.

33. Bentzen AK, Marquard AM, Lyngaa R, Saini SK, Ramskov S, Donia M, et al. Large-scale detection of antigen-specific T cells using peptide-MHC-I multimers labeled with DNA barcodes. Nature Biotechnology. 2016;34(10):1037–45.

34. Bentzen AK, Such L, Jensen KK, Marquard AM, Jessen LE, Miller NJ, et al. T cell receptor fingerprinting enables in-depth characterization of the interactions governing recognition of peptide-MHC complexes. Nat Biotechnol. 2018.

35. Woodham AW, Zeigler SH, Zeyang EL, Kolifrath SC, Cheloha RW, Rashidian M, et al. In vivo detection of antigen-specific CD8(+) T cells by immuno-positron emission tomography. Nat Methods. 2020;17(10):1025–32.

36. Hess SM, Young EF, Miller KR, Vincent BG, Buntzman AS, Collins EJ, et al. Deletion of naive T cells recognizing the minor histocompatibility antigen HY with toxin-coupled peptide-MHC class I tetramers inhibits cognate CTL responses and alters immunodominance. Transpl Immunol. 2013;29(1-4):138–45.

37. Cunningham SC, Dane AP, Spinoulas A, and Alexander IE. Gene Delivery to the Juvenile Mouse Liver Using AAV2/8 Vectors. Mol Ther. 2008;16(6):1081–8.

38. Dolton G, Tungatt K, Lloyd A, Bianchi V, Theaker SM, Trimby A, et al. More tricks with tetramers: a practical guide to staining T cells with peptide-MHC multimers. Immunology. 2015;146(1):11–22.

39. Dudek NL, Perlmutter P, Aguilar MI, Croft NP, and Purcell AW. Epitope discovery and their use in peptide based vaccines. Curr Pharm Des. 2010;16(28):3149–57.

40. Perez-Riverol Y, Csordas A, Bai J, Bernal-Llinares M, Hewapathirana S, Kundu DJ, et al. The PRIDE database and related tools and resources in 2019: improving support for quantification data. Nucleic Acids Res. 2019;47(D1):D442–D50.

41. Faridi P, Li C, Ramarathinam SH, Vivian JP, Illing PT, Mifsud NA, et al. A subset of HLA-I peptides are not genomically templated: Evidence for cis- and trans-spliced peptide ligands. Sci Immunol. 2018;3(28).

42. Andreatta M, and Nielsen M. Gapped sequence alignment using artificial neural networks: application to the MHC class I system. Bioinformatics. 2016;32(4):511–7.

43. Mi H, Muruganujan A, Ebert D, Huang X, and Thomas PD. PANTHER version 14: more genomes, a new PANTHER GO-slim and improvements in enrichment analysis tools. Nucleic Acids Res. 2019;47(D1):D419–D26.

44. Mi H, Muruganujan A, Huang X, Ebert D, Mills C, Guo X, et al. Protocol Update for large-scale genome and gene function analysis with the PANTHER classification system (v.14.0). Nat Protoc. 2019;14(3):703–21.

45. Falth M, Svensson M, Nilsson A, Skold K, Fenyo D, and Andren PE. Validation of endogenous peptide identifications using a database of tandem mass spectra. Journal of Proteome Research. 2008;7(7):3049–53.

