## Supplementary Material for "The self-peptide repertoire plays a critical role in transplant tolerance induction"

#### Methods

##### Mice

Unless otherwise stated, mice were bred at the University of Sydney (Camperdown, Australia). 178.3 mice (originally provided by Drs. W. Health and M. Hoffman, Walter and Eliza Hall Institute, Melbourne, Australia) express the transgenic MHC class I molecule H-2K$^b$ ubiquitously, under the control of its own promoter, on a B10.BR (H-2$^k$) background. Des-TCR mice were provided by Dr. Patrick Bertolino. Des-TCR mice express an alloreactive TCR, which recognises the peptides KVITFIDL, KVLHFYNV and KIITYRNL restricted by H-2K$^b$. Des-TCR is identifiable by a clonotypic mAb (Désiré). Des-RAG mice (CD45.1$^+$) were obtained by crossing Des-TCR mice with *Ptprc$^{a/a}$* *Rag1$^{-/-}$* mice, also on a B10.BR (H-2$^k$) background. These mice were obtained from Dr. Barbara Fazekas at the University of Sydney. OT-I mice carry a TCR which recognises the peptide SIINFEKL presented by H-2K$^b$. OT-I were crossed with *Rag1$^{-/-}$* mice to create the OT-I-RAG line. These mice were provided by Patrick Bertolino, and were bred at the Centenary Institute. C57BL/6J$^{Arc}$ (H-2$^b$) and BALB/c$^{Arc}$ (H-2$^d$) mice (herein termed C57BL/6 and BALB/c) were purchased from the Animal Resources Centre, Perth, Australia. B6.Kd mice(1) express an H-2K$^d$ transgene ubiquitously on a C57BL/6 (H-2$^b$) background. B6.Kd mice were originally developed by R. Pat Bucy at the University of Alabama (Tuscaloosa, Alabama, USA) and were provided by Dr Robert Fairchild, Cleveland Clinic (Cleveland, Ohio, USA). B6.Kd mice were backcrossed for 4 generations to C57BL/6J$^{Arc}$, prior to use. *Tap1*KOHep mice were generated based on the conditional-ready strain 09400, C57BL/6N-*Tap1*<tm2a(EUCOMM)Hmgu>/Ieg, developed as part of the European Conditional Mouse Mutagenesis programme (EUCOMM)(2). Mice heterozygous for the *Tap1*tm2a allele on the C57BL/6N genetic background were obtained from the European Mutant Mouse Archive, based at Helmholtz Zentrum. These mice were backcrossed to C57BL/6J$^{Arc}$ for 3 generations, then intercrossed with FLPo deleter (B6.129S4-*Gt(ROSA)26SOR$^{tm2(FLPo)Sor}$*/J) mice(3) (imported from the Jackson Laboratory, Bar Harbor, ME) to generate mice carrying the *Tap1*tm2c (floxed) allele. FLPo was bred out by backcrossing to C57BL/6J$^{Arc}$ (2 generations), following which the mice were crossed to Albumin-Cre mice (B6.FVB(129)-*Tg(*$^{Alb1-}$

$^{cre)1Dlr}$/J)(4), provided by Dr Patrick Bertolino. *Tap1*KOHep mice are homozygous for the floxed *Tap1* allele (*Tap1*tm2c) and have one copy of *Cre*, which is expressed exclusively in hepatocytes resulting in hepatocyte-specific deletion of the floxed *Tap1* allele. Genotyping and genetic background testing was performed on earpunch tissue, isolated hepatocytes or spleen by Transnetyx (Cordova, TN, USA). The genetic background of *Tap1*KOHep and *Tap1*$^{fl/fl}$ control mice was at least 91.3% C57BL/6J (91.3-97.9%) and these mice did not reject syngeneic skin grafts from C57BL/6J$^{Arc}$ donors (not shown). Further characterisation of this strain is shown in Figure 4 and Supplementary Figure 5. Animals were randomly allocated to treatment groups. Male and female mice aged between 8 and 12 weeks were used in this study. Male mice were used unless stated otherwise. At the termination of each experiment, tissues were collected under general anaesthesia. Frozen tissues were stored at -80°C.

##### Skin Transplantation

Full-thickness grafts of 1x1 cm$^2$ tail skin from donor mice were applied to the dorsum of anaesthetised recipient mice following excision of a 1x1 cm$^2$ area of skin to accommodate the donor skin graft. The graft was fixed using cyanoacrylate tissue adhesive (Dermabond, Ethicon, catalogue# ANX12) and bandaged. Mice received analgesia with buprenorphine (Temgesic, Schering-Plough, 0.05 mg kg$^{-1}$ s.c.), prophylactic ampicillin (Alphapharm, 100 mg kg$^{-1}$ s.c.) and 0.5 ml of warmed saline. The bandage was removed 7-10 days later and the grafts were monitored frequently for up to 100 days post-transplant.

##### Isolation of leukocytes from skin grafts

Skin grafts were collected from recipient mice and the subcutaneous tissue was removed. They were sectioned into 4 mm$^2$ pieces and washed with HBSS (Lonza, catalogue #10-543F) supplemented with 0.5 mM EDTA (MilliporeSigma, catalogue # E6758) and 10% FCS, followed by incubation with 15 mL of HBSS supplemented with 5 mM $CaCl_2$ (MilliporeSigma, catalogue #C5670), 1 mg/mL Collagenase D (11088866001, Roche) and 10% FCS for 1 hour at 37°C while shaking at 150 rpm. Digested tissue was gently pushed through a 70 μm nylon mesh strainer. Dissociated cells were washed with RPMI/FCS2 medium, resuspended in 15 mL PBS and then mixed with 9 mL of isotonic Percoll PLUS (Cytiva, catalogue

#GE17-5445-01). Following centrifugation at 500 *g* for 15 minutes (room temperature) floating debris and excess solution were aspirated and the cell pellet was resuspended in RPMI/FCS2 medium.

##### Histology and immunostaining

For immunohistochemical staining, OCT-embedded frozen tissues were cut into 6µm thick sections. Sections were allowed to air dry for 1 hour at room temperature (RT) prior to fixation in acetone for 8 minutes at RT. Sections were blocked with 20% normal mouse serum (MilliporeSigma, catalogue# M5905) and 5% normal porcine serum (Thermo Fisher Scientific, catalogue# 31890) for 20 minutes at RT and stained with FITC-conjugated primary antibodies or the corresponding isotype controls (listed in Supplementary Table 5) for 30 minutes at RT. Sections were then incubated with horseradish peroxidase-conjugated rabbit-anti-FITC secondary antibody before development with diaminobenzidine (DAB) substrate chromogen system (Dako, catalogue# K3468). Sections were counterstained in Mayer's hematoxylin solution (MilliporeSigma, catalogue# MHS16) for 2 minutes and mounted with Fronine safety mount No.4 (Thermo Fisher Scientific, catalogue# FNNII068). Tissue processing and H&E staining were performed by the Histopathology Laboratory, Discipline of Pathology, Sydney Medical School. For H&E staining, 5µm thick sections from formalin-fixed paraffin-embedded tissues were used.

##### Hybridoma antibody production and purification

Hybridoma cell lines SF1-1.1.10 (anti-H-2K$^d$), K9-178 (anti-H-2K$^b$), Y3 (anti-H-2K$^b$/K$^k$) and 28.14.8s (anti-H-2D$^b$) were cultured in either RF5 or RF10 in a roller bottle (cat # CLS431134, Corning) at 37°C, 5%$CO_2$. Both cells and supernatant were harvested when the optimal density of 1-2×$10^8$ cells was reached and the supernatant was passed through a 0.22 *µ*m filter (cat #CLS431097, Corning). The Profinia Protein Purification System (Bio-Rad) was used for hybridoma supernatant antibody purification. Here, a Protein A agarose affinity column captures IgG, which is then eluted using 0.1 M citrate buffer (pH 3.0) before being passed over a desalting column to recover the purified antibody in PBS.

#### Immunoaffinity Purification

Around 1×$10^8$ purified hepatocytes from 4-5 mice were pooled per sample. Hepatocytes were lysed in 0.5% IGEPAL, 50 mM Tris (pH 8), 150 mM NaCl and protease inhibitors (Roche cOmplete Protease Inhibitor Cocktail; Merck, catalogue# 11836145001). Spleens, skin grafts (d7 post-transplant) or tail skins from 5 - 9 donors were pooled per sample. Spleen and skin samples were ground in a Retsch Mixer Mill MM 400 under cryogenic conditions and then lysed in 0.5% IGEPAL, 50 mM Tris (pH 8), 150 mM NaCl, and protease inhibitors. Lysates were incubated for 1 hour at 4°C, then cleared by ultracentrifugation (40,000 rpm, 30 min) and MHC complexes were isolated from supernatant by immunoaffinity purification using solid-phase-bound monoclonal antibodies SF1-1.1.10 (anti H-2$K^d$), K9-178 (anti H-2$K^b$), Y3 (anti H-2$K^b$/$K^k$) and 28.14.8s (anti H-2$D^b$) as described previously. Peptides were dissociated from the MHC with 10% acetic acid. For purified hepatocyte and spleen samples, the mixture of peptides, class I HC and β2m was fractionated on a 4.6 mm internal diameter × 100 mm monolithic C18 column (Chromolith SpeedROD; Merck Millipore, catalogue# 1021290001) using an ÄKTAmicro RP-HPLC (GE Healthcare) system, running a mobile phase consisting of buffer A (0.1% trifluoroacetic acid; Thermo Fisher Scientific) and buffer B (80% acetonitrile, 0.1% trifluoroacetic acid; Thermo Fisher Scientific), 1 mL $min^{-1}$ with a gradient of B of 2–40% over 4 min, 40–45% over 4 min and 45–99% over 2 min, collecting 500 µL fractions. Peptide-containing fractions were either unpooled or combined into pools, vacuum-concentrated and reconstituted in 0.1% formic acid (Thermo Fisher Scientific) for mass spectrometry analysis. For tail skin samples, the mixture of peptides, class I HC and β2m was purified using Millipore 5 kDa Amicon centrifugal units (Human Metabolome Technologies; catalogue# UFC3LCCNB_HMT) in 0.1% trifluoroacetic acid. Peptides were extracted and desalted from the filtrate using ZipTip C18 pipette tips (Agilent Technologies, catalogue# A57003100K) in a final buffer of 30% acetonitrile, 0.1% trifluoroacetic acid. Peptide samples were vacuum-concentrated and reconstituted in 0.1% formic acid for mass spectrometry analysis.

#### Mass Spectrometry

Reconstituted peptides were trapped on a 2 cm Nanoviper PepMap100 trap column at a flow rate of 15 min using a RSLC nano-HPLC. The trap column was then switched inline to an analytical PepMap100 C18 nanocolumn (75 µm x 50 cm, 3 µm 100 Å pore size) at a flow rate of 250 nL/min using an initial gradient of 2.5% to 7.5% buffer B (0.1% formic acid 80% ACN) in buffer A (0.1% formic acid in water) over 1 min followed with a linear gradient from 7.5% to 32.5% buffer B for 58 min followed by a linear increase to 40% buffer B over 5 min and an additional increase up to 99% buffer B over 5 min. Survey full scan MS spectra (m/z 375–1800) were acquired in the Orbitrap with 70,000 resolution (m/z 200) after the accumulation of ions to a $5\times10^5$ target value with a maximum injection time of 120 ms. For Data Dependant Acquisition (DDA) runs, the 12 most intense multiply charged ions ($z \geq 2$) were sequentially isolated and fragmented by higher-energy collisional dissociation (HCD) at 27% with an injection time of 120 ms, 35,000 resolution and target of $2\times10^5$ counts. An isolation width of 1.8 m/z was applied and underfill ratio was set to 1% and dynamic exclusion to 15 sec. For Data Independent Acquisition (DIA) runs, the MS1 survey scan and fragment ions were acquired using variable windows (Supplementary Table 6) at 35,000 resolution with an automatic gain control (AGC) target of 3e6 ions.

**Validation of peptide identification using retrospectively synthesised peptides**

We validated the identity of a panel of peptides by comparing chromatographic retention and MS/MS spectra of synthesised peptides (GL Biochem, Shanghai) with those of the corresponding eluted peptides. The PKL files of the synthetic and eluted peptides were exported from PEAKS X plus studio software. To evaluate the similarity between two spectra, we predicted all b- and y-ions for each sequence and then extracted the intensity for each ion (with a fragment mass error tolerance of 0.02 Da). The Pearson correlation coefficient and the corresponding p-value between the $\log_{10}$ intensities of identified b- and y-ions in the synthetic and sample-derived spectra were calculated. The closer the correlation coefficient to 1, the greater identity between paired spectra. All tested peptides were found to have a *p*-value of less than 0.05.

**Supplementary Figure 1. Amino acid sequences for the SCT-$K^b$-SIIN, SCT-$K^b$-KIIT, SCT-$K^d$-SYFP and HC-$K^d$-YCAC constructs.**

**Supplementary Figure 2. Expression of SCT-$K^b$-peptide in mouse hepatocytes is robust and persistent.**

B10.BR mice were inoculated with AAV-SCT-$K^b$-KIITYRNL **(A)** or AAV-SCT-$K^b$-SIINFEKL **(B)**. On days 2-100 post-inoculation, tissues were collected for analysis (n=3/interval). Representative immunostained (IHC) and H&E images show transduced liver sections (200 $\mu$m). Robust expression of H2-$K^b$ was present through day 100 post-inoculation. Histologic examination of the liver sections was normal. Levels of aspartate aminotransferase (AST) and alanine aminotransferase (ALT) did not increase significantly from baseline in mice treated with AAV-SCT-$K^b$-KIITYRNL (one-way ANOVA, $p = 0.4$ for AST and $p = 0.21$ for ALT) or in mice receiving AAV-SCT-$K^b$-SIINFEKL (one-way ANOVA, $p = 0.13$ for AST and $p = 0.02$ for ALT, due solely to a decrease in ALT on d4). Minimal infiltration with cells expressing the markers CD4, CD8, CD11c or CD19 was detected. Mean $\pm$ SEM are shown, scale bar = 200 $\mu$m.

**Supplementary Figure 3. Recognition of SCT peptide-MHC ligands in vitro and in vivo.**

**(A)** RMA-S cells were pulsed with different concentrations of the peptides KIITYRNL ($Pcid2_{318-325}$), SIINFEKL ($OVA_{257-264}$) or AAAAFAAL (synthetic negative control), or were untreated. Stabilisation of H-2$K^b$ surface expression was assessed by flow cytometry following staining with a conformation-dependent anti-H-2$K^b$ mAb (clone Y3). **(B)** Flow plots shown are representative of three independent experiments. Peptide concentrations required to achieve equivalent H-2$K^b$ surface expression levels were determined. **(C)** RMA-S cells were transiently transfected with constructs encoding SCT-$K^b$-KIIT, SCT-$K^b$-SIIN and SCT-$K^b$-AAAA using a Lonza-AMAXA Nucleofector 2b. Transgene expression was assessed by flow cytometry (as above) 24 hours after transfection. Flow plots shown are representative of three independent experiments. **(D)** The proportion of cells secreting IFN-$\gamma$ upon recognition of their cognate antigen was determined using ELISPOT assays. Splenocytes from Des-RAG or OT-I-RAG mice were cultured with irradiated stimulators; RMA-S pulsed with selected peptides or expressing SCT constructs after transient transfection. SCT recognition by cognate TCRs mirrored recognition of the native H-2K$^b$-peptide complex. SCT constructs were recognised in a peptide-specific manner *in vitro*. Data from two independent experiments with a total of n = 3 biological replicates per group are shown. **(E)** Des-RAG lymphocytes were labelled with CFSE, adoptively transferred into recipient mice and recovered from the recipient liver two days later. Some recipient mice were treated with AAV encoding SCT-K$^b$-KIIT or SCT-K$^b$-SIIN prior to adoptive transfer, as shown. **(F)** Flow cytometry analysis of CFSE-labelled Des-RAG lymphocytes demonstrates peptide-specific activation and proliferation of adoptively transferred CD8$^+$ Des-RAG T cells upon encounter with their cognate antigen in the liver, confirming that recognition of the SCT-K$^b$-KIIT ligand in vivo was analogous to that of the native pMHC complex. Data from three independent experiments with a total of n = 3 biological replicates per group are shown. **(D, F)** Mean ± SEM are shown, one-way ANOVA in conjunction with Sidak's multiple comparison test: ns, not significant; *$p < 0.05$, ** $p < 0.01$, *** $p < 0.001$, **** $p < 0.0001$.

**Supplementary Figure 4. Recognition of SCT-K$^b$-KIIT in a polyclonal alloreactive population.**

**(A)** Inoculation with SCT-K$^b$-KIIT vector not only activates a clone of transgenic Des-RAG T cells bearing the cognate receptor, but also activates a proportion of the polyclonal T cell repertoire of normal B10.BR mice. B10.BR mice were primed against allogeneic H-2K$^b$ (178.3 skin graft). Approximately 30 days post-graft rejection, some of the primed or naïve B10.BR mice were inoculated with AAV-SCT-K$^b$-KIIT. Liver leukocytes were analysed on day 7 post-inoculation. **(B)** Activated CD8$^+$ T cells, defined as CD44$^+$PD-1$^{hi}$, increased in number following priming or transduction with SCT-K$^b$-KIIT, with a further increase in primed mice receiving SCT-K$^b$-KIIT. **(C-D)** Inoculation of naïve or primed B10.BR mice with AAV-SCT-K$^b$-KIIT generated populations of activated (CD44$^+$PD-1$^{hi}$) CD8$^+$ T cells which bound K$^b$-KIITYRNL dextramers specifically. Dextramers of the syngeneic pMHC K$^k$-EEEPVKKI were used as negative controls. Data from one representative experiment (from n = 3) is shown in **(C)**, while two independent experiments with a total of n = 3 biological replicates per group are shown in **(B, D)**. Data are presented as mean ± SEM, statistical analysis involved two-way analysis of variance (ANOVA) in conjunction with Tukey's multiple comparison test, *$p < 0.05$, ** $p < 0.01$, *** $p < 0.001$, **** $p < 0.0001$).

**Supplementary Figure 5. Characterisation of *Tap1*KOHep mice.**

**(A)** Gating strategy for determination of $K^b$ expression on the surface of hepatocytes, liver leukocytes and splenocytes of *Tap1*KOHep and *Tap1*$^{fl/fl}$ mice. **(B)** Genotyping PCR performed by Transnetyx shows the presence of the Albumin-Cre transgene in hepatocytes, liver leukocytes and spleen from *Tap1*KOHep mice (above). Because Cre activity is restricted to hepatocytes, the recombined *Tap1* sequence specifically detected by the L1L2-Bact-P EX probe is only amplified in hepatocytes and not other tissues from *Tap1*KOHep (below). Data from one experiment with a total of n = 5 biological replicates per group are shown. **(C)** Genetic background analysis was undertaken by Transnetyx. *Tap1*KOHep and *Tap1*$^{fl/fl}$ control mice were at least 91.3% C57BL/6J (91.3-97.9%). Data from one experiment with a total of n = 5 biological replicates per group are shown. **(D)** AST and ALT levels are comparable between *Tap1*KOHep and floxed littermate control mice (n = 3). **(B-D)** Mean ± SEM are shown. **(E)** H-2$K^b$ is expressed at normal levels in the spleen, thymus and lymph node of *Tap1*KOHep and *Tap1*$^{fl/fl}$ mice (scale bar = 100$\mu$m, representative images from n = 3). **(F)** H-2$K^b$ is absent from the hepatocytes of *Tap1*KOHep mice, but detectable on other liver cells (scale bar = 40$\mu$m, representative images from n = 3).

**Supplementary Figure 6. Features of H-2$K^d$-associated peptides.**

The length distribution for H-2$K^d$-associated peptides eluted from transduced hepatocytes in each of four vector/strain combinations is shown in panel **(A)**. Peptides eluted from C57BL/6 mice expressing $K^d$-HC are predominantly nonamers – this preference was less strong for the peptide repertoires of hepatocytes expressing $K^d$-YCAC. **(B-D)** Gene Ontology annotations of the source proteins associated with eluted peptides were analysed using the PANTHER classification system. Function classification analysis and statistical over-representation tests were performed. **(B)** Cellular component and biological process analysis of source proteins corresponding to the same hepatocyte peptide repertoires shown in **(A)**. **(C)** Analysis of the source proteins giving rise to the H-2$K^d$ and $K^b$-associated peptide repertoires of transduced hepatocytes, donor skin grafts and donor spleen. **(D)** A number of Gene Ontology terms were enriched or depleted when hepatocyte source proteins from AAV-HC-$K^d$-YCAC-transduced *Tap1*KOHep mice were compared with those from AAV-HC-$K^d$-treated C57BL/6. The most striking enrichment was in terms associated with mitochondria and mitochondrial metabolism. Significant enrichment was also found for the cellular component terms endoplasmic reticulum, extracellular region and cytoplasm.

**Supplementary Figure 7. Expression of AAV-SCT-$K^d$-SYFPEITHI and AAV-HC-$K^d$-YCAC is strong and durable.**

**(A)** C57BL/6 mice were inoculated with AAV-SCT-$K^d$-SYFPEITHI iv. On days 2, 4, 7, 14, 28 and 100 post-inoculation, tissues were collected for analysis (n = 3 at each interval). Representative IHC and H&E images show transduced liver sections. Robust expression of H-2$K^d$ was present through day 100 post-inoculation. Histologic examination of the liver sections was normal. Levels of AST and ALT did not increase significantly from baseline (one-way ANOVA, $p = 0.14$ for AST and $p = 0.11$ for ALT in mice inoculated with AAV-SCT-$K^d$-SYFPEITHI). Minimal infiltration with cells expressing the markers CD4, CD8, CD11c or CD19 was detected. **(B)** Liver function tests remained within the normal range in mice transduced with AAV-HC-$K^d$-YCAC (here shown on d7 post-inoculation, n = 3). **(A, B)** Mean±SEM are shown. Other expression data for this vector are shown in Figure 4. **(C)** Expression of H-2$K^d$ persisted in transduced livers through to at least d100 following B6.Kd skin transplantation in all vector/strain combinations (scale bar = 200 μm, representative images from n = 6).

**Supplementary Figure 8. Validation of the identity of eluted peptides.**

The identity of a panel of eluted peptides was validated by comparing chromatographic retention and MS/MS spectra with those of the corresponding synthetic peptides. (**A**) Representative spectra for three pairs of synthetic and eluted peptides. (**B**) Pearson correlation coefficients (*r*) between the $\log_{10}$ intensities of identified b- and y-ions in the synthetic and sample-derived spectra are shown. Error bars represent the 95% confidence intervals. The corresponding p-value was $< 0.05$ for each peptide pair.

**Supplementary Figure 9. Tetramer Staining of alloreactive T cell populations.**

(**A**) Gating strategy for identification of alloreactive T cells using a 5-tetramer panel. Here, $CD4^+$ T cells are used as a specificity control for $CD8^+$ T cell staining. The proportion of $CD8^+$ and $CD4^+$ T cells staining with the tetramer panel is shown for (**B**) combined secondary lymphoid organs, (**C**) liver leukocytes and (**D**) skin graft-infiltrating cells on the protocol days indicated. Data from experiments with a total of n = 3 biological replicates per group are shown in **(B-D)**. Data are presented as mean ± SEM.

**Supplementary Figure 10. CD8$^+$ T cell subsets of liver leukocytes and combined secondary lymphoid organs.**

(**A**) Rejection of a primary or secondary skin graft is accompanied by the loss of naïve CD8$^+$ tetramer-positive cells within the liver leukocyte population, and a shift of the majority of CD44$^+$ cells from CD62L$^+$ to CD62L$^-$. Inoculation of primed mice with AAV-K$^b$ results in almost total loss of CD62L$^-$ cells. (**B**) Similar trends are observed in the CD8$^+$tet$^+$ cells from combined secondary lymphoid organs, but in this case there is never a complete loss of naïve or antigen-experienced CD62L$^+$ cells. The secondary lymphoid organs were pooled in order to estimate changes in the total number of CD8$^+$tet$^+$ cells under different transplant conditions; inclusion of both draining and non-draining lymph node groups for the liver and skin grafts means that some CD8$^+$tet$^+$ T cells which do not recirculate to/from these sites are mixed with the recirculating cells. For both the liver leukocytes and the SLOs, changes in the phenotype of CD8$^+$tet$^+$ cells were partially or completely obscured within the bulk CD8$^+$ population. Expression of KLRG1, CD69 and CXCR6 was determined for the CD44$^+$CD62L$^-$ cells from the liver (**C**) and SLOs (**D**). (**A-B**) representative flow plots from experiments with a total of n = 3 biological replicates per group. (**C-D**) Data from experiments with a total of n = 3 biological replicates per group are shown. Data are presented as mean ± SEM.

**Supplementary Figure 11.**

**Expression of coinhibitory receptors by bulk and tetramer-positive CD8$^+$ T cells from liver or combined secondary lymphoid organs.**

Expression of PD-1, TIGIT, Tim-3 and LAG-3 was determined for tet$^+$ and bulk CD8$^+$ T cells in a model of secondary skin graft rejection or tolerance. Modest upregulation of PD-1 was noted in the CD8$^+$tet$^+$ liver leukocytes (**A**) and SLOs (**B**) at all intervals following graft rejection. In contrast, induction of tolerance upon inoculation of recipient mice with AAV-K$^b$ was accompanied by strong expression of all coinhibitory ligands, with expression of LAG-3 and Tim-3 declining to baseline by protocol d84. Changes in the phenotype of $CD8^+tet^+$ cells were much less obvious within the bulk $CD8^+$ populations. (**A-B**) representative flow plots from experiments with a total of n = 3 biological replicates per group.

**Supplementary Data 1. Complete list of identified peptides**

**Supplementary Table 2. Common subset of peptides presented by H-2K$^d$**

1083 $K^d$-binding peptides were identified in either or both replicate samples from C57BL/6 hepatocytes transduced with AAV-HC-$K^d$, B6.Kd donor skin grafts or B6.Kd donor spleen. Features of the common peptides are listed.

**Supplementary Table 3. Common subset of peptides presented by H-2K$^b$**

880 $K^b$-binding peptides were identified in either or both replicate samples from B10.BR hepatocytes transduced with AAV-HC-$K^b$, 178.3 donor skin grafts or 178.3 donor spleen. Features of the common peptides are listed.

**Supplementary Table 4. Tetramer screening of alloreactive CD8$^+$ T cells**

100 peptides were selected for evaluation of immunogenicity by pMHC tetramer staining of activated alloreactive $CD8^+$ T cells. Peptide characteristics and screening results in three different responder populations are shown. Values given for peptide screening results are the mean and the range (min-max). Peptides recognised by > 5% of activated ($CD44^+PD\text{-}1^{hi}$) $CD8^+$ T cells are indicated in red.

**Supplementary Table 5. List of Antibodies used in this study**

**Supplementary Table 6. Variable Window widths used for DIA acquisition**

**Supplementary References**

1. Honjo K, Yan Xu X, Kapp JA, and Bucy RP. Evidence for cooperativity in the rejection of cardiac grafts mediated by CD4 TCR Tg T cells specific for a defined allopeptide. *Am J Transplant.* 2004;4(11):1762-8.
2. Friedel RH, Seisenberger C, Kaloff C, and Wurst W. EUCOMM--the European conditional mouse mutagenesis program. *Brief Funct Genomic Proteomic.* 2007;6(3):180-5.
3. Raymond CS, and Soriano P. High-efficiency FLP and PhiC31 site-specific recombination in mammalian cells. *PLoS One.* 2007;2(1):e162.
4. Yakar S, Liu JL, Stannard B, Butler A, Accili D, Sauer B, et al. Normal growth and development in the absence of hepatic insulin-like growth factor I. *Proc Natl Acad Sci U S A.* 1999;96(13):7324-9.

Supplementary Figure 1

**SCT - single chain trimer**

**Leader sequence----peptide----linker1(2C)----β2m----linker2----heavy chain H-2K$^b$/H-2K$^d$ (Y84C)**

SCT-K$^b$-SIINFEKL
(amino acid)
MARSVTLVFLVLVSLTGLYASIINFEKLGCGASGGGGSGGGGSIQKTPQIQVYSRHPPENGKPNILNCYVTQFHPP
HIEIQMLKNGKKIPKVEMSDMSFSKDWSFYILAHTEFTPTETDTYACRVKHASMAEPKTVYWDRDMGGGGSGGG
GSGGGGSGGGGSGPHSLRYFVTAVSRPGLGEPRYMEVGYVDDTEFVRFDSDAENPRYEPRARWMEQEGPEY
WERETQKAKGNEQSFRVDLRTLLGCYNQSKGGSHTIQVISGCEVGSDGRLLRGYQQYAYDGCDYIALNEDLKT
WTAADMAALITKHKWEQAGEAERLRAYLEGTCVEWLRRYLKNGNATLLRTDSPKAHVTHHSRPEDKVTLRCWAL
GFYPADITLTWQLNGEELIQDMELVETRPAGDGTFQKWASVVVPLGKEQYYTCHVYHQGLPEPLTLRWEPPPST
VSNMATVAVLVVLGAAIVTGAVVAFVMKMRRRNTGGKGGDYALAPGSQTSDLSLPDCKVMVHDPHSLA

SCT-K$^b$-KIITYRNL
(amino acid)
MARSVTLVFLVLVSLTGLYAKIITYRNLGCGASGGGGSGGGGSIQKTPQIQVYSRHPPENGKPNILNCYVTQFHPP
HIEIQMLKNGKKIPKVEMSDMSFSKDWSFYILAHTEFTPTETDTYACRVKHASMAEPKTVYWDRDMGGGGSGGG
GSGGGGSGGGGSGPHSLRYFVTAVSRPGLGEPRYMEVGYVDDTEFVRFDSDAENPRYEPRARWMEQEGPEY
WERETQKAKGNEQSFRVDLRTLLGCYNQSKGGSHTIQVISGCEVGSDGRLLRGYQQYAYDGCDYIALNEDLKT
WTAADMAALITKHKWEQAGEAERLRAYLEGTCVEWLRRYLKNGNATLLRTDSPKAHVTHHSRPEDKVTLRCWAL
GFYPADITLTWQLNGEELIQDMELVETRPAGDGTFQKWASVVVPLGKEQYYTCHVYHQGLPEPLTLRWEPPPST
VSNMATVAVLVVLGAAIVTGAVVAFVMKMRRRNTGGKGGDYALAPGSQTSDLSLPDCKVMVHDPHSLA

SCT-K$^d$-SYFPEITHI
(amino acid)
MAPCTLLLLLAAALAPTQTRASYFPEITHIGCGASGGGGSGGGGSIQKTPQIQVYSRHPPENGKPNILNCYVTQFH
PPHIEIQMLKNGKKIPKVEMSDMSFSKDWSFYILAHTEFTPTETDTYACRVKHASMAEPKTVYWDRDMGGGGSG
GGGSGGGGSGGGGSGPHSLRYFVTAVSRPGLGEPRFIAVGYVDDTQFVRFDSDADNPRFEPRAPWMEQEGPE
YWEEQTQRAKSDEQWFRVSLRTAQRCYNQSKGGSHTFQRMFGCDVGSDWRLLRGYQQFAYDGRDYIALNEDL
KTWTAADTAALITRRKWEQAGDAEYYRAYLEGECVEWLRRYLELGNETLLRTDSPKAHVTYHPRSQVDVTLRCW
ALGFYPADITLTWQLNGEDLTQDMELVETRPAGDGTFQKWAAVVVPLGKEQNYTCHVHHKGLPEPLTLRWKLPP
STVSNTVIIAVLVVLGAAIVTGAVVAFVMKMRRNTGGKGVNYALAPGSQTSDLSLPDGKVMVHDPHSLA

**MHC (+YCAC mutation)**

**Leader sequence----heavy chain H-2K$^d$ (Y84C, A139C)**

HC-K$^d$-YCAC
(amino acid)
MAPCTLLLLLAAALAPTQTRAGPHSLRYFVTAVSRPGLGEPRFIAVGYVDDTQFVRFDSDADNPRFEPRAPWME
QEGPEYWEEQTQRAKSDEQWFRVSLRTAQRCYNQSKGGSHTFQRMFGCDVGSDWRLLRGYQQFAYDGRDYI
ALNEDLKTWTAADTCALITRRKWEQAGDAEYYRAYLEGECVEWLRRYLELGNETLLRTDSPKAHVTYHPRSQVD
VTLRCWALGFYPADITLTWQLNGEDLTQDMELVETRPAGDGTFQKWAAVVVPLGKEQNYTCHVHHKGLPEPLTL
RWKLPPSTVSNTVIIAVLVVLGAAIVTGAVVAFVMKMRRNTGGKGVNYALAPGSQTSDLSLPDGKVMVHDPHSLA

**Supplementary Figure 1.**
**Amino acid sequences for the SCT-K$^b$-SIIN, SCT-K$^b$-KIIT, SCT-K$^d$-SYFP and HC-K$^d$-YCAC constructs.**

Supplementary Figure 2

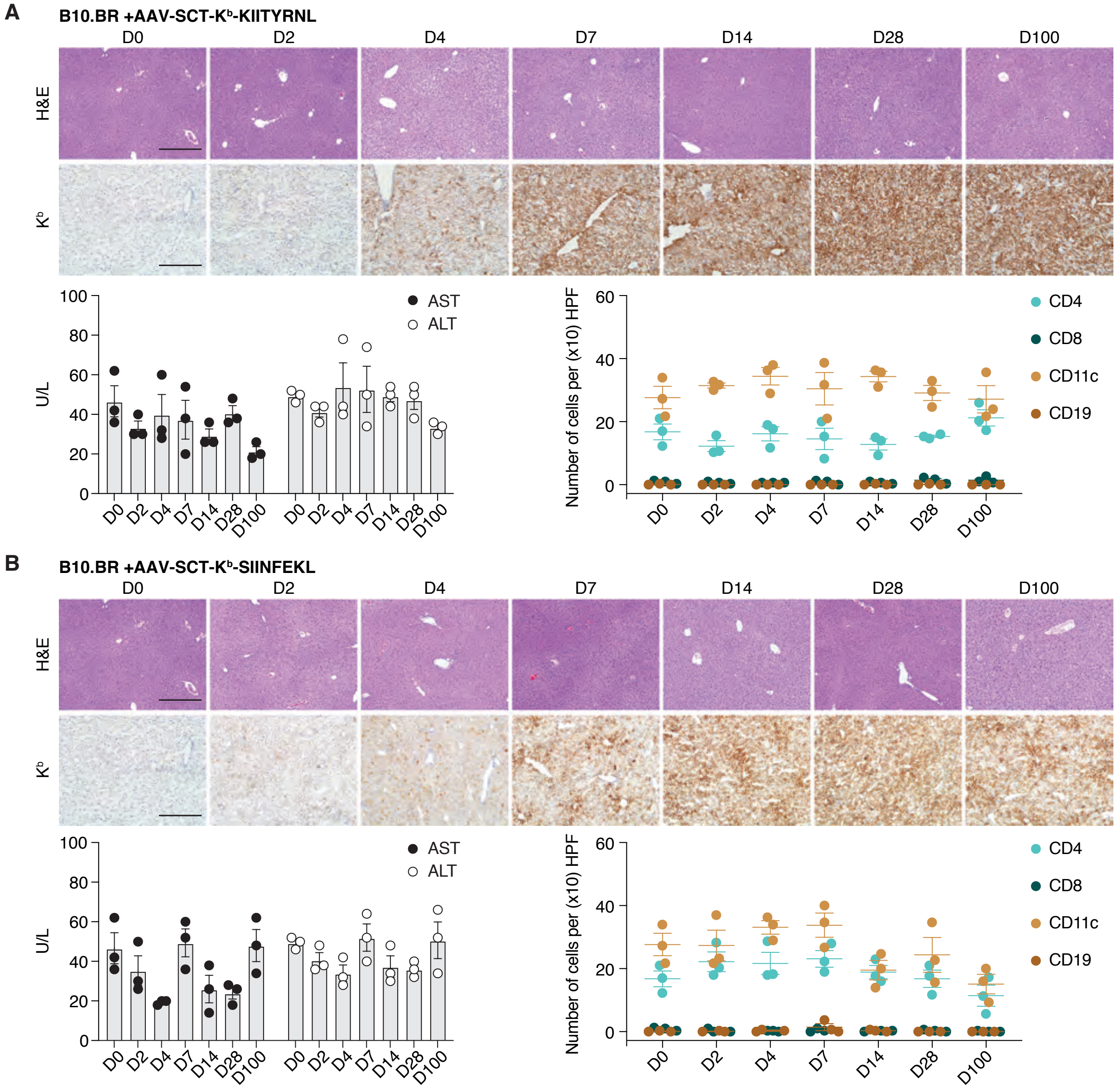

**Supplementary Figure 2.**

**Expression of SCT-K$^b$-peptide in mouse hepatocytes is robust and persistent.**

B10.BR mice were inoculated with AAV-SCT-K$^b$-KIITYRNL **(A)** or AAV-SCT-K$^b$-SIINFEKL **(B)**. On days 2-100 post-inoculation, tissues were collected for analysis (n = 3 /interval). Representative immunostained (IHC) and H&E images show transduced liver sections (scale bar: 200 μm). Robust expression of H2-K$^b$ was present through day 100 post-inoculation. Histologic examination of the liver sections was normal. Levels of aspartate aminotransferase (AST) and alanine aminotransferase (ALT) did not increase significantly from baseline in mice treated with AAV-SCT-K$^b$-KIITYRNL (one-way ANOVA, p = 0.4 for AST and p = 0.21 for ALT) or in mice receiving AAV-SCT-K$^b$-SIINFEKL (one-way ANOVA, p = 0.13 for AST and p = 0.02 for ALT, due solely to a decrease in ALT on d4). Minimal infiltration with cells expressing the markers CD4, CD8, CD11c or CD19 was detected. Mean ± SEM are shown.

Supplementary Figure 3

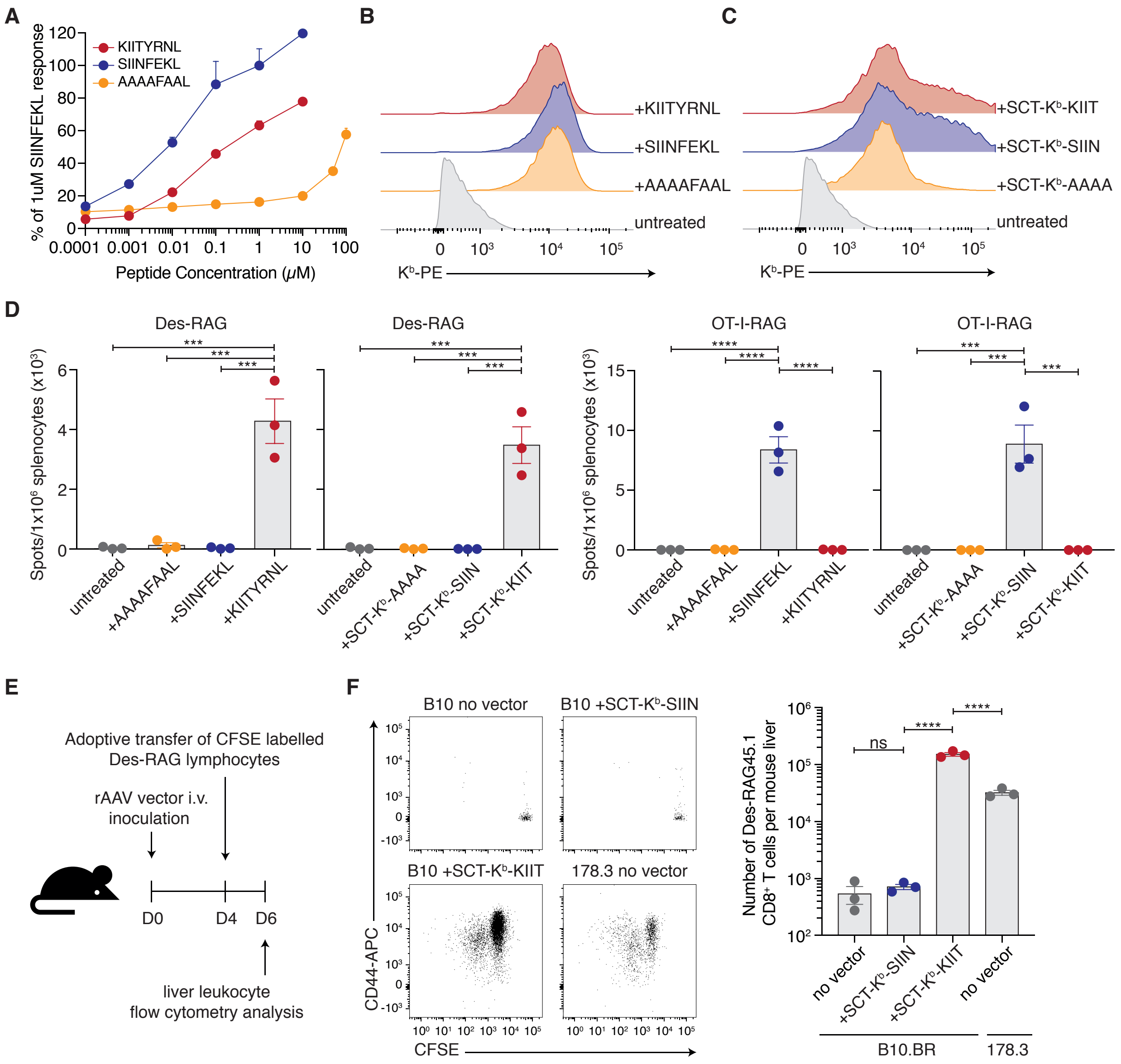

**Supplementary Figure 3.**
**Recognition of SCT peptide-MHC ligands in vitro and in vivo.**
**(A)** RMA-S cells were pulsed with different concentrations of the peptides KIITYRNL (Pcid2$_{318\text{-}325}$), SIINFEKL (OVA$_{257\text{-}264}$) or AAAAFAAL (synthetic negative control), or were untreated. Stabilisation of H-2K$^b$ surface expression was assessed by flow cytometry following staining with a conformation-dependent anti-H-2K$^b$ mAb (clone Y3). **(B)** Flow plots shown are representative of three independent experiments. Peptide concentrations required to achieve equivalent H-2K$^b$ surface expression levels were determined. **(C)** RMA-S cells were transiently transfected with constructs encoding SCT-K$^b$-KIIT, SCT-K$^b$-SIIN and SCT-K$^b$-AAAA using a Lonza-AMAXA Nucleofector 2b. Transgene expression was assessed by flow cytometry (as above) 24 hours after transfection. Flow plots shown are representative of three independent experiments. **(D)** The proportion of cells secreting IFN-γ upon recognition of their cognate antigen was determined using ELISPOT assays. Splenocytes from Des-RAG or OT-I-RAG mice were cultured with irradiated stimulators; RMA-S pulsed with selected peptides or expressing SCT constructs after transient transfection. SCT recognition by cognate TCRs mirrored recognition of the native H-2K$^b$-peptide complex. SCT constructs were recognised in a peptide-specific manner in vitro. Data from two independent experiments with a total of n = 3 biological replicates per group are shown. **(E)** Des-RAG lymphocytes were labelled with CFSE, adoptively transferred into recipient mice and recovered from the recipient liver two days later. Some recipient mice were treated with AAV encoding SCT-K$^b$-KIIT or SCT-K$^b$-SIIN prior to adoptive transfer, as shown. **(F)** Flow cytometry analysis of CFSE-labelled Des-RAG lymphocytes demonstrates peptide-specific activation and proliferation of adoptively transferred CD8$^+$ Des-RAG T cells upon encounter with their cognate antigen in the liver, confirming that recognition of the SCT-K$^b$-KIIT ligand in vivo was analogous to that of the native pMHC complex. Data from three independent experiments with a total of n = 3 biological replicates per group are shown. **(D, F)** Mean ± SEM are shown, one-way ANOVA in conjunction with Sidak's multiple comparison test: ns, not significant; *p < 0.05, ** p < 0.01, *** p < 0.001, **** p < 0.0001.

Supplementary Figure 4

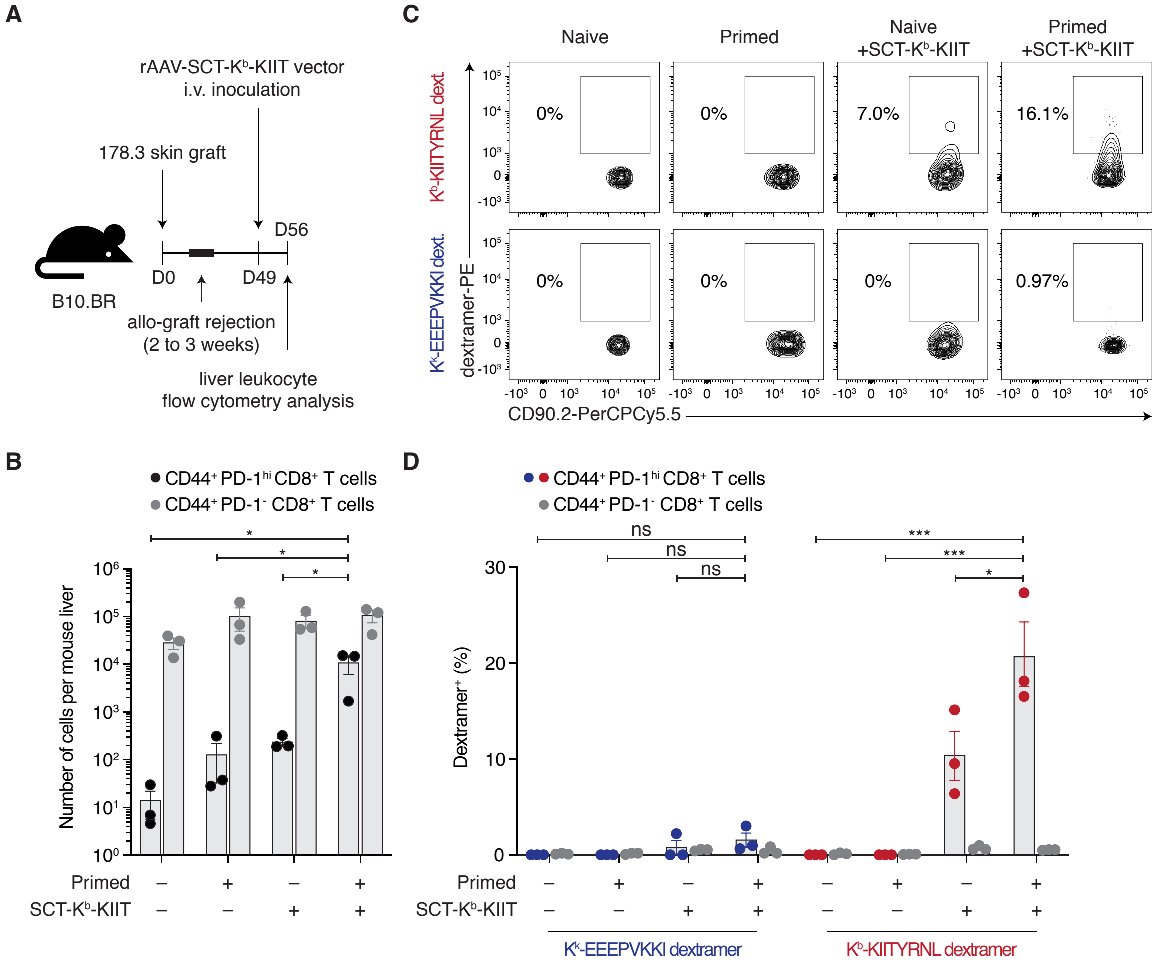

**Supplementary Figure 4.**

**Recognition of SCT-K$^b$-KIIT in a polyclonal alloreactive population.**

**(A)** Inoculation with SCT-K$^b$-KIIT vector not only activates a clone of transgenic Des-RAG T cells bearing the cognate receptor, but also activates a proportion of the polyclonal T cell repertoire of normal B10.BR mice. B10.BR mice were primed against allogeneic H-2K$^b$ (178.3 skin graft). Approximately 30 days post-graft rejection, some of the primed or naïve B10.BR mice were inoculated with AAV-SCT-K$^b$-KIIT. Liver leukocytes were analysed on day 7 post-inoculation. **(B)** Activated CD8$^+$ T cells, defined as CD44$^+$PD-1$^{hi}$, increased in number following priming or transduction with SCT-K$^b$-KIIT, with a further increase in primed mice receiving SCT-K$^b$-KIIT. **(C-D)** Inoculation of naïve or primed B10.BR mice with AAV-SCT-K$^b$-KIIT generated populations of activated (CD44$^+$PD-1$^{hi}$) CD8$^+$ T cells which bound K$^b$-KIITYRNL dextramers specifically. Dextramers of the syngeneic pMHC K$^k$-EEEPVKKI were used as negative controls. Data from one representative experiment (from n = 3) is shown in **(C)**, while two independent experiments with a total of n = 3 biological replicates per group are shown in **(B, D)**. Data are presented as mean ± SEM, statistical analysis involved two-way analysis of variance (ANOVA) in conjunction with Tukey's multiple comparison test, *$p < 0.05$, ** $p < 0.01$, *** $p < 0.001$, **** $p < 0.0001$).

Supplementary Figure 5

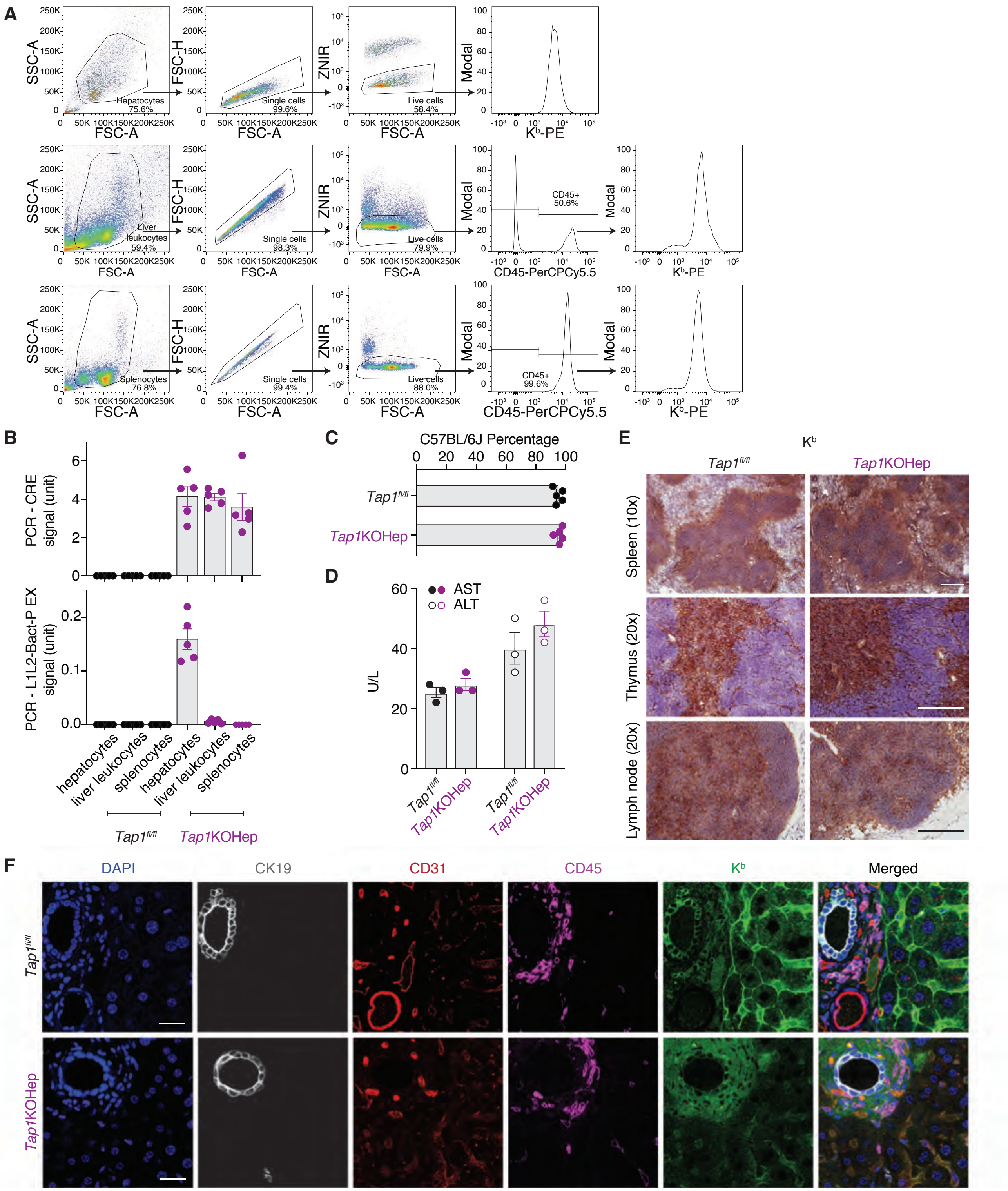

**Supplementary Figure 5.**
**Characterisation of *Tap1*KOHep mice.**
**(A)** Gating strategy for determination of Kᵇ expression on the surface of hepatocytes, liver leukocytes and splenocytes of *Tap1*KOHep and *Tap1*$^{fl/fl}$ mice. **(B)** Genotyping PCR performed by Transnetyx shows the presence of the Albumin-Cre transgene in hepatocytes, liver leukocytes and spleen from *Tap1*KOHep mice (above). Because Cre activity is restricted to hepatocytes, the recombined *Tap1* sequence specifically detected by the L1L2-Bact-P EX probe is only amplified in hepatocytes and not other tissues from *Tap1*KOHep (below). Data from one experiment with a total of n = 5 biological replicates per group are shown. **(C)** Genetic background analysis was undertaken by Transnetyx. *Tap1*KOHep and *Tap1*$^{fl/fl}$ control mice were at least 91.3% C57BL/6J (91.3-97.9%). Data from one experiment with a total of n=5 biological replicates per group are shown. **(D)** AST and ALT levels are comparable between *Tap1*KOHep and floxed littermate control mice (n = 3). **(B-D)** Mean ± SEM are shown. **(E)** H-2Kᵇ is expressed at normal levels in the spleen, thymus and lymph node of *Tap1*KOHep and *Tap1*$^{fl/fl}$ mice (scale bar = 100$\mu$m, representative images from n = 3). **(F)** H-2Kᵇ is absent from the hepatocytes of *Tap1*KOHep mice, but detectable on other liver cells (scale bar = 40$\mu$m, representative images from n = 3).

Supplementary Figure 6

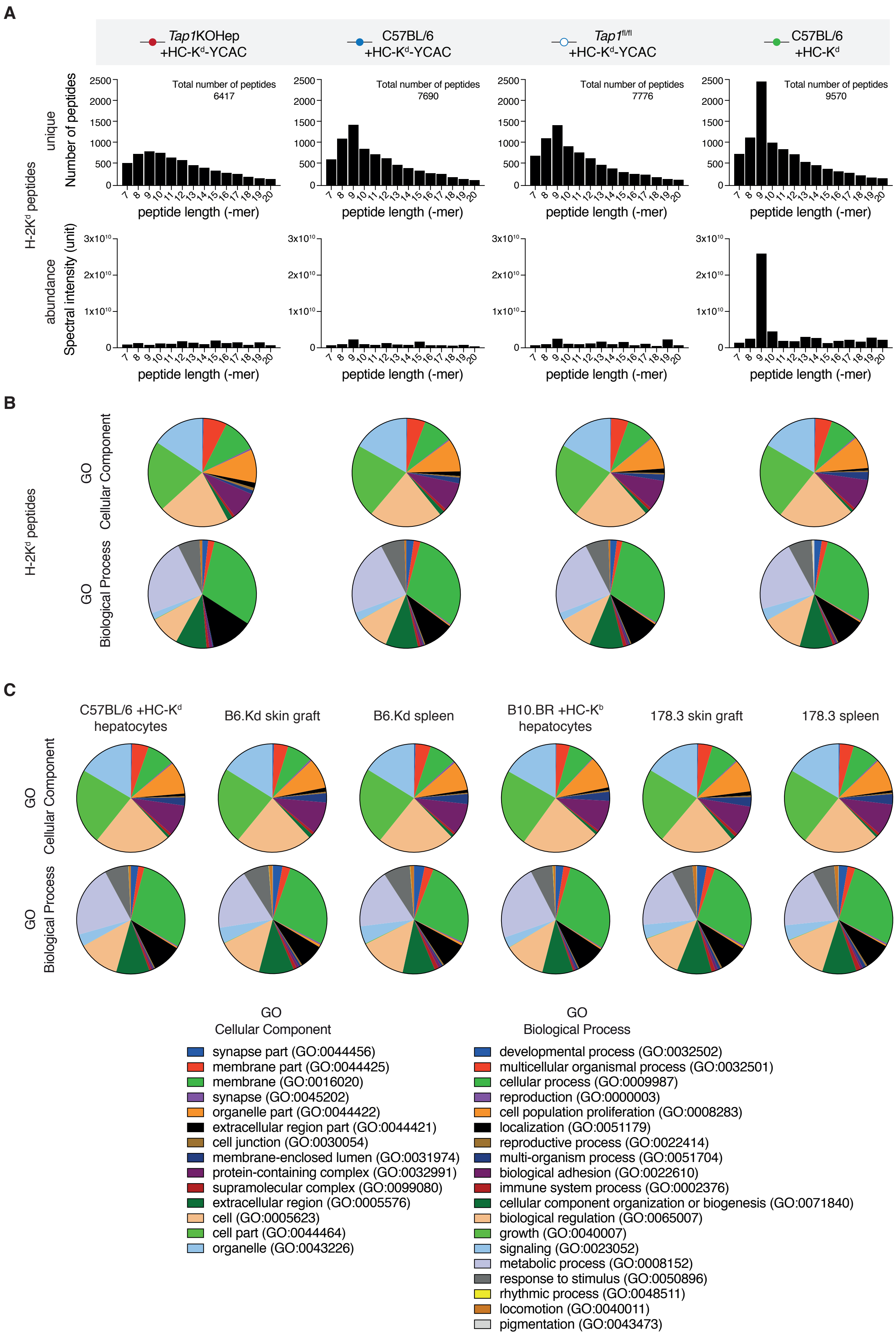

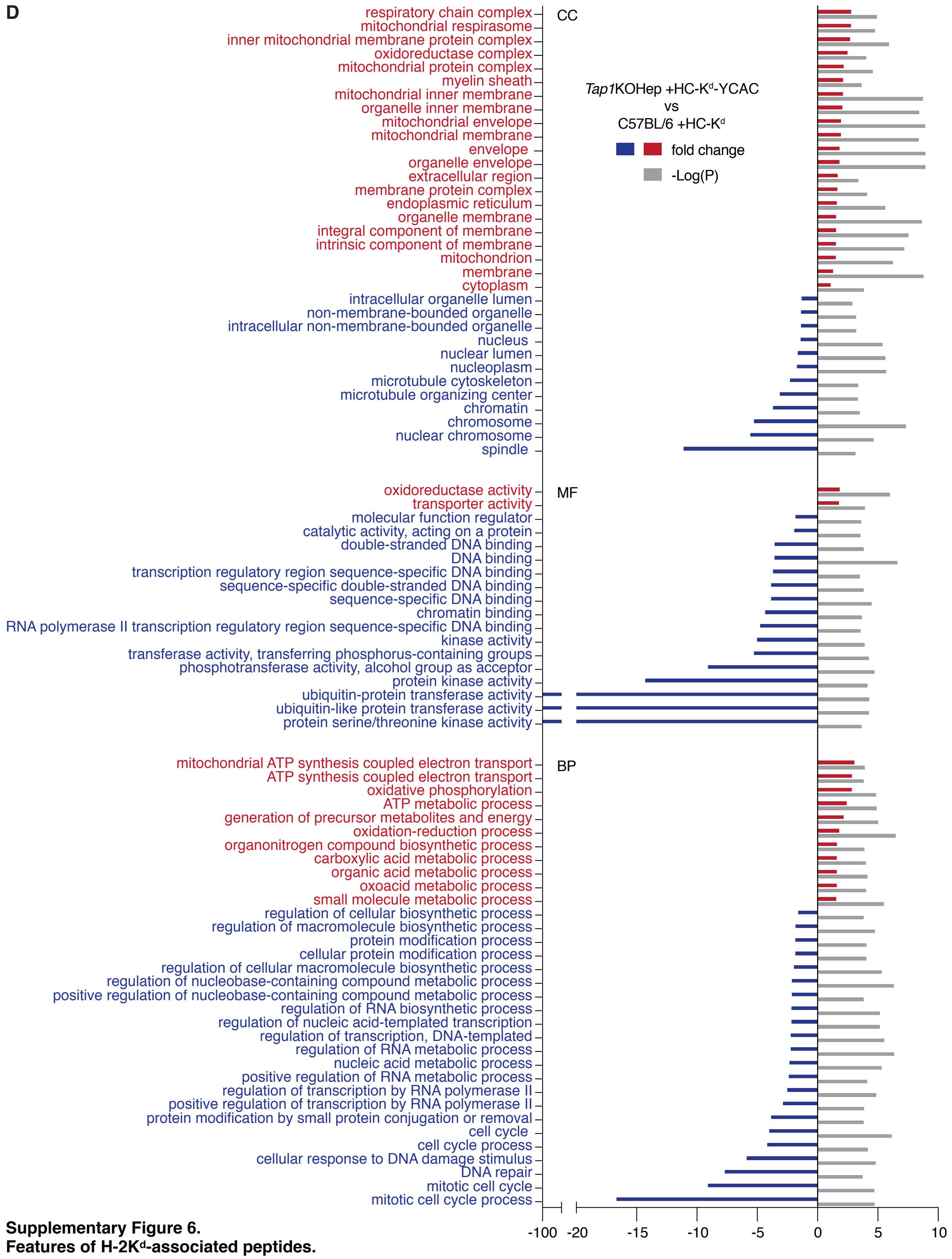

**Supplementary Figure 6.**
**Features of H-2K$^d$-associated peptides.**

The length distribution for H-2K$^d$-associated peptides eluted from transduced hepatocytes in each of four vector/strain combinations is shown in panel **(A)**. Peptides eluted from C57BL/6 mice expressing K$^d$-HC are predominantly nonamers – this preference was less strong for the peptide repertoires of hepatocytes expressing K$^d$-YCAC. **(B-D)** Gene Ontology annotations of the source proteins associated with eluted peptides were analysed using the PANTHER classification system. Function classification analysis and statistical over-representation tests were performed. **(B)** Cellular component and biological process analysis of source proteins corresponding to the same hepatocyte peptide repertoires shown in **(A)**. **(C)** Analysis of the source proteins giving rise to the H-2K$^d$ and K$^b$-associated peptide repertoires of transduced hepatocytes, donor skin grafts and donor spleen. **(D)** A number of Gene Ontology terms were enriched or depleted when hepatocyte source proteins from AAV-HC-K$^d$-YCAC-transduced *Tap1*KOHep mice were compared with those from AAV-HC-K$^d$-treated C57BL/6. The most striking enrichment was in terms associated with mitochondria and mitochondrial metabolism. Significant enrichment was also found for the cellular component terms endoplasmic reticulum, extracellular region and cytoplasm.

Supplementary Figure 7

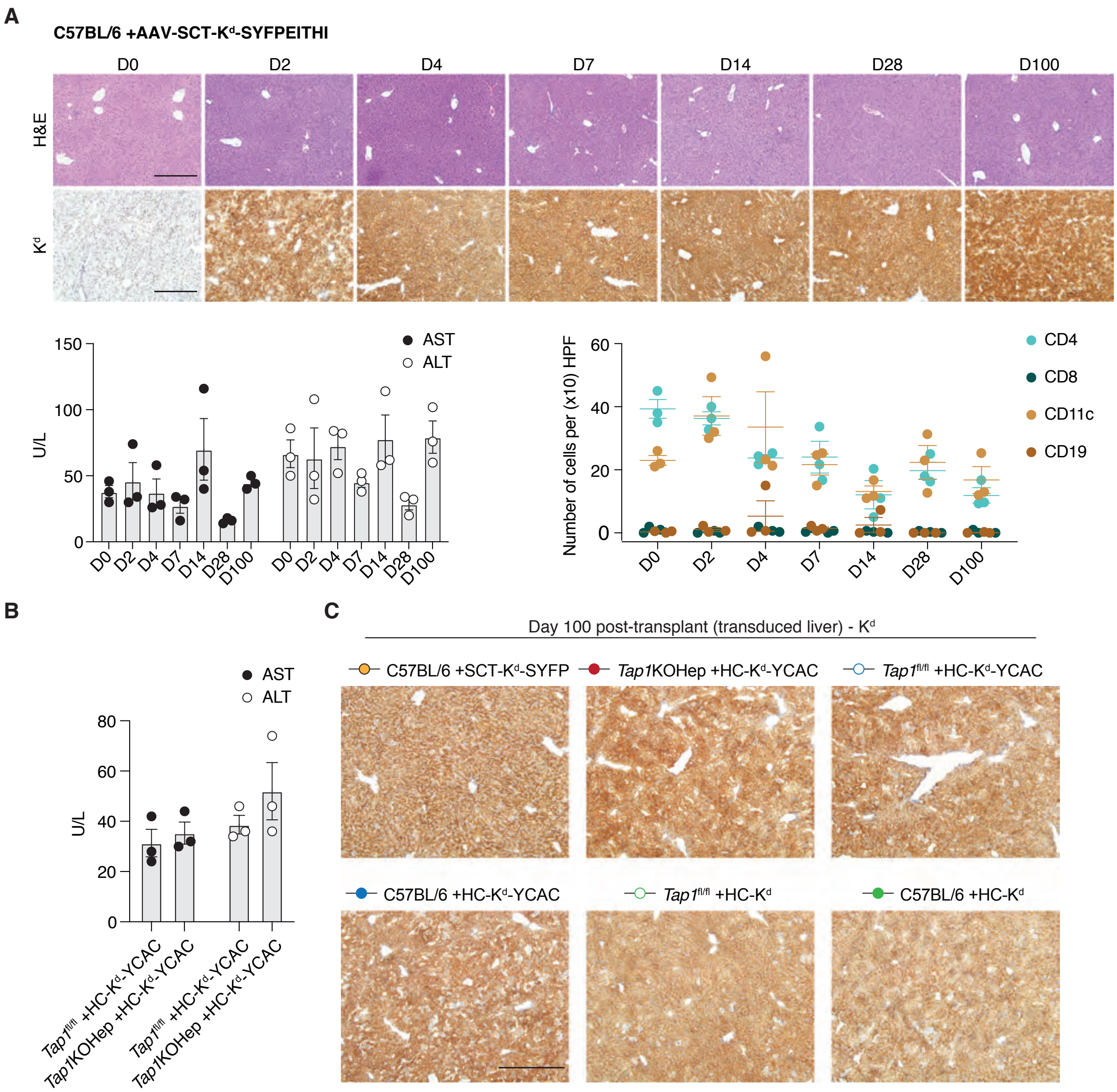

**Supplementary Figure 7.**
**Expression of AAV-SCT-K$^d$-SYFPEITHI and AAV-HC-K$^d$-YCAC is strong and durable.**
**(A)** C57BL/6 mice were inoculated with AAV-SCT-K$^d$-SYFPEITHI iv. On days 2, 4, 7, 14, 28 and 100 post-inoculation, tissues were collected for analysis (n = 3 at each interval). Representative IHC and H&E images show transduced liver sections (scale bar: 200 µm). Robust expression of H-2K$^d$ was present through day 100 post-inoculation. Histologic examination of the liver sections was normal. Levels of AST and ALT did not increase significantly from baseline (one-way ANOVA, p = 0.14 for AST and p = 0.11 for ALT in mice inoculated with AAV-SCT-K$^d$-SYFPEITHI). Minimal infiltration with cells expressing the markers CD4, CD8, CD11c or CD19 was detected. **(B)** Liver function tests remained within the normal range in mice transduced with AAV-HC-K$^d$-YCAC (here shown on d7 post-inoculation, n = 3). **(A, B)** Mean ± SEM are shown. Other expression data for this vector are shown in Figure 4. **(C)** Expression of H-2K$^d$ persisted in transduced livers through to at least d100 following B6.Kd skin transplantation in all vector/strain combinations (scale bar: 200 µm, representative images from n = 6).

Supplementary Figure 8

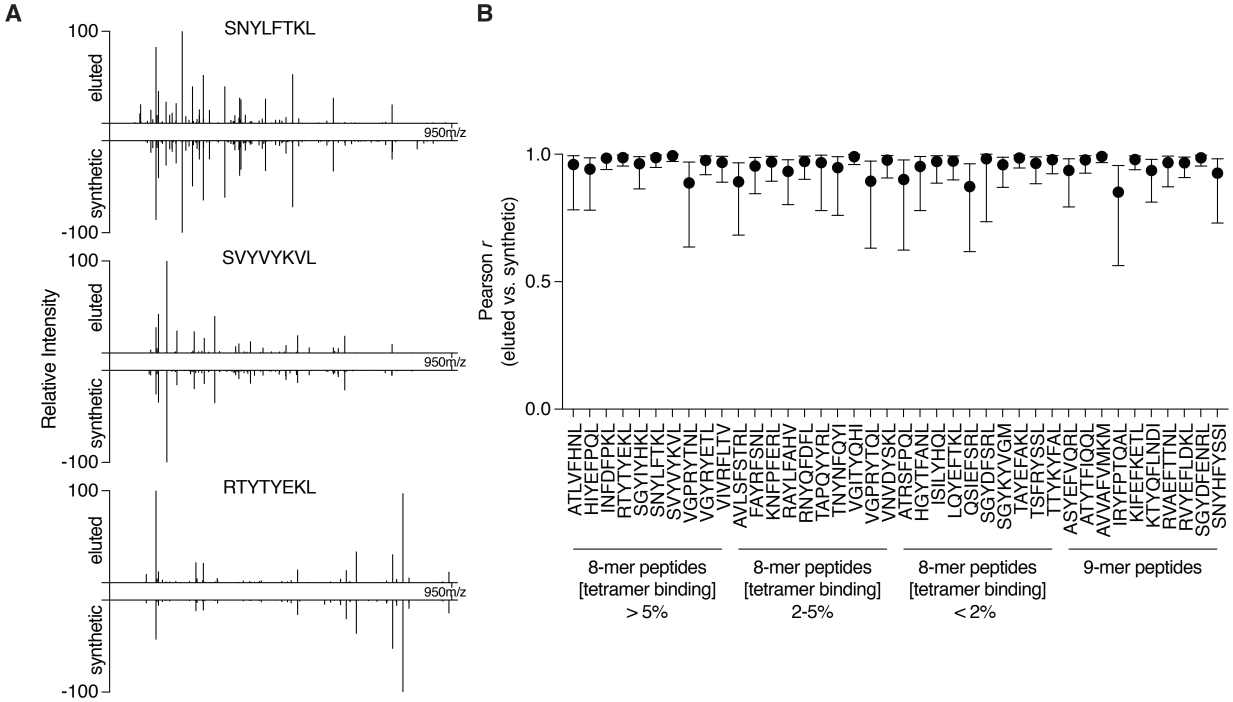

**Supplementary Figure 8.**
**Validation of the identity of eluted peptides.**
The identity of a panel of eluted peptides was validated by comparing chromatographic retention and MS/MS spectra with those of the corresponding synthetic peptides. (**A**) Representative spectra for three pairs of synthetic and eluted peptides. (**B**) Pearson correlation coefficients (*r*) between the $log_{10}$ intensities of identified b- and y-ions in the synthetic and sample-derived spectra are shown. Error bars represent the 95% confidence intervals. The corresponding p-value was $< 0.05$ for each peptide pair.

Supplementary Figure 9

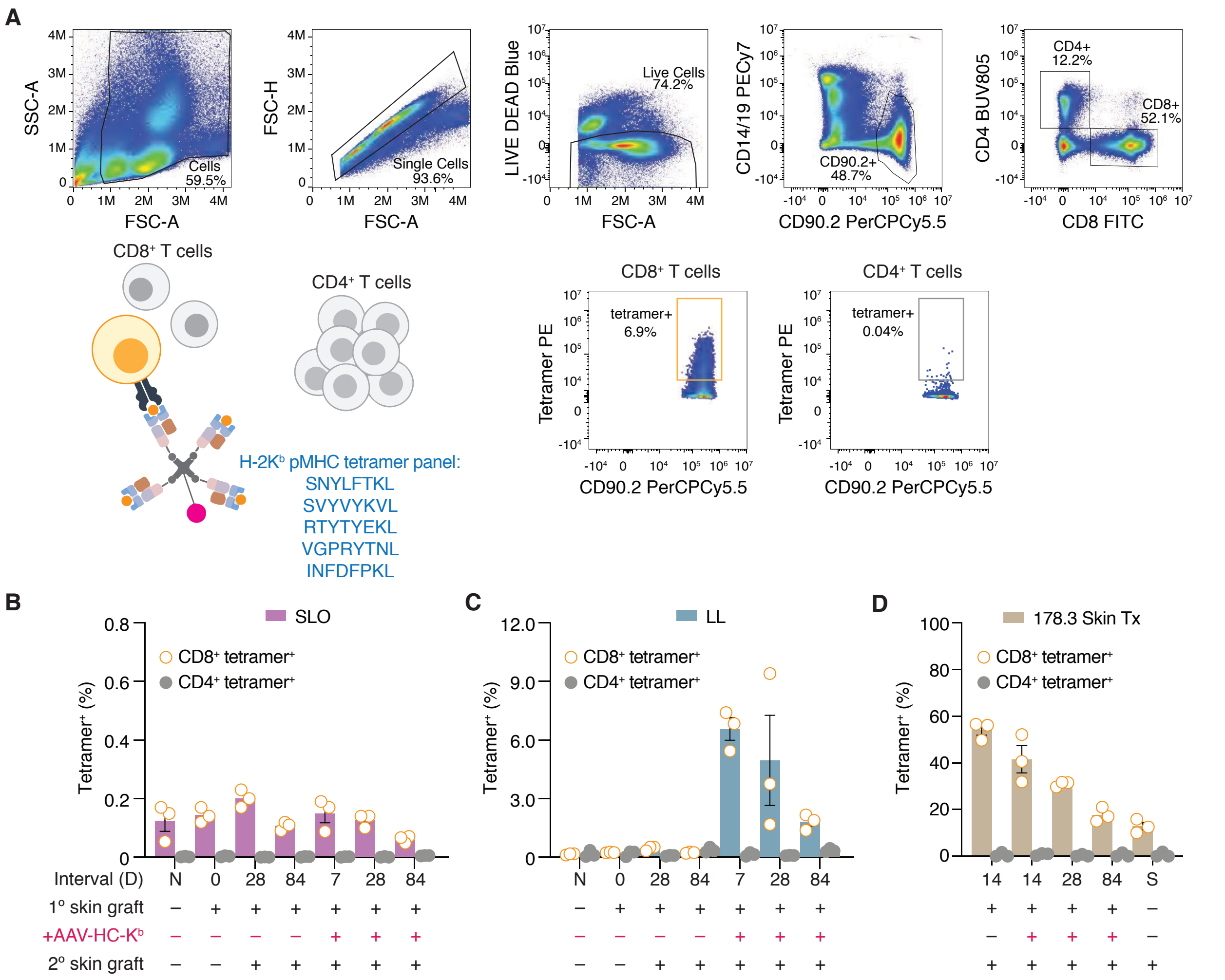

**Supplementary Figure 9.**
**Tetramer Staining of alloreactive T cell populations.**
(**A**) Gating strategy for identification of alloreactive T cells using a 5-tetramer panel. Here, CD4+ T cells are used as a specificity control for CD8+ T cell staining. The proportion of CD8+ and CD4+ T cells staining with the tetramer panel is shown for (**B**) combined secondary lymphoid organs, (**C**) liver leukocytes and (**D**) skin graft-infiltrating cells on the protocol days indicated. Data from experiments with a total of n = 3 biological replicates per group are shown in **(B-D)**. Data are presented as mean ± SEM.

Supplementary Figure 10

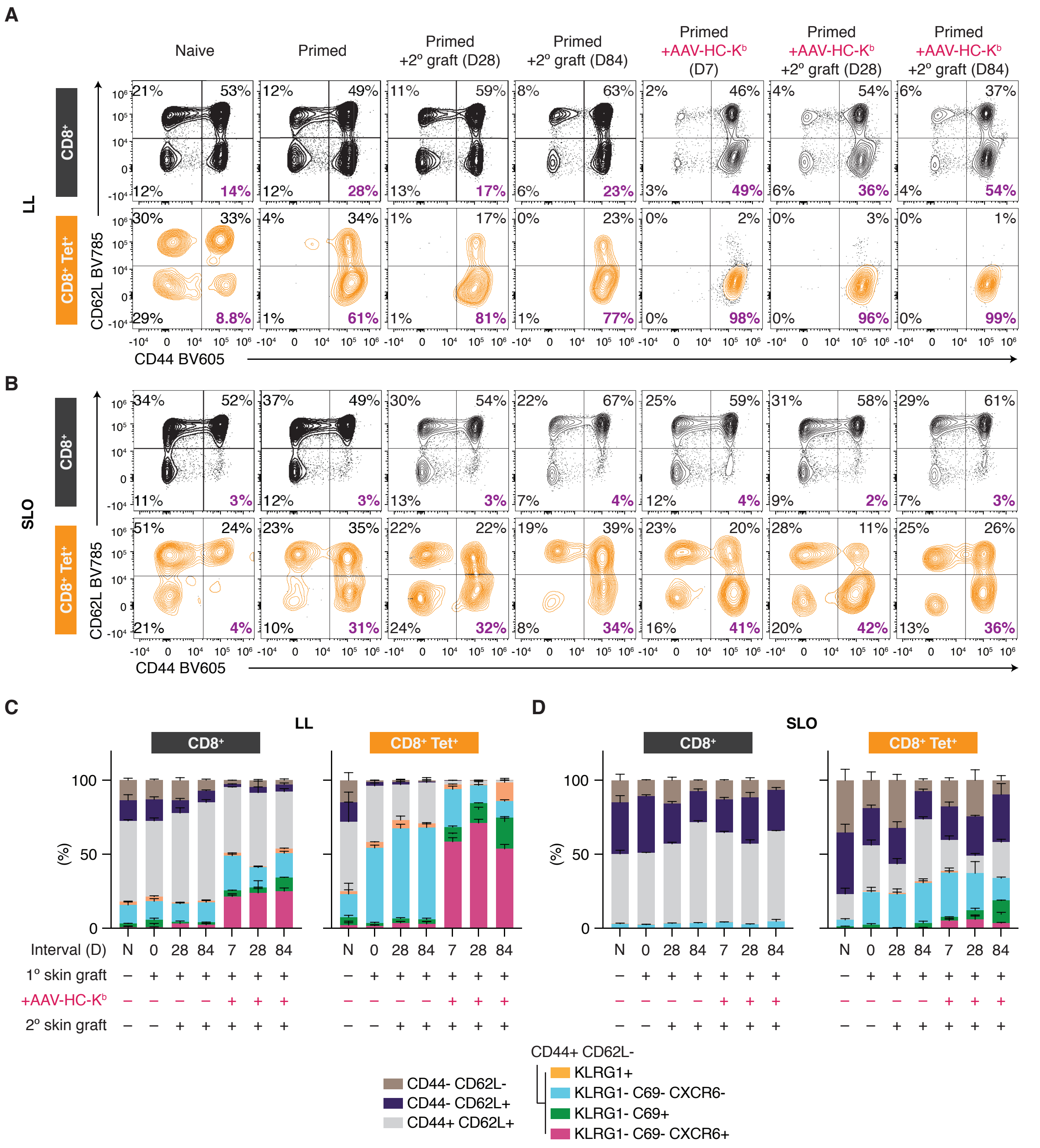

**Supplementary Figure 10.**
**CD8⁺ T cell subsets of liver leukocytes and combined secondary lymphoid organs.**
(**A**) Rejection of a primary or secondary skin graft is accompanied by the loss of naïve CD8⁺ tetramer-positive cells within the liver leukocyte population, and a shift of the majority of CD44⁺ cells from CD62L⁺ to CD62L⁻. Inoculation of primed mice with AAV-Kᵇ results in almost total loss of CD62L⁻ cells. (**B**) Similar trends are observed in the CD8⁺tet⁺ cells from combined secondary lymphoid organs, but in this case there is never a complete loss of naïve or antigen-experienced CD62L⁺ cells. The secondary lymphoid organs were pooled in order to estimate changes in the total number of CD8⁺tet⁺ cells under different transplant conditions; inclusion of both draining and non-draining lymph node groups for the liver and skin grafts means that some CD8⁺tet⁺ T cells which do not recirculate to/from these sites are mixed with the recirculating cells. For both the liver leukocytes and the SLOs, changes in the phenotype of CD8⁺tet⁺ cells were partially or completely obscured within the bulk CD8⁺ population. Expression of KLRG1, CD69 and CXCR6 was determined for the CD44⁺CD62L⁻ cells from the liver (**C**) and SLOs (**D**). (**A-B**) representative flow plots from experiments with a total of n = 3 biological replicates per group. (**C-D**) Data from experiments with a total of n = 3 biological replicates per group are shown. Data are presented as mean ± SEM.

Supplementary Figure 11

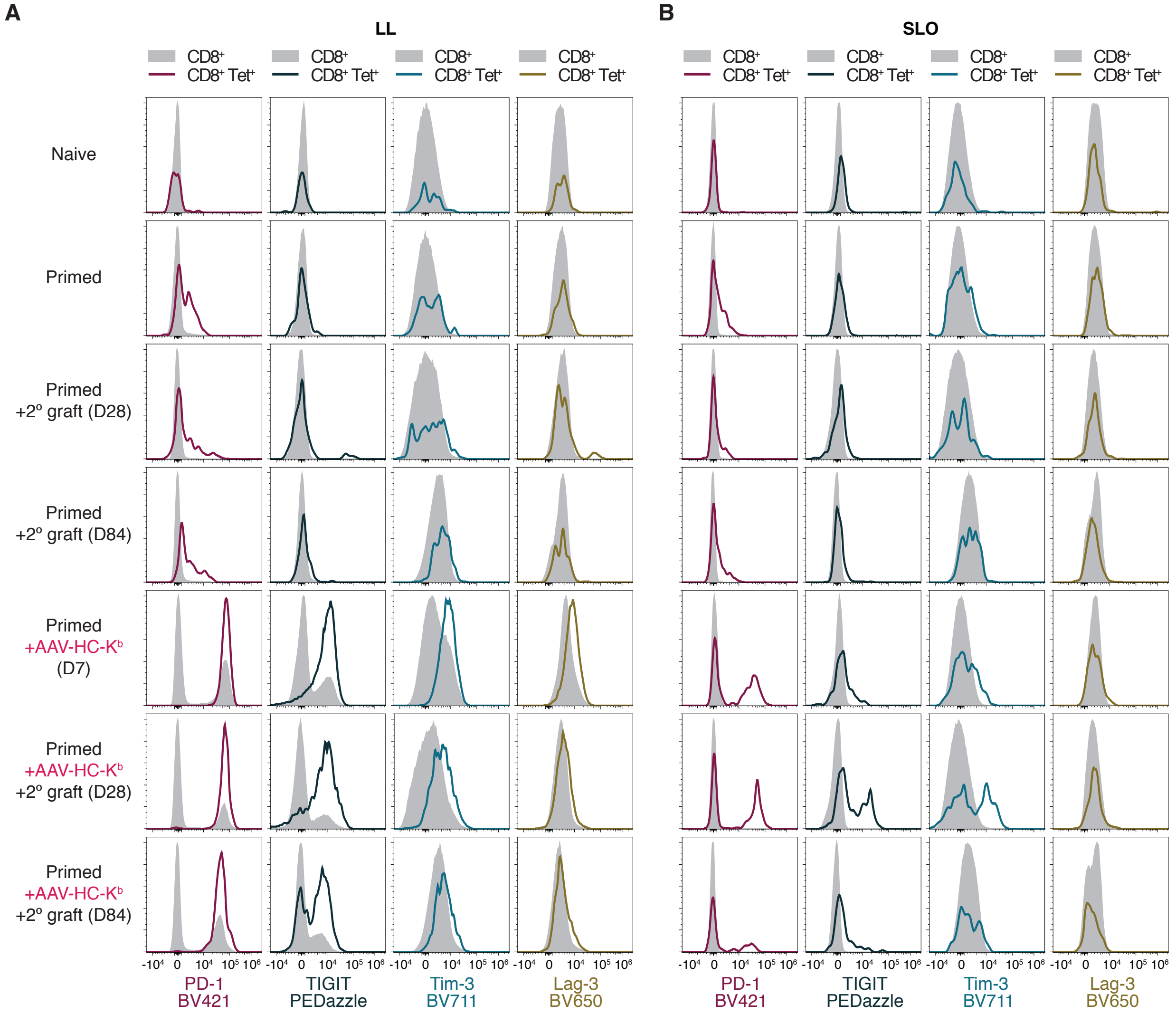

**Supplementary Figure 11.**

**Expression of coinhibitory receptors by bulk and tetramer-positive CD8+ T cells from liver or combined secondary lymphoid organs.**

Expression of PD-1, TIGIT, Tim-3 and LAG-3 was determined for tet+ and bulk CD8+ T cells in a model of secondary skin graft rejection or tolerance. Modest upregulation of PD-1 was noted in the CD8+tet+ liver leukocytes (**A**) and SLOs (**B**) at all intervals following graft rejection. In contrast, induction of tolerance upon inoculation of recipient mice with AAV-Kb was accompanied by strong expression of all coinhibitory ligands, with expression of LAG-3 and Tim-3 declining to baseline by protocol d84. Changes in the phenotype of CD8+tet+ cells were much less obvious within the bulk CD8+ populations. (**A-B**) representative flow plots from experiments with a total of n = 3 biological replicates per group.

Supplementary Table 2

| AA SEQUENCE | Peptide | Length | H-2Kᵈ IC50 (nM) | H-2Kᵈ Rank | Spectral intensity value DDA (hep) | Present in DIA (hep) | Spectral intensity value DDA (skin) | Present in DIA (skin) | Spectral intensity value DDA (spleen) | Present in DIA (spleen) | Accession |
|---|---|---|---|---|---|---|---|---|---|---|---|
| **AFHPSKAYI** | AFHPSKAYI | 9 | 302.7 | 0.3 | 4801600 | YES | 224950 | YES | 27405000 | YES | Q9ERG2 |
| **AFHPVHGTL** | AFHPVHGTL | 9 | 406.9 | 0.4 | | YES | | YES | 2174800 | YES | Q8C570 |
| **AFHSSRTSL** | AFHSSRTSL | 9 | 39.1 | 0.03 | | YES | 1219200 | YES | 32245000 | YES | Q9QXZ0 |
| **AFIPTINAI** | AFIPTINAI | 9 | 202.2 | 0.175 | 386170 | NO | 39941000 | YES | 62111000 | YES | P47930 |
| **AFLSSLTDV** | AFLSSLTDV | 9 | 228.5 | 0.25 | 919540 | NO | | YES | 18927000 | YES | Q3UVL4 |
| **AFQTILTEI** | AFQTILTEI | 9 | 53.5 | 0.05 | 273360 | YES | 8433500 | YES | 10894000 | YES | P62492 |
| **AFVATGTNL** | AFVATGTNL | 9 | 292.2 | 0.3 | 979870 | NO | 766910 | NO | 2340900 | NO | Q08024 |
| **AFVSMLNDI** | AFVSM(+15.99)LNDI | 9 | 227.8 | 0.25 | 12944000 | YES | 6645800 | YES | 41164000 | YES | P19096 |
| **AFVSMLNDI** | AFVSMLNDI | 9 | 227.8 | 0.25 | 2013300 | YES | 1389400 | YES | | YES | P19096 |
| **AGLQFPVGR** | AGLQFPVGR | 9 | 40663.1 | 85 | 699580 | NO | 477890 | YES | 722300 | YES | Q6GSS7:Q64523:C0HKE7:C0HKE6:C0HKE5:C0HKE3:C0HKE9:C0HKE2:C0HKE1:C0HKE4:C0HKE8:Q8BFU2:Q8CGP7:Q8R1M2:P27661:Q64522:P0C0S6 |
| **AISKLYSTI** | AISKLYSTI | 9 | 2926.9 | 1.7 | 1293300 | YES | 1583800 | YES | 23999000 | YES | Q6P5B0 |
| **AKERLLLWT** | AKERLLLWT | 9 | 36992 | 60 | | YES | | YES | | YES | Q91ZU6 |
| **ALLPTITQL** | ALLPTITQL | 9 | 3507.2 | 2 | 400250 | NO | 12164000 | YES | 27669000 | YES | C3VPR6 |
| **ALQANRTAL** | ALQANRTAL | 9 | 1290.7 | 0.9 | 49022 | NO | | YES | 1400400 | YES | Q6PB66 |
| **ALSRLFSSI** | ALSRLFSSI | 9 | 5105.7 | 3 | 423680 | NO | | YES | 7906300 | YES | Q3U308 |
| **ASLVNADKL** | ASLVNADKL | 9 | 21827.1 | 17 | | YES | | YES | | YES | P59016 |
| **ASVLNVNHI** | ASVLNVNHI | 9 | 13866.4 | 8 | | YES | 881800 | YES | 6652900 | YES | Q99NH0 |
| **ASYEFVQRL** | ASYEFVQRL | 9 | 28523.7 | 28 | | YES | 4687600 | YES | | YES | Q9JHU4 |

| | | | | | | | | | | | |
|---|---|---|---|---|---|---|---|---|---|---|---|
| **AVFPSIVGR** | AVFPSIVGR | 9 | 36425.3 | 55 | 988750 | YES | 878000 | YES | 150920 | YES | P60710:P63260:P68033:P62737 |
| **AVLSFSTRL** | AVLSFSTRL | 9 | 6750.8 | 4 | | YES | | YES | 2664100 | YES | P46978 |
| **AWAKALTDI** | AWAKALTDI | 9 | 883.8 | 0.7 | 2017000 | NO | 1174600 | YES | 1154900 | YES | O09117 |
| **AYAPAAATV** | AYAPAAATV | 9 | 31.6 | 0.02 | 15307000 | YES | 1207700 | YES | 42971000 | YES | O88532 |
| **AYAPAIHQI** | AYAPAIHQI | 9 | 25.5 | 0.015 | 8671500 | YES | 3783900 | YES | 71931000 | YES | E9Q7E2 |
| **AYAPSGNFV** | AYAPSGNFV | 9 | 37.7 | 0.03 | 19059000 | YES | 26714000 | YES | 46585000 | YES | P62880:Q61011 |
| **AYAPSGNYV** | AYAPSGNYV | 9 | 22.2 | 0.01 | 4199800 | YES | 19868000 | YES | 91558000 | YES | P62874:P29387 |
| **AYDGVRGSL** | AYDGVRGSL | 9 | 361.9 | 0.4 | 133580 | NO | 315290 | YES | 11411000 | YES | Q8C4J7 |
| **AYFHLLNQI** | AYFHLLNQI | 9 | 51.1 | 0.04 | 4804500 | YES | 91785000 | YES | 159680000 | YES | Q99K51 |
| **AYGSLFNSI** | AYGSLFNSI | 9 | 39 | 0.03 | 73719000 | YES | 2485800 | YES | 15326000 | YES | Q71RI9 |
| **AYGVAVNKL** | AYGVAVNKL | 9 | 649.1 | 0.6 | 2016100 | YES | 403540 | YES | 3307600 | YES | Q80XI6 |
| **AYHGGHLTI** | AYHGGHLTI | 9 | 43.6 | 0.04 | 8511100 | YES | 1852400 | YES | 14381000 | YES | Q9CPX7 |
| **AYHQALSRV** | AYHQALSRV | 9 | 38.5 | 0.03 | 13361000 | YES | 3349900 | YES | 21002000 | YES | P43883 |
| **AYHTQTTPL** | AYHTQTTPL | 9 | 12.8 | 0.01 | 164470 | NO | 212260 | YES | 6796200 | NO | Q9WTP6 |
| **AYIPLNNYL** | AYIPLNNYL | 9 | 23.8 | 0.015 | 7756400 | YES | 9993000 | YES | 27978000 | YES | Q8C878 |
| **AYITGKEDI** | AYITGKEDI | 9 | 19.4 | 0.01 | 421910 | NO | | YES | 10259000 | YES | Q60767 |
| **AYKANRDLI** | AYKANRDLI | 9 | 395.8 | 0.4 | 2276200 | NO | 7942100 | NO | 23426000 | NO | P26231 |
| **AYKAVLNYL** | AYKAVLNYL | 9 | 43.8 | 0.04 | 1311500 | NO | 9871000 | NO | 87473000 | NO | Q3TUU5 |
| **AYKFGKTVV** | AYKFGKTVV | 9 | 113.3 | 0.125 | 14182000 | NO | 4185100 | NO | 36495000 | YES | Q8R3Q0 |
| **AYKVLKTEM** | AYKVLKTEM | 9 | 111 | 0.125 | 301280 | NO | 59993 | NO | | YES | Q8BI84 |
| **AYKWIRTSL** | AYKWIRTSL | 9 | 48.3 | 0.04 | 912120 | NO | 936360 | NO | 15429000 | NO | Q8VE88 |
| **AYLAALTQL** | AYLAALTQL | 9 | 14.7 | 0.01 | 401190 | NO | 16738000 | YES | 99174000 | YES | P54310 |
| **AYLGTITKT** | AYLGTITKT | 9 | 997.6 | 0.8 | 259800 | NO | | YES | 588300 | YES | O88545 |
| **AYLHAQHYI** | AYLHAQHYI | 9 | 28.4 | 0.02 | | YES | 99485 | YES | 1034700 | YES | Q9R117 |
| **AYLHSHNMI** | AYLHSHNM(+15.99)I | 9 | 17.8 | 0.01 | 96867 | NO | 415040 | YES | 4841000 | YES | Q6ZQ29 |
| **AYLHSHTMI** | AYLHSHTM(+15.99)I | 9 | 10 | 0.01 | 547090 | NO | 698980 | YES | 9570100 | YES | Q5F2E8 |
| **AYLHSHTMI** | AYLHSHTMI | 9 | 10 | 0.01 | 586530 | NO | 1169700 | YES | | YES | Q5F2E8 |
| **AYLLNLNHL** | AYLLNLNHL | 9 | 137.5 | 0.125 | 20372000 | YES | 21824000 | YES | 276160000 | YES | Q91V04 |
| **AYLPQTSRL** | AYLPQTSRL | 9 | 65.8 | 0.06 | | YES | 2246600 | YES | 60929000 | YES | Q6NXY1 |

| | | | | | | | | | | | |
|---|---|---|---|---|---|---|---|---|---|---|---|
| **AYLPQYTHM** | AYLPQYTHM(+15.99) | 9 | 40 | 0.03 | 15770000 | YES | 51156000 | YES | 552720000 | NO | Q91W59 |
| **AYLTDLTKL** | AYLTDLTKL | 9 | 75.4 | 0.07 | 669560 | NO | | YES | 20522000 | YES | G5E8P0 |
| **AYLVDIKTI** | AYLVDIKTI | 9 | 120 | 0.125 | 8257400 | YES | 17114000 | YES | 24060000 | YES | Q6VH22 |
| **AYMKSSRYI** | AYM(+15.99)KSSRYI | 9 | 10.6 | 0.01 | 5584800 | NO | 102970 | YES | 4117800 | YES | P10649 |
| **AYMKSSRYI** | AYMKSSRYI | 9 | 10.6 | 0.01 | 719650 | NO | 110900 | YES | | YES | P10649 |
| **AYNILIGEL** | AYNILIGEL | 9 | 766.3 | 0.6 | 198110 | YES | | YES | 26291000 | YES | Q5SV77 |
| **AYNPGQAVP** | AYNPGQAVP | 9 | 12730.9 | 7.5 | 2218500 | NO | | YES | 4581600 | YES | Q8VDP4 |
| **AYNRILDAL** | AYNRILDAL | 9 | 95.8 | 0.09 | 2140500 | YES | 3750600 | NO | 20923000 | YES | Q68FD7 |
| **AYQELLRLI** | AYQELLRLI | 9 | 157.2 | 0.15 | | YES | 3798900 | YES | 1990000 | YES | Q9JLB2 |
| **AYQNNKELL** | AYQNNKELL | 9 | 304.4 | 0.3 | 7911200 | YES | 10425000 | YES | 172590000 | YES | P57080 |
| **AYQRVFHSL** | AYQRVFHSL | 9 | 19.3 | 0.01 | 234370 | NO | 311020 | YES | 1952000 | YES | Q3TUA9 |
| **AYQSIQSYL** | AYQSIQSYL | 9 | 19.1 | 0.01 | 59021000 | YES | 55445000 | YES | 134100000 | YES | Q925U4 |
| **AYRQLRETL** | AYRQLRETL | 9 | 92 | 0.09 | 11090000 | NO | 4210400 | NO | 37984000 | NO | Q9D083 |
| **AYSFGRTTI** | AYSFGRTTI | 9 | 15.1 | 0.01 | 13677000 | YES | 1371500 | NO | 5356100 | NO | Q3TWI9 |
| **AYSGIKNQL** | AYSGIKNQL | 9 | 200.6 | 0.175 | 1395300 | YES | 1009700 | YES | 14226000 | YES | Q9ERG2 |
| **AYSGVKNSL** | AYSGVKNSL | 9 | 60.3 | 0.06 | 1372700 | YES | 876020 | YES | 39980000 | YES | G5E829 |
| **AYSIVIRQI** | AYSIVIRQI | 9 | 377.2 | 0.4 | 5487500 | YES | 13654000 | YES | 33996000 | YES | Q80TP3 |
| **AYSKLGNYV** | AYSKLGNYV | 9 | 104.4 | 0.1 | 15272000 | YES | 4448600 | NO | 57828000 | YES | Q8BJU0 |
| **AYSQAHLTL** | AYSQAHLTL | 9 | 96.6 | 0.09 | 1210300 | NO | | YES | 8048900 | YES | Q9DBE9 |
| **AYSRSMTKL** | AYSRSM(+15.99)TKL | 9 | 47 | 0.04 | 1457900 | YES | 1330500 | YES | 31203000 | YES | Q3UQN2 |
| **AYSRSMTKL** | AYSRSMTKL | 9 | 47 | 0.04 | 119590 | YES | 1887500 | YES | | YES | Q3UQN2 |
| **AYSTLLSHI** | AYSTLLSHI | 9 | 16.6 | 0.01 | 10019000 | YES | 1997600 | YES | 34845000 | YES | Q8CFX1 |
| **AYVAMNERL** | AYVAM(+15.99)NERL | 9 | 43.1 | 0.04 | 2529800 | YES | 3514000 | YES | 271380000 | YES | Q9D7E4 |
| **AYVAMNERL** | AYVAMNERL | 9 | 43.1 | 0.04 | 6731200 | YES | | YES | | YES | Q9D7E4 |
| **AYVAPTNDL** | AYVAPTNDL | 9 | 246.7 | 0.25 | 12962000 | YES | 118820 | NO | 159830 | YES | Q5SW19 |
| **AYVETQDQL** | AYVETQDQL | 9 | 365.7 | 0.4 | 1502500 | NO | 3448500 | YES | 27736000 | YES | F6ZDS4 |
| **AYVHVVTHF** | AYVHVVTHF | 9 | 357.8 | 0.4 | 1261200 | YES | 212630 | YES | 440820 | YES | Q9D2C7 |
| **AYVPGFAHI** | AYVPGFAHI | 9 | 102.1 | 0.1 | 120030000 | YES | 170250000 | YES | 1010600000 | YES | Q9CXX9 |
| **AYVPQQAWI** | AYVPQQAWI | 9 | 507.1 | 0.5 | 20734000 | YES | | YES | 2886600 | YES | Q8VI47:B2RX12:O35379 |

| | | | | | | | | | | | |
|---|---|---|---|---|---|---|---|---|---|---|---|
| **AYVPSHSDA** | AYVPSHSDA | 9 | 288.3 | 0.3 | | YES | | YES | 1951900 | YES | Q8R420 |
| **AYWAGGLHL** | AYWAGGLHL | 9 | 92 | 0.09 | 10670000 | YES | 4115600 | YES | 26656000 | YES | Q8BGH2 |
| **AYWPGLQSL** | AYWPGLQSL | 9 | 53.7 | 0.05 | 20978000 | YES | 1578700 | YES | 10007000 | YES | Q8BJT9 |
| **DFHPSGTVV** | DFHPSGTVV | 9 | 936.9 | 0.7 | 2602900 | YES | | YES | 21906000 | YES | Q3UMY5 |
| **DFITVHTPL** | DFITVHTPL | 9 | 75.3 | 0.07 | | YES | | YES | 1777700 | YES | Q61753 |
| **DLLPSHSTI** | DLLPSHSTI | 9 | 689.9 | 0.6 | 14804000 | YES | 841920 | YES | 1515000 | YES | Q99MR8 |
| **DYHHIHTEI** | DYHHIHTEI | 9 | 35.5 | 0.025 | 448990 | YES | 367660 | YES | 3992400 | YES | Q9EPE9 |
| **DYIALNEDL** | DYIALNEDL | 9 | 237.8 | 0.25 | | YES | 3949300 | YES | 907390 | YES | P01901 |
| **DYIITPHAL** | DYIITPHAL | 9 | 166.8 | 0.15 | 28525000 | YES | 36810000 | YES | 35211000 | YES | Q3TWF6 |
| **DYIYGVTYI** | DYIYGVTYI | 9 | 25.6 | 0.015 | 5728600 | YES | | YES | 2340300 | YES | Q91ZX7 |
| **DYKESFNTI** | DYKESFNTI | 9 | 61.1 | 0.06 | 3436200 | NO | 81121000 | YES | 1310500 | YES | Q921D9 |
| **DYLADKSYI** | DYLADKSYI | 9 | 31.2 | 0.02 | 111990000 | YES | 16793000 | YES | 68925000 | YES | O70251 |
| **DYLGSRQYV** | DYLGSRQYV | 9 | 56.8 | 0.05 | 57315000 | NO | 27100000 | NO | 111590000 | NO | P52332 |
| **DYLNVVNEL** | DYLNVVNEL | 9 | 130.2 | 0.125 | 1776300 | YES | 567220 | YES | 1365400 | YES | Q6ZQB6 |
| **DYLPDRELV** | DYLPDRELV | 9 | 1829.5 | 1.2 | 6299900 | NO | 9177800 | NO | 27360000 | NO | Q5SW75 |
| **DYLPSWQKI** | DYLPSWQKI | 9 | 171.6 | 0.175 | 5536600 | NO | | YES | 327260 | NO | Q91XC9 |
| **DYMEALTRL** | DYM(+15.99)EALTRL | 9 | 34.7 | 0.025 | 1207000 | YES | 4203400 | YES | 54106000 | YES | Q99K70 |
| **DYMEALTRL** | DYMEALTRL | 9 | 34.7 | 0.025 | 4842100 | YES | 5164600 | YES | | YES | Q99K70 |
| **DYNRIGSSL** | DYNRIGSSL | 9 | 45 | 0.04 | 14104000 | YES | 7430600 | YES | 32374000 | YES | Q6P8X1 |
| **DYNTAHNKV** | DYNTAHNKV | 9 | 109.6 | 0.1 | 614960 | YES | | YES | 428560 | YES | Q61510 |
| **DYQALRTSI** | DYQALRTSI | 9 | 8.7 | 0.01 | 44113000 | YES | 36431000 | YES | 802740000 | YES | Q68FD5 |
| **DYQDVRNEI** | DYQDVRNEI | 9 | 42.6 | 0.04 | 763740 | NO | | YES | 4406200 | YES | G5E829 |
| **DYQPGITFI** | DYQPGITFI | 9 | 24 | 0.015 | 1956100 | NO | 9724600 | YES | 65946000 | YES | Q8CJG0 |
| **DYQPGITYI** | DYQPGITYI | 9 | 15.1 | 0.01 | 7258000 | YES | 6303900 | YES | 17776000 | YES | Q8CJG1:Q8CJF9 |
| **DYQRLLQTI** | DYQRLLQTI | 9 | 20.5 | 0.01 | 6339000 | YES | 11135000 | YES | 33648000 | YES | Q6KAQ7 |
| **DYVVGFTDL** | DYVVGFTDL | 9 | 305.1 | 0.3 | 504840 | NO | | YES | 398480 | NO | Q9CQ79 |
| **DYYPDRTYI** | DYYPDRTYI | 9 | 29.8 | 0.02 | 2425300 | NO | 3058400 | YES | 94280000 | YES | Q8JZL7 |
| **EGPVTEQVK** | EGPVTEQVK | 9 | 42038.8 | 90 | 5673300 | NO | | YES | 1071600 | YES | Q7TQH0 |
| **EYEKIKSQL** | EYEKIKSQL | 9 | 82.8 | 0.08 | 211480 | NO | | YES | 469070 | NO | B2RXS4 |

| | | | | | | | | | | | |
|---|---|---|---|---|---|---|---|---|---|---|---|
| **EYEPGKSSI** | EYEPGKSSI | 9 | 23.8 | 0.015 | 2963200 | NO | 83781 | NO | 10092000 | NO | Q3UTJ2 |
| **EYIHALTLL** | EYIHALTLL | 9 | 96.2 | 0.09 | 1131600 | NO | | YES | 10427000 | YES | O35382 |
| **EYIHSKNFI** | EYIHSKNFI | 9 | 26.8 | 0.02 | 145970000 | YES | 59681000 | YES | 377220000 | YES | Q9DC28:Q9JMK2 |
| **EYIKVITGL** | EYIKVITGL | 9 | 78.7 | 0.08 | 9480000 | YES | 17114000 | YES | 35285000 | YES | Q6A070 |
| **EYLEGRNLI** | EYLEGRNLI | 9 | 90.2 | 0.09 | | YES | 6789600 | YES | 12951000 | YES | Q91Z67 |
| **EYLPSQITI** | EYLPSQITI | 9 | 67.2 | 0.06 | 6041700 | YES | 40351000 | YES | 32736000 | YES | Q8R151 |
| **EYMKVQTEI** | EYMKVQTEI | 9 | 13.3 | 0.01 | 2485300 | YES | 1204200 | YES | | YES | Q62073 |
| **EYNDLKTEL** | EYNDLKTEL | 9 | 124.5 | 0.125 | 321160 | NO | 872110 | YES | 5006700 | YES | Q8BU04 |
| **EYNELLTAI** | EYNELLTAI | 9 | 34.3 | 0.025 | 413120 | NO | 6739600 | YES | 49390000 | YES | P70428 |
| **EYVANLTEL** | EYVANLTEL | 9 | 100.4 | 0.1 | 41849000 | YES | | YES | 76108000 | YES | Q3TKT4 |
| **EYVANLTNL** | EYVANLTNL | 9 | 101.7 | 0.1 | | YES | 9411600 | YES | 29909000 | YES | Q6DIC0 |
| **EYVATTFDI** | EYVATTFDI | 9 | 353.3 | 0.3 | 1559600 | NO | | YES | | YES | Q922E6 |
| **EYVHTKNFI** | EYVHTKNFI | 9 | 65.8 | 0.06 | 42833000 | YES | 22258000 | YES | 141560000 | YES | Q8BK63 |
| **EYVRNQQTI** | EYVRNQQTI | 9 | 76.3 | 0.07 | 294130 | NO | | YES | 1334800 | YES | Q9CQC6 |
| **EYWRLIGEL** | EYWRLIGEL | 9 | 458.4 | 0.4 | 1361900 | NO | 874200 | YES | | YES | P97352 |
| **FAYEGRDYI** | FAYEGRDYI | 9 | 4274.6 | 2.5 | | YES | 11493000 | YES | 10900000 | YES | P01899 |
| **FFSEIISSI** | FFSEIISSI | 9 | 477.5 | 0.4 | 489660 | NO | 531820 | YES | 372150 | YES | Q925E7:Q6P1F6 |
| **FFSTIRTEL** | FFSTIRTEL | 9 | 151.6 | 0.15 | 42866000 | YES | 29077000 | YES | 551310000 | YES | P50172 |
| **FFVENVSEL** | FFVENVSEL | 9 | 1873.6 | 1.2 | 551130 | NO | 1560300 | YES | 14915000 | YES | Q6KCD5 |
| **FGPVNHEEL** | FGPVNHEEL | 9 | 27315.3 | 26 | 304380 | NO | | YES | 30507000 | YES | P46414 |
| **FLLPILSQI** | FLLPILSQI | 9 | 1108.5 | 0.8 | 1045300 | YES | 1365900 | YES | 7734300 | YES | Q62167:Q62095 |
| **FYEPQKGSI** | FYEPQKGSI | 9 | 28.1 | 0.02 | 543420 | NO | | YES | | YES | Q61102 |
| **FYFASKLVL** | FYFASKLVL | 9 | 32.7 | 0.025 | | YES | 13400000 | YES | 15821000 | YES | Q07813 |
| **FYFQQKGQL** | FYFQQKGQL | 9 | 43 | 0.04 | 1557800 | NO | 1789400 | YES | | YES | O55229 |
| **FYGDVQTHI** | FYGDVQTHI | 9 | 29.6 | 0.02 | 7727200 | YES | | YES | | YES | Q6ZPE2 |
| **FYHPETTQL** | FYHPETTQL | 9 | 56.8 | 0.05 | 6473600 | YES | | YES | | YES | Q61584 |
| **FYIGLGSRI** | FYIGLGSRI | 9 | 16 | 0.01 | 41374000 | YES | 23817000 | YES | 125600000 | YES | Q8VCV1:Q7M759 |
| **FYLETQQQI** | FYLETQQQI | 9 | 29.4 | 0.02 | 1217300 | NO | | YES | 10760000 | YES | Q99MP8 |
| **FYLGSFDNI** | FYLGSFDNI | 9 | 34.2 | 0.025 | 1094800 | YES | 4021000 | YES | 12358000 | YES | Q6GV12 |

| | | | | | | | | | | | |
|---|---|---|---|---|---|---|---|---|---|---|---|
| **FYLPAPGTL** | FYLPAPGTL | 9 | 18.8 | 0.01 | 781250 | NO | 1851900 | YES | 1631300 | YES | Q80WC3 |
| **FYNPAVSRI** | FYNPAVSRI | 9 | 17 | 0.01 | | YES | 3912800 | YES | 56969000 | YES | Q80XR2 |
| **FYNQVSTPL** | FYNQVSTPL | 9 | 8.6 | 0.01 | 10580000 | YES | | YES | 1776800 | YES | Q61703 |
| **FYNVDISYL** | FYNVDISYL | 9 | 103.8 | 0.1 | 652900 | YES | | YES | 5337700 | YES | P10605 |
| **FYQKADHTL** | FYQKADHTL | 9 | 12.8 | 0.01 | 2423100 | YES | 688220 | YES | 3961700 | YES | Q8K2Q7 |
| **FYQQQAGGL** | FYQQQAGGL | 9 | 119.8 | 0.125 | 114990 | NO | 525240 | YES | 4545800 | YES | Q91YE7 |
| **FYSAGKNYL** | FYSAGKNYL | 9 | 26.1 | 0.015 | 3573000 | YES | | YES | 43423000 | YES | Q8R3N6 |
| **FYSNIQTVI** | FYSNIQTVI | 9 | 54.4 | 0.05 | | YES | 31998000 | YES | 113460000 | YES | P08775 |
| **FYSQDLTHL** | FYSQDLTHL | 9 | 73.6 | 0.07 | 926340 | NO | 3867900 | YES | 4288500 | YES | Q8CHE4 |
| **FYSTTHGAL** | FYSTTHGAL | 9 | 35.9 | 0.025 | 1215300 | YES | | YES | | YES | Q8C0L6 |
| **FYTPIPNGL** | FYTPIPNGL | 9 | 200 | 0.175 | 121730000 | YES | 48789000 | YES | 38209000 | YES | O08573 |
| **FYTQLLQEL** | FYTQLLQEL | 9 | 69.7 | 0.07 | 92251 | NO | 615870 | YES | 809110 | YES | Q5RJH6 |
| **FYVATSRQL** | FYVATSRQL | 9 | 37 | 0.025 | 38324000 | YES | | YES | | YES | Q8VI47:B2RX12 |
| **FYVPSVSQL** | FYVPSVSQL | 9 | 20.6 | 0.01 | 3321900 | YES | | YES | 23206000 | YES | Q62136 |
| **GAVTNVKVI** | GAVTNVKVI | 9 | 17664 | 12 | | YES | | YES | 2167800 | YES | P70372 |
| **GFHPSGSVL** | GFHPSGSVL | 9 | 601.8 | 0.5 | 3094100 | YES | 3561900 | YES | 67444000 | YES | Q7TNG5 |
| **GFHRTISHL** | GFHRTISHL | 9 | 324.5 | 0.3 | 346380 | NO | 201600 | NO | 850400 | YES | Q9DBU5 |
| **GFIASHVIV** | GFIASHVIV | 9 | 2437 | 1.5 | | YES | 1744900 | YES | 2906200 | YES | Q8VDR7 |
| **GFLKSISNV** | GFLKSISNV | 9 | 437.8 | 0.4 | 1660100 | NO | 1935900 | NO | 6027800 | YES | Q8C176 |
| **GFNPALQLI** | GFNPALQLI | 9 | 2641.9 | 1.6 | 20014000 | YES | 96580000 | YES | 140080000 | YES | Q9ER81 |
| **GFNSSISNI** | GFNSSISNI | 9 | 205.1 | 0.2 | | YES | 1127100 | YES | 13727000 | YES | A2AJ88 |
| **GGIQNVGHI** | GGIQNVGHI | 9 | 9776.3 | 5.5 | 113890 | YES | | YES | 17747000 | YES | P24547 |
| **GINEAGISR** | GINEAGISR | 9 | 35325.7 | 50 | | YES | | YES | | YES | P29533 |
| **GWGELQNTI** | GWGELQNTI | 9 | 7006.9 | 4 | 2588600 | NO | 1697500 | YES | 1220900 | YES | P24527 |
| **GYAKLIAEL** | GYAKLIAEL | 9 | 255 | 0.25 | 3698100 | YES | 5673900 | YES | 27228000 | YES | B1AZI6 |
| **GYALPHAIL** | GYALPHAIL | 9 | 1552.7 | 1 | 2056600 | YES | 1755600 | YES | 4444900 | YES | P60710:P63260:Q8BFZ3 |
| **GYATLHHVI** | GYATLHHVI | 9 | 21.8 | 0.01 | | YES | 1018700 | YES | 13564000 | YES | Q925U4 |
| **GYATTYRQL** | GYATTYRQL | 9 | 49.2 | 0.04 | | YES | | YES | 5257900 | YES | Q09200 |
| **GYEETLTRL** | GYEETLTRL | 9 | 81.3 | 0.08 | 2427400 | YES | | YES | 1426400 | YES | Q5XPI3 |

| | | | | | | | | | | | |
|---|---|---|---|---|---|---|---|---|---|---|---|
| **GYERIYNEI** | GYERIYNEI | 9 | 54.1 | 0.05 | 124340000 | YES | 4814800 | YES | 16195000 | YES | Q61823 |
| **GYFEVTHDI** | GYFEVTHDI | 9 | 56.9 | 0.05 | 414680000 | YES | 11397000 | YES | 39391000 | YES | P24270 |
| **GYFGSTQGL** | GYFGSTQGL | 9 | 311.3 | 0.3 | 2587900 | YES | | YES | 12818000 | YES | Q6KAR6 |
| **GYFNTYKLL** | GYFNTYKLL | 9 | 341.6 | 0.3 | 1505800 | NO | 2386400 | YES | 19343000 | YES | Q8BT60 |
| **GYFPNKKQL** | GYFPNKKQL | 9 | 138.3 | 0.125 | 6191400 | YES | 1735700 | YES | 11806000 | YES | P70388 |
| **GYGQGAGTL** | GYGQGAGTL | 9 | 101.9 | 0.1 | 1272900 | NO | | YES | 20702000 | YES | P97315 |
| **GYGRSILTV** | GYGRSILTV | 9 | 127.8 | 0.125 | 4729400 | YES | | YES | 201240 | NO | A2A8Z1 |
| **GYGTQKSSL** | GYGTQKSSL | 9 | 25.1 | 0.015 | 43231 | NO | | YES | 5587100 | YES | Q99KW3 |
| **GYIASLHEL** | GYIASLHEL | 9 | 24.4 | 0.015 | 30330000 | YES | 712280 | YES | 4194500 | YES | Q920R0 |
| **GYIESIQHI** | GYIESIQHI | 9 | 22 | 0.01 | | YES | 806020 | YES | 4452500 | YES | Q9CXC3 |
| **GYIGGKKEI** | GYIGGKKEI | 9 | 37.6 | 0.03 | 1982400 | YES | 1776500 | YES | 7325400 | YES | |
| **GYIGGKKEL** | GYIGGKKEL | 9 | 80 | 0.08 | 1982400 | YES | 1776500 | YES | 7325400 | YES | P97363 |
| **GYIGSHTVL** | GYIGSHTVL | 9 | 22 | 0.01 | 88165000 | YES | 9113400 | YES | 93856000 | YES | Q8R059 |
| **GYILSIHRI** | GYILSIHRI | 9 | 49.6 | 0.04 | 9524200 | YES | 1254500 | YES | 6468000 | YES | Q9Z0M5 |
| **GYIPSASMT** | GYIPSASM(+15.99)T | 9 | 74.7 | 0.07 | 520180 | NO | 308350 | NO | 4624600 | NO | Q9QYF9 |
| **GYIQTGDRL** | GYIQTGDRL | 9 | 92.9 | 0.09 | 2273000 | YES | | YES | | YES | Q69ZS7 |
| **GYISIANGL** | GYISIANGL | 9 | 57.1 | 0.05 | 5062200 | YES | 2643900 | YES | 9879100 | YES | Q99LC2 |
| **GYKAGMTHI** | GYKAGM(+15.99)THI | 9 | 15 | 0.01 | 7341100 | YES | 9721800 | YES | 746490000 | YES | P27659 |
| **GYKAGMTHI** | GYKAGMTHI | 9 | 15 | 0.01 | 6111700 | YES | 11103000 | YES | 2422300 | YES | P27659 |
| **GYKESFSSI** | GYKESFSSI | 9 | 28.2 | 0.02 | 5685800 | NO | 618920 | NO | 17230000 | NO | Q80YE7 |
| **GYLDGRLEP** | GYLDGRLEP | 9 | 5733.6 | 3.5 | 123760000 | YES | 2485300 | YES | 1563600 | YES | P53566 |
| **GYLELLDHV** | GYLELLDHV | 9 | 138.2 | 0.125 | 8150400 | YES | 114840000 | YES | 751550000 | YES | P26039 |
| **GYLGGENRV** | GYLGGENRV | 9 | 210 | 0.2 | 1495300 | YES | | YES | 1061000 | YES | E9Q5F9 |
| **GYLGQVTTI** | GYLGQVTTI | 9 | 15.1 | 0.01 | | YES | 93831000 | YES | 344690000 | YES | Q9Z1F9 |
| **GYLKGYTLV** | GYLKGYTLV | 9 | 45.2 | 0.04 | 6891400 | YES | 32833000 | YES | 114640000 | YES | Q9CWZ3 |
| **GYLKLWDTV** | GYLKLWDTV | 9 | 71.6 | 0.07 | 528650 | NO | 3632000 | YES | 2715700 | YES | P97499 |
| **GYLLDKETL** | GYLLDKETL | 9 | 167.7 | 0.15 | 2045500 | YES | 3492000 | YES | 15829000 | YES | Q80ZJ6 |
| **GYLPGNEKL** | GYLPGNEKL | 9 | 164.4 | 0.15 | 47000000 | YES | 30146000 | YES | 181020000 | YES | P97372 |
| **GYLPLAHVL** | GYLPLAHVL | 9 | 64.3 | 0.06 | 119230000 | YES | 123210000 | YES | 356670000 | YES | Q9QUJ7:Q9CZW4 |

| | | | | | | | | | | | |
|---|---|---|---|---|---|---|---|---|---|---|---|
| **GYLPNKQVL** | GYLPNKQVL | 9 | 126.1 | 0.125 | 1231000000 | YES | 77832000 | YES | | YES | Q05915 |
| **GYLPTQQDV** | GYLPTQQDV | 9 | 193.7 | 0.175 | 1139300 | NO | | YES | | YES | P21278 |
| **GYLPVQTVL** | GYLPVQTVL | 9 | 31.1 | 0.02 | 1492400 | NO | 47236000 | YES | 471550000 | YES | P11531 |
| **GYLSSRVLL** | GYLSSRVLL | 9 | 95.1 | 0.09 | 15237000 | YES | 8758400 | YES | 88262000 | YES | Q5NCS9 |
| **GYLTNPRSL** | GYLTNPRSL | 9 | 66.9 | 0.06 | | YES | 484650 | YES | | YES | Q924Z5 |
| **GYMGHLTRI** | GYM(+15.99)GHLTRI | 9 | 42.3 | 0.04 | 945800 | YES | 182830 | YES | 1829700 | YES | Q922D4 |
| **GYMRLLEFI** | GYM(+15.99)RLLEFI | 9 | 26.7 | 0.02 | 3700600 | NO | 17673000 | YES | 34678000 | NO | Q9ER73 |
| **GYMGHLTRI** | GYMGHLTRI | 9 | 42.3 | 0.04 | | YES | | YES | | YES | Q922D4 |
| **GYNKAKAYI** | GYNKAKAYI | 9 | 33.7 | 0.025 | 40651 | NO | 158340 | YES | 1995900 | YES | Q05909 |
| **GYNKQNTTL** | GYNKQNTTL | 9 | 16.3 | 0.01 | 622260 | YES | | YES | 25708000 | YES | E9Q3L2 |
| **GYNRVLQFL** | GYNRVLQFL | 9 | 77.5 | 0.07 | 2877000 | YES | 26707000 | YES | 6624200 | YES | Q9R269 |
| **GYNSVQHLI** | GYNSVQHLI | 9 | 77.9 | 0.08 | 1264000 | NO | | YES | 22084000 | YES | Q91V83 |
| **GYQRLLELI** | GYQRLLELI | 9 | 73.8 | 0.07 | 1119500 | NO | 16437000 | YES | 36589000 | YES | A2AIV2 |
| **GYQTAFSQL** | GYQTAFSQL | 9 | 14.4 | 0.01 | 1359600 | NO | 4738600 | YES | 21117000 | YES | Q9ERK4 |
| **GYQTVKEAL** | GYQTVKEAL | 9 | 13.3 | 0.01 | 145620000 | YES | 7417000 | YES | | YES | P33267 |
| **GYQVLRSVL** | GYQVLRSVL | 9 | 35.9 | 0.025 | 745030 | NO | 7632300 | YES | 44588000 | YES | Q8R5K4 |
| **GYSHILTNI** | GYSHILTNI | 9 | 61.6 | 0.06 | 1901500 | YES | 164500 | YES | 680790 | NO | Q60866 |
| **GYSKLYDDI** | GYSKLYDDI | 9 | 403 | 0.4 | 4912000 | YES | 268340 | NO | 1581900 | NO | O88822 |
| **GYSMNHQVI** | GYSM(+15.99)NHQVI | 9 | 434.8 | 0.4 | 5764000 | YES | 1507500 | YES | 8976600 | YES | P46935 |
| **GYSMNHQVI** | GYSMNHQVI | 9 | 434.8 | 0.4 | 8292300 | YES | 1878800 | YES | | YES | P46935 |
| **GYSPQLQGL** | GYSPQLQGL | 9 | 1499.7 | 1 | 7576800 | YES | 3711900 | YES | 8228100 | YES | Q8BRG8 |
| **GYTHGMHTL** | GYTHGMHTL | 9 | 64.2 | 0.06 | 2594400 | YES | | YES | | YES | Q9R049 |
| **GYTITPNTL** | GYTITPNTL | 9 | 78 | 0.08 | 1281200 | NO | | YES | 335610 | NO | Q61139 |
| **GYVDNKEFV** | GYVDNKEFV | 9 | 785.1 | 0.6 | 152950000 | YES | 102660000 | YES | 459180000 | YES | P01899:P01897 |
| **GYVETPRGL** | GYVETPRGL | 9 | 929.9 | 0.7 | 15489000 | YES | 7483300 | YES | 68411000 | YES | P27659 |
| **GYVPSQADV** | GYVPSQADV | 9 | 558.5 | 0.5 | 3292300 | YES | 600450 | YES | 10131000 | YES | O70251 |
| **GYVQGINDL** | GYVQGINDL | 9 | 191.6 | 0.175 | 20575000 | YES | 6004500 | YES | 17717000 | YES | Q8R5A6 |
| **GYWNTYTEL** | GYWNTYTEL | 9 | 34.1 | 0.025 | 3198700 | NO | | YES | 66718000 | YES | Q3V2Q8 |
| **GYYNNSTKV** | GYYNNSTKV | 9 | 82.5 | 0.08 | 147360 | NO | 377760 | YES | 53721000 | YES | P25911 |

| | | | | | | | | | | | |
|---|---|---|---|---|---|---|---|---|---|---|---|
| **GYYQANERV** | GYYQANERV | 9 | 102.9 | 0.1 | 153050 | YES | | YES | 8398700 | YES | O70378 |
| **HFIEGGRTV** | HFIEGGRTV | 9 | 819.3 | 0.7 | 6713500 | YES | | YES | 2473000 | YES | Q7TQI3 |
| **HFLPMLQTV** | HFLPM(+15.99)LQTV | 9 | 132.1 | 0.125 | 93378000 | YES | 180070000 | YES | 1572300000 | NO | Q60605 |
| **HFLPMLQTV** | HFLPMLQTV | 9 | 132.1 | 0.125 | 54390000 | YES | 142220000 | YES | 368670 | NO | Q60605 |
| **HFNPVQTQI** | HFNPVQTQI | 9 | 231 | 0.25 | | YES | | YES | 12779000 | YES | E9PZJ8 |
| **HFSTVKTHL** | HFSTVKTHL | 9 | 94.9 | 0.09 | 346980 | NO | 854010 | YES | 3848200 | YES | Q62348 |
| **HFVEGQTVV** | HFVEGQTVV | 9 | 2352.2 | 1.4 | 7210500 | YES | 1698000 | YES | 16198000 | YES | Q6NS46 |
| **HFYSSISSL** | HFYSSISSL | 9 | 121 | 0.125 | 43117000 | YES | 59111000 | YES | 50938000 | YES | Q60795 |
| **HFYSSKSEI** | HFYSSKSEI | 9 | 47 | 0.04 | 357840 | NO | 326900 | YES | 12292000 | YES | P54276 |
| **HYEITKQDI** | HYEITKQDI | 9 | 133.7 | 0.125 | 6781400 | YES | 1472600 | YES | 10157000 | YES | Q62245 |
| **HYFEDKENI** | HYFEDKENI | 9 | 169.8 | 0.15 | 3520100 | YES | 4116300 | YES | 572820 | YES | P53351 |
| **HYGESITNI** | HYGESITNI | 9 | 36.6 | 0.025 | 4370600 | YES | 983690 | YES | 55329000 | YES | Q80YA3 |
| **HYHQLLEKV** | HYHQLLEKV | 9 | 285.8 | 0.25 | 18213000 | YES | 663360 | YES | 8047100 | YES | Q68FL6 |
| **HYLDMNTVL** | HYLDM(+15.99)NTVL | 9 | 21.1 | 0.01 | 3941200 | YES | 42293000 | YES | 75605000 | NO | E9Q555 |
| **HYLDTTTLI** | HYLDTTTLI | 9 | 44.9 | 0.04 | 4970400 | NO | 12857000 | YES | 32580000 | YES | P47941 |
| **HYLHSLSQA** | HYLHSLSQA | 9 | 226.4 | 0.2 | 771260 | NO | 598500 | YES | 6547800 | YES | Q8K1X1 |
| **HYLPSYYHL** | HYLPSYYHL | 9 | 59.3 | 0.06 | 13251000 | YES | 3903400 | YES | 23000000 | YES | Q9DAR7 |
| **HYQNMKHAI** | HYQNM(+15.99)KHAI | 9 | 8.1 | 0.01 | 1213700 | NO | 263900 | YES | 33853000 | YES | Q8K0S9 |
| **HYQNMKHAI** | HYQNMKHAI | 9 | 8.1 | 0.01 | 396380 | NO | 113580 | YES | | YES | Q8K0S9 |
| **HYQQALTSA** | HYQQALTSA | 9 | 37.8 | 0.03 | 137920 | NO | 125550 | YES | 2987600 | YES | Q8C181 |
| **HYQSIGSTL** | HYQSIGSTL | 9 | 10 | 0.01 | 8890500 | YES | 1176300 | YES | 33462000 | YES | A3KGS3 |
| **HYSAIYSLL** | HYSAIYSLL | 9 | 202.4 | 0.175 | 3095800 | YES | 18774000 | YES | | YES | Q6P4S6 |
| **HYVAGLVGI** | HYVAGLVGI | 9 | 276.9 | 0.25 | 9046400 | YES | 2437200 | YES | 3603800 | YES | P53798 |
| **HYVITARAL** | HYVITARAL | 9 | 121.4 | 0.125 | 2100000 | YES | 996850 | YES | 11734000 | YES | Q9EQ61 |
| **HYWPVHNEL** | HYWPVHNEL | 9 | 36.4 | 0.025 | 2287500 | NO | 3738400 | NO | 103740000 | NO | P97471 |
| **HYYETPTGI** | HYYETPTGI | 9 | 43.5 | 0.04 | | YES | 1152500 | YES | 22383000 | YES | Q5NCF2 |
| **IFDRVLTEL** | IFDRVLTEL | 9 | 1876.2 | 1.2 | 90057000 | YES | 70323000 | YES | 880290000 | YES | P28700:P28704 |
| **IFIKIINTI** | IFIKIINTI | 9 | 136.9 | 0.125 | 1783600 | NO | 19772000 | YES | 69929000 | YES | E9Q7G0 |
| **IFLLTNNNL** | IFLLTNNNL | 9 | 1689 | 1.1 | 517410 | NO | 1545400 | NO | 256420 | NO | P46735 |

| | | | | | | | | | | | |
|---|---|---|---|---|---|---|---|---|---|---|---|
| **IFTSVRSEL** | IFTSVRSEL | 9 | 275.9 | 0.25 | 452130 | YES | 8862000 | YES | 177900000 | YES | Q3UWM4 |
| **IGIENIHYL** | IGIENIHYL | 9 | 11116.3 | 6.5 | | YES | 3897700 | YES | 12622000 | YES | Q9DB27 |
| **IGPTYYQRL** | IGPTYYQRL | 9 | 31544 | 36 | | YES | | YES | 2032500 | NO | Q8CFI7 |
| **ISPVNPVAI** | ISPVNPVAI | 9 | 27767 | 27 | | YES | | YES | 552350 | YES | Q91WT8 |
| **IYASSKDAI** | IYASSKDAI | 9 | 12.9 | 0.01 | 1598100 | YES | | YES | 67363000 | YES | P18760:P45591 |
| **IYDKIKTGL** | IYDKIKTGL | 9 | 324.2 | 0.3 | 54964000 | YES | | YES | 7706500 | YES | O70503 |
| **IYEETRGVL** | IYEETRGVL | 9 | 416 | 0.4 | 6790300 | YES | 3419700 | YES | 22551000 | YES | P62806 |
| **IYEGQITAV** | IYEGQITAV | 9 | 131.3 | 0.125 | 276650 | NO | | YES | | YES | Q8K440 |
| **IYERAISTL** | IYERAISTL | 9 | 20.9 | 0.01 | 5633500 | YES | 2001900 | YES | 37566000 | YES | Q99LI7 |
| **IYFPSVTGI** | IYFPSVTGI | 9 | 27.6 | 0.02 | 3868900 | YES | 2355800 | YES | 34081000 | YES | Q9WVL3:Q91V14 |
| **IYGDVISNI** | IYGDVISNI | 9 | 158.4 | 0.15 | 5502900 | YES | 3752500 | YES | 1843500 | YES | Q8CHW4 |
| **IYGGAHQTL** | IYGGAHQTL | 9 | 108.7 | 0.1 | 600170 | NO | 195090 | YES | 3038200 | YES | Q80YV4 |
| **IYGGLTSKV** | IYGGLTSKV | 9 | 840 | 0.7 | 606430 | NO | 96860 | YES | 796360 | YES | Q9WU78 |
| **IYHGLATLL** | IYHGLATLL | 9 | 80.6 | 0.08 | 5710800 | YES | 61807000 | YES | 51936000 | YES | Q9QZQ1 |
| **IYHPNVDKL** | IYHPNVDKL | 9 | 1042.5 | 0.8 | 1732200 | NO | 475400 | YES | 488630 | NO | P61089 |
| **IYKALQTAL** | IYKALQTAL | 9 | 54.4 | 0.05 | 497990 | NO | 2884200 | YES | 62823000 | YES | Q3U0V2 |
| **IYKDSSTFL** | IYKDSSTFL | 9 | 92.3 | 0.09 | 13480000 | YES | 12467000 | YES | 46644000 | YES | Q3UIK4 |
| **IYKGVIQAI** | IYKGVIQAI | 9 | 122.2 | 0.125 | 3535600 | YES | 48411000 | YES | 123410000 | YES | Q99P72 |
| **IYKNSISKI** | IYKNSISKI | 9 | 84.2 | 0.08 | 1266200 | NO | 600750 | NO | 5588600 | NO | Q8VCY6 |
| **IYKPGLSRL** | IYKPGLSRL | 9 | 148.8 | 0.15 | 1015400 | NO | 481980 | YES | | YES | P61406 |
| **IYLPAAQTM** | IYLPAAQTM(+15.99) | 9 | 26.8 | 0.02 | 11065000 | NO | 1321500 | NO | 3559900 | NO | P58281 |
| **IYNQVKQII** | IYNQVKQII | 9 | 51.1 | 0.04 | 3757000 | NO | 678260 | NO | 7437200 | NO | Q811D0 |
| **IYNRINNDL** | IYNRINNDL | 9 | 121.3 | 0.125 | | YES | | YES | 2543000 | YES | Q6P4T0 |
| **IYQDIRHEA** | IYQDIRHEA | 9 | 788 | 0.6 | 238160 | NO | 94679 | YES | 242900 | NO | P35821 |
| **IYQKVNERI** | IYQKVNERI | 9 | 26.4 | 0.015 | 91931 | NO | 539670 | YES | 6077200 | YES | Q60931 |
| **IYRELEQSI** | IYRELEQSI | 9 | 225.9 | 0.2 | 35338000 | YES | 13543000 | YES | 104510000 | YES | Q9WVE8 |
| **IYRPTINKL** | IYRPTINKL | 9 | 681.4 | 0.6 | 1599400 | NO | | YES | 2267300 | YES | Q7TMQ7 |
| **IYTSSVNRL** | IYTSSVNRL | 9 | 67.2 | 0.06 | 1314000 | YES | | YES | | YES | O55029 |
| **IYVEQKQYI** | IYVEQKQYI | 9 | 39.1 | 0.03 | 31403000 | YES | 6139700 | NO | 88326000 | YES | Q62018 |

| | | | | | | | | | | | |
|---|---|---|---|---|---|---|---|---|---|---|---|
| **IYWPNGLTI** | IYWPNGLTI | 9 | 76.7 | 0.07 | 3951400 | YES | 994390 | YES | 5098800 | YES | Q91VN0 |
| **IYWPNGLTL** | IYWPNGLTL | 9 | 168.5 | 0.15 | 3951400 | YES | 994390 | YES | 5098800 | YES | O88572 |
| **KALINADEL** | KALINADEL | 9 | 17699.8 | 12 | 194420 | YES | 3831800 | YES | 49146000 | YES | P16546 |
| **KAPDNRETL** | KAPDNRETL | 9 | 24861.9 | 21 | 374430 | YES | | YES | 5525400 | YES | Q3TDQ1 |
| **KAPTNEFYA** | KAPTNEFYA | 9 | 36086.8 | 55 | | YES | | YES | | YES | O35988 |
| **KFAEGITKI** | KFAEGITKI | 9 | 329.4 | 0.3 | 17105000 | YES | 4155600 | YES | 47263000 | YES | Q9CZU3 |
| **KFDKVLTAL** | KFDKVLTAL | 9 | 524.6 | 0.5 | 1558700 | NO | 3556800 | YES | 22754000 | YES | Q9QZS8 |
| **KFFQTPEGL** | KFFQTPEGL | 9 | 3524.9 | 2 | 12967000 | YES | 21036000 | YES | 33950000 | YES | Q60980 |
| **KFHPNGSTL** | KFHPNGSTL | 9 | 162.3 | 0.15 | 1256600 | YES | 1319400 | YES | 42569000 | YES | Q6PE01 |
| **KFIATLQYI** | KFIATLQYI | 9 | 67.3 | 0.06 | 15393000 | YES | 42028000 | YES | 200480000 | YES | Q9QUR6 |
| **KFIPLYSKV** | KFIPLYSKV | 9 | 674.1 | 0.6 | | YES | 320010 | YES | 1346500 | YES | P52479 |
| **KFITDCTGL** | KFITDC(+119.00)TGL | 9 | 301.1 | 0.3 | 3738800 | NO | 1749800 | NO | 28541000 | NO | Q8CCJ3 |
| **KFLAAGTHL** | KFLAAGTHL | 9 | 39.7 | 0.03 | 1826700 | YES | 5135600 | YES | 131970000 | YES | P14206 |
| **KFLPSSQEL** | KFLPSSQEL | 9 | 207.6 | 0.2 | 2362200 | NO | 11512000 | YES | 117980000 | YES | O70472 |
| **KFLQDKDFL** | KFLQDKDFL | 9 | 3521 | 2 | 20327000 | YES | 427820 | NO | 1275000 | NO | Q64471 |
| **KFNEVVSAL** | KFNEVVSAL | 9 | 410.2 | 0.4 | 909230 | NO | 1143200 | YES | | YES | Q64331 |
| **KFNPVETFL** | KFNPVETFL | 9 | 288.6 | 0.3 | 763950 | NO | 2062100 | YES | 1373000 | NO | Q6PAC3 |
| **KFNQILTAL** | KFNQILTAL | 9 | 184.4 | 0.175 | 4664800 | YES | | YES | 131450000 | YES | Q9QZK2 |
| **KFQGMISEL** | KFQGM(+15.99)ISEL | 9 | 141.5 | 0.15 | 9388900 | YES | 6474800 | YES | 5951800 | YES | P39061 |
| **KFQGMISEL** | KFQGMISEL | 9 | 141.5 | 0.15 | 6815500 | YES | 15583000 | YES | | YES | P39061 |
| **KFREDRSLL** | KFREDRSLL | 9 | 10221.8 | 6 | 312340 | NO | 328680 | NO | 41071000 | NO | Q4LDD4 |
| **KFREIQQEL** | KFREIQQEL | 9 | 8047.7 | 4.5 | 635480 | NO | 43470 | NO | 1905000 | NO | Q9QX47 |
| **KFSETATAI** | KFSETATAI | 9 | 113.1 | 0.125 | 761180 | NO | 695290 | YES | 16586000 | YES | Q80YV3 |
| **KFSQQYSTI** | KFSQQYSTI | 9 | 205 | 0.2 | 156180 | NO | 560390 | YES | 11405000 | YES | Q9QXZ0 |
| **KFTETLTNI** | KFTETLTNI | 9 | 183.5 | 0.175 | | YES | 147280000 | YES | 253400000 | YES | Q80SU7 |
| **KFTNSLTTV** | KFTNSLTTV | 9 | 74.3 | 0.07 | | YES | 2257500 | YES | 17519000 | YES | Q9D281 |
| **KFVDGVSTV** | KFVDGVSTV | 9 | 602.1 | 0.5 | 188300 | NO | 3277000 | YES | 57981000 | YES | Q8CG48 |
| **KFVSTRSLI** | KFVSTRSLI | 9 | 100.2 | 0.1 | 3495900 | NO | 11130000 | NO | 135200000 | YES | P56383 |
| **KFVTDIDEL** | KFVTDIDEL | 9 | 4542.4 | 2.5 | 2034800 | YES | 9059000 | YES | 1030900 | YES | Q64261 |

| | | | | | | | | | | | |
|---|---|---|---|---|---|---|---|---|---|---|---|
| **KIADTKSSI** | KIADTKSSI | 9 | 770 | 0.6 | 1840000 | NO | 38576 | YES | 1536100 | YES | Q8BPM0 |
| **KITESLSLL** | KITESLSLL | 9 | 8417.5 | 4.5 | 368210 | NO | | YES | | YES | P58281 |
| **KLGPAGTTI** | KLGPAGTTI | 9 | 461.8 | 0.4 | 6754100 | YES | | YES | 24397000 | YES | Q6GQV7 |
| **KLIANNTTV** | KLIANNTTV | 9 | 410.1 | 0.4 | 1811800 | YES | 297300 | YES | 2777100 | YES | Q9Z2N8 |
| **KLIESKENL** | KLIESKENL | 9 | 3387.3 | 2 | 123190 | NO | | YES | 5438200 | YES | Q61334 |
| **KLIESKHEV** | KLIESKHEV | 9 | 2078.9 | 1.3 | 745260 | YES | 98357 | YES | 1008100 | YES | P62257 |
| **KLIPGLNNL** | KLIPGLNNL | 9 | 3679.7 | 2.5 | 8751100 | YES | | YES | 2299700 | YES | Q9JLV2 |
| **KLKGTLSFI** | KLKGTLSFI | 9 | 6251.1 | 3.5 | 157840 | NO | | YES | | YES | O54774 |
| **KLLDIRSYL** | KLLDIRSYL | 9 | 2349.2 | 1.4 | 757470 | NO | 5622600 | YES | 7239600 | YES | P26516 |
| **KLLETKNEL** | KLLETKNEL | 9 | 2982.4 | 1.8 | 473070 | NO | | YES | 12559000 | YES | Q9JJA4 |
| **KLLPGFTTL** | KLLPGFTTL | 9 | 621.8 | 0.5 | 894110 | NO | 5541500 | YES | 40444000 | YES | Q3UQU0 |
| **KLLPSVTEL** | KLLPSVTEL | 9 | 1243.1 | 0.9 | 2801600 | YES | | YES | 47099000 | YES | Q80UV9 |
| **KLQEAFTKI** | KLQEAFTKI | 9 | 1665.7 | 1.1 | 439900 | NO | | YES | | YES | Q8VDR9 |
| **KLVEGRTHI** | KLVEGRTHI | 9 | 423.7 | 0.4 | 16709000 | YES | 2212100 | YES | 27276000 | YES | P01029 |
| **KLVPLKETI** | KLVPLKETI | 9 | 994.6 | 0.8 | 11935000 | YES | 5163100 | YES | 18665000 | YES | P56480 |
| **KLVTTVTEI** | KLVTTVTEI | 9 | 439.2 | 0.4 | 3909300 | YES | 15372000 | YES | 302690000 | YES | P40124 |
| **KMPILISKI** | KM(+15.99)PILISKI | 9 | 16285.6 | 10 | 1390600 | YES | 111330 | YES | 7200300 | YES | Q99L88 |
| **KMTQLFTKV** | KM(+15.99)TQLFTKV | 9 | 5547.4 | 3 | 932730 | YES | 6418200 | YES | 25834000 | YES | Q61221 |
| **KMPILISKI** | KMPILISKI | 9 | 16285.6 | 10 | 1201300 | YES | 401030 | YES | 55761 | YES | Q99L88 |
| **KMTQLFTKV** | KMTQLFTKV | 9 | 5547.4 | 3 | 912440 | YES | 7513200 | YES | | YES | Q61221 |
| **KQLEDGRTL** | KQLEDGRTL | 9 | 8166.6 | 4.5 | | YES | | YES | 500360 | YES | P62983 |
| **KWSSLLQVI** | KWSSLLQVI | 9 | 5162.7 | 3 | 916820 | YES | | YES | 2138600 | YES | Q6A009 |
| **KYAHMINGF** | KYAHM(+15.99)INGF | 9 | 846.8 | 0.7 | 3595400 | YES | 1967500 | YES | 19030000 | YES | P46664 |
| **KYAHMINGF** | KYAHMINGF | 9 | 846.8 | 0.7 | 1240600 | YES | 2200200 | YES | | YES | P46664 |
| **KYAMMFAEL** | KYAM(+15.99)M(+15.99)FA EL | 9 | 58 | 0.05 | 151630 | NO | 1196500 | YES | 126590000 | NO | Q7TMY8 |
| **KYAPSGFYI** | KYAPSGFYI | 9 | 15.2 | 0.01 | | YES | 28637000 | YES | 72935000 | YES | O88342 |
| **KYATGENTV** | KYATGENTV | 9 | 12.1 | 0.01 | 4109600 | YES | | YES | 25514000 | YES | A2A8Z1 |
| **KYDEAASYI** | KYDEAASYI | 9 | 22.1 | 0.01 | 4288800 | YES | 24519000 | YES | 209880000 | YES | P08752 |
| **KYDELKNDL** | KYDELKNDL | 9 | 333.2 | 0.3 | 1723700 | YES | 2111100 | YES | 1871000 | YES | E9Q555 |

| | | | | | | | | | | | |
|---|---|---|---|---|---|---|---|---|---|---|---|
| **KYDEVLHMV** | KYDEVLHMV | 9 | 205.1 | 0.2 | 1251200 | NO | 601500 | YES | | YES | Q8CIE6 |
| **KYDPINSML** | KYDPINSM(+15.99)L | 9 | 64.3 | 0.06 | 37307000 | YES | 11325000 | YES | 42934000 | YES | Q91VC9 |
| **KYDPINSML** | KYDPINSML | 9 | 64.3 | 0.06 | 27689000 | YES | 16885000 | YES | | YES | Q91VC9 |
| **KYEAAGTLV** | KYEAAGTLV | 9 | 29.3 | 0.02 | 4103100 | YES | | YES | 6086800 | YES | Q9JIF7 |
| **KYEALKTYA** | KYEALKTYA | 9 | 42.5 | 0.04 | 1148800 | YES | 550810 | YES | 7120200 | YES | Q9D1R1 |
| **KYEELLQVI** | KYEELLQVI | 9 | 90.6 | 0.09 | 2116500 | NO | 2639500 | YES | 7138300 | YES | P97478 |
| **KYEEVARKL** | KYEEVARKL | 9 | 163.3 | 0.15 | 1083200 | NO | 542080 | YES | 1251100 | NO | P21107:P58774 |
| **KYEHAFNSI** | KYEHAFNSI | 9 | 26.4 | 0.015 | 1492900 | NO | 511820 | NO | 1613400 | NO | Q8BZ60 |
| **KYEIIASDL** | KYEIIASDL | 9 | 98 | 0.09 | | YES | 3492000 | NO | 15829000 | YES | Q64435 |
| **KYEPIFQDI** | KYEPIFQDI | 9 | 97.7 | 0.09 | 2144900 | NO | 1920400 | YES | 13220000 | YES | Q64700 |
| **KYESVIATL** | KYESVIATL | 9 | 32.3 | 0.02 | 3530900 | NO | | YES | 601550 | YES | O35643 |
| **KYFEVPSVL** | KYFEVPSVL | 9 | 27.7 | 0.02 | 4198400 | YES | 23378000 | YES | 110500000 | YES | Q9QZB7 |
| **KYFKGLMHV** | KYFKGLM(+15.99)HV | 9 | 38 | 0.03 | 170640 | NO | 1580100 | NO | 8790700 | NO | Q9CRC9 |
| **KYFKMGDHV** | KYFKMGDHV | 9 | 16.2 | 0.01 | 503600 | NO | 265210 | YES | | YES | O55201 |
| **KYFPSRVSI** | KYFPSRVSI | 9 | 12.5 | 0.01 | 64830000 | YES | 29452000 | YES | 411230000 | YES | Q6PEB6 |
| **KYGIPFSRI** | KYGIPFSRI | 9 | 225.8 | 0.2 | 1000600 | NO | | YES | | YES | Q91VY5 |
| **KYGPVVSLL** | KYGPVVSLL | 9 | 107.2 | 0.1 | 1775300 | YES | 3917000 | YES | 78927000 | YES | P97343 |
| **KYGVTMEQI** | KYGVTM(+15.99)EQI | 9 | 75.1 | 0.07 | 1140300 | NO | 2601100 | YES | 21818000 | YES | Q9D7V2:Q9D0E3 |
| **KYGVVLDEI** | KYGVVLDEI | 9 | 80.5 | 0.08 | 362310000 | YES | 196050000 | YES | 508340000 | YES | O89079 |
| **KYHGNVTLL** | KYHGNVTLL | 9 | 130 | 0.125 | 1022700 | YES | 752400 | YES | 2604700 | YES | P19096 |
| **KYHLLLQEI** | KYHLLLQEI | 9 | 40.7 | 0.03 | 868430 | NO | 2609100 | YES | 64714000 | YES | Q4VAC9 |
| **KYHLLLQEL** | KYHLLLQEL | 9 | 88.4 | 0.08 | 868430 | NO | 2609100 | YES | 64714000 | YES | P27870:Q9R0C8 |
| **KYHSQYHTV** | KYHSQYHTV | 9 | 16.4 | 0.01 | 458080 | NO | 101700 | YES | 11023000 | YES | E9Q3L2 |
| **KYIAEKTEF** | KYIAEKTEF | 9 | 189.1 | 0.175 | 119740 | NO | | YES | 467640 | YES | Q8BL99 |
| **KYIDQKFVL** | KYIDQKFVL | 9 | 226.7 | 0.2 | 30001000 | NO | 14492000 | YES | 52504000 | YES | Q60996:Q61151 |
| **KYIDVGNTI** | KYIDVGNTI | 9 | 9.4 | 0.01 | 127610 | YES | | YES | | YES | A2RTF1 |
| **KYIEDKDVF** | KYIEDKDVF | 9 | 3421.8 | 2 | 4346200 | NO | 127640 | NO | 443310 | NO | Q9WTX6 |
| **KYIEGVSDF** | KYIEGVSDF | 9 | 585.7 | 0.5 | 50458000 | YES | 1585000 | YES | 250520 | NO | P28575 |
| **KYIHSADII** | KYIHSADII | 9 | 23.6 | 0.015 | 5792100 | YES | 1989400 | YES | 76240000 | YES | P47811 |

| | | | | | | | | | | | |
|---|---|---|---|---|---|---|---|---|---|---|---|
| **KYIHSANVL** | KYIHSANVL | 9 | 19.8 | 0.01 | 128430000 | YES | 46794000 | YES | 569820000 | YES | P63085:Q63844:Q61532:Q6P5G0 |
| **KYIKSHYKV** | KYIKSHYKV | 9 | 56.5 | 0.05 | 2691500 | NO | 24387 | NO | 691450 | NO | P10922 |
| **KYIPAARHL** | KYIPAARHL | 9 | 15.8 | 0.01 | 146950 | YES | 320280 | YES | 3086300 | YES | O54692 |
| **KYIPNRGPL** | KYIPNRGPL | 9 | 27.1 | 0.02 | | YES | 477050 | YES | 4455800 | YES | Q6ZWV3 |
| **KYITNTDVL** | KYITNTDVL | 9 | 57.9 | 0.05 | 4298900 | YES | | YES | 2729200 | YES | Q9CXE7 |
| **KYKASENAI** | KYKASENAI | 9 | 16.9 | 0.01 | 3994900 | YES | 176790 | YES | 24457000 | YES | P84091 |
| **KYKASIAAL** | KYKASIAAL | 9 | 78.8 | 0.08 | 753930 | NO | 10417000 | NO | 127400000 | NO | Q8VDD5 |
| **KYKDIYTEL** | KYKDIYTEL | 9 | 91.9 | 0.09 | 55831000 | YES | 99603000 | YES | 1278200000 | YES | P97434 |
| **KYKDSETRL** | KYKDSETRL | 9 | 84.7 | 0.08 | 12132000 | YES | | YES | 2718400 | YES | Q91XD7 |
| **KYKELGEKL** | KYKELGEKL | 9 | 502.3 | 0.5 | 668000 | NO | 967910 | YES | 7419300 | YES | P27773 |
| **KYLATLERL** | KYLATLERL | 9 | 39.1 | 0.03 | 687950 | YES | 559870 | YES | 7942600 | YES | Q9R117 |
| **KYLENPNAL** | KYLENPNAL | 9 | 50.9 | 0.04 | 25230000 | YES | 8442500 | YES | 148310000 | YES | Q8BX02 |
| **KYLGLMENL** | KYLGLMENL | 9 | 82.1 | 0.08 | 387670 | NO | 25237000 | YES | | YES | Q9WTI7 |
| **KYLGQLHYL** | KYLGQLHYL | 9 | 40 | 0.03 | 1780700 | NO | 6151600 | YES | 15720000 | YES | Q6PB44 |
| **KYLGQLTSI** | KYLGQLTSI | 9 | 10.8 | 0.01 | 2096600 | YES | 2465400 | YES | 198840000 | YES | Q8K2Y9 |
| **KYLPLLDRA** | KYLPLLDRA | 9 | 795.7 | 0.6 | 1887300 | YES | | YES | 3155900 | YES | Q8C4V4 |
| **KYLQNDLYI** | KYLQNDLYI | 9 | 49.3 | 0.04 | 4228000 | NO | 17689000 | YES | 64621000 | YES | O89050 |
| **KYLQSKEDL** | KYLQSKEDL | 9 | 27.2 | 0.02 | 364980 | NO | 344200 | YES | 6356500 | YES | Q6ZWQ0 |
| **KYLSDNVHL** | KYLSDNVHL | 9 | 101.4 | 0.1 | 15529000 | YES | 9586500 | YES | 172790000 | YES | Q61081 |
| **KYLSQKNVV** | KYLSQKNVV | 9 | 23.2 | 0.01 | 455110 | NO | | YES | 6687300 | YES | A2AN08 |
| **KYLSVQGQL** | KYLSVQGQL | 9 | 31.5 | 0.02 | 20312000 | YES | 21373000 | YES | 67291000 | YES | Q791T5 |
| **KYLVGQRLV** | KYLVGQRLV | 9 | 295.2 | 0.3 | 2283300 | YES | 553460 | YES | 6194800 | YES | Q6ZQ89 |
| **KYMEPLQEI** | KYM(+15.99)EPLQEI | 9 | 28 | 0.02 | 586610 | NO | 557240 | YES | 8041700 | YES | Q80TY5 |
| **KYMETIEKL** | KYM(+15.99)ETIEKL | 9 | 78 | 0.08 | 1769600 | YES | 1033600 | YES | 87331000 | YES | Q3TDD9 |
| **KYMEPLQEI** | KYMEPLQEI | 9 | 28 | 0.02 | 2141700 | NO | 1091600 | YES | | YES | Q80TY5 |
| **KYMETIEKL** | KYMETIEKL | 9 | 78 | 0.08 | 151640 | YES | 2155900 | YES | | YES | Q3TDD9 |
| **KYNDVSHQL** | KYNDVSHQL | 9 | 51.5 | 0.04 | 1879100 | YES | 619550 | YES | 8231400 | YES | O09053 |
| **KYNGAVNEI** | KYNGAVNEI | 9 | 34.8 | 0.025 | 1072200 | NO | 447080 | YES | 22999000 | YES | Q3URD3 |
| **KYNILIATL** | KYNILIATL | 9 | 137.3 | 0.125 | 359210 | NO | | YES | 7498900 | YES | Q99J21 |

| | | | | | | | | | | | |
|---|---|---|---|---|---|---|---|---|---|---|---|
| **KYNIMLVRL** | KYNIM(+15.99)LVRL | 9 | 186.1 | 0.175 | 10205000 | YES | 11276000 | YES | 135910000 | YES | P42932 |
| **KYNIMLVRL** | KYNIMLVRL | 9 | 186.1 | 0.175 | 6567900 | YES | 15476000 | YES | | YES | P42932 |
| **KYNNGSTEL** | KYNNGSTEL | 9 | 16.3 | 0.01 | | YES | | YES | 27875000 | YES | Q7TSH2 |
| **KYNPDKHYI** | KYNPDKHYI | 9 | 29.8 | 0.02 | 221180 | NO | 1268200 | YES | 11074000 | YES | Q61194 |
| **KYNRGLTVV** | KYNRGLTVV | 9 | 24.5 | 0.015 | 64903000 | YES | 2547500 | YES | 31519000 | YES | Q920E5 |
| **KYPPSATTL** | KYPPSATTL | 9 | 19.7 | 0.01 | 695690 | NO | 69661 | YES | 5573200 | YES | Q7TMY8 |
| **KYQAAMERL** | KYQAAMERL | 9 | 24.9 | 0.015 | 6411400 | YES | | YES | | YES | Q8BGC4 |
| **KYQDILNEI** | KYQDILNEI | 9 | 27.8 | 0.02 | 1280700000 | YES | 31909000 | YES | 660100000 | YES | Q3UQ44 |
| **KYQDSLQSI** | KYQDSLQSI | 9 | 14.7 | 0.01 | 13373000 | YES | 3150000 | YES | 140550000 | YES | Q6ZWR6 |
| **KYQEALDVI** | KYQEALDVI | 9 | 31 | 0.02 | 4777600 | YES | 25132000 | YES | 61705000 | YES | Q8BWZ3 |
| **KYQEVTNNL** | KYQEVTNNL | 9 | 22.6 | 0.01 | 85935000 | YES | 50002000 | YES | 1585700000 | YES | Q60865 |
| **KYQHTGAVL** | KYQHTGAVL | 9 | 130.3 | 0.125 | 100780 | NO | 209010 | YES | 3414300 | YES | Q9WVA3 |
| **KYQIAVTKV** | KYQIAVTKV | 9 | 33.9 | 0.025 | 7111400 | YES | 8255600 | YES | 83978000 | YES | Q9CU62 |
| **KYQKGFSLW** | KYQKGFSLW | 9 | 519.5 | 0.5 | 10105000 | YES | | YES | 130820 | NO | Q91V04 |
| **KYQKTFTVI** | KYQKTFTVI | 9 | 9.3 | 0.01 | 1930500 | YES | 5090500 | YES | 87079000 | YES | O35459 |
| **KYQRILERL** | KYQRILERL | 9 | 43.2 | 0.04 | 5444200 | NO | 1567400 | NO | 10802000 | NO | P58281 |
| **KYQRLLHEV** | KYQRLLHEV | 9 | 24.3 | 0.015 | 15740000 | YES | | YES | 20266000 | NO | Q99KJ8 |
| **KYQSQNEKL** | KYQSQNEKL | 9 | 47.3 | 0.04 | 1170000 | YES | | YES | 21778000 | YES | Q921T2 |
| **KYQTVIDDI** | KYQTVIDDI | 9 | 16 | 0.01 | 4300500 | YES | 14042000 | YES | 30874000 | YES | P53798 |
| **KYQVSSNGI** | KYQVSSNGI | 9 | 15.8 | 0.01 | | YES | 2063700 | YES | 43548000 | YES | Q3UQN2 |
| **KYRHVDGNL** | KYRHVDGNL | 9 | 547.4 | 0.5 | 211350 | NO | | YES | 4324800 | YES | P49962 |
| **KYSEVFEAI** | KYSEVFEAI | 9 | 53.5 | 0.05 | 7639900 | YES | 34091000 | YES | 77407000 | YES | Q60737:O54833 |
| **KYSGVLSSI** | KYSGVLSSI | 9 | 18.1 | 0.01 | 9785200 | YES | 38576000 | YES | 193160000 | YES | Q791T5 |
| **KYSNVIQLL** | KYSNVIQLL | 9 | 327.6 | 0.3 | 415640 | NO | | YES | 11216000 | YES | Q9Z329 |
| **KYSPQRVGL** | KYSPQRVGL | 9 | 866.1 | 0.7 | 5714400 | YES | 2222000 | YES | 33791000 | YES | Q9QXB9 |
| **KYSSLYENL** | KYSSLYENL | 9 | 110.4 | 0.1 | 275680 | NO | 1248500 | YES | 9424500 | YES | Q8K4E0 |
| **KYSTSLSWI** | KYSTSLSWI | 9 | 13.6 | 0.01 | 363830 | NO | | YES | 19394000 | YES | Q8K0F1 |
| **KYSVLDSPL** | KYSVLDSPL | 9 | 49.4 | 0.04 | 2082000 | YES | | YES | 765730 | YES | P26187 |
| **KYTAQNREL** | KYTAQNREL | 9 | 74.9 | 0.07 | 1526200 | NO | | YES | 1766300 | NO | Q61817 |

| | | | | | | | | | | | |
|---|---|---|---|---|---|---|---|---|---|---|---|
| **KYTEGVQSL** | KYTEGVQSL | 9 | 68.4 | 0.07 | 1101800 | YES | | YES | 1775000 | YES | Q920B9 |
| **KYTGNASAL** | KYTGNASAL | 9 | 44.5 | 0.04 | 37914000 | YES | | YES | | YES | P07759:Q03734:Q91WP6 |
| **KYVAVYNLI** | KYVAVYNLI | 9 | 23.4 | 0.01 | 3822800 | YES | 37984000 | YES | 454220000 | YES | Q80U87 |
| **KYVENFGLI** | KYVENFGLI | 9 | 391.8 | 0.4 | 2346700 | NO | 3013100 | YES | 123840000 | YES | Q9CYN2 |
| **KYVNSIWDL** | KYVNSIWDL | 9 | 60.7 | 0.06 | 11665000 | YES | 19953000 | YES | 12521000 | YES | Q9JLV5 |
| **KYVPLVTGL** | KYVPLVTGL | 9 | 61.8 | 0.06 | 20549000 | YES | 28772000 | YES | 206470000 | YES | Q8BMI0 |
| **KYVPQQDAL** | KYVPQQDAL | 9 | 105 | 0.1 | 179750 | NO | 440510 | YES | 4584800 | YES | F8VPU2 |
| **KYVYVVTEL** | KYVYVVTEL | 9 | 25.9 | 0.015 | 4426100 | YES | 124920000 | YES | 354330000 | YES | P18654 |
| **KYWKGQHVI** | KYWKGQHVI | 9 | 41.4 | 0.03 | 54139000 | YES | 1328800 | NO | 51120000 | YES | Q9D1P2 |
| **KYWPDEYAL** | KYWPDEYAL | 9 | 480.1 | 0.4 | | YES | 2985700 | YES | 5958500 | YES | P35235 |
| **KYYVQLEQL** | KYYVQLEQL | 9 | 116 | 0.125 | 1986400 | YES | 78528000 | YES | 70394000 | YES | Q9Z131 |
| **LAGNEQVTR** | LAGNEQVTR | 9 | 43453.5 | 99 | 365890 | YES | | YES | 7393.1 | YES | P56183 |
| **LFQPVISQV** | LFQPVISQV | 9 | 714.6 | 0.6 | 5810700 | YES | 1923000 | YES | | YES | O88895 |
| **LLLPGELAK** | LLLPGELAK | 9 | 32390.6 | 39 | 5723900 | YES | 3390500 | YES | 2995200 | YES | Q6ZWY9:P10853:Q64525:Q8CGP1:Q8CGP0:Q9D2U9:Q64524 |
| **LYDPVISKL** | LYDPVISKL | 9 | 758.6 | 0.6 | 31740000 | YES | 75829000 | YES | 301920000 | YES | Q99L90 |
| **LYEAVREVL** | LYEAVREVL | 9 | 183.3 | 0.175 | 6988300 | YES | 10532000 | YES | 6061600 | YES | P53026 |
| **LYERLKTEL** | LYERLKTEL | 9 | 37.4 | 0.025 | 168420000 | YES | 18590000 | YES | 199370000 | YES | Q9CQF9 |
| **LYHEAGQQL** | LYHEAGQQL | 9 | 120.5 | 0.125 | 677400 | YES | 1332700 | YES | 8388700 | YES | Q99JG7 |
| **LYIDSRQSL** | LYIDSRQSL | 9 | 42.5 | 0.04 | | YES | | YES | 24405000 | YES | Q8BUR3 |
| **LYIGHLTAL** | LYIGHLTAL | 9 | 113.6 | 0.125 | 2436900 | NO | 4715300 | YES | 17770000 | YES | Q99MI1 |
| **LYIPSVDLL** | LYIPSVDLL | 9 | 300.3 | 0.3 | 787740 | YES | | YES | | YES | P97432 |
| **LYIQAQNNL** | LYIQAQNNL | 9 | 70.5 | 0.07 | 77948 | NO | 1076800 | NO | 6049600 | YES | Q6ZPU9 |
| **LYKDVRNLL** | LYKDVRNLL | 9 | 995.1 | 0.8 | 4034300 | NO | | YES | 4603000 | YES | Q80UU1 |
| **LYKEQLAKL** | LYKEQLAKL | 9 | 5054.5 | 3 | 318280 | NO | 725770 | YES | 1293900 | YES | Q8VDD5 |
| **LYKESLTKL** | LYKESLTKL | 9 | 181.6 | 0.175 | 1921000 | NO | 1132600 | YES | 2082000 | YES | Q61879 |
| **LYLDNRKEI** | LYLDNRKEI | 9 | 226.3 | 0.2 | 2470000 | YES | | YES | | YES | Q8BXC6 |
| **LYLPNKAET** | LYLPNKAET | 9 | 3577.1 | 2.5 | 4492200 | YES | 515710 | YES | 4265500 | YES | Q8BFQ9 |
| **LYQEVFGRL** | LYQEVFGRL | 9 | 281.9 | 0.25 | 4065400 | YES | 39008000 | YES | 45459000 | YES | Q61985 |

| | | | | | | | | | | | |
|---|---|---|---|---|---|---|---|---|---|---|---|
| **LYQNQRAVL** | LYQNQRAVL | 9 | 387.5 | 0.4 | 580370 | NO | 496710 | YES | 3739100 | YES | Q921M4 |
| **LYQPSAESL** | LYQPSAESL | 9 | 24.2 | 0.015 | 5001800 | NO | | YES | | YES | Q9QZR0 |
| **LYQPTGGQL** | LYQPTGGQL | 9 | 90.4 | 0.09 | 29885000 | YES | 34273000 | YES | 37939000 | YES | P36371 |
| **LYRQSLEII** | LYRQSLEII | 9 | 445.1 | 0.4 | 53423000 | YES | 85278000 | YES | 96446000 | YES | P97287 |
| **LYSEQKTQL** | LYSEQKTQL | 9 | 130.2 | 0.125 | 1160400 | YES | 1332100 | YES | 13755000 | YES | Q8K1N2 |
| **LYSPVRSKL** | LYSPVRSKL | 9 | 119.2 | 0.125 | | YES | 617240 | YES | 4022100 | YES | Q9ERA6 |
| **LYVPALSAL** | LYVPALSAL | 9 | 32.3 | 0.02 | | YES | 3854600 | YES | 44526000 | YES | P27600 |
| **MFIEDLHNL** | M(+15.99)FIEDLHNL | 9 | 10516.5 | 6 | 2855600 | NO | 507840 | YES | 4056700 | YES | Q9CPY1 |
| **MLPSILNQL** | M(+15.99)LPSILNQL | 9 | 28205.5 | 28 | 12821000 | YES | 606170 | YES | 3272700 | YES | Q64152 |
| **MFIEDLHNL** | MFIEDLHNL | 9 | 10516.5 | 6 | 3402200 | NO | 2370400 | YES | | YES | Q9CPY1 |
| **MLPSILNQL** | MLPSILNQL | 9 | 28205.5 | 28 | 4195100 | YES | 394980 | YES | | YES | Q64152 |
| **MYNSVSQRL** | MYNSVSQRL | 9 | 94.3 | 0.09 | 671790 | YES | 541010 | YES | | YES | Q9DBC3 |
| **NFIGTKTVI** | NFIGTKTVI | 9 | 163.5 | 0.15 | 103190000 | NO | 12253000 | NO | 70647000 | YES | Q9R1J0 |
| **NFNPTVNYI** | NFNPTVNYI | 9 | 567.2 | 0.5 | 25704000 | YES | 12527000 | YES | 6265200 | YES | Q9DCF9 |
| **NLLTTRNYI** | NLLTTRNYI | 9 | 391.4 | 0.4 | 7307000 | NO | 1899400 | NO | 2452600 | YES | Q9DCV3 |
| **NSIRNLDTI** | NSIRNLDTI | 9 | 5225.2 | 3 | 2226200 | YES | 1121900 | YES | 25661000 | YES | P28658 |
| **NYARGHYTI** | NYARGHYTI | 9 | 23.3 | 0.01 | 6186800 | YES | 1617100 | YES | 5455700 | YES | P68373:P05213:P68368 |
| **NYDDIRTEL** | NYDDIRTEL | 9 | 290.9 | 0.3 | 5010100 | NO | 3111000 | YES | 68593000 | YES | Q3UMY5 |
| **NYFPSKQDI** | NYFPSKQDI | 9 | 25.9 | 0.015 | 82847000 | YES | 19257000 | YES | 362780000 | YES | P27600 |
| **NYFYDQQRI** | NYFYDQQRI | 9 | 788.7 | 0.6 | 570230 | YES | | YES | | YES | O70126 |
| **NYGDLLQTV** | NYGDLLQTV | 9 | 414.4 | 0.4 | 7639400 | YES | | YES | 513110 | YES | Q6GQT6 |
| **NYISGIQTI** | NYISGIQTI | 9 | 12.2 | 0.01 | 38469000 | YES | 51459000 | YES | 118330000 | YES | Q921M3 |
| **NYITPQTQI** | NYITPQTQI | 9 | 14.8 | 0.01 | 2996100 | NO | | YES | 2296600 | YES | Q5DTW7 |
| **NYLDIKGLL** | NYLDIKGLL | 9 | 615.9 | 0.5 | 104970000 | YES | 82733000 | YES | 125160000 | YES | Q9WTX5 |
| **NYLFSASAI** | NYLFSASAI | 9 | 11.8 | 0.01 | | YES | | YES | 6214700 | YES | Q68FL6 |
| **NYLPAINGI** | NYLPAINGI | 9 | 60.4 | 0.06 | 101270000 | YES | 22833000 | YES | 43986000 | YES | Q9CQC9 |
| **NYNSVNTRM** | NYNSVNTRM(+15.99) | 9 | 32.1 | 0.02 | 92792 | NO | 46245 | NO | 708110 | NO | P70452 |
| **NYQEALRYI** | NYQEALRYI | 9 | 27.1 | 0.02 | 13753000 | YES | 26366000 | YES | 47681000 | YES | Q91W86 |
| **NYQNVVHKL** | NYQNVVHKL | 9 | 106.8 | 0.1 | 556950 | YES | 666130 | YES | 1934300 | YES | A2A791 |

| | | | | | | | | | | | |
|---|---|---|---|---|---|---|---|---|---|---|---|
| **NYQPAGIAV** | NYQPAGIAV | 9 | 81 | 0.08 | 2361200 | NO | | YES | 3066300 | YES | Q6ZQF0 |
| **NYTNTPSVI** | NYTNTPSVI | 9 | 56.7 | 0.05 | 1437000 | NO | 2224400 | YES | 6353400 | YES | A2A6Q5 |
| **NYVDLVSSL** | NYVDLVSSL | 9 | 121.5 | 0.125 | 2013000 | YES | | YES | | YES | Q9WUR2 |
| **NYVNGKTFL** | NYVNGKTFL | 9 | 25.5 | 0.015 | 100230000 | YES | 34093000 | YES | 354240000 | YES | P49722 |
| **NYVRAGTLI** | NYVRAGTLI | 9 | 19.4 | 0.01 | 18874000 | YES | 2486300 | YES | 36332000 | YES | Q924Z4 |
| **NYYGSLTQA** | NYYGSLTQA | 9 | 219 | 0.2 | 934120 | NO | 278990 | NO | 2805000 | YES | Q9JHU9 |
| **NYYPVNTRI** | NYYPVNTRI | 9 | 12.4 | 0.01 | 8190400 | YES | 6059700 | YES | 353920000 | YES | O09159 |
| **NYYSSRTLL** | NYYSSRTLL | 9 | 15.8 | 0.01 | | YES | 567760 | YES | 11550000 | YES | Q6P8H8 |
| **PYFAGISAL** | PYFAGISAL | 9 | 40.3 | 0.03 | 249430 | NO | | YES | 1503800 | YES | Q9CZ42 |
| **PYFPIVNFL** | PYFPIVNFL | 9 | 220.5 | 0.2 | | YES | | YES | 2414600 | NO | Q8C2E7 |
| **PYIASGNNL** | PYIASGNNL | 9 | 33.4 | 0.025 | 30457000 | YES | 98142 | NO | | YES | Q61586 |
| **PYIESNSKL** | PYIESNSKL | 9 | 67.1 | 0.06 | | YES | | YES | | YES | Q9D0V8 |
| **PYLPSAHRV** | PYLPSAHRV | 9 | 68.2 | 0.07 | 7184300 | YES | | YES | | YES | P83887 |
| **PYLPSGESL** | PYLPSGESL | 9 | 53 | 0.05 | 4785800 | NO | | YES | 785550 | YES | Q9JKV1 |
| **QEKVTYQEL** | QEKVTYQEL | 9 | 23825.1 | 19 | 1022000 | YES | | YES | 650930 | YES | Q8R0W0 |
| **QFEKALTQI** | QFEKALTQI | 9 | 478.7 | 0.4 | 9099700 | YES | | YES | 14132000 | YES | Q91VE6 |
| **QFISVFSNL** | QFISVFSNL | 9 | 484.8 | 0.4 | 178640 | NO | | YES | 2993800 | YES | Q80ZJ6 |
| **QFITSVTAL** | QFITSVTAL | 9 | 91.1 | 0.09 | 1137700 | NO | 2566200 | YES | 26642000 | YES | Q8BGA7 |
| **QFLPDNINI** | QFLPDNINI | 9 | 9574.3 | 5.5 | 1175700 | NO | 763660 | NO | 1922100 | NO | Q6PIP5 |
| **QFNSSLHNI** | QFNSSLHNI | 9 | 644.9 | 0.5 | | YES | 2434100 | YES | 91158000 | YES | O35166 |
| **QILSDFPKL** | QILSDFPKL | 9 | 33951.5 | 44 | 585370 | YES | | YES | 9020200 | YES | Q99ME9 |
| **QWSQLKEQI** | QWSQLKEQI | 9 | 10837.8 | 6 | 1447000 | NO | | YES | 1543600 | NO | P11031 |
| **QYASNITSV** | QYASNITSV | 9 | 33.1 | 0.025 | 4655700 | YES | | YES | 1080500 | YES | Q5SWU9:E9Q4Z2 |
| **QYHDILHAL** | QYHDILHAL | 9 | 231 | 0.25 | 4925800 | NO | 1568600 | YES | 1037400 | NO | Q9D0K0 |
| **QYLENLEKL** | QYLENLEKL | 9 | 1517.9 | 1 | 1954700 | NO | 2922100 | YES | 7450300 | YES | Q8BKX6 |
| **QYLKMLQKL** | QYLKMLQKL | 9 | 80.5 | 0.08 | 686010 | NO | 703380 | NO | 98616 | YES | Q8BHG1 |
| **QYLKNQTVL** | QYLKNQTVL | 9 | 123.9 | 0.125 | 13182000 | YES | 5072100 | NO | 41298000 | NO | Q8BM75 |
| **QYNAGGLTV** | QYNAGGLTV | 9 | 111.6 | 0.125 | 2273000 | NO | | YES | 776120 | YES | P97493 |
| **QYNKLRNLL** | QYNKLRNLL | 9 | 220.1 | 0.2 | 2367500 | NO | 2643500 | YES | 38004000 | NO | Q8C0S1 |

| | | | | | | | | | | | |
|---|---|---|---|---|---|---|---|---|---|---|---|
| **QYNPSRQTL** | QYNPSRQTL | 9 | 24.2 | 0.015 | 60095000 | YES | 9680400 | YES | 112880000 | YES | Q9ESV0 |
| **QYNPVKQQL** | QYNPVKQQL | 9 | 73.3 | 0.07 | 204930000 | YES | | YES | 103070000 | YES | Q9DBL7 |
| **QYQANASQL** | QYQANASQL | 9 | 37.6 | 0.03 | 98457 | NO | | YES | 8069000 | YES | B1AVY7 |
| **QYQNIKNNL** | QYQNIKNNL | 9 | 53.4 | 0.05 | 808520 | NO | 1826500 | YES | 32689000 | YES | Q6ZWQ0 |
| **QYQQIINRL** | QYQQIINRL | 9 | 124.7 | 0.125 | 681010 | NO | 15087000 | YES | 1746500 | YES | Q5I1X5 |
| **QYQQSQHNL** | QYQQSQHNL | 9 | 67.2 | 0.06 | 1757600 | YES | 52659 | YES | 4924400 | YES | E9Q4N7 |
| **QYQSLLRSL** | QYQSLLRSL | 9 | 48.6 | 0.04 | 173280 | NO | 15714000 | YES | 17045000 | YES | Q91ZU6 |
| **QYRDTQTSI** | QYRDTQTSI | 9 | 138.7 | 0.125 | 243490 | YES | 980200 | YES | 43717000 | YES | P47758 |
| **QYSKVLNEL** | QYSKVLNEL | 9 | 168.5 | 0.15 | 17173000 | YES | 22573000 | YES | 163590000 | YES | Q9EP71 |
| **QYSTGKTTF** | QYSTGKTTF | 9 | 98.2 | 0.09 | 635610 | YES | | YES | 5441800 | YES | Q9QXY6:Q9WVK4:Q9EQP2 |
| **QYVCQQTGL** | QYVC(+119.00)QQTGL | 9 | 346.4 | 0.3 | 2121300 | NO | 468880 | NO | 12809000 | NO | Q8R404 |
| **QYVDFYSQL** | QYVDFYSQL | 9 | 1235.5 | 0.9 | 619310 | NO | 621140 | YES | 464230 | NO | P35689 |
| **QYVSAFSKL** | QYVSAFSKL | 9 | 98.8 | 0.1 | 9197900 | YES | 16079000 | YES | 49982000 | YES | Q99JB2 |
| **QYWTTVSSL** | QYWTTVSSL | 9 | 25.3 | 0.015 | 2705800 | NO | | YES | 10164000 | YES | P28575 |
| **RFFESYHEV** | RFFESYHEV | 9 | 1508.5 | 1 | 980820 | NO | | YES | 3103500 | NO | Q9D8B4 |
| **RFKDDITTI** | RFKDDITTI | 9 | 2938.2 | 1.8 | 2759000 | YES | | YES | 13776000 | YES | Q8C147 |
| **RFNPSISMI** | RFNPSISMI | 9 | 242.1 | 0.25 | | YES | 4923300 | YES | | YES | Q9D4H8 |
| **RLLASKSLL** | RLLASKSLL | 9 | 2952.7 | 1.8 | | YES | 2327700 | YES | 10469000 | YES | Q9CPW5 |
| **RLVPSVNGI** | RLVPSVNGI | 9 | 10613 | 6 | | YES | | YES | 535210 | YES | Q922D8 |
| **RNYEYLIRL** | RNYEYLIRL | 9 | 31765.6 | 37 | | YES | | YES | 4510200 | NO | Q8R3L2 |
| **RSIKNVTEL** | RSIKNVTEL | 9 | 6909.6 | 4 | | YES | 361630 | YES | 7855200 | YES | Q99K95 |
| **RSIQNAQFL** | RSIQNAQFL | 9 | 11914.5 | 7 | | YES | | YES | 3446500 | YES | P24638 |
| **RSLLLLAPL** | RSLLLLAPL | 9 | 17348.4 | 11 | | YES | | YES | | YES | Q9R0E2 |
| **RVTPTRTEI** | RVTPTRTEI | 9 | 1494 | 1 | 1483800 | NO | 730240 | NO | 8617100 | NO | P62908 |
| **RYAALRELI** | RYAALRELI | 9 | 48.8 | 0.04 | 4083500 | NO | 4094600 | NO | 108460000 | YES | Q9DBT3 |
| **RYAPSLHEL** | RYAPSLHEL | 9 | 34.4 | 0.025 | 10022000 | YES | | YES | 750970 | NO | Q66JZ4 |
| **RYASINTHL** | RYASINTHL | 9 | 12 | 0.01 | 568360 | YES | 6489400 | YES | 130560000 | YES | Q9DC28:Q9JMK2 |
| **RYFPVFEKI** | RYFPVFEKI | 9 | 89.4 | 0.09 | 3365400 | YES | 477960 | YES | 2091000 | NO | P24472 |
| **RYHAALAVI** | RYHAALAVI | 9 | 63.1 | 0.06 | 4402400 | YES | 3467200 | YES | 9075800 | YES | Q9JJW0 |

| | | | | | | | | | | | |
|---|---|---|---|---|---|---|---|---|---|---|---|
| **RYIANTVEL** | RYIANTVEL | 9 | 268.6 | 0.25 | 89638000 | YES | 25714000 | YES | 80816000 | YES | Q9DBG6 |
| **RYIENNSVV** | RYIENNSVV | 9 | 84.9 | 0.08 | 503980 | NO | | YES | 4579100 | YES | O88566 |
| **RYIQGILNV** | RYIQGILNV | 9 | 180.8 | 0.175 | 5557300 | NO | 506510 | YES | 563530 | YES | Q8C0S1 |
| **RYISQTQGL** | RYISQTQGL | 9 | 152.3 | 0.15 | 136150 | NO | | YES | 1910800 | YES | O88904:Q9ERH7 |
| **RYKEGRVIL** | RYKEGRVIL | 9 | 1260.6 | 0.9 | 8817200 | NO | 3560300 | NO | 58635000 | NO | P62700 |
| **RYKGTLSML** | RYKGTLSM(+15.99)L | 9 | 122.3 | 0.125 | 3079600 | YES | 2217200 | YES | 25145000 | YES | Q9DB90 |
| **RYKGTLSML** | RYKGTLSML | 9 | 122.3 | 0.125 | 887480 | YES | 1466800 | YES | | YES | Q9DB90 |
| **RYKQLLTYI** | RYKQLLTYI | 9 | 19.7 | 0.01 | 16597000 | YES | 17099000 | YES | 280570000 | YES | P61202 |
| **RYLEQLHQL** | RYLEQLHQL | 9 | 120.3 | 0.125 | 52456000 | YES | 33871000 | YES | 131480000 | YES | P42227 |
| **RYLGLLENV** | RYLGLLENV | 9 | 222.8 | 0.2 | 9347800 | YES | 16641000 | YES | 8504200 | YES | P46735 |
| **RYLPPATQV** | RYLPPATQV | 9 | 27.8 | 0.02 | 21702000 | YES | 4012500 | YES | 65516000 | YES | Q91VC3 |
| **RYLQTLTTI** | RYLQTLTTI | 9 | 7.2 | 0.01 | 32013000 | YES | 198650000 | YES | 348490000 | YES | P54116 |
| **RYLSLKEKL** | RYLSLKEKL | 9 | 70.7 | 0.07 | 12822000 | YES | | YES | 1291400 | NO | Q8CC88 |
| **RYMELYTHV** | RYMELYTHV | 9 | 17.5 | 0.01 | | YES | 5275200 | YES | | YES | Q9WTX6 |
| **RYNPGSESI** | RYNPGSESI | 9 | 28 | 0.02 | 544980 | YES | | YES | 7760800 | YES | Q9Z0Y9 |
| **RYQAGGLTV** | RYQAGGLTV | 9 | 30 | 0.02 | 49917000 | YES | | YES | 1224200 | YES | Q8R0W0 |
| **RYQEALSEL** | RYQEALSEL | 9 | 24.8 | 0.015 | 3872500 | YES | 5593300 | YES | 83003000 | YES | Q8R092 |
| **RYQEVIQEL** | RYQEVIQEL | 9 | 65.4 | 0.06 | | YES | 2383900 | YES | 43062000 | YES | Q5DU37 |
| **RYSGMLETV** | RYSGM(+15.99)LETV | 9 | 68.8 | 0.07 | 1932900 | NO | 23121000 | YES | 54579000 | NO | F8VQB6 |
| **RYSPAYAHL** | RYSPAYAHL | 9 | 254.7 | 0.25 | | YES | | YES | 957370 | YES | O09012 |
| **RYTESISMV** | RYTESISM(+15.99)V | 9 | 56.2 | 0.05 | 3012500 | YES | 262240 | YES | 17907000 | YES | O35657 |
| **RYTESISMV** | RYTESISMV | 9 | 56.2 | 0.05 | 834010 | YES | 494120 | YES | | YES | O35657 |
| **RYTNSSTEI** | RYTNSSTEI | 9 | 11.9 | 0.01 | 2896200 | YES | 110790 | YES | 3934200 | YES | P83940 |
| **RYYGAISKL** | RYYGAISKL | 9 | 114.6 | 0.125 | 10038000 | YES | 3848700 | YES | 124160000 | YES | Q6KCD5 |
| **SAVKNLQQL** | SAVKNLQQL | 9 | 14213.6 | 8.5 | 713980 | YES | 101410 | YES | 4478100 | YES | Q8VIJ8 |
| **SAVVDKDFL** | SAVVDKDFL | 9 | 21369.6 | 16 | 1421200 | YES | | YES | 4195300 | YES | P55264 |
| **SFATSGHLI** | SFATSGHLI | 9 | 100.7 | 0.1 | 270590 | NO | | YES | | YES | Q9WV76 |
| **SFEPVKSHL** | SFEPVKSHL | 9 | 98.4 | 0.09 | 7365200 | YES | 2476900 | YES | 25714000 | YES | E9QAT4 |
| **SFGVTLHEL** | SFGVTLHEL | 9 | 2536.7 | 1.5 | 60696000 | YES | 33964000 | YES | 31009000 | YES | P52332 |

| | | | | | | | | | | | |
|---|---|---|---|---|---|---|---|---|---|---|---|
| **SFHPSGDFI** | SFHPSGDFI | 9 | 251.9 | 0.25 | 26202000 | YES | 15136000 | YES | 85486000 | YES | Q99LC2 |
| **SFHPSGNYL** | SFHPSGNYL | 9 | 109.5 | 0.1 | 3608600 | YES | 1555100 | YES | | YES | Q8JZX3 |
| **SFHSSFSEI** | SFHSSFSEI | 9 | 47.7 | 0.04 | 2061300 | YES | 1709700 | YES | 31514000 | YES | Q8K4J0 |
| **SFHSTQTDL** | SFHSTQTDL | 9 | 206.1 | 0.2 | 746340 | YES | 2541800 | YES | 52704000 | YES | Q9EP71 |
| **SFHVSGTWL** | SFHVSGTWL | 9 | 179.5 | 0.175 | 627210 | NO | | YES | 18013000 | YES | Q5SUQ9 |
| **SFIKGKCTV** | SFIKGKC(+119.00)TV | 9 | 146.9 | 0.15 | 226310 | NO | 1089500 | NO | 1765500 | NO | E9Q555 |
| **SFIPAVNDL** | SFIPAVNDL | 9 | 591.6 | 0.5 | 4392900 | YES | 7613000 | YES | 54993000 | YES | Q8BHX1 |
| **SFLEDLTKM** | SFLEDLTKM(+15.99) | 9 | 4337.1 | 2.5 | 1587000 | YES | 2594200 | YES | 2986100 | NO | Q5U430 |
| **SFLESFGRL** | SFLESFGRL | 9 | 1731.4 | 1.1 | | YES | | YES | 2504300 | YES | Q3V2Q8 |
| **SFLETVNQL** | SFLETVNQL | 9 | 754.8 | 0.6 | 5575500 | YES | 73794000 | YES | 338000000 | YES | Q9EP52 |
| **SFLPAPTQL** | SFLPAPTQL | 9 | 104.3 | 0.1 | 2931100 | YES | 8640900 | YES | 36573000 | YES | Q9CSN1 |
| **SFLPSGSEI** | SFLPSGSEI | 9 | 42.5 | 0.04 | | YES | | YES | 5593100 | YES | Q9JKC7 |
| **SFMKGLTEL** | SFM(+15.99)KGLTEL | 9 | 56.8 | 0.05 | 8490500 | YES | 709040 | NO | 43896000 | NO | O35099 |
| **SFNALLREL** | SFNALLREL | 9 | 1504.2 | 1 | | YES | 891090 | YES | 1224500 | YES | Q8C547 |
| **SFNNVKQWL** | SFNNVKQWL | 9 | 907.9 | 0.7 | 20603000 | YES | 2361500 | YES | 2893900 | YES | P62821 |
| **SFNPAISNI** | SFNPAISNI | 9 | 131 | 0.125 | 1008300 | NO | | YES | 9382600 | YES | Q6P3Y5 |
| **SFQHLLQTL** | SFQHLLQTL | 9 | 280.1 | 0.25 | 8777200 | YES | 2518600 | YES | 1325400 | YES | Q00897:P22599:Q00898:Q00896:P07758 |
| **SFQKIFSEL** | SFQKIFSEL | 9 | 210.8 | 0.2 | 2740600 | NO | 3181700 | YES | 10383000 | YES | Q80YR4 |
| **SFSHSFSAL** | SFSHSFSAL | 9 | 803 | 0.6 | 2291800 | NO | 1777500 | YES | 8594000 | YES | Q3UYV9 |
| **SFTDVRTAI** | SFTDVRTAI | 9 | 72.7 | 0.07 | | YES | 2123200 | YES | 2528200 | YES | P11276 |
| **SFTGTKTSL** | SFTGTKTSL | 9 | 108.7 | 0.1 | | YES | 1599900 | YES | 97941000 | YES | A2A4P0 |
| **SFVGTLQYL** | SFVGTLQYL | 9 | 805.5 | 0.6 | 374550 | NO | | YES | | YES | O88351 |
| **SFVGTRSYM** | SFVGTRSYM(+15.99) | 9 | 287.2 | 0.25 | 11224000 | NO | 5854900 | NO | 408150000 | NO | P31938:Q63932 |
| **SFVNTMTSL** | SFVNTM(+15.99)TSL | 9 | 81.4 | 0.08 | 5144900 | YES | 26784000 | YES | 118000000 | YES | P28660 |
| **SFVNTMTSL** | SFVNTMTSL | 9 | 81.4 | 0.08 | 4042500 | YES | 16995000 | YES | | YES | P28660 |
| **SFVSVLHAL** | SFVSVLHAL | 9 | 144.8 | 0.15 | 648520 | YES | 5493100 | YES | 25028000 | YES | Q91V83 |
| **SFYNVKTKL** | SFYNVKTKL | 9 | 129.2 | 0.125 | | YES | 142130 | YES | 666630 | NO | Q8K3K7 |
| **SFYPSLTVV** | SFYPSLTVV | 9 | 128.5 | 0.125 | 3711000 | NO | 4880100 | YES | 40477000 | YES | Q8R316 |
| **SGYDFENRL** | SGYDFENRL | 9 | 24474.6 | 20 | 73167000 | YES | | YES | 84138 | NO | Q8VE09 |

| | | | | | | | | | | | |
|---|---|---|---|---|---|---|---|---|---|---|---|
| **SIAAFIQRL** | SIAAFIQRL | 9 | 13280.4 | 8 | | YES | | YES | 1448500 | YES | Q9Z1R2 |
| **SIINFIERL** | SIINFIERL | 9 | 15477.5 | 9.5 | | YES | 170040 | YES | | YES | Q9QZ09 |
| **SLIGSKTQI** | SLIGSKTQI | 9 | 220.7 | 0.2 | 23251000 | NO | 2297400 | NO | 45963000 | NO | Q9R1P4 |
| **SMSTTRTYI** | SMSTTRTYI | 9 | 78.9 | 0.08 | 513670 | YES | | YES | | YES | Q6ZQ38 |
| **SQPVNPHSL** | SQPVNPHSL | 9 | 16601.8 | 11 | | YES | 62800 | YES | | YES | Q3UHH1 |
| **STLRLLTTI** | STLRLLTTI | 9 | 233.8 | 0.25 | 340530 | NO | 2223900 | YES | 26576000 | YES | Q4VA53 |
| **SYADLCSTI** | SYADLC(+119.00)STI | 9 | 7.5 | 0.01 | 1369500 | NO | 854610 | NO | 5568600 | NO | P51944 |
| **SYADLITRA** | SYADLITRA | 9 | 374.3 | 0.4 | 747590 | NO | | YES | 614350 | YES | Q9WVH4 |
| **SYAEQLSML** | SYAEQLSM(+15.99)L | 9 | 25.5 | 0.015 | 633020 | NO | 5403800 | YES | 26060000 | YES | E9Q7G0 |
| **SYAETPLQL** | SYAETPLQL | 9 | 99.7 | 0.1 | 3555600 | YES | 1620200 | YES | 1022600 | YES | Q7TQI7 |
| **SYAKNGELL** | SYAKNGELL | 9 | 161 | 0.15 | 760870 | NO | | YES | 1161200 | YES | Q9Z2A0 |
| **SYALSRHDV** | SYALSRHDV | 9 | 33.5 | 0.025 | 2664300 | YES | 121580 | NO | 711210 | NO | Q8CD26 |
| **SYAMANTGI** | SYAM(+15.99)ANTGI | 9 | 12.6 | 0.01 | 44918000 | YES | 1493500 | YES | 17884000 | NO | Q8R1S9:Q8CFE6 |
| **SYANVKQWL** | SYANVKQWL | 9 | 26.6 | 0.02 | 17064000 | YES | | YES | 8035600 | YES | Q9D1G1 |
| **SYAQNAKVI** | SYAQNAKVI | 9 | 38.4 | 0.03 | | YES | 168040 | NO | 5587000 | NO | Q8VDP2 |
| **SYASQHSQL** | SYASQHSQL | 9 | 13.2 | 0.01 | 1308200 | YES | | YES | 5336300 | YES | Q68FH0 |
| **SYASQQSKL** | SYASQQSKL | 9 | 36 | 0.025 | 1841000 | YES | | YES | 24726000 | YES | Q78T81 |
| **SYAVSVNHV** | SYAVSVNHV | 9 | 19.8 | 0.01 | 7939600 | YES | 3345100 | YES | 106950000 | YES | Q8K387 |
| **SYDPQKQLI** | SYDPQKQLI | 9 | 200.6 | 0.175 | 5842800 | YES | | YES | 1216600 | NO | Q9D358 |
| **SYDPVKDVL** | SYDPVKDVL | 9 | 204 | 0.2 | 3481600 | YES | 2885500 | YES | 3564200 | YES | Q8CDG3 |
| **SYEAAASAL** | SYEAAASAL | 9 | 10.7 | 0.01 | 589670 | NO | 267120 | YES | 10466000 | YES | Q61033 |
| **SYEEAKNTL** | SYEEAKNTL | 9 | 14.3 | 0.01 | 1542500 | YES | 998700 | YES | 55857000 | YES | Q9WV54 |
| **SYEKQDTLL** | SYEKQDTLL | 9 | 80.1 | 0.08 | 8608200 | YES | | YES | 4113000 | YES | P46978 |
| **SYENMVTEI** | SYENM(+15.99)VTEI | 9 | 9.7 | 0.01 | 18254000 | YES | 29515000 | YES | 205070000 | YES | P54728 |
| **SYENMVTEI** | SYENMVTEI | 9 | 9.7 | 0.01 | 23440000 | YES | 28611000 | YES | | YES | P54728 |
| **SYESTIQSL** | SYESTIQSL | 9 | 25 | 0.015 | 549550 | NO | | YES | 895180 | YES | Q6TXD4 |
| **SYFKDRAHI** | SYFKDRAHI | 9 | 39.8 | 0.03 | 2013200 | NO | 511080 | NO | 11298000 | NO | O88559 |
| **SYFKGASLL** | SYFKGASLL | 9 | 19.8 | 0.01 | 370730 | NO | 2846600 | YES | 19398000 | YES | Q8C129 |
| **SYFKNNAYL** | SYFKNNAYL | 9 | 53.8 | 0.05 | 17984000 | YES | 4467900 | YES | 58046000 | YES | Q922B2 |

| | | | | | | | | | | | |
|---|---|---|---|---|---|---|---|---|---|---|---|
| **SYFPEITHI** | SYFPEITHI | 9 | 21.8 | 0.01 | 669520000 | YES | 1313300000 | YES | 9931300000 | YES | P52332 |
| **SYFPTVNDI** | SYFPTVNDI | 9 | 23.4 | 0.01 | 2598800 | NO | 3872900 | YES | 7854100 | YES | Q8VHI4 |
| **SYGDILHVI** | SYGDILHVI | 9 | 193.3 | 0.175 | 11318000 | YES | 5972600 | YES | 8824000 | YES | P70175 |
| **SYGDLKNAI** | SYGDLKNAI | 9 | 38.5 | 0.03 | 123190000 | NO | 29885000 | NO | 908840000 | YES | O35326 |
| **SYGKVKEVL** | SYGKVKEVL | 9 | 72.1 | 0.07 | 552100 | NO | 872200 | NO | 7024600 | NO | Q9WTK7 |
| **SYGLTPRLL** | SYGLTPRLL | 9 | 1078.6 | 0.8 | | YES | 3584000 | YES | 15339000 | YES | Q8K224 |
| **SYGPGRQSL** | SYGPGRQSL | 9 | 31.3 | 0.02 | | YES | 998210 | YES | 29374000 | YES | Q62137 |
| **SYGQNKTAF** | SYGQNKTAF | 9 | 263.3 | 0.25 | 111490 | NO | 39886 | YES | 1941700 | YES | Q6PFD9 |
| **SYGSVFKAI** | SYGSVFKAI | 9 | 24.4 | 0.015 | 2414600 | YES | 4834300 | YES | 4050300 | YES | Q9JI10 |
| **SYGSVYKAI** | SYGSVYKAI | 9 | 21.1 | 0.01 | 6326100 | NO | 2128300 | YES | 80968000 | YES | Q9JI11 |
| **SYGTAVTHI** | SYGTAVTHI | 9 | 8.1 | 0.01 | 8724400 | YES | 5794600 | YES | 206830000 | YES | P27808 |
| **SYGVTVWEL** | SYGVTVWEL | 9 | 141 | 0.125 | 120120 | NO | | YES | | YES | Q01279:P70424:Q61526 |
| **SYHPALNAI** | SYHPALNAI | 9 | 9.9 | 0.01 | 69395000 | YES | 22983000 | YES | 408990000 | YES | O88738 |
| **SYHPSGLSL** | SYHPSGLSL | 9 | 22.3 | 0.01 | 8303700 | YES | 5898500 | YES | 84080000 | YES | Q8BXQ8 |
| **SYHSQAVHI** | SYHSQAVHI | 9 | 18.7 | 0.01 | 295410 | NO | | YES | 4727400 | YES | Q8BXQ2 |
| **SYHTDINML** | SYHTDINM(+15.99)L | 9 | 40.8 | 0.03 | 1150400 | YES | 3028800 | YES | 18284000 | YES | P13864 |
| **SYHTDINML** | SYHTDINML | 9 | 40.8 | 0.03 | | YES | 4013700 | YES | | YES | P13864 |
| **SYHVIKGNL** | SYHVIKGNL | 9 | 45.9 | 0.04 | 1704200 | YES | 2161800 | YES | 31336000 | YES | B1AZI6 |
| **SYIFDINTI** | SYIFDINTI | 9 | 31.2 | 0.02 | 258120 | NO | | YES | 4468800 | YES | Q9QYY0 |
| **SYIGANVRL** | SYIGANVRL | 9 | 84.7 | 0.08 | | YES | 2739800 | YES | 12039000 | YES | P40336 |
| **SYIGGHEGL** | SYIGGHEGL | 9 | 122.8 | 0.125 | 33439000 | YES | 4707400 | YES | 101260000 | YES | P40201 |
| **SYIGSPRAV** | SYIGSPRAV | 9 | 31.1 | 0.02 | 28283000 | YES | 10988000 | YES | 184600000 | YES | Q8BJS4 |
| **SYIKDLSVV** | SYIKDLSVV | 9 | 53.9 | 0.05 | 6154400 | YES | 11071000 | YES | 13454000 | NO | Q3TP92 |
| **SYIYGAQHL** | SYIYGAQHL | 9 | 19.3 | 0.01 | 365150 | NO | 277330 | YES | | YES | Q9WVG9 |
| **SYKAGIYSV** | SYKAGIYSV | 9 | 113.4 | 0.125 | 6326100 | YES | 2128300 | YES | 86673000 | YES | Q8K2Z8 |
| **SYKDGKMNI** | SYKDGKM(+15.99)NI | 9 | 117 | 0.125 | 3327500 | YES | 399100 | YES | 23918000 | YES | Q8R3N1 |
| **SYKDGKMNI** | SYKDGKMNI | 9 | 117 | 0.125 | 2535600 | YES | 630200 | YES | | YES | Q8R3N1 |
| **SYKENIMRL** | SYKENIM(+15.99)RL | 9 | 1059.8 | 0.8 | 27755000 | YES | 209980 | YES | 470660 | NO | Q63886:Q64435:Q62452:Q6ZQM8 |

| | | | | | | | | | | | |
|---|---|---|---|---|---|---|---|---|---|---|---|
| **SYKNGFLNL** | SYKNGFLNL | 9 | 389.9 | 0.4 | 12597000 | YES | 1785100 | NO | 1430500 | YES | Q02053 |
| **SYKPHASNL** | SYKPHASNL | 9 | 93.8 | 0.09 | 1073100 | YES | 104540 | YES | 5772200 | YES | P70265 |
| **SYKSVQTTL** | SYKSVQTTL | 9 | 10.1 | 0.01 | 257430 | YES | 1438400 | YES | 42127000 | YES | Q8BFZ9 |
| **SYKTIYREL** | SYKTIYREL | 9 | 77.7 | 0.08 | 11691000 | NO | 827590 | NO | 4253800 | YES | Q9WVE8 |
| **SYLDGKGNL** | SYLDGKGNL | 9 | 63.6 | 0.06 | 2596500 | NO | 1757900 | YES | 63171000 | YES | Q8R2K4 |
| **SYLDQGTQI** | SYLDQGTQI | 9 | 16.6 | 0.01 | 5927600 | NO | | YES | | YES | Q9EPL4 |
| **SYLDVKQRL** | SYLDVKQRL | 9 | 41.6 | 0.03 | 82778000 | YES | 6852400 | YES | 45632000 | YES | P61222 |
| **SYLEDKDLV** | SYLEDKDLV | 9 | 473.3 | 0.4 | 1366300 | YES | 14867000 | YES | 83798000 | YES | P52332 |
| **SYLEDKVYL** | SYLEDKVYL | 9 | 158.1 | 0.15 | 2075200 | NO | 1122600 | YES | 1272400 | YES | Q9D1M4 |
| **SYLEMGHDI** | SYLEMGHDI | 9 | 13.3 | 0.01 | | YES | 2346000 | YES | | YES | P08775 |
| **SYLESKGLL** | SYLESKGLL | 9 | 47.1 | 0.04 | 2520200 | YES | 7724800 | YES | 83082000 | YES | Q99ME2 |
| **SYLFSHVPL** | SYLFSHVPL | 9 | 44.8 | 0.04 | 2905200 | YES | 43976000 | YES | 67729000 | YES | Q9D7G0:Q9CS42 |
| **SYLGGNSTI** | SYLGGNSTI | 9 | 8 | 0.01 | 1892900 | NO | 3639200 | NO | 14089000 | NO | P19091 |
| **SYLHSLQEV** | SYLHSLQEV | 9 | 34.1 | 0.025 | 4061600 | YES | 5307000 | YES | 179190000 | YES | Q5SSZ5 |
| **SYLIGRQKI** | SYLIGRQKI | 9 | 37 | 0.025 | 12728000 | YES | 6231400 | YES | 162590000 | YES | Q91ZX7 |
| **SYLKQLPHF** | SYLKQLPHF | 9 | 1488.6 | 1 | 1741600 | NO | 4682800 | YES | 40511000 | YES | Q6P4T2 |
| **SYLKSELGL** | SYLKSELGL | 9 | 121.1 | 0.125 | 488980 | NO | 2031900 | YES | | YES | Q922B9 |
| **SYLLSIHKV** | SYLLSIHKV | 9 | 81.1 | 0.08 | 1989200 | YES | 1019400 | YES | 3523200 | YES | Q8BX17 |
| **SYLNSVFQL** | SYLNSVFQL | 9 | 64.5 | 0.06 | 677420 | NO | 1007100 | YES | 1646500 | YES | B2RVL6 |
| **SYLNSVQQL** | SYLNSVQQL | 9 | 29 | 0.02 | 8710200 | YES | | YES | 41207000 | YES | Q8CIC2 |
| **SYLPEKLQI** | SYLPEKLQI | 9 | 95 | 0.09 | 874490 | NO | | YES | | YES | Q6PD31 |
| **SYLPGVREL** | SYLPGVREL | 9 | 61 | 0.06 | 1610100 | YES | 5412500 | YES | | YES | Q9D5E4 |
| **SYLPPGTSL** | SYLPPGTSL | 9 | 12.1 | 0.01 | 166660000 | YES | 10950000 | NO | 60470000 | YES | Q8VCF0 |
| **SYLTSASSL** | SYLTSASSL | 9 | 5.6 | 0.01 | 1170600 | NO | 2309600 | YES | 23258000 | YES | Q80TP3 |
| **SYLVSKQEL** | SYLVSKQEL | 9 | 18.5 | 0.01 | 2366300 | YES | 1946900 | YES | 102310000 | YES | Q3UM18 |
| **SYMIPTNDL** | SYM(+15.99)IPTNDL | 9 | 99.9 | 0.1 | 6227400 | YES | 2100000 | YES | 11583000 | YES | P00405 |
| **SYMPPSTVL** | SYM(+15.99)PPSTVL | 9 | 12.6 | 0.01 | 13315000 | NO | 4605000 | NO | 6171300 | NO | Q9DB77 |
| **SYMPQQVTV** | SYM(+15.99)PQQVTV | 9 | 37.3 | 0.025 | 4332500 | NO | 2654900 | YES | 19520000 | NO | Q5SSH7 |
| **SYMPTVSHL** | SYM(+15.99)PTVSHL | 9 | 10 | 0.01 | 4860700 | YES | 2574400 | YES | 86717000 | YES | Q9Z2E9 |

| | | | | | | | | | | | |
|---|---|---|---|---|---|---|---|---|---|---|---|
| **SYMIPTNDL** | SYMIPTNDL | 9 | 99.9 | 0.1 | 3345900 | YES | 1513900 | YES | | YES | P00405 |
| **SYMPTVSHL** | SYMPTVSHL | 9 | 10 | 0.01 | 2927900 | YES | 5001200 | YES | | YES | Q9Z2E9 |
| **SYNIAITRA** | SYNIAITRA | 9 | 272.9 | 0.25 | 263550 | YES | | YES | 16355000 | YES | E9Q286 |
| **SYNKAISYL** | SYNKAISYL | 9 | 13.5 | 0.01 | 19324000 | YES | 4738700 | YES | 32776000 | YES | A2ALW5 |
| **SYNKVYKSL** | SYNKVYKSL | 9 | 23.9 | 0.015 | 1474000 | YES | 125770 | YES | 3483100 | YES | Q6WKZ7 |
| **SYNLTVREL** | SYNLTVREL | 9 | 177.5 | 0.175 | 4703300 | YES | 7121400 | YES | 80676000 | YES | Q9ESE1 |
| **SYNPAENAV** | SYNPAENAV | 9 | 26.9 | 0.02 | 8908800 | YES | 564990 | YES | 5024000 | YES | Q8CIE6 |
| **SYNPSGQGL** | SYNPSGQGL | 9 | 69.3 | 0.07 | 17399000 | YES | | YES | 57198000 | YES | Q8CI61 |
| **SYNPSSQAL** | SYNPSSQAL | 9 | 15.1 | 0.01 | 3160700 | NO | 1621100 | YES | 87359000 | YES | Q2NL51 |
| **SYNPVTHQL** | SYNPVTHQL | 9 | 28.5 | 0.02 | 1080500 | YES | 7308000 | YES | 83692000 | YES | Q6NWV3 |
| **SYNTVAQEL** | SYNTVAQEL | 9 | 13.7 | 0.01 | 10353000 | YES | 4228300 | YES | 34171000 | YES | Q9QX47 |
| **SYNWLQETL** | SYNWLQETL | 9 | 83.1 | 0.08 | 2300500 | YES | 1017400 | YES | 1361500 | YES | Q8BM55 |
| **SYQDLASQI** | SYQDLASQI | 9 | 12.5 | 0.01 | 17078000 | YES | | YES | 16264000 | YES | P48410 |
| **SYQDLRSAL** | SYQDLRSAL | 9 | 14.2 | 0.01 | 1082200 | NO | | YES | 7215900 | YES | Q9CQE2 |
| **SYQEGLARL** | SYQEGLARL | 9 | 106 | 0.1 | 4924400 | YES | 3194900 | YES | 29072000 | YES | Q9D2V5 |
| **SYQEMIANL** | SYQEM(+15.99)IANL | 9 | 37 | 0.025 | 2757800 | YES | 8528900 | YES | 97464000 | YES | Q64674 |
| **SYQEMIANL** | SYQEMIANL | 9 | 37 | 0.025 | 3077400 | YES | 7864600 | YES | | YES | Q64674 |
| **SYQESTKQL** | SYQESTKQL | 9 | 29.4 | 0.02 | 1495700 | YES | | YES | 369850 | YES | Q5DU02:Q8CEG8 |
| **SYQGRNEII** | SYQGRNEII | 9 | 110 | 0.1 | 28055000 | YES | 343430 | YES | 1019700 | YES | Q9QYF1 |
| **SYQPIVDYI** | SYQPIVDYI | 9 | 17.3 | 0.01 | 2054800 | NO | 17768000 | YES | 31441000 | YES | Q8C650 |
| **SYQQALLRI** | SYQQALLRI | 9 | 23.7 | 0.015 | | YES | 621850 | YES | 35334000 | YES | Q9ERC3 |
| **SYQSLVSLP** | SYQSLVSLP | 9 | 531.2 | 0.5 | 5502900 | NO | 3752500 | NO | 5114200 | NO | Q9D8Y1 |
| **SYQSQINQI** | SYQSQINQI | 9 | 12.6 | 0.01 | 27692000 | YES | 15538000 | YES | 236990000 | YES | Q99MJ9 |
| **SYSATKETL** | SYSATKETL | 9 | 17.9 | 0.01 | 19873000 | YES | 8202500 | YES | 359450000 | YES | P09405 |
| **SYSDMKRAL** | SYSDM(+15.99)KRAL | 9 | 48.7 | 0.04 | 841230 | YES | 1097200 | YES | 63545000 | YES | Q3TWW8 |
| **SYSDMKRAL** | SYSDMKRAL | 9 | 48.7 | 0.04 | 1068300 | YES | | YES | 201680 | YES | Q3TWW8 |
| **SYSEVKSDL** | SYSEVKSDL | 9 | 40 | 0.03 | 141880 | NO | 1871000 | YES | 76319000 | YES | O35245 |
| **SYSGSIQSL** | SYSGSIQSL | 9 | 62.6 | 0.06 | 36520000 | YES | | YES | 2090400 | YES | B2RQE8 |
| **SYSKGASVI** | SYSKGASVI | 9 | 19.7 | 0.01 | 793220 | NO | | YES | 5405800 | YES | Q11011 |

| | | | | | | | | | | | |
|---|---|---|---|---|---|---|---|---|---|---|---|
| **SYSQGRSFA** | SYSQGRSFA | 9 | 216.4 | 0.2 | 147310 | YES | | YES | 1333000 | YES | A2AN08 |
| **SYSQLITLV** | SYSQLITLV | 9 | 89.4 | 0.09 | 1154800 | NO | | YES | 16956000 | YES | Q6PAR5 |
| **SYSQSKQFL** | SYSQSKQFL | 9 | 37 | 0.025 | 10402000 | YES | 1472600 | YES | 14476000 | YES | Q9CR62 |
| **SYSSIIREV** | SYSSIIREV | 9 | 134.3 | 0.125 | 21671000 | YES | | YES | | YES | Q60991 |
| **SYSSLIRNL** | SYSSLIRNL | 9 | 126.2 | 0.125 | 20130000 | YES | | YES | 6686500 | YES | Q64324 |
| **SYSSSRSDL** | SYSSSRSDL | 9 | 17.7 | 0.01 | | YES | | YES | 57247000 | YES | Q91VM5;Q9WV02 |
| **SYTPSKISV** | SYTPSKISV | 9 | 37.9 | 0.03 | 387610 | YES | | YES | 7485600 | YES | Q8K2H6 |
| **SYTSVLSRL** | SYTSVLSRL | 9 | 16.8 | 0.01 | | YES | 1692600 | YES | 19112000 | YES | Q80TA9 |
| **SYTVGQSEL** | SYTVGQSEL | 9 | 47.4 | 0.04 | 2956700 | NO | 1165300 | YES | 8399000 | YES | Q7TMY8 |
| **SYTYPPSSL** | SYTYPPSSL | 9 | 51.3 | 0.04 | 8541200 | NO | | YES | 13084000 | YES | P59326 |
| **SYVAIINKS** | SYVAIINKS | 9 | 1229.9 | 0.9 | 634300 | YES | 155230 | YES | | YES | P39054 |
| **SYVDIHTGL** | SYVDIHTGL | 9 | 77.3 | 0.07 | 127960000 | YES | 26715000 | YES | 82168000 | YES | Q9DCC4 |
| **SYVGSHREL** | SYVGSHREL | 9 | 53.6 | 0.05 | 9450900 | YES | 2560200 | YES | 54019000 | YES | Q8CHI8 |
| **SYVLTRVGL** | SYVLTRVGL | 9 | 378.7 | 0.4 | 789920 | YES | | YES | 6986100 | YES | Q8R3I3 |
| **SYVPARSLP** | SYVPARSLP | 9 | 371.7 | 0.4 | 3499400 | NO | 6081000 | NO | 105680000 | NO | Q9QZE5 |
| **SYVPVNGRL** | SYVPVNGRL | 9 | 47 | 0.04 | 2495600 | YES | 1191800 | YES | 34170000 | YES | Q9WUP7 |
| **SYVTTSTRT** | SYVTTSTRT | 9 | 38 | 0.03 | 29213 | NO | 8155600 | YES | 33984000 | YES | P20152 |
| **SYWLVRTEL** | SYWLVRTEL | 9 | 15.7 | 0.01 | | YES | 5702200 | YES | 66642000 | YES | P42859 |
| **SYWSVGETI** | SYWSVGETI | 9 | 8.4 | 0.01 | 1459000 | NO | 6599800 | NO | 8664400 | NO | Q78IS1 |
| **SYYADKHEA** | SYYADKHEA | 9 | 352.7 | 0.3 | 265480 | YES | | YES | 326700 | YES | Q8K1N2 |
| **SYYAVAHAV** | SYYAVAHAV | 9 | 11.2 | 0.01 | | YES | 249100 | YES | 6634300 | YES | Q3TKT4 |
| **SYYGPLNLL** | SYYGPLNLL | 9 | 241 | 0.25 | 15221000 | YES | 36613000 | YES | 24608000 | YES | O09005 |
| **SYYTVAHAI** | SYYTVAHAI | 9 | 6.3 | 0.01 | | YES | 7430600 | YES | 154180000 | YES | Q6DIC0 |
| **TDPVTIENK** | TDPVTIENK | 9 | 38755 | 70 | | YES | | YES | | YES | P01872 |
| **TFASTLSHL** | TFASTLSHL | 9 | 144.4 | 0.15 | 4379500 | YES | 5374800 | YES | 30859000 | YES | Q8CFI7 |
| **TFHPTISGL** | TFHPTISGL | 9 | 2069.5 | 1.3 | 1701900 | YES | 1072300 | YES | 2547500 | YES | A2AH22 |
| **TFINLMTHI** | TFINLM(+15.99)THI | 9 | 69.8 | 0.07 | 13991000 | YES | 36427000 | YES | 378730000 | NO | Q8BVE3 |
| **TFINLMTHI** | TFINLMTHI | 9 | 69.8 | 0.07 | 13480000 | YES | 49247000 | YES | 494870 | NO | Q8BVE3 |
| **TFITSKEDL** | TFITSKEDL | 9 | 334.4 | 0.3 | 3384100 | YES | | YES | 14686000 | YES | Q9Z1D1 |

| | | | | | | | | | | | |
|---|---|---|---|---|---|---|---|---|---|---|---|
| **TFLPAKALL** | TFLPAKALL | 9 | 2987.9 | 1.8 | 7521700 | YES | 5081900 | YES | 54932000 | YES | Q78JE5 |
| **TFLPSRGIL** | TFLPSRGIL | 9 | 1191.3 | 0.9 | 9738400 | NO | 22953000 | NO | 28388000 | NO | F8VQB6 |
| **TFLQTATLI** | TFLQTATLI | 9 | 157.3 | 0.15 | 579700 | NO | | YES | 15114000 | YES | Q9JLV2 |
| **TFQEAQSRL** | TFQEAQSRL | 9 | 986.9 | 0.8 | 60157 | NO | 2161800 | YES | 26838000 | YES | P26039 |
| **TFQPVNNNL** | TFQPVNNNL | 9 | 312.6 | 0.3 | 4366000 | NO | 570020 | YES | 3681800 | YES | Q91V09 |
| **TFVPVANEL** | TFVPVANEL | 9 | 601.4 | 0.5 | 2470800 | YES | 31300000 | YES | 8875500 | YES | Q5SYD0 |
| **TGIRNLEWL** | TGIRNLEWL | 9 | 17090.2 | 11 | | YES | | YES | 236640 | NO | Q8BMD6 |
| **TINVGLTSI** | TINVGLTSI | 9 | 1704.8 | 1.1 | | YES | | YES | 7078100 | YES | O35381 |
| **TNQDFIQRL** | TNQDFIQRL | 9 | 22944.4 | 18 | | YES | | YES | | YES | Q80TM9 |
| **TSPVNEKTL** | TSPVNEKTL | 9 | 28217.1 | 28 | | YES | | YES | | YES | P11859 |
| **TWNKLLTTI** | TWNKLLTTI | 9 | 305.3 | 0.3 | 18436000 | YES | 10438000 | YES | 76108000 | YES | Q9D4H8 |
| **TYDEVQTRL** | TYDEVQTRL | 9 | 292.1 | 0.3 | 625510 | YES | 420770 | YES | 1437000 | YES | Q80XL1 |
| **TYDQMYNDL** | TYDQM(+15.99)YNDL | 9 | 235.3 | 0.25 | 1650500 | NO | 222440 | YES | 1268500 | NO | Q9CY97 |
| **TYDYAKTIL** | TYDYAKTIL | 9 | 166.4 | 0.15 | 52832000 | YES | 7122800 | YES | 15985000 | YES | Q91V92 |
| **TYETSLSEI** | TYETSLSEI | 9 | 11.9 | 0.01 | 16610000 | YES | | YES | 12291000 | YES | Q5HZI1 |
| **TYFFGATHV** | TYFFGATHV | 9 | 20.1 | 0.01 | 5533300 | YES | | YES | 7401400 | YES | Q7TT23 |
| **TYFPTWEGL** | TYFPTWEGL | 9 | 476.3 | 0.4 | 2166500 | YES | 1120500 | YES | 3893800 | YES | Q99PV0 |
| **TYFSGMVLI** | TYFSGM(+15.99)VLI | 9 | 108.3 | 0.1 | 1724200 | NO | 6979800 | NO | 4548800 | NO | O35678 |
| **TYGALVTQL** | TYGALVTQL | 9 | 69.3 | 0.07 | 16552000 | YES | 41652000 | YES | 91409000 | YES | O55013 |
| **TYGITVAEL** | TYGITVAEL | 9 | 1878.3 | 1.2 | 7720600 | NO | 5131400 | NO | 11026000 | NO | Q6P4T2 |
| **TYHASGTEL** | TYHASGTEL | 9 | 12.1 | 0.01 | 7531600 | YES | 3843500 | YES | 643700000 | YES | Q3U1N2 |
| **TYHTAASTL** | TYHTAASTL | 9 | 7.3 | 0.01 | 1949100 | YES | 1107600 | YES | 106630000 | YES | Q9R0X0 |
| **TYIESSTKV** | TYIESSTKV | 9 | 20.6 | 0.01 | 8565800 | YES | 1686800 | YES | 63500000 | YES | Q8VBZ3 |
| **TYKALNTFI** | TYKALNTFI | 9 | 21.2 | 0.01 | 1133600 | NO | 9566200 | YES | 62959000 | YES | Q8VBZ3 |
| **TYKDSGVDI** | TYKDSGVDI | 9 | 830.1 | 0.7 | 439810 | NO | | YES | 2080400 | YES | Q64737 |
| **TYKPNPNQI** | TYKPNPNQI | 9 | 153 | 0.15 | 171740 | NO | | YES | 2028500 | YES | Q924K8 |
| **TYKRQVVEL** | TYKRQVVEL | 9 | 1144.4 | 0.8 | 218260 | NO | | YES | 1103000 | YES | Q8BUK6 |
| **TYLAALETL** | TYLAALETL | 9 | 19.2 | 0.01 | 914380 | YES | 346240 | YES | 5872600 | YES | P47738 |
| **TYLDSKSQL** | TYLDSKSQL | 9 | 25.9 | 0.015 | 742000 | NO | 854570 | YES | 31777000 | YES | Q8K389 |

| | | | | | | | | | | | |
|---|---|---|---|---|---|---|---|---|---|---|---|
| **TYLKDLEVI** | TYLKDLEVI | 9 | 212.6 | 0.2 | 13803000 | YES | 16622000 | YES | 29377000 | YES | F8VPU2:Q91VS8 |
| **TYLPAGQSV** | TYLPAGQSV | 9 | 27.7 | 0.02 | 98829000 | YES | 8522300 | YES | 39578000 | YES | P67778 |
| **TYLPGIVGL** | TYLPGIVGL | 9 | 772.9 | 0.6 | 654170 | YES | | YES | 6942500 | YES | Q3TIX9 |
| **TYLPQSYLI** | TYLPQSYLI | 9 | 154.6 | 0.15 | 8195500 | YES | 6918600 | YES | 16747000 | YES | O89051 |
| **TYLVSKESI** | TYLVSKESI | 9 | 14.1 | 0.01 | 1321300 | YES | 1446800 | YES | 130850000 | YES | Q6PNC0 |
| **TYNHLSSWL** | TYNHLSSWL | 9 | 96.9 | 0.09 | 3147800 | NO | | YES | 10432000 | NO | Q91V41 |
| **TYNMVLNLL** | TYNM(+15.99)VLNLL | 9 | 173.9 | 0.175 | 1510000 | NO | 11200000 | YES | 115970000 | NO | Q9CZU3 |
| **TYNMAPSAL** | TYNMAPSAL | 9 | 32.5 | 0.02 | | YES | 2286900 | YES | | YES | Q61985 |
| **TYNNILTVL** | TYNNILTVL | 9 | 55.3 | 0.05 | 9060700 | YES | | YES | 45529000 | YES | Q9EQH3 |
| **TYNPVPGVM** | TYNPVPGVM(+15.99) | 9 | 446.6 | 0.4 | 9602400 | NO | 362890 | NO | 1386400 | NO | Q99L13 |
| **TYQAMVHEL** | TYQAMVHEL | 9 | 17.8 | 0.01 | | YES | | YES | | YES | P97390 |
| **TYQDIQNTI** | TYQDIQNTI | 9 | 27.9 | 0.02 | 34845000 | YES | 26484000 | YES | 154470000 | YES | P08775 |
| **TYQENLTDL** | TYQENLTDL | 9 | 72.6 | 0.07 | 530820 | NO | 1345200 | YES | 12291000 | YES | Q9EPW0 |
| **TYQLGFHSI** | TYQLGFHSI | 9 | 27.4 | 0.02 | 4337400 | YES | 3779400 | YES | 5434700 | YES | Q60692 |
| **TYQNTAQTV** | TYQNTAQTV | 9 | 16 | 0.01 | 2229500 | YES | | YES | 6260600 | YES | Q3UA37 |
| **TYQQVQQTL** | TYQQVQQTL | 9 | 20.1 | 0.01 | 20283000 | YES | 10132000 | YES | 150790000 | YES | Q91V81 |
| **TYRELFNSI** | TYRELFNSI | 9 | 104.5 | 0.1 | 697880 | NO | 17704000 | YES | 44182000 | YES | P59764 |
| **TYRNLINKL** | TYRNLINKL | 9 | 929.4 | 0.7 | 15999000 | NO | 7207300 | YES | 48475000 | NO | Q9JI13 |
| **TYSPPLNKL** | TYSPPLNKL | 9 | 880 | 0.7 | 136470000 | YES | 88076000 | YES | 1049300000 | YES | P02340 |
| **TYSPSRVLI** | TYSPSRVLI | 9 | 222.7 | 0.2 | 34866000 | YES | 6540700 | YES | 126060000 | YES | Q91XU0 |
| **TYSSVYDSI** | TYSSVYDSI | 9 | 35.3 | 0.025 | 805950 | NO | 3097400 | NO | 4782200 | YES | Q8R1B4 |
| **TYTSARTLL** | TYTSARTLL | 9 | 22.1 | 0.01 | | YES | 8778800 | YES | 287270000 | YES | Q61881 |
| **TYTSLKTKL** | TYTSLKTKL | 9 | 28.6 | 0.02 | 6072300 | YES | | YES | 3707300 | YES | Q8K1N1 |
| **TYVHSSATI** | TYVHSSATI | 9 | 24.3 | 0.015 | 3433700 | YES | 690280 | YES | 19164000 | YES | Q8BGQ7 |
| **VAYWRQAGL** | VAYWRQAGL | 9 | 38465.9 | 70 | | YES | | YES | | YES | P56382 |
| **VFIGNLNTL** | VFIGNLNTL | 9 | 1690.7 | 1.1 | 7802800 | YES | 3611900 | YES | 12274000 | YES | Q9Z204 |
| **VFIPAGTHV** | VFIPAGTHV | 9 | 160.2 | 0.15 | 1471800 | YES | | YES | 223380 | NO | Q8VCA8 |
| **VFVDSLTKV** | VFVDSLTKV | 9 | 3673.7 | 2.5 | 205500 | NO | 29854000 | YES | 15831000 | YES | Q8BTM8 |
| **VGFDYKERL** | VGFDYKERL | 9 | 31254.2 | 35 | | YES | 1270700 | YES | 1202400 | NO | Q60598 |

| | | | | | | | | | | | |
|---|---|---|---|---|---|---|---|---|---|---|---|
| **VGVNNPVFL** | VGVNNPVFL | 9 | 28002.7 | 27 | 344840 | YES | | YES | 781790 | YES | Q9DBN5 |
| **VQVLVPLPQ** | VQVLVPLPQ | 9 | 26981.6 | 25 | 163590 | YES | | YES | | YES | Q62419 |
| **VSFELFADK** | VSFELFADK | 9 | 39546.7 | 75 | 4173300 | NO | | YES | 1182200 | YES | P17742 |
| **VYAGTPTKV** | VYAGTPTKV | 9 | 126.5 | 0.125 | 219230 | NO | | YES | | YES | P97440 |
| **VYAHAGTTL** | VYAHAGTTL | 9 | 16.2 | 0.01 | 872270 | YES | 348800 | YES | 9539100 | YES | Q99JN2 |
| **VYDLLKTNL** | VYDLLKTNL | 9 | 306.3 | 0.3 | 8745000 | YES | 5480700 | YES | 10263000 | YES | Q3TXS7 |
| **VYESLISHI** | VYESLISHI | 9 | 26.3 | 0.015 | 12880000 | YES | 21960000 | YES | 111880000 | YES | Q61037 |
| **VYFPALTSL** | VYFPALTSL | 9 | 16 | 0.01 | 620160 | YES | | YES | 5965600 | YES | Q8VCL5 |
| **VYGAMHVEI** | VYGAM(+15.99)HVEI | 9 | 98.3 | 0.09 | 18325000 | YES | 1942700 | YES | 43710000 | NO | Q5SW19 |
| **VYGSLASVL** | VYGSLASVL | 9 | 114.9 | 0.125 | 29445000 | NO | 15442000 | NO | 14181000 | NO | Q8JZK9 |
| **VYHNLKNVI** | VYHNLKNVI | 9 | 50.1 | 0.04 | 9648100 | YES | | YES | 11670000 | YES | P17182 |
| **VYIITKPEL** | VYIITKPEL | 9 | 434.4 | 0.4 | 2124200 | YES | 1007400 | YES | 1147100 | YES | Q9WUE4 |
| **VYIPAHGRL** | VYIPAHGRL | 9 | 102 | 0.1 | 5848500 | YES | 866430 | YES | 2709200 | YES | Q920I9 |
| **VYIPSKTDL** | VYIPSKTDL | 9 | 18.6 | 0.01 | 13760000 | YES | 4171000 | YES | | YES | Q9CRB2 |
| **VYISNGQVL** | VYISNGQVL | 9 | 137.1 | 0.125 | | YES | 2127300 | YES | 2188600 | YES | Q9JLJ1 |
| **VYKASLNLI** | VYKASLNLI | 9 | 108.7 | 0.1 | 1160400 | NO | 6667900 | NO | 28504000 | NO | P52293 |
| **VYKELKNLI** | VYKELKNLI | 9 | 302.4 | 0.3 | 172960 | NO | 692830 | NO | 689010 | NO | P09055 |
| **VYKGQITAI** | VYKGQITAI | 9 | 88.6 | 0.09 | 8653200 | YES | | YES | 33398000 | YES | Q8K442 |
| **VYLENKEQV** | VYLENKEQV | 9 | 286.6 | 0.25 | 440560 | NO | | YES | 6974300 | YES | Q9CY66 |
| **VYLPNIQSL** | VYLPNIQSL | 9 | 148.7 | 0.15 | 444610 | NO | 1096700 | YES | 14938000 | YES | Q62240 |
| **VYLTPKTSV** | VYLTPKTSV | 9 | 13.4 | 0.01 | 397660 | NO | 987700 | NO | 8266600 | NO | Q8BFX3 |
| **VYNASNNEL** | VYNASNNEL | 9 | 41.8 | 0.04 | 10668000 | YES | 3778400 | YES | 208710000 | YES | P62242 |
| **VYNPMPFEL** | VYNPMPFEL | 9 | 186.4 | 0.175 | 452940 | NO | 795920 | YES | | YES | Q3U0M1 |
| **VYNVTQHAV** | VYNVTQHAV | 9 | 190.8 | 0.175 | 1527200 | YES | | YES | 6468000 | YES | O09167 |
| **VYQETRERL** | VYQETRERL | 9 | 97.2 | 0.09 | 3398500 | NO | 1080700 | NO | 28503000 | NO | Q9CWK3 |
| **VYQQTASLL** | VYQQTASLL | 9 | 44.6 | 0.04 | 2092500 | NO | 7440600 | YES | 21786000 | YES | Q3U269 |
| **VYSNTIQSI** | VYSNTIQSI | 9 | 104.4 | 0.1 | 4107300 | NO | 28545000 | YES | 145670000 | YES | P08752:Q9DC51 |
| **VYSNTIQSL** | VYSNTIQSL | 9 | 253.3 | 0.25 | 4107300 | NO | 28545000 | YES | 145670000 | YES | |
| **VYSRTFTWL** | VYSRTFTWL | 9 | 135.6 | 0.125 | 864320 | NO | 58806000 | YES | 15045000 | YES | Q9WTI7 |

| | | | | | | | | | | | |
|---|---|---|---|---|---|---|---|---|---|---|---|
| **VYTPVINGI** | VYTPVINGI | 9 | 164 | 0.15 | 2417700 | NO | 911270 | NO | 1844300 | YES | Q8BYR8 |
| **VYTTSYQQI** | VYTTSYQQI | 9 | 26.9 | 0.02 | 193740 | NO | | YES | 7334000 | YES | P63139 |
| **VYTTTRSSL** | VYTTTRSSL | 9 | 14.3 | 0.01 | 1186400 | YES | 1945100 | YES | 158170000 | YES | Q8BXA5 |
| **VYTTTVHWL** | VYTTTVHWL | 9 | 151 | 0.15 | 2207500 | NO | | YES | 6367000 | YES | Q99PV0 |
| **VYVAGGQHL** | VYVAGGQHL | 9 | 65.3 | 0.06 | 79450 | NO | 58705 | NO | | YES | Q8BGY4 |
| **VYVDGKEEI** | VYVDGKEEI | 9 | 144.9 | 0.15 | 31211000 | YES | 5814400 | YES | 187030000 | YES | Q9WUD8 |
| **VYWKIYNSI** | VYWKIYNSI | 9 | 33.7 | 0.025 | 18832000 | YES | 8523700 | YES | 59729000 | YES | Q99NB9 |
| **VYWPTPSAL** | VYWPTPSAL | 9 | 42.2 | 0.04 | 3229700 | NO | | YES | 14604000 | YES | Q8BYH8 |
| **VYYFSKGAL** | VYYFSKGAL | 9 | 71.8 | 0.07 | 559000 | NO | 1156800 | YES | 6842400 | YES | Q8VEE4 |
| **VYYPVRHHL** | VYYPVRHHL | 9 | 25.2 | 0.015 | 2854400 | YES | 3167300 | YES | 75638000 | YES | Q80YV3 |
| **WFTDSNNAI** | WFTDSNNAI | 9 | 502.9 | 0.5 | 1578500 | NO | 1147700 | YES | 1396900 | YES | A2AH22 |
| **WYDPNASLL** | WYDPNASLL | 9 | 502.8 | 0.5 | 1421000 | NO | | YES | 4378100 | YES | Q9QXL8 |
| **WYIGDQNPM** | WYIGDQNPM | 9 | 274.2 | 0.25 | 1341200000 | YES | 23636000 | YES | | YES | P70274 |
| **WYIGDQNPM** | WYIGDQNPM(+15.99) | 9 | 274.2 | 0.25 | 3269900000 | YES | 69189000 | YES | 70015000 | YES | P70274 |
| **WYKSNMNGV** | WYKSNM(+15.99)NGV | 9 | 450.9 | 0.4 | 8245800 | YES | 614630 | YES | 10953000 | NO | Q9EQQ9 |
| **WYNPILNRV** | WYNPILNRV | 9 | 84.2 | 0.08 | 4565100 | YES | 22636000 | YES | 54569000 | YES | Q9Z222 |
| **WYQPSFHGV** | WYQPSFHGV | 9 | 58.4 | 0.05 | 5365700 | YES | 5918200 | YES | 1278900 | YES | Q9WVG6 |
| **YFISSTTRI** | YFISSTTRI | 9 | 29.6 | 0.02 | 358740 | NO | | YES | 14212000 | YES | Q3UMC0 |
| **YFKSSLTTI** | YFKSSLTTI | 9 | 30.5 | 0.02 | 6094400 | NO | 1553300 | YES | 54607000 | YES | Q9EPT5 |
| **YFNWIKTQL** | YFNWIKTQL | 9 | 499.3 | 0.5 | 637810 | NO | 655180 | YES | 323670 | NO | Q8BVE3 |
| **YFQPAISRL** | YFQPAISRL | 9 | 217.9 | 0.2 | 7952000 | YES | 697070 | YES | 1136700 | YES | Q9WU79 |
| **YFRQSLSYL** | YFRQSLSYL | 9 | 504.5 | 0.5 | 13220000 | YES | 28638000 | YES | 72366000 | YES | Q9Z2X8 |
| **YFVPAFSGL** | YFVPAFSGL | 9 | 1215.1 | 0.9 | 695670 | YES | | YES | 249100 | NO | Q64516 |
| **YVHVNRDTL** | YVHVNRDTL | 9 | 6981.2 | 4 | | YES | | YES | 1114100 | YES | Q9JLV6 |
| **YYDKAFDRI** | YYDKAFDRI | 9 | 231.5 | 0.25 | 661030 | NO | 166900 | YES | 610480 | NO | O70194 |
| **YYDPMISKL** | YYDPMISKL | 9 | 88.6 | 0.09 | 10750000 | YES | 505480 | YES | | YES | Q91ZA3 |
| **YYFEVVQKL** | YYFEVVQKL | 9 | 111.9 | 0.125 | 859130 | NO | 2442100 | YES | 4273900 | YES | Q8BT60 |
| **YYFPVKNVI** | YYFPVKNVI | 9 | 12.2 | 0.01 | 8743000 | YES | 82641000 | YES | 1275700000 | YES | Q921M3 |
| **YYGILQEKI** | YYGILQEKI | 9 | 800.1 | 0.6 | 28189000 | YES | 22661000 | YES | 46171000 | YES | P97858 |

| | | | | | | | | | | | |
|---|---|---|---|---|---|---|---|---|---|---|---|
| **YYHLLAEKI** | YYHLLAEKI | 9 | 72.4 | 0.07 | 469160 | NO | 2145500 | YES | 19897000 | YES | P45481:B2RWS6 |
| **YYINGKTGL** | YYINGKTGL | 9 | 18.7 | 0.01 | 22579000 | NO | 7268400 | NO | 273280000 | YES | Q6ZQ93 |
| **YYKASVTRL** | YYKASVTRL | 9 | 24.6 | 0.015 | 7597600 | YES | | YES | 27561000 | YES | Q9QVP9 |
| **YYKQGIGHL** | YYKQGIGHL | 9 | 137.7 | 0.125 | 585950 | NO | | YES | 725700 | NO | Q8R1X6 |
| **YYLNDLDRI** | YYLNDLDRI | 9 | 170.1 | 0.15 | 42360000 | YES | 54185000 | YES | 23757000 | YES | Q9DC51 |
| **YYLNDLDRL** | YYLNDLDRL | 9 | 439.8 | 0.4 | 42360000 | YES | 54185000 | YES | 23757000 | YES | |
| **YYLNDLERI** | YYLNDLERI | 9 | 145.1 | 0.15 | 19988000 | YES | 303410000 | YES | 412770000 | YES | P08752 |
| **YYLPLGKTL** | YYLPLGKTL | 9 | 20.9 | 0.01 | | YES | 4972700 | YES | 24733000 | YES | Q8BL99 |
| **YYLTDVDRI** | YYLTDVDRI | 9 | 76.4 | 0.07 | 1221400 | NO | 3952800 | YES | 359890 | YES | P21278 |
| **YYNELETRV** | YYNELETRV | 9 | 55.9 | 0.05 | 3376500 | YES | 3580500 | YES | 44220000 | YES | Q8K2T8 |
| **YYNMLLKKL** | YYNM(+15.99)LLKKL | 9 | 298 | 0.3 | 5689000 | YES | 207870 | NO | 5311200 | YES | Q9DBG3 |
| **YYQDTPKQI** | YYQDTPKQI | 9 | 49.4 | 0.04 | | YES | | YES | 250170 | NO | Q9CPT5 |
| **YYQGLYETL** | YYQGLYETL | 9 | 30.2 | 0.02 | 41368000 | YES | 37748000 | YES | 33764000 | YES | Q6NZJ6 |
| **YYQGNTSRL** | YYQGNTSRL | 9 | 62.5 | 0.06 | 42020 | NO | | YES | 641000 | YES | Q68FH4 |
| **YYQGVIQQI** | YYQGVIQQI | 9 | 21.8 | 0.01 | 2140800 | NO | 4002900 | YES | 6643700 | YES | Q8K0T4 |
| **YYQSGRMLL** | YYQSGRM(+15.99)LL | 9 | 43.3 | 0.04 | 354030000 | YES | 7617000 | YES | 34241000 | NO | P55096 |
| **YYRNQQQGL** | YYRNQQQGL | 9 | 2058.6 | 1.3 | 126800 | NO | 209130 | YES | 1160400 | YES | P23949 |
| **YYSGLKHFI** | YYSGLKHFI | 9 | 67.4 | 0.06 | 144430 | NO | 270490 | NO | 6305200 | YES | Q8C092 |
| **YYSPTKNEI** | YYSPTKNEI | 9 | 21.2 | 0.01 | 72119000 | YES | 28448000 | NO | 41759000 | YES | Q4PZA2 |
| **YYTNSLEKL** | YYTNSLEKL | 9 | 136.8 | 0.125 | 673160 | YES | | YES | 646470 | YES | Q61329 |
| **YYTPQRVDV** | YYTPQRVDV | 9 | 348.8 | 0.3 | 79815 | NO | | YES | 482530 | YES | Q8K1R7 |
| **YYVGAAHGL** | YYVGAAHGL | 9 | 103.3 | 0.1 | 11363000 | YES | 1856400 | YES | 13291000 | YES | O89112 |
| **YYVRILSTI** | YYVRILSTI | 9 | 8.5 | 0.01 | 5697200 | YES | 2384600 | YES | 7239900 | YES | P54775 |

### Supplementary Table 3

| AA SEQUENCE | Peptide | Length | H-2K$^b$ IC50 (nM) | H-2K$^b$ Rank | Spectral intensity value DDA (hep) | Present in DIA (hep) | Spectral intensity value DDA (skin) | Present in DIA (skin) | Spectral intensity value DDA (spleen) | Present in DIA (spleen) | Accession |
|---|---|---|---|---|---|---|---|---|---|---|---|
| **AAFVFRKL** | AAFVFR KL | 8 | 4.9 | 0.01 | 294570 | YES | 6990200 | YES | 6187400 | YES | Q8K0S9 |
| **AAILFSERL** | AAILFSERL | 9 | 39.3 | 0.125 | | YES | | YES | | YES | Q3KQJ0 |
| **AAIRFKDL** | AAIRFKDL | 8 | 60.2 | 0.175 | 275657.526 | NO | | YES | | YES | Q7TQE6 |
| **AALDFKNV** | AALDFKNV | 8 | 146.1 | 0.4 | 116260 | NO | 6603200 | YES | | YES | Q8CG47 |
| **AALEFLNRF** | AALEFLNRF | 9 | 123.9 | 0.4 | 232820 | NO | 2852800 | YES | 231350 | NO | Q60864 |
| **AALIYGKL** | AALIYGKL | 8 | 28.4 | 0.09 | 53770.4931 | NO | 6327900 | YES | 3991800 | YES | Q9DC23 |
| **AALRFLSQL** | AALRFLSQL | 9 | 11.4 | 0.04 | 48180.7613 | NO | 2522300 | YES | | YES | Q9D2N9 |
| **AAPVLVRL** | AAPVLVRL | 8 | 399.5 | 0.9 | 94546.3206 | NO | | YES | | YES | Q8C0Q3 |
| **AAVKFHNL** | AAVKFHNL | 8 | 15.8 | 0.05 | 182840 | YES | 4538200 | YES | 6557300 | YES | Q64521 |
| **AAVVYHKL** | AAVVYHKL | 8 | 94.7 | 0.3 | 348390 | YES | 583970 | YES | | YES | Q9WTN3 |
| **AAYAYSAL** | AAYAYSAL | 8 | 2.7 | 0.01 | | YES | | YES | | YES | Q9CQ22 |
| **AAYEFTTL** | AAYEFTTL | 8 | 3.3 | 0.01 | 211837.201 | YES | 19091000 | YES | | YES | P32233 |
| **AAYGFRNI** | AAYGFRNI | 8 | 9.6 | 0.03 | 10593612.9 | YES | 18562000 | YES | | YES | Q9CYQ7 |
| **AAYSFYNV** | AAYSFYNV | 8 | 3.3 | 0.01 | 438680 | YES | 9605800 | YES | 3362200 | YES | O70310 |
| **AFYKISTL** | AFYKISTL | 8 | 187.2 | 0.5 | | YES | 866640 | NO | 325800 | NO | Q91YW3 |
| **AFYQFVNNL** | AFYQFVNNL | 9 | 10.8 | 0.03 | 224450 | NO | 11748000 | YES | 510730 | YES | Q9WTV7 |
| **AFYYIHNL** | AFYYIHNL | 8 | 87.6 | 0.25 | 1032800 | YES | 68357000 | YES | 13818000 | YES | Q9QYC0 |
| **AFYYPSRL** | AFYYPSRL | 8 | 89 | 0.25 | 155445.182 | YES | 245850 | YES | | YES | Q6P5C7 |
| **AGFDFKQL** | AGFDFKQL | 8 | 27.2 | 0.08 | 750932.749 | NO | 501750 | YES | | YES | Q0VEE6 |
| **AGIGFYQHL** | AGIGFYQHL | 9 | 7.8 | 0.02 | 58087 | NO | | YES | 2049700 | YES | Q6ZPY2 |
| **AGLSYSKI** | AGLSYSKI | 8 | 316 | 0.8 | 64494.2085 | YES | | YES | | YES | F8VPZ5 |

| | | | | | | | | | | | |
|---|---|---|---|---|---|---|---|---|---|---|---|
| **AGPEYKGL** | AGPEYKGL | 8 | 249.8 | 0.6 | 145500 | YES | | YES | 431740 | YES | Q07113 |
| **AGPWYRNL** | AGPWYRNL | 8 | 16.7 | 0.05 | 112300 | YES | 6784700 | YES | 2133200 | YES | P98195 |
| **AGYMYTQL** | AGYM(+15.99)YTQL | 8 | 3.1 | 0.01 | 956440 | NO | 11525000 | YES | 26600000 | NO | A2AN08 |
| **AGYSFEKL** | AGYSFEKL | 8 | 10.7 | 0.03 | | YES | | YES | 1421300 | YES | Q91Y86 |
| **AIFNFQSL** | AIFNFQSL | 8 | 4 | 0.01 | | YES | 44311000 | YES | | YES | Q9CR64 |
| **AIHEFQETL** | AIHEFQETL | 9 | 296 | 0.7 | 110937.673 | NO | 2561500 | YES | | YES | Q64282 |
| **AILERFPTI** | AILERFPTI | 9 | 147.7 | 0.4 | 50915.2503 | NO | 3915500 | YES | | YES | P50652 |
| **AIRVFANI** | AIRVFANI | 8 | 11.5 | 0.04 | 454774.346 | NO | | YES | 512020 | NO | Q8QZY1 |
| **AIVNFVSKV** | AIVNFVSKV | 9 | 183.2 | 0.5 | 172350.713 | NO | 4391300 | YES | 7011900 | YES | Q5SSZ5 |
| **AIVSFAHV** | AIVSFAHV | 8 | 10.5 | 0.03 | 792690.188 | YES | | YES | | YES | P43247 |
| **AIVTFITKV** | AIVTFITKV | 9 | 441.4 | 1 | | YES | 16513000 | YES | 8339500 | YES | Q8CGB6 |
| **AIYAFSHL** | AIYAFSHL | 8 | 2.2 | 0.01 | 400084.566 | YES | 1908100 | YES | | YES | Q9JJ59 |
| **AIYEFIHNF** | AIYEFIHNF | 9 | 46.3 | 0.15 | 1540000 | NO | 4882700 | YES | | YES | Q9WVL1 |
| **ALVRFVNL** | ALVRFVNL | 8 | 39.5 | 0.125 | 1723468.17 | NO | 6226700 | YES | | YES | A2BE28 |
| **AMYIFLHTV** | AM(+15.99)YIFLHTV | 9 | 31.7 | 0.09 | 5706700 | YES | 25153000 | YES | 15170000 | YES | Q9CQZ0 |
| **AMYIFLHTV** | AMYIFLHTV | 9 | 31.7 | 0.09 | 3655400 | YES | 59652000 | YES | 1244300 | YES | Q9CQZ0 |
| **ANFSFAPVTKL** | ANFSFAPVTKL | 11 | 315.4 | 0.8 | 50350.4342 | NO | | YES | 1055800 | YES | Q8BJ34 |
| **ANIDFYAQV** | ANIDFYAQV | 9 | 7.5 | 0.02 | 172820 | YES | 20108000 | YES | 867440 | YES | P16882 |
| **ANILFTREL** | ANILFTREL | 9 | 55.1 | 0.175 | 208191.216 | NO | 507260 | YES | 111340 | YES | Q9ERI6 |
| **ANIQFRTI** | ANIQFRTI | 8 | 128.1 | 0.4 | 190432.619 | YES | 490680 | YES | 2203400 | YES | F8VPQ2 |
| **ANLIYYSL** | ANLIYYSL | 8 | 9.2 | 0.025 | 1624200 | YES | 7724900 | YES | 5338000 | YES | Q91VR2 |
| **ANLKYLSL** | ANLKYLSL | 8 | 25.3 | 0.08 | 1446318.76 | NO | 2597200 | YES | 1559700 | YES | Q8R1T4 |
| **ANLLFTREL** | ANLLFTREL | 9 | 56 | 0.175 | 208191.216 | NO | 507260 | YES | 111340 | YES | |
| **ANRYFTTV** | ANRYFTTV | 8 | 130.1 | 0.4 | | YES | 1687100 | NO | | YES | Q6PAR5 |
| **ANVVFTQL** | ANVVFTQL | 8 | 13.4 | 0.04 | | YES | 9351100 | YES | 1901700 | YES | Q8VHH5 |
| **ANYQRDGPM** | ANYQRDGPM | 9 | 214.7 | 0.6 | 139990 | YES | 375930 | YES | 27928 | YES | P24270 |
| **ANYQRDGPM** | ANYQRDGPM(+15.99) | 9 | 214.7 | 0.6 | 17986000 | YES | 143040 | YES | 265360 | YES | P24270 |
| **AQFEHTILL** | AQFEHTILL | 9 | 206.7 | 0.5 | 286751.293 | NO | 2350100 | YES | | YES | O08663 |
| **AQFEHTLL** | AQFEHTLL | 8 | 132.4 | 0.4 | | YES | 4416500 | YES | | YES | Q8BP48 |

| | | | | | | | | | | | |
|---|---|---|---|---|---|---|---|---|---|---|---|
| **AQFKFTVL** | AQFKFTVL | 8 | 25.6 | 0.08 | 3074500 | NO | 25223000 | NO | 431260 | YES | P50580 |
| **AQFNFQNV** | AQFNFQNV | 8 | 13.1 | 0.04 | 1019863.12 | NO | | YES | | YES | Q91YI6 |
| **AQQLFQKL** | AQQLFQKL | 8 | 435.6 | 1 | 605015.43 | NO | | YES | | YES | Q8R420 |
| **AQYHFPKL** | AQYHFPKL | 8 | 14.5 | 0.05 | 597970 | YES | 6229000 | NO | 143760 | NO | P39061 |
| **AQYKFIYV** | AQYKFIYV | 8 | 18.9 | 0.06 | 2079700 | NO | 28620000 | NO | 1191600000 | YES | P29351 |
| **AQYNFILV** | AQYNFILV | 8 | 25.3 | 0.08 | | YES | | YES | | YES | Q9D0R2 |
| **AQYRFIYM** | AQYRFIYM | 8 | 8.7 | 0.025 | 3406503.31 | YES | 32786000 | YES | 4627700 | YES | P35235 |
| **AQYRFIYM** | AQYRFIYM(+15.99) | 8 | 8.7 | 0.025 | 2208900 | YES | 15790000 | YES | 44105000 | YES | P35235 |
| **AQYSFDKL** | AQYSFDKL | 8 | 36.3 | 0.125 | 125810 | YES | 16345000 | YES | 5552400 | YES | Q8C547 |
| **ASITFEHM** | ASITFEHM(+15.99) | 8 | 22 | 0.07 | 847980 | NO | 9001500 | YES | 1327000 | NO | Q91WK2 |
| **ASLRYLGL** | ASLRYLGL | 8 | 5.5 | 0.015 | 1118998.08 | YES | 6380400 | YES | 6640200 | YES | Q8VC16 |
| **ASPEFTKL** | ASPEFTKL | 8 | 20.9 | 0.06 | 17233000 | YES | 84586000 | YES | 70477000 | YES | Q3TJZ6 |
| **ASPIFTHV** | ASPIFTHV | 8 | 17.1 | 0.05 | 1096800 | YES | 35247000 | YES | 120870000 | YES | Q8C5N3 |
| **ASVKFHVL** | ASVKFHVL | 8 | 64 | 0.2 | 1058738.05 | YES | 4419800 | YES | | YES | Q6PDQ2 |
| **ASVRLAALL** | ASVRLAALL | 9 | 122.9 | 0.4 | 651850 | YES | 4436600 | YES | | YES | Q91ZU9 |
| **ASYEFTTL** | ASYEFTTL | 8 | 2.6 | 0.01 | 8478500 | YES | 16574000 | YES | 10134000 | YES | Q9QXB9 |
| **ASYEFVQRL** | ASYEFVQRL | 9 | 4.2 | 0.01 | 21688000 | YES | 865700000 | YES | 154000000 | YES | Q9JHU4 |
| **ASYEVKEL** | ASYEVKEL | 8 | 215.3 | 0.6 | 287588.779 | NO | 2098400 | YES | 380350 | NO | Q60967 |
| **ASYLFRGL** | ASYLFRGL | 8 | 2.1 | 0.01 | 1195890.98 | YES | 15684000 | YES | 21413000 | YES | Q7TN60 |
| **ASYLLAAL** | ASYLLAAL | 8 | 3.4 | 0.01 | 749490 | NO | 3658600 | YES | | YES | P99027 |
| **ASYNHPVL** | ASYNHPVL | 8 | 28.9 | 0.09 | 26479.358 | NO | 1582600 | YES | 1912800 | YES | Q6NZM9 |
| **ATIFFTRL** | ATIFFTRL | 8 | 9.5 | 0.03 | 182270 | YES | 9519200 | YES | | YES | Q61194 |
| **ATIRVTNL** | ATIRVTNL | 8 | 84 | 0.25 | 1400600 | YES | 13367000 | YES | 6405500 | YES | Q9Z1D1 |
| **ATLAYTKL** | ATLAYTKL | 8 | 22.2 | 0.07 | | YES | 6899700 | YES | | YES | Q8BUR4 |
| **ATLEFEERL** | ATLEFEERL | 9 | 107.1 | 0.3 | 61720.7667 | NO | 665310 | YES | 377830 | YES | Q5SW75 |
| **ATLVFHNL** | ATLVFHNL | 8 | 6.9 | 0.02 | 37932000 | YES | 1110600000 | YES | 1629400000 | YES | P42227 |
| **ATPIFSKM** | ATPIFSKM(+15.99) | 8 | 103.3 | 0.3 | 2370100 | YES | 2628300 | NO | 4524800 | NO | P10649 |
| **ATQVYPKL** | ATQVYPKL | 8 | 118.9 | 0.4 | 14029000 | YES | 67874000 | YES | 122660000 | YES | Q8K3W0 |
| **ATRSFPQL** | ATRSFPQL | 8 | 37.4 | 0.125 | 513200 | YES | 245240000 | YES | 159290000 | YES | Q07076 |

| | | | | | | | | | | | |
|---|---|---|---|---|---|---|---|---|---|---|---|
| **ATYIFLQTF** | ATYIFLQTF | 9 | 39 | 0.125 | | YES | 8473900 | YES | 718510 | YES | Q8K1A5 |
| **ATYIFNGL** | ATYIFNGL | 8 | 3.1 | 0.01 | 468950 | NO | 7482700 | YES | 988130 | YES | Q9QXK3 |
| **ATYSYKEAL** | ATYSYKEAL | 9 | 27.7 | 0.08 | 601050 | YES | 19085000 | YES | 2846800 | YES | Q8R349 |
| **ATYTFIQQL** | ATYTFIQQL | 9 | 7.6 | 0.02 | 9016900 | YES | 57050000 | YES | 17251000 | YES | Q9R0N0 |
| **AVDRFQTL** | AVDRFQTL | 8 | 481.1 | 1 | 352490 | YES | 5790900 | YES | 1288200 | YES | Q9WVJ2 |
| **AVFTWTNL** | AVFTWTNL | 8 | 12 | 0.04 | 76152 | NO | 5669100 | YES | | YES | Q3UH60:Q8BWT5 |
| **AVIDFSEAHL** | AVIDFSEAHL | 10 | 199.7 | 0.5 | 206991.469 | NO | 2303100 | YES | 2062800 | NO | P0CB42 |
| **AVIHFAGL** | AVIHFAGL | 8 | 5.9 | 0.015 | | YES | 6541500 | YES | 4090200 | YES | Q8R059 |
| **AVIKFLEL** | AVIKFLEL | 8 | 70.5 | 0.2 | | YES | 50551000 | YES | | YES | P43247 |
| **AVIQFLERI** | AVIQFLERI | 9 | 236.9 | 0.6 | | YES | 8257900 | YES | | YES | Q571H0 |
| **AVLKFAAA** | AVLKFAAA | 8 | 212.1 | 0.6 | 42594.9248 | NO | 727220 | YES | 271940 | YES | P14206 |
| **AVLKYYKV** | AVLKYYKV | 8 | 177 | 0.5 | 600613.759 | YES | 2701700 | YES | 2032500 | YES | P62983 |
| **AVLRYTKL** | AVLRYTKL | 8 | 14 | 0.04 | 1539416.03 | YES | 13493000 | YES | 22335000 | YES | Q9WUK4 |
| **AVLSFSTRL** | AVLSFSTRL | 9 | 20.1 | 0.06 | 2384400 | YES | 90025000 | YES | 151070000 | YES | P46978 |
| **AVPEFQGL** | AVPEFQGL | 8 | 40 | 0.125 | 3529477.41 | YES | 13494000 | YES | | YES | Q9QZE5 |
| **AVPVFKTL** | AVPVFKTL | 8 | 76.1 | 0.25 | 58943 | NO | 1792700 | YES | 620180 | YES | Q8K2G4 |
| **AVVAFVMKM** | AVVAFVM(+15.99)KM | 9 | 339.2 | 0.8 | 372850 | YES | 421170000 | YES | 40534000 | YES | P01902:P01901:P04223 |
| **AVVAFVMKM** | AVVAFVM(+15.99)KM(+15.99) | 9 | 339.2 | 0.8 | 13066000 | YES | 568050000 | YES | 633310000 | YES | P01902:P01901:P04223 |
| **AVVAFVMKM** | AVVAFVMKM | 9 | 339.2 | 0.8 | 13818000 | YES | 1161900000 | YES | | YES | P01902:P01901:P04223 |
| **AVVAFVMKM** | AVVAFVMKM(+15.99) | 9 | 339.2 | 0.8 | 7300000 | YES | 645460000 | YES | 74529000 | YES | P01902:P01901:P04223 |
| **AVVRFINRF** | AVVRFINRF | 9 | 216.3 | 0.6 | 1122815.46 | NO | 2212800 | NO | 759340 | NO | Q6ZPE2 |
| **AVVSFKEL** | AVVSFKEL | 8 | 178.9 | 0.5 | | YES | | YES | | YES | Q8K394 |
| **AVYQFGSAL** | AVYQFGSAL | 9 | 10.4 | 0.03 | | YES | | YES | | YES | Q80ZE4 |
| **AVYSFEAL** | AVYSFEAL | 8 | 4.4 | 0.01 | 2395400 | NO | | YES | | YES | Q99KC8 |
| **AVYTYLRL** | AVYTYLRL | 8 | 3.7 | 0.01 | 167070 | YES | 2234200 | YES | 3955300 | YES | Q7TPD0 |
| **AWYQKQELL** | AWYQKQELL | 9 | 259.4 | 0.7 | 203715.535 | NO | 369610 | YES | 40471 | NO | Q6NXY1 |
| **AYFTHSNL** | AYFTHSNL | 8 | 44.4 | 0.15 | 4911562.94 | YES | 14987000 | YES | 5398100 | YES | Q8CIE6 |
| **EALSFVSL** | EALSFVSL | 8 | 474.8 | 1 | 405490 | NO | | YES | | YES | P52332:Q62120 |

| | | | | | | | | | | | |
|---|---|---|---|---|---|---|---|---|---|---|---|
| **EIISFQHL** | EIISFQHL | 8 | 146.9 | 0.4 | 1037200 | YES | 38617000 | YES | | YES | Q8BGR2:Q5DU41:Q80WG5 |
| **EQYKFYSV** | EQYKFYSV | 8 | 269.9 | 0.7 | 12983758.9 | NO | 6982900 | NO | 4638800 | NO | Q62425 |
| **ESFKFVRL** | ESFKFVRL | 8 | 13.9 | 0.04 | 727911.623 | NO | 1760800 | NO | 616150 | NO | Q99LC8 |
| **ESFQFYDRL** | ESFQFYDRL | 9 | 13.2 | 0.04 | 204175.179 | NO | 899810 | YES | 144360 | NO | Q8K284 |
| **ETPVYANL** | ETPVYANL | 8 | 42.8 | 0.125 | 2748700 | YES | 12346000 | NO | 33253000 | YES | P15066 |
| **ETYKYFSL** | ETYKYFSL | 8 | 23.6 | 0.07 | 984532.899 | NO | 7780800 | NO | 5289000 | NO | Q9ET30 |
| **EVYLFERI** | EVYLFERI | 8 | 288.1 | 0.7 | 847925.307 | YES | 7195400 | YES | | YES | P97390 |
| **FAPIYADL** | FAPIYADL | 8 | 34.3 | 0.1 | 670080 | YES | 5140300 | YES | | YES | Q9D710 |
| **FAYRFSNL** | FAYRFSNL | 8 | 2 | 0.01 | 1584800 | NO | 29455000 | NO | 67624000 | NO | Q8BU03 |
| **FQFTFKHL** | FQFTFKHL | 8 | 15.8 | 0.05 | 964790 | NO | 8642600 | YES | 1155300 | NO | Q9QYJ0 |
| **FSPSFINHI** | FSPSFINHI | 9 | 161.1 | 0.5 | 258200 | NO | 4297100 | YES | 663560 | NO | Q9JIX9 |
| **FSQEYINL** | FSQEYINL | 8 | 15.7 | 0.05 | 493376.605 | YES | 7226600 | YES | 1486900 | YES | Q9D8V0 |
| **FSVRPFAL** | FSVRPFAL | 8 | 147.3 | 0.4 | | YES | | YES | | YES | O08852 |
| **FTFEYRYL** | FTFEYRYL | 8 | 5.8 | 0.015 | 142540 | NO | 12504000 | YES | | YES | Q8K0V4 |
| **FTFQFNNL** | FTFQFNNL | 8 | 3.6 | 0.01 | 197767.437 | NO | 4755500 | NO | | YES | Q8R5H1 |
| **FTYDYHTL** | FTYDYHTL | 8 | 12.9 | 0.04 | 7739600 | YES | 8368700 | YES | 1324000 | YES | Q9Z239 |
| **FVYIFQEV** | FVYIFQEV | 8 | 19.9 | 0.06 | | YES | 110510000 | YES | 1307900 | YES | Q99K51 |
| **GAVDFSHL** | GAVDFSHL | 8 | 328.3 | 0.8 | | YES | | YES | | YES | O88487 |
| **GLYLHALL** | GLYLHALL | 8 | 167.4 | 0.5 | 65471.9247 | NO | | YES | | YES | Q5IXF8 |
| **GMYIFLHTV** | GMYIFLHTV | 9 | 172.6 | 0.5 | 124271.229 | YES | 5013700 | YES | | YES | Q9CPZ6 |
| **GNYLFHYI** | GNYLFHYI | 8 | 69.4 | 0.2 | 38380 | NO | 1249800 | YES | | YES | P70280 |
| **GQLEFRALL** | GQLEFRALL | 9 | 352.8 | 0.8 | 142123.313 | NO | 2037100 | YES | | YES | P11499:P07901 |
| **GQYEFHSL** | GQYEFHSL | 8 | 91.9 | 0.25 | 1430700 | YES | 16506000 | YES | 561140 | YES | Q61805 |
| **GTYDYTQL** | GTYDYTQL | 8 | 31.7 | 0.09 | 765630 | NO | | YES | 996050 | YES | Q8K2A8 |
| **GTYEFLYTV** | GTYEFLYTV | 9 | 170.8 | 0.5 | 401690 | NO | 2134200 | YES | | YES | Q9ESN6 |
| **GVLKFARL** | GVLKFARL | 8 | 15.5 | 0.05 | | YES | | YES | | YES | O35127 |
| **GVLRFVNL** | GVLRFVNL | 8 | 30.8 | 0.09 | 2488200 | YES | 34288000 | YES | | YES | Q3UGP8 |
| **HAVVFAQL** | HAVVFAQL | 8 | 20.2 | 0.06 | 325969.007 | NO | 743660 | YES | 1101400 | YES | Q9EPL9 |
| **HAYIISSL** | HAYIISSL | 8 | 303.5 | 0.8 | | YES | 2198200 | YES | 1986100 | YES | Q9QXY6 |

| | | | | | | | | | | | |
|---|---|---|---|---|---|---|---|---|---|---|---|
| **HAYIISYL** | HAYIISYL | 8 | 160.8 | 0.4 | | YES | 8362800 | YES | | YES | Q8BH64 |
| **HGVSYVSL** | HGVSYVSL | 8 | 451.6 | 1 | | YES | | YES | 515100 | YES | Q99L04 |
| **HGYIFSSL** | HGYIFSSL | 8 | 6.4 | 0.015 | 1757707.72 | YES | 26834000 | YES | 19580000 | YES | Q9DAA6 |
| **HGYTFANL** | HGYTFANL | 8 | 3 | 0.01 | 1415800 | YES | 72188000 | YES | 125310000 | YES | Q9R1T2 |
| **HIYEFPQL** | HIYEFPQL | 8 | 7.9 | 0.02 | 40560000 | YES | 276940000 | YES | 343060000 | YES | Q8C4B4 |
| **HIYQFEYM** | HIYQFEYM | 8 | 33.8 | 0.1 | 487533.68 | NO | 3111100 | YES | | YES | Q8C079 |
| **HIYQFEYM** | HIYQFEYM(+15.99) | 8 | 33.8 | 0.1 | 368489.922 | NO | 3152900 | YES | 1809900 | YES | Q8C079 |
| **HSALIYSNL** | HSALIYSNL | 9 | 107.8 | 0.3 | 405320 | YES | 28194000 | YES | 3452300 | YES | O55013 |
| **HSIRFVTL** | HSIRFVTL | 8 | 20.3 | 0.06 | 1343483.27 | NO | 1184500 | NO | 361920 | NO | Q91XU0 |
| **HSPAFVQL** | HSPAFVQL | 8 | 52.2 | 0.175 | 802973.736 | NO | | YES | | YES | Q61191 |
| **HSYLYGLL** | HSYLYGLL | 8 | 4.2 | 0.01 | 740220.719 | YES | | YES | | YES | Q9QYC7 |
| **HTFTYTGL** | HTFTYTGL | 8 | 10.7 | 0.03 | 429377.098 | NO | 6812100 | YES | | YES | A2AGH6 |
| **HTYDFEKL** | HTYDFEKL | 8 | 90.4 | 0.25 | 2210534.42 | YES | 39527000 | YES | 3549200 | YES | P07742 |
| **HTYVHATL** | HTYVHATL | 8 | 15 | 0.05 | | YES | 773030 | YES | 691870 | YES | Q9EQJ0 |
| **HVYYFAHL** | HVYYFAHL | 8 | 2.7 | 0.01 | | YES | 31254000 | YES | 1038200 | YES | Q80SU7 |
| **IAAVFHTL** | IAAVFHTL | 8 | 32.3 | 0.1 | 163547.372 | NO | 3174600 | YES | 875050 | YES | Q9JHZ2 |
| **IAFGFHQL** | IAFGFHQL | 8 | 5.4 | 0.015 | 2926136.95 | YES | 4662900 | YES | 1004200 | YES | Q9R0A0 |
| **IAFSYELSKL** | IAFSYELSKL | 10 | 54 | 0.175 | 138855.17 | NO | | YES | | YES | Q9D1P2 |
| **IAGPYNRL** | IAGPYNRL | 8 | 95.3 | 0.3 | 755540 | YES | 9045100 | YES | 2444000 | YES | Q8VDJ3 |
| **IALRYVAL** | IALRYVAL | 8 | 4.2 | 0.01 | 3105600 | YES | 28335000 | YES | 69440000 | YES | Q9JIF7 |
| **IAPEYFEKL** | IAPEYFEKL | 9 | 17.2 | 0.05 | 1792648.41 | YES | 280820 | NO | 451760 | NO | Q9D9V7 |
| **IAPSFVKGF** | IAPSFVKGF | 9 | 193.5 | 0.5 | 602930 | YES | 7042200 | YES | 1899200 | YES | O88967 |
| **IAVIFKQL** | IAVIFKQL | 8 | 13.3 | 0.04 | 189010 | NO | | YES | | YES | Q6PGC1 |
| **IAVSFREL** | IAVSFREL | 8 | 10.8 | 0.03 | 4228600 | YES | 50840000 | YES | 67920000 | YES | Q8BJW5 |
| **IAYAFFHL** | IAYAFFHL | 8 | 2 | 0.01 | | YES | 12616000 | YES | 1095200 | YES | Q8K4P7 |
| **IAYKFGKTV** | IAYKFGKTV | 9 | 30.1 | 0.09 | 4634997.81 | YES | 4537100 | YES | 1444400 | YES | Q8R3Q0 |
| **IAYKFGKTVV** | IAYKFGKTVV | 10 | 276 | 0.7 | 272650.367 | NO | 266690 | NO | 116460 | NO | Q8R3Q0 |
| **IAYLYDRL** | IAYLYDRL | 8 | 2.7 | 0.01 | 98394 | NO | 2716300 | YES | 1267700 | YES | Q9CZ15 |
| **IDFDFTHL** | IDFDFTHL | 8 | 48.1 | 0.15 | 422209.776 | NO | | YES | | YES | O70566 |

| | | | | | | | | | | | |
|---|---|---|---|---|---|---|---|---|---|---|---|
| **IDYQYQLL** | IDYQYQLL | 8 | 26.2 | 0.08 | 126990 | YES | 10378000 | YES | 3196500 | YES | Q9D8X5 |
| **IDYSFPSL** | IDYSFPSL | 8 | 21.7 | 0.07 | 307740 | YES | | YES | | YES | Q91VE6 |
| **IFYFVNKL** | IFYFVNKL | 8 | 79.5 | 0.25 | 67271.5457 | NO | 3859900 | YES | | YES | Q8BH24 |
| **IFYYVQKL** | IFYYVQKL | 8 | 52.7 | 0.175 | 3267207.97 | YES | 31918000 | YES | 21433000 | YES | Q69ZR2 |
| **IGFDEYMNL** | IGFDEYMNL | 9 | 10.7 | 0.03 | 39394 | NO | 3296000 | YES | | YES | P62305 |
| **IGIAYNRL** | IGIAYNRL | 8 | 6.1 | 0.015 | 1069100 | YES | 10650000 | YES | | YES | Q8CIB5 |
| **IGIVKQAGL** | IGIVKQAGL | 9 | 76.6 | 0.25 | 439738.552 | YES | 6112800 | YES | 5997900 | YES | Q9Z2X1 |
| **IGPEYKSM** | IGPEYKSM | 8 | 212.5 | 0.6 | 418236.587 | YES | 1554500 | NO | 84240 | NO | Q99K01 |
| **IGPEYKSM** | IGPEYKSM(+15.99) | 8 | 212.5 | 0.6 | 5935100 | YES | 688020 | NO | 1546300 | NO | Q99K01 |
| **IGPRFKLL** | IGPRFKLL | 8 | 14 | 0.04 | 1431900 | YES | | YES | 1862300 | YES | E9Q3L2 |
| **IGPRFSNL** | IGPRFSNL | 8 | 4.2 | 0.01 | 2796501.91 | NO | 7480700 | NO | 2699400 | NO | P46460 |
| **IGPTYYQRL** | IGPTYYQRL | 9 | 3.9 | 0.01 | 467700 | YES | 41366000 | YES | 208580 | YES | Q8CFI7 |
| **IGYEHEVL** | IGYEHEVL | 8 | 64 | 0.2 | 322550 | YES | | YES | 1912900 | YES | Q3U4G0 |
| **IHYDRITSL** | IHYDRITSL | 9 | 80.6 | 0.25 | 364668.649 | NO | 772650 | YES | | YES | Q8CFQ3 |
| **IIFETPLRV** | IIFETPLRV | 9 | 417.4 | 0.9 | 352207.637 | NO | 2833700 | YES | | YES | Q6NZJ6 |
| **IIHKYPSL** | IIHKYPSL | 8 | 10.6 | 0.03 | 863000.054 | NO | 9896600 | YES | 1128000 | YES | Q3TEA8 |
| **IILKNFEKL** | IILKNFEKL | 9 | 115.1 | 0.3 | 758645.41 | NO | 1786900 | NO | 2466900 | YES | P97371 |
| **IILKYIGM** | IILKYIGM | 8 | 15.3 | 0.05 | 3091608.59 | NO | | YES | | YES | Q9JKW0 |
| **IILKYIGM** | IILKYIGM(+15.99) | 8 | 15.3 | 0.05 | 1668700 | NO | 12168000 | YES | 68646000 | YES | Q9JKW0 |
| **IITRFYQL** | IITRFYQL | 8 | 9.1 | 0.025 | 105440 | NO | 476280 | NO | 1018700 | NO | Q69ZT1 |
| **IIVQFRYI** | IIVQFRYI | 8 | 23 | 0.07 | 48989 | NO | 10144000 | YES | | YES | Q8BHK1 |
| **IIWVKTNQL** | IIWVKTNQL | 9 | 130.8 | 0.4 | 1036184.36 | NO | 2659500 | YES | 1389900 | YES | Q8C3P7 |
| **IIYDRKFLM** | IIYDRKFLM(+15.99) | 9 | 84.1 | 0.25 | 183534.072 | NO | 6929900 | YES | 9705500 | NO | Q60876 |
| **IIYEFESSTQM** | IIYEFESSTQM(+15.99) | 11 | 101.2 | 0.3 | 125410 | NO | 8198900 | YES | 45969000 | YES | Q9DB00 |
| **IIYNPKNL** | IIYNPKNL | 8 | 25.5 | 0.08 | 1628800 | YES | 21285000 | YES | 4971000 | YES | Q9D554 |
| **ILSSFESRL** | ILSSFESRL | 9 | 244.7 | 0.6 | 304490.414 | NO | | YES | | YES | O35250 |
| **INAEFVTQL** | INAEFVTQL | 9 | 40.7 | 0.125 | 331270.488 | NO | 8932700 | YES | 4395100 | YES | Q8CFQ3 |
| **INAIFERL** | INAIFERL | 8 | 25.9 | 0.08 | 41399.0728 | NO | 527920 | NO | 472550 | YES | Q8C5N3 |
| **INFDFNTI** | INFDFNTI | 8 | 19.8 | 0.06 | 1897500 | YES | | YES | | YES | Q8CDD8 |

| | | | | | | | | | | | |
|---|---|---|---|---|---|---|---|---|---|---|---|
| **INFDFPKL** | INFDFPKL | 8 | 6.2 | 0.015 | 286070000 | YES | 1772900000 | YES | 277580000 | YES | P54823 |
| **INFSHDSSFL** | INFSHDSSFL | 10 | 187.8 | 0.5 | 366748.73 | NO | 404440 | NO | 66207 | NO | Q91VM3 |
| **ININSLRL** | ININSLRL | 8 | 159.7 | 0.4 | 88960.4841 | YES | 12526000 | YES | 2967400 | YES | P16460 |
| **INLEFVKV** | INLEFVKV | 8 | 37 | 0.125 | 235270 | NO | | YES | | YES | A2AAE1 |
| **INLEHSVPM** | INLEHSVPM | 9 | 84.3 | 0.25 | | YES | | YES | | YES | Q80TY5 |
| **INLNYKDL** | INLNYKDL | 8 | 38.1 | 0.125 | 778010 | YES | 15517000 | YES | | YES | Q9CY50 |
| **INQRFEEL** | INQRFEEL | 8 | 40.2 | 0.125 | 520293.014 | YES | 3531700 | NO | 1393000 | NO | P42230 |
| **INYDYVHEL** | INYDYVHEL | 9 | 21.2 | 0.06 | 162160 | NO | 14779000 | YES | | YES | E9Q784 |
| **INYQPPTV** | INYQPPTV | 8 | 237 | 0.6 | 46852 | NO | 511000 | YES | 338810 | YES | P68373:P05213:P68368:Q9JJZ2 |
| **INYSFPAKGKL** | INYSFPAKGKL | 11 | 35.6 | 0.125 | 343077.092 | NO | 50256 | NO | | YES | Q8K4L0 |
| **INYVIKQL** | INYVIKQL | 8 | 33.2 | 0.1 | 409783.822 | YES | | YES | | YES | P98192 |
| **INYVVPRV** | INYVVPRV | 8 | 24.6 | 0.07 | 44916.5138 | NO | | YES | 289900 | YES | Q80U59 |
| **IPPEYRHL** | IPPEYRHL | 8 | 132.1 | 0.4 | 1278899.47 | YES | 9571700 | YES | 6949400 | YES | P51125 |
| **IQLEFREL** | IQLEFREL | 8 | 25.1 | 0.08 | 3984095.94 | NO | | YES | 1736900 | YES | A2AJ15 |
| **IQQQYAQV** | IQQQYAQV | 8 | 53.6 | 0.175 | | YES | | YES | 3182500 | YES | Q6Y7W8 |
| **IRYFPTQAL** | IRYFPTQAL | 9 | 437.1 | 1 | 366530 | YES | 29716000 | YES | 10250000 | YES | P51881:P48962 |
| **ISARFVQL** | ISARFVQL | 8 | 6.4 | 0.015 | 1912800 | NO | 19232000 | NO | 14575000 | NO | Q7TN98:Q812E0:Q7TN99 |
| **ISATFKML** | ISATFKM(+15.99)L | 8 | 16.8 | 0.05 | 297973.605 | NO | 3271200 | YES | 4595400 | NO | Q9CZX9 |
| **ISATFKML** | ISATFKML | 8 | 16.8 | 0.05 | 631737.074 | NO | 6350500 | YES | 207500 | NO | Q9CZX9 |
| **ISFEFRSL** | ISFEFRSL | 8 | 2.2 | 0.01 | 18539212.2 | YES | 38297000 | YES | | YES | Q3THF9 |
| **ISFEFRSLL** | ISFEFRSLL | 9 | 4.3 | 0.01 | | YES | 9432300 | YES | | YES | Q3THF9 |
| **ISFKFDHL** | ISFKFDHL | 8 | 2.7 | 0.01 | 4034500 | YES | 120960000 | YES | 473140000 | YES | P47753 |
| **ISILYHQL** | ISILYHQL | 8 | 3.7 | 0.01 | 341950 | NO | 75767000 | NO | 4447000 | NO | Q9D2V5 |
| **ISLDYHQL** | ISLDYHQL | 8 | 7.9 | 0.02 | | YES | | YES | | YES | Q7TPV2 |
| **ISLDYQHL** | ISLDYQHL | 8 | 4.4 | 0.01 | 247110 | YES | 11531000 | YES | | YES | Q5SYL3 |
| **ISLEFRNL** | ISLEFRNL | 8 | 2.5 | 0.01 | 16607000 | YES | 12365000 | YES | 6056600 | YES | P53798 |
| **ISLRFTHL** | ISLRFTHL | 8 | 2.1 | 0.01 | 307341.762 | NO | | YES | 1943200 | NO | Q6P549 |
| **ISPEWKQQL** | ISPEWKQQL | 9 | 83.6 | 0.25 | 45855.2771 | NO | 1387300 | YES | 59327 | NO | Q9WTI7 |

| | | | | | | | | | | | |
|---|---|---|---|---|---|---|---|---|---|---|---|
| **ISPFFPNGV** | ISPFFPNGV | 9 | 53.6 | 0.175 | 120660 | YES | 5108300 | YES | 399880 | YES | Q640L3 |
| **ISPNFNFM** | ISPNFNFM(+15.99) | 8 | 16.2 | 0.05 | 302106.501 | NO | 7797500 | YES | 4367300 | YES | Q9DBB1:Q91Z46 |
| **ISPPIPHL** | ISPPIPHL | 8 | 270.5 | 0.7 | 50042 | NO | 4546500 | YES | 3900200 | YES | Q5SVR0 |
| **ISPRFDVQL** | ISPRFDVQL | 9 | 17.4 | 0.05 | 32432130.2 | YES | 329740000 | YES | 62385000 | YES | P62245 |
| **ISSRFQNL** | ISSRFQNL | 8 | 5.1 | 0.015 | 1166987.98 | NO | 8631600 | NO | 4834000 | NO | Q9QXZ0 |
| **ISTIFKSL** | ISTIFKSL | 8 | 17 | 0.05 | 445520 | YES | 3015400 | YES | | YES | Q9ES00 |
| **ISVRFHNL** | ISVRFHNL | 8 | 2.9 | 0.01 | | YES | 3825300 | YES | | YES | Q8CFL8 |
| **ISVSFYHV** | ISVSFYHV | 8 | 6 | 0.015 | 660140 | YES | 44932000 | YES | 37833000 | YES | Q8JZQ9 |
| **ISYAWKEL** | ISYAWKEL | 8 | 9.4 | 0.03 | 1266707.23 | YES | | YES | 494690 | YES | Q9JIK5 |
| **ISYLYNKL** | ISYLYNKL | 8 | 2.1 | 0.01 | 1060400 | YES | | YES | 3008400 | YES | Q61471:Q9JM55 |
| **ITALHIKL** | ITALHIKL | 8 | 422.1 | 0.9 | 173032.387 | NO | 28797000 | YES | 1863900 | NO | P62264 |
| **ITFIFKSL** | ITFIFKSL | 8 | 3.7 | 0.01 | | YES | 44803000 | YES | 681380 | YES | Q9JLN9 |
| **ITFSYVNNM** | ITFSYVNNM(+15.99) | 9 | 9.4 | 0.03 | 343879.521 | NO | 2516100 | YES | 6404800 | YES | Q8C172 |
| **ITPPGYSHV** | ITPPGYSHV | 9 | 197.3 | 0.5 | 765000 | NO | 20186000 | NO | 14664000 | NO | Q8BLB7 |
| **ITYVHNEL** | ITYVHNEL | 8 | 8.1 | 0.025 | | YES | 5421600 | YES | | YES | Q80YA7 |
| **ITYYFDNV** | ITYYFDNV | 8 | 3.4 | 0.01 | 865921.517 | NO | | YES | | YES | Q6ZQ58 |
| **IVELFRNL** | IVELFRNL | 8 | 17.8 | 0.05 | 738273.077 | YES | 9835200 | YES | | YES | P42227 |
| **IVLTFRQL** | IVLTFRQL | 8 | 4.6 | 0.01 | 431180 | YES | | YES | | YES | Q8BGF9 |
| **IVWEFEQL** | IVWEFEQL | 8 | 9.6 | 0.03 | | YES | 13599000 | YES | | YES | Q62158 |
| **IWIRVASL** | IWIRVASL | 8 | 44.9 | 0.15 | 4903100 | YES | 37280000 | YES | 2278600 | YES | Q8BRG6 |
| **IYFKVTHV** | IYFKVTHV | 8 | 308.1 | 0.8 | 116624.787 | NO | | YES | | YES | Q80SY5 |
| **KAFASLRM** | KAFASLRM | 8 | 497.1 | 1.1 | 295784.456 | YES | 1938000 | YES | 601580 | NO | P47963 |
| **KAFASLRM** | KAFASLRM(+15.99) | 8 | 497.1 | 1.1 | 978884.738 | YES | 738210 | YES | 3830700 | NO | P47963 |
| **KAFDYPSRL** | KAFDYPSRL | 9 | 32.4 | 0.1 | 3783177.22 | YES | 5843200 | YES | 5456200 | YES | Q6P5C7 |
| **KAFEHLQQL** | KAFEHLQQL | 9 | 61.4 | 0.175 | 385364.29 | NO | 854210 | YES | 145090 | NO | O88708 |
| **KAFGFNVL** | KAFGFNVL | 8 | 59.5 | 0.175 | 171750.839 | NO | | YES | | YES | O88712 |
| **KAFHFPSL** | KAFHFPSL | 8 | 9.8 | 0.03 | 417496.483 | YES | 1614500 | YES | 678590 | NO | Q6P5C7 |
| **KAFTYINL** | KAFTYINL | 8 | 4.2 | 0.01 | 305580 | YES | 5796100 | YES | 20891000 | YES | Q62351 |
| **KALEYLKL** | KALEYLKL | 8 | 88.9 | 0.25 | 160064.989 | NO | 2212400 | YES | 1107600 | YES | A2AGT5 |

| | | | | | | | | | | | |
|---|---|---|---|---|---|---|---|---|---|---|---|
| **KALLFVNTL** | KALLFVNTL | 9 | 64 | 0.2 | 106072.464 | NO | 639890 | YES | | YES | Q9D0R4 |
| **KALQFKQV** | KALQFKQV | 8 | 236 | 0.6 | 222557.021 | YES | 94649 | NO | | YES | Q8R164 |
| **KALQFLEQV** | KALQFLEQV | 9 | 353 | 0.8 | 285570 | YES | 18801000 | YES | 5818300 | YES | P80317 |
| **KALTLSNL** | KALTLSNL | 8 | 93.2 | 0.25 | 195001.786 | YES | | YES | | YES | Q8VI47 |
| **KALTYEKL** | KALTYEKL | 8 | 119.7 | 0.4 | 267629.347 | NO | 2129400 | YES | 3160500 | YES | O08800 |
| **KAPGFAHL** | KAPGFAHL | 8 | 13 | 0.04 | 235956.797 | YES | | YES | 40184 | NO | Q91WD5 |
| **KAPVFMEKL** | KAPVFM(+15.99)EKL | 9 | 135.3 | 0.4 | 1804412.17 | YES | 2654900 | NO | 1457500 | NO | Q9ET54 |
| **KAPVFMEKL** | KAPVFMEKL | 9 | 135.3 | 0.4 | 3241187.48 | YES | 5452100 | NO | 447130 | NO | Q9ET54 |
| **KAVTFIDL** | KAVTFIDL | 8 | 205.6 | 0.5 | 964280 | YES | | YES | | YES | Q91YP3 |
| **KAYSFKEQI** | KAYSFKEQI | 9 | 64.5 | 0.2 | 5602975.75 | YES | 2653200 | YES | 430200 | YES | E9PVA8 |
| **KEFIFPNM** | KEFIFPNM | 8 | 277.2 | 0.7 | 2920021.35 | YES | 1033800 | NO | | YES | Q61471 |
| **KGFEFTLM** | KGFEFTLM | 8 | 20.2 | 0.06 | 1129086.87 | YES | 115880000 | YES | | YES | P42208 |
| **KGFGFIKL** | KGFGFIKL | 8 | 52.9 | 0.175 | 206610 | YES | 1292000 | YES | | YES | Q8VIJ6 |
| **KGFTFSAL** | KGFTFSAL | 8 | 4.2 | 0.01 | | YES | | YES | | YES | Q9CYV5 |
| **KGFYFAKL** | KGFYFAKL | 8 | 3.8 | 0.01 | 1273700 | YES | 6887100 | YES | 22488000 | NO | Q9ERU9 |
| **KGIIYRDL** | KGIIYRDL | 8 | 152.1 | 0.4 | | YES | 3719800 | YES | | YES | P28867 |
| **KGLDFALL** | KGLDFALL | 8 | 15.2 | 0.05 | | YES | 24567000 | YES | 6028200 | YES | Q9Z1M8 |
| **KGPQYGTL** | KGPQYGTL | 8 | 148.3 | 0.4 | 1976100 | YES | | YES | | YES | Q99JB8 |
| **KGVAYVYL** | KGVAYVYL | 8 | 65 | 0.2 | 1614049.66 | NO | | YES | | YES | Q8K1H1 |
| **KGYAYTFI** | KGYAYTFI | 8 | 27.4 | 0.08 | 196380 | YES | 5100200 | YES | 18883000 | YES | Q569Z5 |
| **KGYGFAEYM** | KGYGFAEYM | 9 | 106.5 | 0.3 | 309507.539 | NO | | YES | | YES | Q9CW46 |
| **KGYGYQAL** | KGYGYQAL | 8 | 9.7 | 0.03 | | YES | | YES | | YES | Q99LI9 |
| **KGYIFLTL** | KGYIFLTL | 8 | 7.6 | 0.02 | | YES | 23538000 | YES | 10935000 | YES | Q80VI1 |
| **KIFEFKETL** | KIFEFKETL | 9 | 63.4 | 0.2 | 895720.436 | NO | 23426000 | YES | 212030000 | YES | Q8BW96 |
| **KIFTASNV** | KIFTASNV | 8 | 171.6 | 0.5 | 230164.51 | NO | | YES | 956870 | YES | Q80X50 |
| **KIIPFNRL** | KIIPFNRL | 8 | 17.3 | 0.05 | | YES | | YES | | YES | Q8VCM7 |
| **KIITYRNL** | KIITYRNL | 8 | 7.5 | 0.02 | | YES | | YES | | YES | Q8BFV2 |
| **KILTFDQL** | KILTFDQL | 8 | 66.5 | 0.2 | 3864400 | YES | 55083000 | YES | 35815000 | NO | P35980 |
| **KIQSFINRM** | KIQSFINRM | 9 | 202.8 | 0.5 | 2194719.57 | YES | 14078000 | YES | | YES | Q8C1Y8 |

| | | | | | | | | | | | |
|---|---|---|---|---|---|---|---|---|---|---|---|
| **KIVPFFKL** | KIVPFFKL | 8 | 139.2 | 0.4 | 1664143.01 | YES | 21855000 | YES | 177590 | YES | Q9JHU4 |
| **KIYQWINEL** | KIYQWINEL | 9 | 89.5 | 0.25 | 2946700 | YES | 80391000 | YES | | YES | Q9JKY0 |
| **KNFAFLEF** | KNFAFLEF | 8 | 247.3 | 0.6 | | YES | 5128900 | YES | 498150 | NO | P26369 |
| **KNFAFTLV** | KNFAFTLV | 8 | 14.6 | 0.05 | | YES | 22810000 | YES | | YES | A3FIN4;Q148W0 |
| **KNFDKLSFL** | KNFDKLSFL | 9 | 146.9 | 0.4 | 876220 | NO | 3378800 | YES | 628630 | YES | Q8CIE6 |
| **KNFPFERL** | KNFPFERL | 8 | 11.2 | 0.04 | 1317400 | YES | 29532000 | YES | 6481900 | YES | Q6P4T2 |
| **KNFVYRTL** | KNFVYRTL | 8 | 5.9 | 0.015 | 993725.768 | YES | | YES | | YES | A2RSX7 |
| **KNHEFIATF** | KNHEFIATF | 9 | 233.3 | 0.6 | 570736.935 | NO | 5348400 | YES | 16647000 | YES | P28867 |
| **KNIDRFIPV** | KNIDRFIPV | 9 | 82.8 | 0.25 | 183553.548 | NO | 470870 | YES | 201630 | NO | Q9CX30 |
| **KNIIYRFL** | KNIIYRFL | 8 | 25.2 | 0.08 | 1012300 | YES | 4915500 | YES | | YES | Q9D4H2 |
| **KNIRFPLM** | KNIRFPLM | 8 | 35.4 | 0.125 | 830840 | NO | 25938000 | YES | 3992100 | YES | Q6ZPT1 |
| **KNIRFPLM** | KNIRFPLM(+15.99) | 8 | 35.4 | 0.125 | 514910 | NO | 21403000 | YES | 47522000 | YES | Q6ZPT1 |
| **KNIRYVAL** | KNIRYVAL | 8 | 10.9 | 0.03 | 3511909.68 | YES | 1677900 | YES | 9440700 | YES | P22892 |
| **KNLNYLHL** | KNLNYLHL | 8 | 14.7 | 0.05 | 4629154.88 | YES | 6068500 | YES | 1621800 | YES | Q99PV0 |
| **KNLVFVVGL** | KNLVFVVGL | 9 | 42.5 | 0.125 | | YES | 6228700 | YES | 1592400 | YES | Q8BT14 |
| **KNNQFQAL** | KNNQFQAL | 8 | 149.7 | 0.4 | 82680 | YES | 336060 | YES | 527540 | NO | P17225:Q8BHD7 |
| **KNNQFQALL** | KNNQFQALL | 9 | 78.7 | 0.25 | 1133371.68 | NO | 1785100 | YES | 1219900 | YES | P17225:Q8BHD7 |
| **KNPGYIKL** | KNPGYIKL | 8 | 416.9 | 0.9 | 1776327.17 | YES | 540580 | YES | 159330 | NO | O35129 |
| **KNVLFSHL** | KNVLFSHL | 8 | 7.9 | 0.02 | 2590519.32 | YES | 24861000 | YES | 25310000 | YES | P12849;Q9DBC7 |
| **KNVTFEHV** | KNVTFEHV | 8 | 302.9 | 0.7 | 760410 | YES | 4106700 | YES | 4897800 | YES | O88967 |
| **KNVVYERV** | KNVVYERV | 8 | 337.2 | 0.8 | 173680 | YES | 6426700 | YES | | YES | Q8K4L4 |
| **KNVVYRDL** | KNVVYRDL | 8 | 172.5 | 0.5 | 1482233.27 | YES | 6426700 | YES | 10218000 | YES | P31750 |
| **KNWEFMTI** | KNWEFMTI | 8 | 83 | 0.25 | 351210.444 | NO | 2357400 | YES | | YES | Q04592 |
| **KNYDFAQV** | KNYDFAQV | 8 | 5.5 | 0.015 | 547730 | YES | 1174400 | YES | 1235900 | YES | Q8BGF7 |
| **KNYDFAQVL** | KNYDFAQVL | 9 | 29.5 | 0.09 | 4638300 | YES | 26337000 | YES | 25694000 | YES | Q8BGF7 |
| **KNYGFVHI** | KNYGFVHI | 8 | 23.9 | 0.07 | | YES | 11400000 | YES | | YES | Q8VE92 |
| **KNYGYVRV** | KNYGYVRV | 8 | 19.7 | 0.06 | 380680 | YES | 421160 | YES | 582900 | YES | Q61586 |
| **KNYLLPIL** | KNYLLPIL | 8 | 132.1 | 0.4 | | YES | 5267500 | YES | 2686000 | YES | P50172 |
| **KNYSFPLNNL** | KNYSFPLNNL | 10 | 25 | 0.08 | 176680 | YES | 15035000 | YES | 29275000 | YES | Q8CDG3 |

| | | | | | | | | | | | |
|---|---|---|---|---|---|---|---|---|---|---|---|
| **KQFAFVHM** | KQFAFVHM | 8 | 32.1 | 0.1 | 387167.806 | NO | 153870 | NO | 57969 | NO | Q8C2Q3 |
| **KQFEYIEV** | KQFEYIEV | 8 | 309.5 | 0.8 | 63090 | YES | | YES | | YES | P42859 |
| **KQFSYTHI** | KQFSYTHI | 8 | 49.5 | 0.15 | 707889.866 | YES | | YES | 243110 | NO | Q99LC5 |
| **KSFDYGNL** | KSFDYGNL | 8 | 4.3 | 0.01 | 219027.895 | NO | | YES | 9577200 | YES | Q3UHF7 |
| **KSFEWLSQM** | KSFEWLSQM | 9 | 38.9 | 0.125 | 2019626.58 | YES | 43332000 | YES | | YES | Q9JHU4 |
| **KSFEWLSQM** | KSFEWLSQM(+15.99) | 9 | 38.9 | 0.125 | 1101600 | YES | 24455000 | YES | 459290 | YES | Q9JHU4 |
| **KSFLFSAL** | KSFLFSAL | 8 | 2.4 | 0.01 | | YES | 66535000 | YES | 1457600 | YES | Q920L5 |
| **KSIAFPSI** | KSIAFPSI | 8 | 68.3 | 0.2 | | YES | | YES | 2635400 | YES | Q9QZQ8 |
| **KSITFSKL** | KSITFSKL | 8 | 7.5 | 0.02 | 13769437.6 | YES | 9271700 | YES | 7622800 | YES | O35459 |
| **KSLAFQKL** | KSLAFQKL | 8 | 13.2 | 0.04 | 3467776.12 | YES | | YES | 1125400 | YES | Q8BIL5 |
| **KSLERATQL** | KSLERATQL | 9 | 327.8 | 0.8 | | YES | | YES | | YES | Q63829 |
| **KSLSFPKL** | KSLSFPKL | 8 | 13.4 | 0.04 | 373873.203 | NO | | YES | 3151300 | YES | Q8BH48 |
| **KSPEYESL** | KSPEYESL | 8 | 99.2 | 0.3 | 2194485.85 | YES | 2308100 | YES | 9309200 | YES | Q3V1L4 |
| **KSYLMNKL** | KSYLM(+15.99)NKL | 8 | 29.4 | 0.09 | 243350.045 | NO | 94214000 | YES | 6447900 | NO | Q9Z0E6:Q8CFB4:Q01514 |
| **KSYLMNKL** | KSYLMNKL | 8 | 29.4 | 0.09 | 254591.833 | NO | 199660000 | YES | 1684400 | NO | Q9Z0E6:Q8CFB4:Q01514 |
| **KSYLMNRL** | KSYLMNRL | 8 | 9.9 | 0.03 | 829929.098 | YES | 56274000 | YES | 2711700 | NO | Q61107 |
| **KSYSFDEV** | KSYSFDEV | 8 | 26.2 | 0.08 | 729510 | YES | 9297900 | YES | 1468200 | YES | Q99KQ4 |
| **KSYSFIARM** | KSYSFIARM | 9 | 4.4 | 0.01 | 3179875.71 | YES | 12133000 | YES | 130240 | NO | Q8CGZ0 |
| **KSYSFIARM** | KSYSFIARM(+15.99) | 9 | 4.4 | 0.01 | 1361362.62 | YES | 5948400 | YES | 1743900 | NO | Q8CGZ0 |
| **KTFDFKGL** | KTFDFKGL | 8 | 19.5 | 0.06 | 531978.865 | NO | 3453100 | YES | 2483500 | YES | Q924W7 |
| **KTFLFSATM** | KTFLFSATM | 9 | 17.4 | 0.05 | 230550.143 | NO | | YES | 1120300 | YES | Q9CWX9 |
| **KTFLFSATM** | KTFLFSATM(+15.99) | 9 | 17.4 | 0.05 | 322560 | NO | 4847700 | YES | 24311000 | YES | Q9CWX9 |
| **KTFSYAGF** | KTFSYAGF | 8 | 16.7 | 0.05 | 667028.343 | NO | 3334400 | YES | | YES | P80313 |
| **KTLVLSNL** | KTLVLSNL | 8 | 71.3 | 0.2 | 1051843.4 | YES | 2703600 | YES | | YES | P09405 |
| **KTVCFQNL** | KTVC(+119.00)FQNL | 8 | 21.2 | 0.06 | 516943.07 | NO | 17106000 | NO | 1960800 | NO | Q7TMB8 |
| **KTVEYTRL** | KTVEYTRL | 8 | 24.6 | 0.07 | 139891.316 | YES | 491770 | YES | | YES | A8C756 |
| **KTVIFENL** | KTVIFENL | 8 | 33.4 | 0.1 | 1172869.86 | NO | 5900500 | YES | 5046500 | YES | Q99KZ6 |
| **KTWRFSNM** | KTWRFSNM | 8 | 6.1 | 0.015 | 5095420.32 | YES | 7395700 | YES | 1389300 | YES | Q8CIB5 |

| | | | | | | | | | | | |
|---|---|---|---|---|---|---|---|---|---|---|---|
| **KTWRFSNM** | KTWRFSNM(+15.99) | 8 | 6.1 | 0.015 | 5972248.63 | YES | 4587600 | YES | 7300200 | YES | Q8CIB5:Q8K1B8 |
| **KTYEHFNAM** | KTYEHFNAM | 9 | 16.8 | 0.05 | 4290265.22 | YES | 2736100 | YES | 505990 | YES | P37040 |
| **KTYEHFNAM** | KTYEHFNAM(+15.99) | 9 | 16.8 | 0.05 | 4691300 | YES | 1624100 | YES | 12153000 | YES | P37040 |
| **KTYQFLNDI** | KTYQFLNDI | 9 | 89.3 | 0.25 | 792620 | YES | 15599000 | YES | 18020000 | YES | Q3UFY0 |
| **KTYSFLITL** | KTYSFLITL | 9 | 15.3 | 0.05 | | YES | | YES | | YES | P27656 |
| **KVFEYHNV** | KVFEYHNV | 8 | 18.2 | 0.06 | | YES | 9587700 | YES | 15290000 | YES | Q9DAX9 |
| **KVFQFLNA** | KVFQFLNA | 8 | 143.9 | 0.4 | 775550 | YES | 6631400 | YES | 4279000 | NO | Q8BP67 |
| **KVIEFKKL** | KVIEFKKL | 8 | 122.5 | 0.4 | 438219.391 | YES | 487110 | YES | 1507900 | NO | Q8R4D1 |
| **KVITFIDL** | KVITFIDL | 8 | 135.5 | 0.4 | 820930.993 | YES | 38822000 | YES | 14261000 | YES | O08582 |
| **KVLEFERV** | KVLEFERV | 8 | 211.9 | 0.6 | 5696462.56 | YES | 12869000 | YES | 8690800 | YES | Q8CEC6 |
| **KVLIFSQM** | KVLIFSQM(+15.99) | 8 | 42.8 | 0.125 | 29475 | NO | 766590 | YES | 8739700 | YES | Q09XV5:Q8BYH8 |
| **KVLRFIAEV** | KVLRFIAEV | 9 | 107.3 | 0.3 | 506542.663 | NO | 15107000 | YES | 5355900 | NO | Q9QYH6 |
| **KVQEFQRL** | KVQEFQRL | 8 | 59.9 | 0.175 | | YES | 1583700 | YES | 610770 | YES | Q62172 |
| **KVQEFVLL** | KVQEFVLL | 8 | 198.9 | 0.5 | 844630 | YES | | YES | | YES | Q3UHQ6 |
| **KVVDHFGRL** | KVVDHFGRL | 9 | 53.1 | 0.175 | 653706.473 | YES | | YES | 488000 | NO | Q8VCC1 |
| **KVVEFSEL** | KVVEFSEL | 8 | 65.2 | 0.2 | | YES | 4187700 | YES | | YES | Q3UFM5 |
| **KVYLYTHL** | KVYLYTHL | 8 | 2.7 | 0.01 | | YES | 8677600 | YES | | YES | Q8R0A7 |
| **KVYNYNHL** | KVYNYNHL | 8 | 4 | 0.01 | 122070 | YES | 11420000 | YES | 46183000 | YES | P61358 |
| **KVYTFNSV** | KVYTFNSV | 8 | 10.5 | 0.03 | 350154.822 | NO | 1761700 | YES | 2781200 | YES | G5E829:Q9R0K7:Q6Q477 |
| **LAPHFNSL** | LAPHFNSL | 8 | 87.3 | 0.25 | 163330 | YES | 1633500 | YES | | YES | Q6P6J9 |
| **LAPVFQRV** | LAPVFQRV | 8 | 64.3 | 0.2 | 248310 | YES | | YES | | YES | Q9JJA2 |
| **LAPVYQRL** | LAPVYQRL | 8 | 17.7 | 0.05 | 3609900 | YES | 98327000 | YES | | YES | P82198 |
| **LGYKYVGM** | LGYKYVGM | 8 | 10.2 | 0.03 | 446672.156 | NO | 4696300 | YES | 773390 | YES | Q9CX30 |
| **LGYKYVGM** | LGYKYVGM(+15.99) | 8 | 10.2 | 0.03 | 367700 | NO | 4401900 | YES | 6116600 | YES | Q9CX30 |
| **LGYQYPSL** | LGYQYPSL | 8 | 5.3 | 0.015 | 6405600 | YES | 3584700 | YES | 2941100 | YES | Q9QYG0 |
| **LIYKFLNV** | LIYKFLNV | 8 | 5.8 | 0.015 | 6060300 | YES | 37941000 | YES | 871200 | YES | Q6P5F9 |
| **LQYEFTKL** | LQYEFTKL | 8 | 10.8 | 0.03 | 1714119.49 | YES | 168370000 | YES | 273580000 | YES | A2APV2;Q6ZPF4 |
| **LQYIFAHV** | LQYIFAHV | 8 | 4.7 | 0.01 | | YES | | YES | 9228300 | YES | Q99PV0 |
| **LSLPFEARL** | LSLPFEARL | 9 | 10.9 | 0.03 | 1331914.28 | YES | 8527400 | YES | | YES | Q9JL15 |

| | | | | | | | | | | | |
|---|---|---|---|---|---|---|---|---|---|---|---|
| **LSPKYIKM** | LSPKYIKM(+15.99) | 8 | 152.1 | 0.4 | 320140 | YES | 1832800 | YES | 1747400 | NO | P60843 |
| **LSPPSYSKL** | LSPPSYSKL | 9 | 148.2 | 0.4 | 2030260.7 | NO | 7906600 | NO | 5303000 | NO | P98195 |
| **LSPSHYALL** | LSPSHYALL | 9 | 5.4 | 0.015 | 6995929.13 | YES | 7231100 | YES | 9300300 | YES | Q8C7X2 |
| **LSYDYSGRFL** | LSYDYSGRFL | 10 | 91.5 | 0.25 | 108265.509 | NO | | YES | 155860 | YES | Q80UJ9 |
| **LSYSYQSRF** | LSYSYQSRF | 9 | 28.8 | 0.09 | 648330.982 | YES | 4388600 | YES | 3918200 | YES | Q921M3 |
| **LTQQYHQL** | LTQQYHQL | 8 | 206.2 | 0.5 | 103110 | NO | | YES | 608060 | YES | P10404:P11370 |
| **LVAIFTHL** | LVAIFTHL | 8 | 58.8 | 0.175 | | YES | 37825000 | YES | 2915000 | YES | Q9JHU4 |
| **LVYKNFPQL** | LVYKNFPQL | 9 | 17.5 | 0.05 | 386380.959 | NO | 7943600 | YES | 2906300 | YES | Q8K1A5 |
| **LVYQFKEM** | LVYQFKEM | 8 | 27 | 0.08 | 356161.349 | NO | 7286300 | YES | 1157800 | NO | Q60775 |
| **LVYQFKEM** | LVYQFKEM(+15.99) | 8 | 27 | 0.08 | 159426.162 | NO | 3676000 | YES | 25991000 | NO | Q60775 |
| **MSFQFAHL** | M(+15.99)SFQFAHL | 8 | 1.8 | 0.01 | 1345800 | YES | 7670100 | YES | 2189500 | YES | Q80TY5 |
| **MAYLFRNI** | MAYLFRNI | 8 | 3.1 | 0.01 | 598210 | YES | | YES | | YES | Q9D0M1 |
| **MSFQFAHL** | MSFQFAHL | 8 | 1.8 | 0.01 | 1312200 | YES | 20512000 | YES | | YES | Q80TY5 |
| **MSYLFRNI** | MSYLFRNI | 8 | 2.4 | 0.01 | 39090 | YES | 3542300 | YES | | YES | Q8R574 |
| **NIFMFSKV** | NIFMFSKV | 8 | 55.6 | 0.175 | 299120 | YES | 2348400 | YES | | YES | Q9D710 |
| **NMVPFPRL** | NMVPFPRL | 8 | 315 | 0.8 | | YES | | YES | | YES | P68372 |
| **NNPIFRYL** | NNPIFRYL | 8 | 160.9 | 0.5 | 2092818.95 | YES | 1952300 | YES | | YES | Q8VCH6 |
| **NNYVFKNAL** | NNYVFKNAL | 9 | 27.4 | 0.08 | 98714.2739 | NO | 1492600 | YES | | YES | Q9JJJ7 |
| **NNYVYAGL** | NNYVYAGL | 8 | 3.9 | 0.01 | 400870 | YES | 5269200 | YES | 4679900 | YES | Q8CIK8 |
| **NSFRYNGL** | NSFRYNGL | 8 | 9.3 | 0.025 | 495930 | YES | 29392000 | YES | 5432100 | YES | P41105 |
| **NSPEFQKL** | NSPEFQKL | 8 | 386.5 | 0.9 | 307090 | YES | 2015800 | YES | 559980 | YES | P42859 |
| **NSPEYQRL** | NSPEYQRL | 8 | 126.3 | 0.4 | 2523300 | YES | 2664600 | YES | 5966800 | YES | Q9CVD2 |
| **NTHEFVNL** | NTHEFVNL | 8 | 116.6 | 0.4 | 479790 | YES | 16164000 | YES | | YES | P40336 |
| **NTPKYAKL** | NTPKYAKL | 8 | 171.4 | 0.5 | 257790 | NO | 148120 | NO | | YES | Q9R020 |
| **NTYKYAKI** | NTYKYAKI | 8 | 37.2 | 0.125 | 638610 | YES | 579670 | YES | | YES | Q7TQI7 |
| **NTYSYQKV** | NTYSYQKV | 8 | 134.4 | 0.4 | 1247300 | YES | | YES | 2852700 | YES | Q3TA59 |
| **QAFDFEFTHV** | QAFDFEFTHV | 10 | 101.1 | 0.3 | 1175207.03 | NO | 11974000 | YES | | YES | Q8C0E2 |
| **QAIDYHEL** | QAIDYHEL | 8 | 442.9 | 1 | 728418.01 | NO | | YES | 488040 | NO | O35648 |
| **QALKYFNL** | QALKYFNL | 8 | 11.4 | 0.04 | 19960000 | YES | 100420000 | YES | 70000000 | YES | Q9Z2G6 |

| | | | | | | | | | | | |
|---|---|---|---|---|---|---|---|---|---|---|---|
| **QALSRFPVM** | QALSRFPVM | 9 | 235.4 | 0.6 | 97775.5106 | NO | 2015900 | YES | 66467 | YES | Q91VH2 |
| **QGGQIYVHL** | QGGQIYVHL | 8 | 99.7 | 0.3 | | YES | | YES | | YES | Q922H4 |
| **QGYTVARI** | QGYTVARI | 8 | 157.2 | 0.4 | | YES | | YES | | YES | Q9Z2G6 |
| **QIIPFKTL** | QIIPFKTL | 8 | 139.6 | 0.4 | 37205410.6 | YES | 137410000 | YES | 50332000 | YES | Q8BVY0 |
| **QIVSFYRV** | QIVSFYRV | 8 | 98.7 | 0.3 | 499725.917 | NO | 3389100 | NO | 714870 | YES | Q8BUR4 |
| **QIYARQYYM** | QIYARQYYM | 9 | 162.3 | 0.5 | 594069.683 | NO | 426660 | YES | | YES | Q9JJK5 |
| **QIYDIFQKL** | QIYDIFQKL | 9 | 76.4 | 0.25 | 212811.022 | YES | 95679000 | YES | 136110 | NO | P60843 |
| **QIYYYHNV** | QIYYYHNV | 8 | 7 | 0.02 | | YES | | YES | | YES | Q6WKZ8 |
| **QNAVYINL** | QNAVYINL | 8 | 52.3 | 0.175 | | YES | 6268600 | YES | 422970 | YES | Q3U0M1 |
| **QNHVFPLL** | QNHVFPLL | 8 | 48.9 | 0.15 | 3090907.44 | YES | 51268000 | YES | 10677000 | YES | Q7TMY7 |
| **QNPNYYNL** | QNPNYYNL | 8 | 16.9 | 0.05 | 1875500 | YES | 2084800 | YES | 2953300 | YES | Q6P4T2 |
| **QNPRFSKL** | QNPRFSKL | 8 | 28 | 0.08 | | YES | | YES | | YES | Q8BFT2 |
| **QNYEMPNL** | QNYEM(+15.99)PNL | 8 | 34.5 | 0.1 | 5676400 | YES | 23478000 | YES | 75330000 | YES | P0DP99 |
| **QNYEMPNL** | QNYEMPNL | 8 | 34.5 | 0.1 | 483130 | YES | 64019000 | YES | 15229000 | YES | P0DP99 |
| **QNYLFGCEL** | QNYLFGC(+119.00)EL | 9 | 36.3 | 0.125 | 142247.962 | NO | 9810500 | NO | 1933000 | NO | Q61937 |
| **QQFIYEKL** | QQFIYEKL | 8 | 114.5 | 0.3 | 5746711.71 | YES | 25431000 | YES | | YES | Q8K2V6 |
| **QQIAFKNL** | QQIAFKNL | 8 | 82.2 | 0.25 | 202285.966 | NO | 1261100 | NO | 949770 | NO | Q08639:Q64163 |
| **QQYLFDRL** | QQYLFDRL | 8 | 13.1 | 0.04 | 393189.914 | NO | 3615700 | YES | 3796800 | NO | Q8BHG1 |
| **QQYRFSVI** | QQYRFSVI | 8 | 59.4 | 0.175 | 1704615 | NO | 10986000 | NO | 6041400 | NO | Q0GNC1 |
| **QQYRFSVIM** | QQYRFSVIM(+15.99) | 9 | 181.2 | 0.5 | 359200 | NO | 2008800 | NO | 2414200 | NO | Q0GNC1 |
| **QQYVFINQM** | QQYVFINQM | 9 | 58.4 | 0.175 | 1778469.58 | NO | 1481400 | NO | | YES | Q8BGR2 |
| **QRVEFAAL** | QRVEFAAL | 8 | 381 | 0.9 | | YES | | YES | | YES | E9Q7G0 |
| **QSIAFISRL** | QSIAFISRL | 9 | 13.5 | 0.04 | 18119690.2 | YES | 73708000 | YES | 66159000 | YES | Q9CR67 |
| **QSIEFSRL** | QSIEFSRL | 8 | 5.5 | 0.015 | 5254200 | YES | 188430000 | YES | 22150000 | YES | P23116 |
| **QSLAFHTL** | QSLAFHTL | 8 | 19.7 | 0.06 | 507360 | YES | 21435000 | YES | | YES | Q91WG4 |
| **QSPAFTRQL** | QSPAFTRQL | 9 | 69 | 0.2 | | YES | 536170 | YES | 208770 | YES | Q6ZQ93 |
| **QSPEFQSL** | QSPEFQSL | 8 | 32.7 | 0.1 | 944420 | YES | | YES | 7284600 | YES | Q8K1J6 |
| **QSPGFYRNV** | QSPGFYRNV | 9 | 10.7 | 0.03 | 523097.618 | YES | 5525900 | YES | 1784200 | YES | P32921 |
| **QSYEFFHL** | QSYEFFHL | 8 | 2.6 | 0.01 | 463850 | YES | 63805000 | YES | | YES | Q8BHB0 |

| | | | | | | | | | | | |
|---|---|---|---|---|---|---|---|---|---|---|---|
| **QTFVFHVV** | QTFVFHVV | 8 | 139.8 | 0.4 | 770430 | NO | | YES | 1294100 | NO | A2AAE1 |
| **QTLKYLAV** | QTLKYLAV | 8 | 287.6 | 0.7 | 957499.632 | NO | 5258500 | YES | | YES | Q9DBG6 |
| **QTYDYRNI** | QTYDYRNI | 8 | 20.8 | 0.06 | 330510 | NO | 14079000 | NO | 41363000 | NO | Q9JHI7 |
| **QVVEFKKL** | QVVEFKKL | 8 | 449.9 | 1 | 1110233.7 | YES | 1734000 | YES | 433180 | NO | Q62559 |
| **QVVQFNRL** | QVVQFNRL | 8 | 36.9 | 0.125 | 2988071.96 | YES | 26351000 | YES | | YES | O88653 |
| **QVYGFLEV** | QVYGFLEV | 8 | 191.7 | 0.5 | 768960 | NO | 36730000 | YES | | YES | Q9CQE7 |
| **RAFDYFNL** | RAFDYFNL | 8 | 5.8 | 0.015 | 3266312.05 | YES | 23216000 | YES | | YES | Q9Z2G6 |
| **RAFEFTYV** | RAFEFTYV | 8 | 10.4 | 0.03 | | YES | 30650000 | YES | 29351000 | NO | Q8VDR7 |
| **RAFSFRTV** | RAFSFRTV | 8 | 15.3 | 0.05 | 1576400 | YES | 45215000 | YES | 3954900 | NO | P46735 |
| **RAFVFDVL** | RAFVFDVL | 8 | 52.6 | 0.175 | | YES | 2349400 | YES | | YES | Q9DC50 |
| **RAIAFQHL** | RAIAFQHL | 8 | 11.2 | 0.04 | 2728918.75 | YES | 19771000 | YES | 19342000 | YES | Q7TPV4 |
| **RALNYTHL** | RALNYTHL | 8 | 8.8 | 0.025 | | YES | 6634100 | YES | 3045500 | YES | Q6PFD9 |
| **RAPAFHQL** | RAPAFHQL | 8 | 65 | 0.2 | 1423687.16 | YES | 11701000 | YES | 3290100 | YES | Q3UHH1 |
| **RAPSYRTL** | RAPSYRTL | 8 | 60.3 | 0.175 | 1515654.8 | YES | 3742000 | YES | | YES | Q924W7 |
| **RAPVYARI** | RAPVYARI | 8 | 25 | 0.08 | 32588720.6 | YES | 4311100 | YES | 3352300 | YES | P52840 |
| **RAVEYNTL** | RAVEYNTL | 8 | 140.1 | 0.4 | 42497.5427 | NO | | YES | | YES | |
| **RAVLFVGL** | RAVLFVGL | 8 | 31.2 | 0.09 | 20940 | YES | 3377300 | YES | | YES | P47758 |
| **RAYFFVEV** | RAYFFVEV | 8 | 40.5 | 0.125 | 96190.1302 | NO | | YES | | YES | |
| **RAYLFAHV** | RAYLFAHV | 8 | 2.5 | 0.01 | 2292900 | YES | 58967000 | YES | 13272000 | YES | Q5SWD9 |
| **RAYLFNSV** | RAYLFNSV | 8 | 7 | 0.02 | 763340 | YES | 34049000 | YES | 31992000 | YES | Q65Z95:Q65Z93 |
| **RGFEFTLM** | RGFEFTLM | 8 | 18.5 | 0.06 | | YES | 54087000 | YES | | YES | O55131 |
| **RGLDYFSSL** | RGLDYFSSL | 9 | 14.6 | 0.05 | 309920 | YES | | YES | | YES | B1AZA5 |
| **RGLDYYTGV** | RGLDYYTGV | 9 | 32.3 | 0.1 | 208160.054 | NO | | YES | 107980 | YES | Q61035:Q99KK9 |
| **RGPTYVNM** | RGPTYVNM | 8 | 78.8 | 0.25 | 514527.995 | YES | 7860700 | YES | 3316600 | YES | Q8K2K6 |
| **RGYAFVTF** | RGYAFVTF | 8 | 99.6 | 0.3 | 572020 | YES | | YES | | YES | Q5YD48:Q7TMK9 |
| **RGYDFAAV** | RGYDFAAV | 8 | 6.8 | 0.02 | | YES | 2320700 | YES | | YES | Q9QXY6 |
| **RGYEFIVRL** | RGYEFIVRL | 9 | 14.8 | 0.05 | 316510 | NO | 8938600 | YES | | YES | Q9JKK8 |
| **RGYEFLGV** | RGYEFLGV | 8 | 15.9 | 0.05 | | YES | 29871000 | YES | 6673800 | YES | Q8K3B1 |
| **RGYIFSLV** | RGYIFSLV | 8 | 7.5 | 0.02 | | YES | | YES | | YES | Q8VDR9 |

| | | | | | | | | | | | |
|---|---|---|---|---|---|---|---|---|---|---|---|
| **RGYSFTTT** | RGYSFTTT | 8 | 277.1 | 0.7 | 1137000 | YES | 13746000 | YES | 14272000 | YES | P60710:P63260 |
| **RGYSFTTTA** | RGYSFTTTA | 9 | 360.9 | 0.8 | 335450 | YES | | YES | 14615000 | YES | P60710:P63260 |
| **RGYSYDLKV** | RGYSYDLKV | 9 | 330.6 | 0.8 | | YES | 1653900 | YES | | YES | O55234 |
| **RIFDFQGL** | RIFDFQGL | 8 | 7.2 | 0.02 | 1180426.7 | YES | 2014700 | YES | | YES | Q9DBL1 |
| **RIILFDRL** | RIILFDRL | 8 | 29 | 0.09 | 520430 | NO | 8262800 | YES | | YES | Q8BM85 |
| **RIYDITNV** | RIYDITNV | 8 | 379.5 | 0.9 | 34625.5644 | NO | | YES | | YES | Q61502:Q8R0K9 |
| **RIYGFTAV** | RIYGFTAV | 8 | 13.3 | 0.04 | 2799300 | YES | | YES | | YES | Q69ZR2 |
| **RIYGKFLGL** | RIYGKFLGL | 9 | 19.2 | 0.06 | 13972000 | NO | 45415000 | NO | 2017800 | YES | Q60996:Q61151:Q6PD03 |
| **RNFIFSRL** | RNFIFSRL | 8 | 2.9 | 0.01 | 376720 | NO | 40867000 | YES | 6231000 | YES | Q60848 |
| **RNIHYNSV** | RNIHYNSV | 8 | 386.2 | 0.9 | | YES | 336660 | YES | 482900 | YES | Q3U2S4 |
| **RNLEFHEL** | RNLEFHEL | 8 | 79.3 | 0.25 | 8936559.35 | YES | 31724000 | YES | 8032400 | YES | Q8CCN5 |
| **RNLQFVGV** | RNLQFVGV | 8 | 66.8 | 0.2 | 105890 | YES | 5556900 | YES | | YES | Q8BX02 |
| **RNPQFQKL** | RNPQFQKL | 8 | 93.2 | 0.25 | 1537468.39 | YES | 2131900 | YES | | YES | P06745 |
| **RNPTFKVL** | RNPTFKVL | 8 | 415.2 | 0.9 | | YES | | YES | | YES | Q08288 |
| **RNPTFMGL** | RNPTFM(+15.99)GL | 8 | 12.1 | 0.04 | 6459300 | YES | 12637000 | YES | 18217000 | NO | P17427 |
| **RNQVYTQL** | RNQVYTQL | 8 | 49 | 0.15 | 9031200 | YES | 4265100 | YES | 22223000 | YES | Q91ZV0 |
| **RNVESYTKL** | RNVESYTKL | 9 | 327.9 | 0.8 | 101370 | YES | 6159700 | YES | 3649200 | YES | Q91YQ5 |
| **RNYEYLIRL** | RNYEYLIRL | 9 | 11.9 | 0.04 | 18009000 | YES | 144190000 | YES | 12631000 | YES | Q8R3L2 |
| **RNYLHYSL** | RNYLHYSL | 8 | 6.6 | 0.02 | 523175.524 | NO | 7273400 | YES | | YES | P14685 |
| **RNYQFDFL** | RNYQFDFL | 8 | 11.1 | 0.04 | 7670600 | YES | 245290000 | YES | 2347000 | YES | Q8K4Z5 |
| **RNYSYEKL** | RNYSYEKL | 8 | 11.6 | 0.04 | 2583200 | YES | 232040000 | YES | 156990000 | YES | Q02257 |
| **RQYIFSKL** | RQYIFSKL | 8 | 6.8 | 0.02 | 1073400 | YES | 185760000 | YES | 13740000 | YES | Q80SU7 |
| **RQYMFSSL** | RQYMFSSL | 8 | 4 | 0.01 | 8404074.1 | YES | 9799300 | YES | 3452900 | NO | P27641 |
| **RSFDFIHL** | RSFDFIHL | 8 | 4 | 0.01 | 4750687.73 | YES | 26675000 | YES | | YES | A2RSY6 |
| **RSIDQFANL** | RSIDQFANL | 9 | 10.9 | 0.03 | 348980 | YES | 7410300 | YES | | YES | Q8VC85 |
| **RSISFSNM** | RSISFSNM(+15.99) | 8 | 6.2 | 0.015 | 39677 | NO | 334580 | YES | 7228800 | YES | O35242 |
| **RSIWFQQL** | RSIWFQQL | 8 | 7.2 | 0.02 | 197520 | YES | 4651600 | YES | | YES | Q9CWP6 |
| **RSLKFYSL** | RSLKFYSL | 8 | 5.7 | 0.015 | 4659700 | NO | 59177000 | NO | 25266000 | NO | Q99KP6 |
| **RSLQFPEL** | RSLQFPEL | 8 | 15.3 | 0.05 | | YES | | YES | | YES | Q8CHI8 |

| | | | | | | | | | | | |
|---|---|---|---|---|---|---|---|---|---|---|---|
| **RSPAFTSRL** | RSPAFTSRL | 9 | 19 | 0.06 | 1022044.48 | NO | 6719300 | YES | 3207300 | NO | Q7TMY8 |
| **RSPEYLSL** | RSPEYLSL | 8 | 28.5 | 0.09 | 12999729.5 | YES | 60241000 | YES | 19620000 | YES | Q9JLV5 |
| **RSPKYLEL** | RSPKYLEL | 8 | 58 | 0.175 | 4948957.66 | YES | | YES | | YES | Q7TPV4 |
| **RSPWFTTL** | RSPWFTTL | 8 | 11 | 0.03 | 5107106.17 | YES | 13490000 | YES | | YES | P10404 |
| **RSTIFYYV** | RSTIFYYV | 8 | 160.4 | 0.4 | 578722.266 | NO | 3427500 | YES | | YES | Q69ZR2 |
| **RSYDFEFM** | RSYDFEFM | 8 | 21.5 | 0.07 | 12712000 | YES | 164560000 | YES | 9099500 | YES | P40336 |
| **RSYDFEFM** | RSYDFEFM(+15.99) | 8 | 21.5 | 0.07 | 9694000 | YES | 108350000 | YES | 4269300 | YES | P40336 |
| **RSYLFLGGI** | RSYLFLGGI | 9 | 9.4 | 0.025 | | YES | 24464000 | YES | 681470 | YES | Q9D2C7 |
| **RSYNMPSL** | RSYNMPSL | 8 | 23 | 0.07 | 1023875.26 | YES | 2064200 | YES | | YES | Q91ZV0 |
| **RSYQQALL** | RSYQQALL | 8 | 13.6 | 0.04 | | YES | 6597600 | YES | 2410500 | YES | Q9ERC3 |
| **RSYSFLNSSL** | RSYSFLNSSL | 10 | 18.3 | 0.06 | 180990.451 | NO | 7783200 | YES | 9934300 | YES | Q9D7J6 |
| **RSYSFQKV** | RSYSFQKV | 8 | 6.4 | 0.015 | | YES | 1321500 | YES | 5133000 | YES | Q8CI75 |
| **RSYVFSSL** | RSYVFSSL | 8 | 2.2 | 0.01 | | YES | 3773300 | YES | | YES | Q91VM4 |
| **RTFEFQLM** | RTFEFQLM | 8 | 16.3 | 0.05 | 240327.305 | NO | 8332600 | YES | | YES | Q8C5N5 |
| **RTGTYRQL** | RTGTYRQL | 8 | 371.7 | 0.9 | 444310 | YES | 1043600 | YES | | YES | P68373:P05213 |
| **RTLIYITL** | RTLIYITL | 8 | 61.4 | 0.175 | 11658000 | YES | | YES | | YES | Q9JM76 |
| **RTTEFTNL** | RTTEFTNL | 8 | 37.5 | 0.125 | 109050 | YES | 1858700 | YES | | YES | O35231 |
| **RTYSFLNL** | RTYSFLNL | 8 | 2.9 | 0.01 | 1877700 | YES | 18415000 | YES | | YES | Q810L4 |
| **RTYTYEKL** | RTYTYEKL | 8 | 9.4 | 0.03 | 15970000 | YES | 857150000 | YES | 1299800000 | YES | Q02248 |
| **RVAEFTTNL** | RVAEFTTNL | 9 | 132.4 | 0.4 | 173640 | YES | 101240000 | YES | 119620000 | YES | Q8VDD5 |
| **RVDVFTNL** | RVDVFTNL | 8 | 115.5 | 0.3 | 2554098.42 | YES | 16254000 | YES | 5811000 | YES | Q5U4D9 |
| **RVFNYNTL** | RVFNYNTL | 8 | 10.5 | 0.03 | 286096.886 | NO | | YES | | YES | O55029 |
| **RVIDFFTV** | RVIDFFTV | 8 | 373.1 | 0.9 | | YES | 38810000 | YES | 1509700 | YES | Q80Y17 |
| **RVIDFTVL** | RVIDFTVL | 8 | 246.4 | 0.6 | 9923000 | YES | | YES | | YES | Q3TJ91 |
| **RVIDFVAQV** | RVIDFVAQV | 9 | 107.5 | 0.3 | 169390.297 | NO | 3963600 | YES | 3535200 | YES | E9PYK3 |
| **RVLEYLAV** | RVLEYLAV | 8 | 359.9 | 0.8 | 359420 | YES | 628370 | YES | | YES | Q9DBN5 |
| **RVLIFSQM** | RVLIFSQM | 8 | 42.3 | 0.125 | | YES | 56619000 | YES | 22278000 | YES | Q6PDQ2 |
| **RVLIFSQM** | RVLIFSQM(+15.99) | 8 | 42.3 | 0.125 | 1802100 | YES | 26779000 | YES | 128460000 | YES | Q6PDQ2:Q91ZW3:Q6PGB8:A2AJK6:P40201 |
| **RVLLFSQM** | RVLLFSQM | 8 | 31.8 | 0.09 | | YES | 56619000 | YES | 22278000 | YES | Q9CXF7 |

| | | | | | | | | | | | |
|---|---|---|---|---|---|---|---|---|---|---|---|
| **RVLLFSQM** | RVLLFSQM(+15.99) | 8 | 31.8 | 0.09 | 1802100 | YES | 26779000 | YES | 128460000 | YES | Q9CXF7 |
| **RVMEYINRL** | RVMEYINRL | 9 | 78.4 | 0.25 | 418950 | YES | 74227000 | YES | | YES | Q68FD5 |
| **RVYEFLDKL** | RVYEFLDKL | 9 | 34.9 | 0.125 | 14797000 | YES | 99013000 | YES | 1817000 | YES | P14685 |
| **SAARFALL** | SAARFALL | 8 | 5.9 | 0.015 | 414870 | NO | | YES | 67638000 | YES | Q924N4 |
| **SAFEFNEL** | SAFEFNEL | 8 | 5.6 | 0.015 | | YES | | YES | | YES | Q9Z2C4 |
| **SAFIFRVL** | SAFIFRVL | 8 | 5.5 | 0.015 | | YES | 10932000 | YES | | YES | Q9R0A1 |
| **SAFRFAVQL** | SAFRFAVQL | 9 | 8.5 | 0.025 | 516397.731 | NO | 19095000 | YES | | YES | Q9Z2W9 |
| **SAFSFRTL** | SAFSFRTL | 8 | 3.3 | 0.01 | 5522200 | YES | 42711000 | YES | 209650000 | YES | Q8BMI0 |
| **SAHIFSNL** | SAHIFSNL | 8 | 4.3 | 0.01 | 1432700 | YES | | YES | | YES | Q9DBB9 |
| **SAIHAVNL** | SAIHAVNL | 8 | 214.4 | 0.6 | 204770 | YES | 3539100 | YES | 1732900 | YES | Q9JJ28 |
| **SALAFGQGL** | SALAFGQGL | 9 | 30.4 | 0.09 | 73527.3709 | NO | 788180 | NO | | YES | Q80U78 |
| **SALFFHYL** | SALFFHYL | 8 | 10.4 | 0.03 | | YES | 14824000 | YES | 826850 | YES | Q6WKZ8 |
| **SALKYYQL** | SALKYYQL | 8 | 6.4 | 0.015 | 760870 | YES | 20860000 | YES | 4017300 | YES | Q6ZPU9 |
| **SALPFVKL** | SALPFVKL | 8 | 19.3 | 0.06 | 95329.2725 | NO | | YES | | YES | Q0VGY8 |
| **SALRFLNL** | SALRFLNL | 8 | 3.3 | 0.01 | 347710 | YES | 36995000 | YES | 36290000 | YES | Q3TAA7 |
| **SALTFAGL** | SALTFAGL | 8 | 2.8 | 0.01 | 2855600 | YES | 3855700 | YES | 447120 | YES | Q9CXV1 |
| **SALVFTRL** | SALVFTRL | 8 | 3.6 | 0.01 | 4060833.03 | YES | | YES | 109510000 | YES | Q9JHJ3 |
| **SAMVFSAM** | SAM(+15.99)VFSAM(+15.99) | 8 | 8.4 | 0.025 | 1023500 | NO | 13695000 | NO | 61537000 | NO | P63082 |
| **SANIFRTL** | SANIFRTL | 8 | 52.8 | 0.175 | | YES | | YES | 2409800 | YES | Q6PD03 |
| **SAPIYKRI** | SAPIYKRI | 8 | 197.1 | 0.5 | | YES | 1088800 | YES | | YES | Q9CSH3 |
| **SAPKFPSSGL** | SAPKFPSSGL | 10 | 272 | 0.7 | 552351.197 | YES | | YES | | YES | P70295 |
| **SAPLYTNL** | SAPLYTNL | 8 | 4.6 | 0.01 | 1435700 | NO | 2188100 | NO | 12994000 | YES | Q8K4S1 |
| **SAPRFLTAF** | SAPRFLTAF | 9 | 56.3 | 0.175 | 476782.698 | YES | 8143600 | YES | | YES | Q3TQR0 |
| **SAPTFINF** | SAPTFINF | 8 | 67.5 | 0.2 | | YES | 8805500 | YES | | YES | Q9CQY5 |
| **SAPVFDRL** | SAPVFDRL | 8 | 22.3 | 0.07 | | YES | 2263300 | YES | | YES | A2AIV2 |
| **SAPVFKEKL** | SAPVFKEKL | 9 | 118.3 | 0.4 | 776407.903 | YES | 1985900 | YES | 2081900 | YES | Q6PDN3 |
| **SAPWYLNRV** | SAPWYLNRV | 9 | 53.6 | 0.175 | 18719 | NO | 2089200 | YES | 598660 | YES | P29416 |
| **SASHFSQL** | SASHFSQL | 8 | 56.7 | 0.175 | 79561.165 | NO | | YES | | YES | Q3TLH4 |
| **SATAFQRI** | SATAFQRI | 8 | 245.8 | 0.6 | 1050000 | YES | | YES | | YES | Q91VS7 |

| | | | | | | | | | | | |
|---|---|---|---|---|---|---|---|---|---|---|---|
| **SATTFRLL** | SATTFRLL | 8 | 31.7 | 0.09 | 789350 | YES | 5742100 | YES | | YES | P01027 |
| **SATVFRTV** | SATVFRTV | 8 | 178 | 0.5 | 440244.939 | YES | 4323100 | YES | 3979600 | YES | Q8CDD8 |
| **SAVIFRTL** | SAVIFRTL | 8 | 13.2 | 0.04 | 5686600 | YES | 66144000 | YES | 147100000 | YES | A2AT37 |
| **SAVSFHSL** | SAVSFHSL | 8 | 17.6 | 0.05 | | YES | | YES | 2834100 | YES | Q8BZT9 |
| **SAYEFYHA** | SAYEFYHA | 8 | 13 | 0.04 | 3024142.28 | YES | 39530000 | YES | 31735000 | YES | P97481 |
| **SAYEFYHAL** | SAYEFYHAL | 9 | 2.8 | 0.01 | 1093200 | YES | 596940000 | YES | 304430000 | YES | P97481 |
| **SAYEVIKL** | SAYEVIKL | 8 | 56.8 | 0.175 | 4865800 | YES | 45863000 | YES | 2072600 | YES | P06151:P16125 |
| **SAYLFVKL** | SAYLFVKL | 8 | 3 | 0.01 | 1378462.92 | YES | 18493000 | YES | | YES | B9EJ80 |
| **SAYLYKQGF** | SAYLYKQGF | 9 | 26.3 | 0.08 | 29469 | YES | 622800 | YES | | YES | Q9D8U2 |
| **SAYNAFNRF** | SAYNAFNRF | 9 | 43.1 | 0.15 | 399071.792 | YES | 121840 | YES | | YES | Q9D1C8 |
| **SAYNYAEQTM** | SAYNYAEQTM | 10 | 123.5 | 0.4 | | YES | 1025900 | YES | | YES | Q8VE92 |
| **SAYNYAEQTM** | SAYNYAEQTM(+15.99) | 10 | 123.5 | 0.4 | 1831500 | YES | 1909700 | YES | 3082000 | YES | Q8VE92 |
| **SAYQRGESL** | SAYQRGESL | 9 | 233.2 | 0.6 | 290970 | YES | 1419000 | YES | 4757600 | YES | O35382 |
| **SAYRFSGV** | SAYRFSGV | 8 | 2.5 | 0.01 | 1681515.97 | NO | 1449100 | NO | 2430800 | NO | Q8CJF7 |
| **SDYVYPSL** | SDYVYPSL | 8 | 39.5 | 0.125 | 6086.5 | NO | | YES | 11981000 | YES | Q9WTU0 |
| **SFYEHIITV** | SFYEHIITV | 9 | 141.5 | 0.4 | 488000 | NO | 29851000 | YES | 4511200 | YES | Q8BGZ3 |
| **SFYNELRV** | SFYNELRV | 8 | 464.1 | 1 | | YES | 6560700 | YES | 536460 | YES | P63268 |
| **SGFSFRGV** | SGFSFRGV | 8 | 6.9 | 0.02 | 596562.665 | YES | | YES | | YES | P23188 |
| **SGFVFTRL** | SGFVFTRL | 8 | 2.5 | 0.01 | 6137300 | YES | 38898000 | YES | 2912400 | YES | Q6PCN7 |
| **SGIDFKQL** | SGIDFKQL | 8 | 68.8 | 0.2 | 6127900 | YES | 115270000 | YES | 11335000 | YES | O55222 |
| **SGLIFNKV** | SGLIFNKV | 8 | 97.9 | 0.3 | 94128 | YES | 2912400 | YES | 724800 | YES | P70279 |
| **SGLIFTKI** | SGLIFTKI | 8 | 97.7 | 0.3 | | YES | 5732000 | YES | | YES | P55284 |
| **SGLKYVAV** | SGLKYVAV | 8 | 39.3 | 0.125 | 2102600 | YES | 7001900 | NO | 42053000 | YES | Q923D2 |
| **SGLKYVNV** | SGLKYVNV | 8 | 14.6 | 0.05 | 14281000 | YES | 75444000 | YES | 65593000 | YES | Q8VCP8 |
| **SGLLFRSL** | SGLLFRSL | 8 | 6.5 | 0.02 | 7230300 | YES | | YES | | YES | Q9R1S7 |
| **SGLLFTHL** | SGLLFTHL | 8 | 4.1 | 0.01 | 1609647.99 | YES | 6220800 | YES | | YES | P58158 |
| **SGLTYIKI** | SGLTYIKI | 8 | 186.5 | 0.5 | 5041665.4 | YES | 15119000 | YES | 6071100 | YES | O88738 |
| **SGLVFVQV** | SGLVFVQV | 8 | 32.7 | 0.1 | | YES | | YES | | YES | O35604 |
| **SGPEYLKRL** | SGPEYLKRL | 9 | 121.3 | 0.4 | 71186.3055 | NO | 537930 | NO | 222540 | NO | Q02395 |

| | | | | | | | | | | | |
|---|---|---|---|---|---|---|---|---|---|---|---|
| **SGPTYIKL** | SGPTYIKL | 8 | 33.3 | 0.1 | 1526300 | YES | | YES | 23130000 | YES | Q6NSR3 |
| **SGVDYRGV** | SGVDYRGV | 8 | 431.9 | 1 | 1967600 | YES | 698820 | NO | 1520400 | NO | P62313 |
| **SGVEYTRL** | SGVEYTRL | 8 | 14.1 | 0.04 | 366880 | YES | 1422400 | YES | 2921000 | YES | Q62178 |
| **SGVVYSVGM** | SGVVYSVGM(+15.99) | 9 | 137.1 | 0.4 | 385360 | NO | 101610 | NO | 92725 | NO | P11276 |
| **SGYDFENRL** | SGYDFENRL | 9 | 14.2 | 0.04 | 59060000 | YES | 65547000 | YES | 15536000 | YES | Q8VE09 |
| **SGYDFSRL** | SGYDFSRL | 8 | 3.6 | 0.01 | 6539700 | YES | 26764000 | YES | 45159000 | YES | Q6GQT6 |
| **SGYDYYHV** | SGYDYYHV | 8 | 6.9 | 0.02 | 995946.079 | YES | | YES | 450710 | YES | O08785 |
| **SGYFHPLL** | SGYFHPLL | 8 | 12.1 | 0.04 | 5902912.58 | YES | 4343800 | YES | 4480500 | YES | P10518 |
| **SGYHYGLL** | SGYHYGLL | 8 | 3.7 | 0.01 | 4264556.35 | YES | 4061800 | YES | 18907000 | YES | P45448 |
| **SGYHYNAL** | SGYHYNAL | 8 | 4.2 | 0.01 | 30537000 | YES | | YES | | YES | Q60641 |
| **SGYIYHKL** | SGYIYHKL | 8 | 4.8 | 0.01 | 684360 | YES | 53924000 | YES | 134540000 | YES | Q9EPU0 |
| **SGYKFFSL** | SGYKFFSL | 8 | 2.6 | 0.01 | 48314000 | YES | 105830000 | YES | 2688800 | YES | Q80W47 |
| **SGYKFGVL** | SGYKFGVL | 8 | 4.6 | 0.01 | 2956800 | NO | 140980000 | NO | 148490000 | YES | Q9D902 |
| **SGYKYVGM** | SGYKYVGM | 8 | 4.8 | 0.01 | 796140 | YES | 20298000 | YES | 1583400 | YES | Q91XB7 |
| **SGYKYVGM** | SGYKYVGM(+15.99) | 8 | 4.8 | 0.01 | 5634600 | YES | 9435100 | YES | 57970000 | YES | Q91XB7 |
| **SGYQFIHA** | SGYQFIHA | 8 | 25.9 | 0.08 | 1033535.57 | YES | 1586700 | YES | | YES | P30561 |
| **SGYQYKRL** | SGYQYKRL | 8 | 4.4 | 0.01 | 216060 | YES | 1117200 | YES | 204340 | YES | O88974 |
| **SGYSFTHI** | SGYSFTHI | 8 | 4.5 | 0.01 | 708570 | YES | 7118300 | YES | 2177000 | YES | Q9D864 |
| **SHYDFGLRAL** | SHYDFGLRAL | 10 | 147.1 | 0.4 | 1339821.7 | YES | 30271000 | YES | 11295000 | NO | Q9JHU4 |
| **SIAAFIQRL** | SIAAFIQRL | 9 | 47.7 | 0.15 | | YES | 253990000 | YES | 24752000 | YES | Q9Z1R2 |
| **SIANFTNV** | SIANFTNV | 8 | 31.2 | 0.09 | 13411000 | YES | 15193000 | YES | 5616400 | YES | Q91V92 |
| **SIFEFVHA** | SIFEFVHA | 8 | 36.6 | 0.125 | 169980 | NO | | YES | | YES | Q91YE6 |
| **SIIALVRTL** | SIIALVRTL | 9 | 448.5 | 1 | | YES | 594990 | YES | | YES | Q9CYV5 |
| **SIIKATNL** | SIIKATNL | 8 | 83.5 | 0.25 | 2365021.36 | YES | 8149900 | YES | 10832000 | YES | Q80X82 |
| **SIINFIERL** | SIINFIERL | 9 | 21.3 | 0.06 | 680440 | YES | 33760000 | YES | | YES | Q9QZ09 |
| **SILALTHL** | SILALTHL | 8 | 47.9 | 0.15 | 183990 | NO | | YES | | YES | Q6P5B0 |
| **SILQYSNV** | SILQYSNV | 8 | 6.8 | 0.02 | 288520 | NO | 646740 | NO | 1596300 | YES | Q9WUE4 |
| **SILTYSRI** | SILTYSRI | 8 | 12.8 | 0.04 | | YES | | YES | | YES | Q8VD65 |
| **SIMAFHKL** | SIMAFHKL | 8 | 14.8 | 0.05 | 66006 | YES | 4815200 | YES | | YES | Q7TPM9 |

| | | | | | | | | | | | |
|---|---|---|---|---|---|---|---|---|---|---|---|
| **SITKFLNRI** | SITKFLNRI | 9 | 195.8 | 0.5 | 332719.534 | YES | 3509900 | YES | 4012700 | NO | Q7TNB8 |
| **SITSFPRL** | SITSFPRL | 8 | 15.6 | 0.05 | | YES | | YES | 4986200 | YES | Q99JP7 |
| **SIVQFYYM** | SIVQFYYM | 8 | 20.1 | 0.06 | 523840 | YES | 17244000 | YES | | YES | Q9R190 |
| **SIVQFYYM** | SIVQFYYM(+15.99) | 8 | 20.1 | 0.06 | 1600400 | YES | 29424000 | YES | 23029000 | YES | Q9R190 |
| **SIVSYNHL** | SIVSYNHL | 8 | 10.2 | 0.03 | 2361400 | YES | 23392000 | YES | 64640000 | YES | Q9JHU9 |
| **SIYAPARL** | SIYAPARL | 8 | 7.8 | 0.02 | | YES | | YES | 5800200 | YES | P51944 |
| **SIYAREALI** | SIYAREALI | 9 | 53.6 | 0.175 | 998906.495 | NO | 14465000 | YES | | YES | Q9CU62 |
| **SIYEKLIQF** | SIYEKLIQF | 9 | 89.8 | 0.25 | 6771171.27 | YES | 28208000 | YES | | YES | Q02614 |
| **SIYEYYHAL** | SIYEYYHAL | 9 | 3.2 | 0.01 | 42700.0975 | NO | 6213200 | YES | 453050 | YES | Q61221 |
| **SIYIPRGV** | SIYIPRGV | 8 | 224.8 | 0.6 | 1345625.68 | YES | 10569000 | YES | | YES | P50516 |
| **SIYLPQKL** | SIYLPQKL | 8 | 67.2 | 0.2 | 64955 | NO | | YES | | YES | Q8CFG9:Q8CG16 |
| **SIYPAPQV** | SIYPAPQV | 8 | 333.3 | 0.8 | 1482400 | YES | | YES | 215330 | NO | P68373:P05213:P68368 |
| **SLILFSTRL** | SLILFSTRL | 9 | 47 | 0.15 | | YES | 8325200 | YES | 14659000 | YES | Q9QUK4 |
| **SLNERFTNM** | SLNERFTNM | 9 | 311.4 | 0.8 | 1302699.65 | NO | 1036000 | NO | | YES | Q9CY57 |
| **SLVEFVHV** | SLVEFVHV | 8 | 356.1 | 0.8 | 541870 | YES | | YES | 117170 | YES | Q9D665 |
| **SLVIFMQL** | SLVIFM(+15.99)QL | 8 | 46.8 | 0.15 | 853480 | YES | 2818700 | YES | 113460 | YES | Q9R049 |
| **SLVIFMQL** | SLVIFMQL | 8 | 46.8 | 0.15 | | YES | | YES | | YES | Q9R049 |
| **SLVTFRTL** | SLVTFRTL | 8 | 85.4 | 0.25 | 680340 | YES | 59377000 | YES | 8219400 | YES | Q9QXS1 |
| **SLYSLPKL** | SLYSLPKL | 8 | 202.4 | 0.5 | 377526.979 | NO | | YES | 1673300 | YES | Q8C726 |
| **SMYVPGKL** | SM(+15.99)YVPGKL | 8 | 85.4 | 0.25 | 3819700 | YES | 46563000 | YES | 498200 | YES | Q9WU28 |
| **SMFEFSEKL** | SMFEFSEKL | 9 | 22 | 0.07 | 254821.655 | NO | 1793400 | YES | | YES | Q60674 |
| **SMVSLRAL** | SMVSLRAL | 8 | 280.7 | 0.7 | 1229974.71 | YES | | YES | | YES | P48758 |
| **SMYVPGKL** | SMYVPGKL | 8 | 85.4 | 0.25 | 4934155.58 | YES | 43835000 | YES | 4375000 | YES | Q9WU28 |
| **SNFHFAVL** | SNFHFAVL | 8 | 3.4 | 0.01 | 600020 | NO | 14741000 | YES | | YES | Q80TY5 |
| **SNFNYERSL** | SNFNYERSL | 9 | 20 | 0.06 | | YES | | YES | 959480 | YES | O35841 |
| **SNFQPPKL** | SNFQPPKL | 8 | 415.5 | 0.9 | 335566.986 | NO | 3280400 | YES | 391730 | NO | Q8BPM2:Q99JP0 |
| **SNHVFNAL** | SNHVFNAL | 8 | 11 | 0.04 | 1783900 | YES | 48367000 | YES | 19966000 | YES | Q9DBU3 |
| **SNIHYHTL** | SNIHYHTL | 8 | 23.9 | 0.07 | 25335.3133 | NO | 1614300 | YES | 192770 | YES | Q9D0N7 |
| **SNIQYITRF** | SNIQYITRF | 9 | 115 | 0.3 | 909431.833 | YES | 3853100 | YES | 3129000 | YES | Q9JM13 |

| | | | | | | | | | | | |
|---|---|---|---|---|---|---|---|---|---|---|---|
| **SNIRAGNL** | SNIRAGNL | 8 | 70.7 | 0.2 | 167230 | YES | | YES | | YES | Q78PY7 |
| **SNLELHSL** | SNLELHSL | 8 | 485 | 1.1 | 376170 | YES | 10935000 | NO | 1782800 | NO | P70195 |
| **SNLKYILV** | SNLKYILV | 8 | 44.9 | 0.15 | 714940.329 | YES | 32359000 | YES | 46739000 | YES | A2AWL7 |
| **SNLRYLSL** | SNLRYLSL | 8 | 6.4 | 0.015 | 48725 | NO | 11804000 | YES | 4033300 | NO | Q99MB1 |
| **SNLYYKYL** | SNLYYKYL | 8 | 9.4 | 0.03 | 319884.574 | NO | 6849800 | YES | 3354200 | YES | O88845 |
| **SNPEFAFL** | SNPEFAFL | 8 | 6.3 | 0.015 | 494233.568 | YES | 125970000 | YES | | YES | Q62077 |
| **SNPEFRQL** | SNPEFRQL | 8 | 17.3 | 0.05 | 2551900 | YES | 29116000 | YES | 60359000 | YES | Q3UWM4 |
| **SNPEFSSV** | SNPEFSSV | 8 | 44.6 | 0.15 | 1656000 | NO | 2817500 | YES | 5802900 | YES | P26039 |
| **SNPEYAKI** | SNPEYAKI | 8 | 69.8 | 0.2 | | YES | | YES | | YES | Q99PP2 |
| **SNTMYARL** | SNTMYARL | 8 | 5.3 | 0.015 | | YES | | YES | 155670 | YES | Q8JZN5 |
| **SNTQYARL** | SNTQYARL | 8 | 7.7 | 0.02 | 2148900 | YES | 980380 | YES | 1472100 | YES | P50544 |
| **SNVAREAAL** | SNVAREAAL | 9 | 201.8 | 0.5 | 39291.7244 | NO | | YES | 261870 | YES | Q99JY0 |
| **SNVDFLLRL** | SNVDFLLRL | 9 | 32.9 | 0.1 | | YES | 2292800 | YES | 25806 | YES | Q99MR8 |
| **SNVKHVINF** | SNVKHVINF | 9 | 436.4 | 1 | 9959850.32 | YES | 9310000 | YES | 1473600 | YES | Q62167 |
| **SNVKYVML** | SNVKYVM(+15.99)L | 8 | 16.8 | 0.05 | 5565970.56 | NO | 9099000 | YES | 22452000 | YES | Q61115 |
| **SNVKYVML** | SNVKYVML | 8 | 16.8 | 0.05 | 697450.507 | NO | 3204700 | YES | | YES | Q61115 |
| **SNVLYQHNL** | SNVLYQHNL | 9 | 35.7 | 0.125 | 65885 | YES | 4445500 | YES | 1143800 | YES | P42227 |
| **SNYDHAYL** | SNYDHAYL | 8 | 5.3 | 0.015 | 227293.686 | NO | 1753300 | YES | | YES | Q9QXN0 |
| **SNYERLESL** | SNYERLESL | 9 | 64.4 | 0.2 | 304930 | YES | 17591000 | YES | 869350 | YES | P43883 |
| **SNYHFYSSI** | SNYHFYSSI | 9 | 3.9 | 0.01 | 3121900 | YES | 163830000 | YES | 4199100 | YES | Q60795 |
| **SNYLFTKL** | SNYLFTKL | 8 | 2.4 | 0.01 | 556550000 | YES | 3308400000 | YES | 4708700000 | YES | P97481 |
| **SNYLHRVV** | SNYLHRVV | 8 | 57.5 | 0.175 | 119980 | YES | 2430200 | YES | 5207700 | YES | Q78JE5 |
| **SNYLYREV** | SNYLYREV | 8 | 5.5 | 0.015 | 745050.871 | YES | | YES | 1470500 | YES | Q9EPL0 |
| **SNYNFEKPF** | SNYNFEKPF | 9 | 33.6 | 0.1 | 191632.366 | NO | 1135700 | YES | | YES | P62827 |
| **SNYQHITNF** | SNYQHITNF | 9 | 98.7 | 0.3 | 913490 | YES | 20981000 | YES | | YES | O54774 |
| **SNYQMHLL** | SNYQMHLL | 8 | 20.2 | 0.06 | 584721.003 | NO | 2884900 | YES | 699330 | NO | Q3TEA8 |
| **SNYRFEGL** | SNYRFEGL | 8 | 2.9 | 0.01 | 646910 | NO | 8535700 | NO | 2296500 | YES | F8VPZ5 |
| **SNYSYPQV** | SNYSYPQV | 8 | 5.6 | 0.015 | 4328200 | YES | 7480300 | YES | 50569000 | YES | Q61545 |
| **SNYVFVFL** | SNYVFVFL | 8 | 2.6 | 0.01 | 5415600 | YES | | YES | | YES | Q9Z0S9 |

| | | | | | | | | | | | |
|---|---|---|---|---|---|---|---|---|---|---|---|
| **SQFKYALV** | SQFKYALV | 8 | 6.4 | 0.015 | 2603607.48 | NO | 12248000 | NO | 5488900 | YES | Q9QXB9 |
| **SQHNFNNL** | SQHNFNNL | 8 | 28.6 | 0.09 | 147120 | YES | 590980 | YES | | YES | Q69ZB8 |
| **SQLEFRQNL** | SQLEFRQNL | 9 | 23.5 | 0.07 | 30034.9728 | NO | | YES | | YES | Q80UK0 |
| **SQQLYRHI** | SQQLYRHI | 8 | 459.7 | 1 | 463149.205 | YES | 1532800 | YES | 3223600 | YES | Q6ZQ18 |
| **SQQLYRHL** | SQQLYRHL | 8 | 36.4 | 0.125 | 463149.205 | YES | 1532800 | YES | 3223600 | YES | Q8BM75 |
| **SQQTYYRV** | SQQTYYRV | 8 | 234.3 | 0.6 | | YES | 564190 | YES | | YES | P48410 |
| **SQQYYHSL** | SQQYYHSL | 8 | 180 | 0.5 | 30400 | NO | 2005100 | YES | 1197000 | YES | Q69Z38 |
| **SQYLFPKL** | SQYLFPKL | 8 | 4.5 | 0.01 | 1910519.69 | YES | 13704000 | YES | 4180300 | YES | P35396 |
| **SQYQRFTYL** | SQYQRFTYL | 9 | 5.6 | 0.015 | 260508.768 | NO | 1743000 | YES | 277170 | YES | O88942 |
| **SQYRFEHL** | SQYRFEHL | 8 | 3.8 | 0.01 | 1241855.33 | NO | 12725000 | YES | 8898200 | NO | Q3UJK4 |
| **SQYVFTEM** | SQYVFTEM(+15.99) | 8 | 7.8 | 0.02 | 2131600 | NO | 21620000 | NO | 96687000 | NO | Q5SYH2 |
| **SRIVFRHL** | SRIVFRHL | 8 | 44.5 | 0.15 | 2009498.84 | YES | 7202800 | YES | 9002700 | NO | Q9CR88 |
| **SRYQFRNL** | SRYQFRNL | 8 | 6.2 | 0.015 | 1914687.64 | YES | 3104300 | YES | 264880 | NO | Q8R035 |
| **SSATTFRL** | SSATTFRL | 8 | 309.9 | 0.8 | 143290 | YES | 1611600 | NO | | YES | P01027 |
| **SSFAHAQV** | SSFAHAQV | 8 | 7.1 | 0.02 | 419132.502 | YES | | YES | | YES | Q9EQQ9 |
| **SSFEPRSL** | SSFEPRSL | 8 | 100.8 | 0.3 | 959018.792 | YES | | YES | | YES | Q99MJ9 |
| **SSFSHYSGL** | SSFSHYSGL | 9 | 2.9 | 0.01 | | YES | | YES | | YES | Q9CY58 |
| **SSFSRVTNF** | SSFSRVTNF | 9 | 66.3 | 0.2 | 249169.598 | NO | | YES | 201360 | NO | Q8BYH7 |
| **SSFSSPHM** | SSFSSPHM(+15.99) | 8 | 88.2 | 0.25 | 507360.673 | NO | 487160 | YES | 12962000 | NO | O88291 |
| **SSFVFSTV** | SSFVFSTV | 8 | 2.9 | 0.01 | 12189000 | YES | 35991000 | YES | | YES | Q01237 |
| **SSHSFPQL** | SSHSFPQL | 8 | 8.5 | 0.025 | 6365672.26 | YES | | YES | 39601000 | YES | Q8K371 |
| **SSHSFVNV** | SSHSFVNV | 8 | 8 | 0.025 | 229360 | YES | 3747400 | YES | 5198900 | YES | E9Q5F9 |
| **SSIFFREL** | SSIFFREL | 8 | 6.5 | 0.02 | 29841766.7 | YES | 21128000 | YES | | YES | P45448 |
| **SSINFLTRV** | SSINFLTRV | 9 | 10 | 0.03 | 506425.805 | NO | 2470200 | YES | 4395200 | YES | Q8K409 |
| **SSIRQPSL** | SSIRQPSL | 8 | 238.9 | 0.6 | | YES | | YES | 1930700 | YES | Q9R190 |
| **SSIRYFEI** | SSIRYFEI | 8 | 20.7 | 0.06 | 642760.727 | NO | | YES | 1844300 | YES | Q9WUM4:Q9WUM3:O89053 |
| **SSISHSVL** | SSISHSVL | 8 | 77.1 | 0.25 | | YES | 483270 | YES | | YES | Q99K28 |
| **SSIVFAEL** | SSIVFAEL | 8 | 2.7 | 0.01 | 1979400 | YES | 2774200 | YES | | YES | Q9Z2R9 |
| **SSLHFSFL** | SSLHFSFL | 8 | 4.4 | 0.01 | | YES | | YES | | YES | Q80TN4 |

| | | | | | | | | | | | |
|---|---|---|---|---|---|---|---|---|---|---|---|
| **SSLHPMGGL** | SSLHPM(+15.99)GGL | 9 | 369 | 0.9 | 1168200 | NO | 3524900 | YES | 5266600 | NO | P28659 |
| **SSLHPMGGL** | SSLHPMGGL | 9 | 369 | 0.9 | 70476 | NO | 4683000 | YES | 484870 | NO | P28659 |
| **SSLLFVKL** | SSLLFVKL | 8 | 5.6 | 0.015 | 714784.518 | YES | 28252000 | YES | 25985000 | YES | Q80ZV0 |
| **SSLPKRLAL** | SSLPKRLAL | 9 | 287.6 | 0.7 | 87791.899 | NO | 1396500 | NO | 469880 | NO | Q8R1F0 |
| **SSLSFNTRL** | SSLSFNTRL | 9 | 10.9 | 0.03 | 210060 | YES | 5006300 | YES | | YES | Q5SSN7 |
| **SSLVKVNL** | SSLVKVNL | 8 | 85.4 | 0.25 | 356640.469 | NO | 1547400 | YES | 578280 | YES | Q9QYC0 |
| **SSLYFRDL** | SSLYFRDL | 8 | 7 | 0.02 | 1679139.85 | YES | | YES | 1659100 | YES | Q9DCM7 |
| **SSMAYPNL** | SSM(+15.99)AYPNL | 8 | 3.8 | 0.01 | 665390 | NO | 586060 | NO | 2868500 | NO | Q61249 |
| **SSPAFSKV** | SSPAFSKV | 8 | 26.3 | 0.08 | 396440 | YES | 193390 | YES | 422310 | YES | Q99M02 |
| **SSPEYEAL** | SSPEYEAL | 8 | 20.8 | 0.06 | | YES | | YES | | YES | Q8R242 |
| **SSPHYTTL** | SSPHYTTL | 8 | 12.2 | 0.04 | 18334000 | YES | 11403000 | YES | 33812000 | YES | P27046 |
| **SSPKFSEI** | SSPKFSEI | 8 | 53.5 | 0.175 | 11409000 | NO | 151370000 | NO | 51956000 | NO | Q91VM3 |
| **SSPKFSEL** | SSPKFSEL | 8 | 9.3 | 0.025 | 11409000 | NO | 151370000 | NO | 51956000 | NO | A2AGT5 |
| **SSPKYDYL** | SSPKYDYL | 8 | 27.6 | 0.08 | 2941133.79 | YES | | YES | 4066700 | YES | Q7TQG1 |
| **SSPVFKAM** | SSPVFKAM | 8 | 22.9 | 0.07 | 53924 | NO | 1125000 | YES | 225100 | NO | Q9Z2X8 |
| **SSPVFKAM** | SSPVFKAM(+15.99) | 8 | 22.9 | 0.07 | 559910 | NO | 500250 | YES | 1487500 | NO | Q9Z2X8 |
| **SSPVFKAMF** | SSPVFKAM(+15.99)F | 9 | 37.9 | 0.125 | 907940 | NO | 7465000 | YES | 10939000 | NO | Q9Z2X8 |
| **SSPVFKAMF** | SSPVFKAMF | 9 | 37.9 | 0.125 | 202210 | NO | 13474000 | YES | 1651300 | NO | Q9Z2X8 |
| **SSPVYIDL** | SSPVYIDL | 8 | 15.2 | 0.05 | | YES | 3301900 | YES | | YES | P13439 |
| **SSTHFATL** | SSTHFATL | 8 | 7.4 | 0.02 | 147550 | NO | 1736300 | YES | 1616300 | YES | Q6S5J6 |
| **SSTYFHQL** | SSTYFHQL | 8 | 15.4 | 0.05 | 337580 | YES | 4094100 | YES | 560340 | YES | Q8BN78 |
| **SSVEYIHRI** | SSVEYIHRI | 9 | 182.6 | 0.5 | 294440.583 | NO | 183490 | YES | 53238 | NO | Q8K301 |
| **SSVEYNHRL** | SSVEYNHRL | 9 | 119.5 | 0.4 | 25896 | NO | | YES | 137400 | NO | P49935 |
| **SSVKYSKI** | SSVKYSKI | 8 | 63.6 | 0.2 | 112300 | NO | 87328 | NO | 456110 | NO | Q64511 |
| **SSVLYSRV** | SSVLYSRV | 8 | 7 | 0.02 | 1623700 | YES | 4247300 | YES | | YES | Q8CGC7 |
| **SSVRFSYM** | SSVRFSYM | 8 | 5.1 | 0.015 | 378738.413 | NO | | YES | 1179300 | YES | Q8R5K4 |
| **SSVRPVNL** | SSVRPVNL | 8 | 95.8 | 0.3 | 62326 | YES | | YES | 179580 | YES | Q8CDG3 |
| **SSVSFKERL** | SSVSFKERL | 9 | 28.3 | 0.09 | 165078.218 | NO | | YES | 181650 | NO | Q8BTJ4 |
| **SSVYFRSV** | SSVYFRSV | 8 | 14.4 | 0.04 | 1068900 | YES | 16084000 | YES | 3159900 | YES | Q9D6Y4 |

| | | | | | | | | | | | |
|---|---|---|---|---|---|---|---|---|---|---|---|
| **SSYAYTKV** | SSYAYTKV | 8 | 3.7 | 0.01 | 76713.7128 | NO | | YES | 2014600 | YES | Q8BG87 |
| **SSYFFGKV** | SSYFFGKV | 8 | 4 | 0.01 | 1481700 | YES | 6839600 | YES | 754810 | YES | Q7TMS5 |
| **SSYKFNHL** | SSYKFNHL | 8 | 2.3 | 0.01 | 2775584.24 | YES | 18568000 | YES | 17621000 | YES | Q8K2I1 |
| **SSYLHSLL** | SSYLHSLL | 8 | 3.2 | 0.01 | 64357.8736 | NO | 1735100 | YES | 2890500 | NO | Q8BWZ3 |
| **SSYNTFRL** | SSYNTFRL | 8 | 8.6 | 0.025 | | YES | | YES | | YES | Q9EPZ6 |
| **SSYNYIRV** | SSYNYIRV | 8 | 2.7 | 0.01 | 551500 | YES | 47041000 | YES | 5415000 | YES | Q5SYD0 |
| **SSYNYRVV** | SSYNYRVV | 8 | 5.4 | 0.015 | 638710 | YES | 8930000 | YES | | YES | P11881 |
| **SSYQHTSV** | SSYQHTSV | 8 | 11.8 | 0.04 | 565100 | YES | 93776 | YES | 429620 | YES | Q9CRT8 |
| **SSYRFVQNV** | SSYRFVQNV | 9 | 3.1 | 0.01 | 694256.374 | NO | 2285200 | NO | 3669800 | NO | P42128 |
| **SSYSFRHL** | SSYSFRHL | 8 | 2.1 | 0.01 | 11488749.1 | YES | 7244900 | YES | 5886900 | YES | O35448 |
| **SSYSFRHLL** | SSYSFRHLL | 9 | 4.5 | 0.01 | 204062.216 | NO | 1069900 | YES | 404220 | NO | O35448 |
| **SSYTFPKM** | SSYTFPKM | 8 | 3.4 | 0.01 | 1337600 | YES | 43571000 | YES | 6079500 | YES | Q9WUR2 |
| **SSYTFPKM** | SSYTFPKM(+15.99) | 8 | 3.4 | 0.01 | 17705000 | YES | 39878000 | YES | 81483000 | YES | Q9WUR2 |
| **SSYTFPKMM** | SSYTFPKM(+15.99)M(+15.99) | 9 | 13.7 | 0.04 | 675170 | YES | 2123600 | YES | 4546800 | YES | Q9WUR2 |
| **SSYTFPKMM** | SSYTFPKMM | 9 | 13.7 | 0.04 | 4857808.02 | YES | 4715300 | YES | 58298 | YES | Q9WUR2 |
| **SSYTFPKMM** | SSYTFPKMM(+15.99) | 9 | 13.7 | 0.04 | 4362717.49 | YES | 1066600 | YES | 951970 | YES | Q9WUR2 |
| **SSYVHSNL** | SSYVHSNL | 8 | 3.2 | 0.01 | 995460 | YES | 1668900 | YES | 13793000 | YES | Q6P4T1 |
| **SSYVVKKV** | SSYVVKKV | 8 | 193.7 | 0.5 | 21685.8222 | NO | 51785 | YES | 154110 | YES | Q4U2R1 |
| **STFEFHSI** | STFEFHSI | 8 | 16.8 | 0.05 | | YES | | YES | 42125000 | YES | O70481 |
| **STFFYPKL** | STFFYPKL | 8 | 6.9 | 0.02 | 673611.372 | YES | | YES | 929860 | YES | Q91ZX6 |
| **STFSFTKV** | STFSFTKV | 8 | 7.7 | 0.02 | 171330 | YES | 30304000 | YES | 9882700 | YES | Q3TZZ7 |
| **STFSHKTV** | STFSHKTV | 8 | 278.5 | 0.7 | 32656.4986 | NO | 58292 | NO | 54070 | YES | Q78IS1 |
| **STFTFADL** | STFTFADL | 8 | 2.7 | 0.01 | 961620 | YES | 1780300 | YES | | YES | Q9ERU9 |
| **STFVYNSM** | STFVYNSM | 8 | 7 | 0.02 | | YES | 93050000 | YES | 1482100 | YES | Q61107 |
| **STFVYNSM** | STFVYNSM(+15.99) | 8 | 7 | 0.02 | 3258600 | YES | 114800000 | YES | 63565000 | YES | Q61107 |
| **STIEFKNM** | STIEFKNM(+15.99) | 8 | 20.7 | 0.06 | 851710 | NO | 2471900 | YES | 4413300 | YES | Q6NXI6 |
| **STIVYYKL** | STIVYYKL | 8 | 8.2 | 0.025 | 116280 | YES | 5397200 | YES | 197140 | YES | Q8R3W5 |
| **STLIYRNM** | STLIYRNM | 8 | 8.3 | 0.025 | | YES | 4143400 | NO | 1361200 | NO | Q9D8N6 |
| **STLIYRNM** | STLIYRNM(+15.99) | 8 | 8.3 | 0.025 | 1349200 | YES | 2357700 | NO | 8092200 | NO | Q9D8N6 |

| | | | | | | | | | | | |
|---|---|---|---|---|---|---|---|---|---|---|---|
| **STLSYRSL** | STLSYRSL | 8 | 8.3 | 0.025 | 1980206.31 | YES | | YES | | YES | Q80W22 |
| **STLTYSRM** | STLTYSRM | 8 | 8.3 | 0.025 | | YES | 35731000 | YES | 2924300 | YES | Q9WVC6 |
| **STLTYSRM** | STLTYSRM(+15.99) | 8 | 8.3 | 0.025 | 857975.139 | YES | 20508000 | YES | 55938000 | YES | Q9WVC6 |
| **STPEFYQV** | STPEFYQV | 8 | 40.3 | 0.125 | 1347300 | NO | 26781000 | NO | 10381000 | YES | Q99JY9 |
| **STPKYQRL** | STPKYQRL | 8 | 13.3 | 0.04 | | YES | | YES | 824260 | YES | Q6PGC1 |
| **STRLFAVL** | STRLFAVL | 8 | 9.8 | 0.03 | 5883825.69 | YES | 27109000 | YES | | YES | P46978 |
| **STTVFHSL** | STTVFHSL | 8 | 44.2 | 0.15 | 5173325.99 | YES | | YES | | YES | P19096 |
| **STVEFTNL** | STVEFTNL | 8 | 5.4 | 0.015 | | YES | 9748800 | YES | | YES | O54692 |
| **STVLLQRL** | STVLLQRL | 8 | 164.7 | 0.5 | 488624.359 | YES | | YES | 455990 | YES | O88907 |
| **STVQFHIL** | STVQFHIL | 8 | 68.1 | 0.2 | 157710 | YES | | YES | 284380 | NO | Q9CR50 |
| **STYEYVRFI** | STYEYVRFI | 9 | 31.4 | 0.09 | 54463.8536 | NO | 2380100 | YES | 387580 | YES | O08785 |
| **STYFPHTAI** | STYFPHTAI | 9 | 111.7 | 0.3 | 3909306.5 | YES | 6559500 | YES | | YES | P70255 |
| **STYIKFVNL** | STYIKFVNL | 9 | 4.1 | 0.01 | 120520.071 | NO | 824830 | YES | 148640 | NO | P17427 |
| **STYKFFEV** | STYKFFEV | 8 | 6.5 | 0.02 | 58662969.2 | YES | 451740000 | YES | 189500000 | YES | Q9CZM2 |
| **STYSHSAL** | STYSHSAL | 8 | 6.1 | 0.015 | 337919.737 | YES | 252820 | YES | 9890900 | YES | Q61687 |
| **STYSVAKM** | STYSVAKM(+15.99) | 8 | 33.1 | 0.1 | 375390 | NO | 369240 | YES | 16475000 | YES | O09159 |
| **SVFAFGENKM** | SVFAFGENKM | 10 | 377.8 | 0.9 | 782874.073 | NO | 10787000 | YES | 1785500 | NO | Q8BK67 |
| **SVFAFGENKM** | SVFAFGENKM(+15.99) | 10 | 377.8 | 0.9 | 73913 | NO | 5453600 | YES | 11256000 | NO | Q8BK67 |
| **SVIKFENL** | SVIKFENL | 8 | 7.1 | 0.02 | 30541000 | YES | 71650000 | YES | 103480000 | YES | Q9CPV7 |
| **SVISVIHL** | SVISVIHL | 8 | 187.5 | 0.5 | 41921.0408 | NO | | YES | | YES | Q8VDP6 |
| **SVITVKNL** | SVITVKNL | 8 | 319.1 | 0.8 | 131550 | YES | | YES | | YES | Q8BGQ4 |
| **SVLLFMQL** | SVLLFMQL | 8 | 5 | 0.01 | | YES | | YES | | YES | Q9Z0E8 |
| **SVLQFLGL** | SVLQFLGL | 8 | 8.6 | 0.025 | 3989200 | YES | 2064500 | YES | | YES | Q9CQC9 |
| **SVNIFRTL** | SVNIFRTL | 8 | 62.6 | 0.2 | 388176.684 | NO | 6813600 | YES | | YES | Q6PD28 |
| **SVPKFKHL** | SVPKFKHL | 8 | 19.3 | 0.06 | 1750267.72 | YES | 190830 | YES | 361930 | YES | Q9QZW0 |
| **SVRLAALL** | SVRLAALL | 8 | 270.7 | 0.7 | | YES | | YES | 391860 | YES | Q91ZU9 |
| **SVVAFHNL** | SVVAFHNL | 8 | 8.7 | 0.025 | 6466700 | YES | 60109000 | YES | 29296000 | YES | Q71B07 |
| **SVVALHNL** | SVVALHNL | 8 | 299 | 0.7 | 17220.6587 | NO | | YES | | YES | P26516 |
| **SVVAYNNL** | SVVAYNNL | 8 | 11 | 0.03 | 1735800 | NO | 3440200 | NO | 2624900 | NO | Q99LM9 |

| | | | | | | | | | | | |
|---|---|---|---|---|---|---|---|---|---|---|---|
| **SVVDYCNRL** | SVVDYC(+119.00)NRL | 9 | 67.6 | 0.2 | 298495.573 | NO | 2642800 | NO | 426500 | NO | Q8BU14 |
| **SVVEYSRL** | SVVEYSRL | 8 | 8.7 | 0.025 | 342399.313 | NO | | YES | | YES | G5E8F4 |
| **SVVRYVQL** | SVVRYVQL | 8 | 11 | 0.03 | 851860 | YES | 30668000 | YES | 15728 | YES | O35900 |
| **SVYLVRQL** | SVYLVRQL | 8 | 16.2 | 0.05 | 59777 | YES | | YES | | YES | Q8C5D8:O54714 |
| **SVYQPAQL** | SVYQPAQL | 8 | 13.6 | 0.04 | 2847300 | YES | 10213000 | YES | | YES | B2RVL6 |
| **SVYTHSYL** | SVYTHSYL | 8 | 4.6 | 0.01 | 918450 | YES | 17020000 | YES | 14553000 | YES | Q8K0L2:Q3TZX8 |
| **SVYVYKVL** | SVYVYKVL | 8 | 6.7 | 0.02 | 16062000 | YES | 910460000 | YES | 855000000 | YES | Q6ZWY9:P10853:Q64525:Q8CGP1 |
| **SYFKGASL** | SYFKGASL | 8 | 270.1 | 0.7 | 173215.466 | NO | 1756100 | YES | 1300000 | YES | Q8C129 |
| **TAFGYKGL** | TAFGYKGL | 8 | 23.5 | 0.07 | 378411.209 | NO | 581840 | YES | | YES | Q8BY71 |
| **TAFKFKAL** | TAFKFKAL | 8 | 6.5 | 0.02 | | YES | 9336800 | YES | 4782800 | YES | Q04690 |
| **TAFRFSEL** | TAFRFSEL | 8 | 3.3 | 0.01 | 14826000 | YES | 48645000 | NO | 33178000 | NO | A3KGB4 |
| **TAHAFVNV** | TAHAFVNV | 8 | 41.4 | 0.125 | 108250 | YES | 362360 | YES | 94368 | YES | P19096 |
| **TAILFQRI** | TAILFQRI | 8 | 33.8 | 0.1 | 1480800 | YES | | YES | | YES | E9Q414 |
| **TALAFRTL** | TALAFRTL | 8 | 13.3 | 0.04 | 675481.108 | NO | 7635900 | YES | 4884700 | YES | Q8CIP4 |
| **TALDHYSEL** | TALDHYSEL | 9 | 36.8 | 0.125 | 592862.145 | NO | 3513200 | YES | 1507400 | YES | Q6ZWQ0 |
| **TALRFLEL** | TALRFLEL | 8 | 21.7 | 0.07 | | YES | 28940000 | YES | 2125400 | YES | Q8VD65 |
| **TALRYIQSL** | TALRYIQSL | 9 | 32.6 | 0.1 | 115522.422 | NO | 1123600 | YES | 627840 | YES | O08789 |
| **TAPHYQLL** | TAPHYQLL | 8 | 72 | 0.2 | 384150 | YES | 6562000 | YES | 1033300 | YES | Q9D5R2 |
| **TAPQYYRL** | TAPQYYRL | 8 | 11.9 | 0.04 | 5644400 | YES | 28696000 | YES | 43130000 | YES | Q6PJN8 |
| **TAYAFHFL** | TAYAFHFL | 8 | 4.5 | 0.01 | | YES | 39382000 | YES | 2128800 | YES | Q9R0H0 |
| **TAYEFAKL** | TAYEFAKL | 8 | 2.4 | 0.01 | 15600000 | YES | 22428000 | YES | 27740000 | YES | Q9EQ06 |
| **TAYHFSLV** | TAYHFSLV | 8 | 6.3 | 0.015 | 821910 | YES | 6521300 | YES | | YES | Q8R0L1 |
| **TAYLFSRF** | TAYLFSRF | 8 | 8.1 | 0.025 | 1188600 | YES | | YES | | YES | Q9CRT8 |
| **TEYVFTHL** | TEYVFTHL | 8 | 15.1 | 0.05 | 1134900 | YES | 128070000 | YES | 25324000 | YES | Q6NWW9 |
| **TGATYPHL** | TGATYPHL | 8 | 111.7 | 0.3 | 135890 | YES | 379150 | YES | | YES | Q9DBA6 |
| **TGIKFVVL** | TGIKFVVL | 8 | 107.1 | 0.3 | 2039531.48 | YES | | YES | | YES | Q9ES56 |
| **TGPKYIHL** | TGPKYIHL | 8 | 26.7 | 0.08 | 5749438.41 | YES | 11527000 | YES | 130220000 | YES | Q60805 |
| **TGYNFQRV** | TGYNFQRV | 8 | 6.8 | 0.02 | 1416300 | YES | | YES | 119030 | YES | Q922V4 |
| **THFQPAQL** | THFQPAQL | 8 | 345.9 | 0.8 | 363605.236 | NO | 1538900 | YES | 66558 | NO | P54822 |

| | | | | | | | | | | | |
|---|---|---|---|---|---|---|---|---|---|---|---|
| **THYSFLATL** | THYSFLATL | 9 | 7.1 | 0.02 | 294290 | NO | 1102900 | YES | | YES | Q5U419 |
| **TIIIFHSL** | TIIIFHSL | 8 | 18.4 | 0.06 | 4184313.51 | YES | | YES | | YES | O08575;P97480 |
| **TIILFTKV** | TIILFTKV | 8 | 58.7 | 0.175 | 1239300 | YES | | YES | | YES | Q8K2V6 |
| **TILEFSQNM** | TILEFSQNM | 9 | 34.2 | 0.1 | 1127918.28 | NO | 20337000 | YES | 3643900 | YES | Q6P5F9 |
| **TILEFSQNM** | TILEFSQNM(+15.99) | 9 | 34.2 | 0.1 | 735920 | NO | 17792000 | YES | 59154000 | YES | Q6P5F9 |
| **TITSFPRL** | TITSFPRL | 8 | 32.3 | 0.1 | | YES | | YES | | YES | P97494 |
| **TIYERFVLV** | TIYERFVLV | 9 | 24.9 | 0.08 | 40382.4038 | NO | | YES | | YES | P63154 |
| **TIYRFLKL** | TIYRFLKL | 8 | 3.7 | 0.01 | 1936700 | YES | 10735000 | YES | | YES | Q8CEF1 |
| **TNIDFAFKRL** | TNIDFAFKRL | 10 | 136.9 | 0.4 | 186946.34 | NO | 9278900 | YES | 4783900 | NO | P29416 |
| **TNINFPNL** | TNINFPNL | 8 | 6.2 | 0.015 | 496414.926 | NO | 5379500 | YES | 4129800 | YES | Q8BRH4 |
| **TNISFTNM** | TNISFTNM(+15.99) | 8 | 10.5 | 0.03 | 839680 | NO | 7431100 | NO | 16721000 | NO | Q9CYY7 |
| **TNLIYQQV** | TNLIYQQV | 8 | 63.8 | 0.2 | 1322300 | YES | 16126000 | YES | 7374200 | YES | P97479 |
| **TNLQRVSYL** | TNLQRVSYL | 9 | 162.2 | 0.5 | 189513.332 | NO | 377600 | YES | 262690 | YES | Q810A7 |
| **TNLRYLAL** | TNLRYLAL | 8 | 9.1 | 0.025 | 260120 | YES | 15581000 | YES | 28988000 | YES | P17427:P17426 |
| **TNLVPYPRI** | TNLVPYPRI | 9 | 121 | 0.4 | 785110 | YES | 50228000 | YES | 24822000 | YES | P68373:P05213:P68368:Q9JJZ2 |
| **TNLVYPAL** | TNLVYPAL | 8 | 10.4 | 0.03 | | YES | | YES | 6421600 | YES | Q8BLR9 |
| **TNPSFDGRL** | TNPSFDGRL | 9 | 88.6 | 0.25 | 4145700 | YES | 16339000 | YES | | YES | Q6PA06 |
| **TNQDFIQRL** | TNQDFIQRL | 9 | 149.4 | 0.4 | 1766900 | YES | 231070000 | YES | 4415800 | YES | Q80TM9 |
| **TNVEYAHL** | TNVEYAHL | 8 | 6.6 | 0.02 | | YES | 310790 | YES | 402950 | YES | Q8K370 |
| **TNVKFLAI** | TNVKFLAI | 8 | 284.6 | 0.7 | | YES | | YES | 2471900 | YES | Q02248 |
| **TNVLFNHL** | TNVLFNHL | 8 | 11.1 | 0.04 | 782010 | YES | 68713000 | YES | 14588000 | YES | Q9D706 |
| **TNVQYSNL** | TNVQYSNL | 8 | 9.8 | 0.03 | 1048000 | NO | | YES | 3082200 | YES | Q45VK7 |
| **TNVTFSKV** | TNVTFSKV | 8 | 111.7 | 0.3 | 1067200 | YES | 1094100 | YES | | YES | Q6P5B0 |
| **TNYIFDSL** | TNYIFDSL | 8 | 6.3 | 0.015 | 1429500 | YES | 12847000 | YES | 2465400 | YES | Q9CXF4 |
| **TNYNFQYI** | TNYNFQYI | 8 | 7 | 0.02 | 2145300 | YES | 28333000 | YES | 21076000 | YES | Q8K2C8 |
| **TNYNFQYISL** | TNYNFQYISL | 10 | 9.9 | 0.03 | 423430 | YES | 3189300 | YES | | YES | Q8K2C8 |
| **TNYTFENV** | TNYTFENV | 8 | 5.6 | 0.015 | 176787.441 | NO | 2045600 | YES | 1362000 | YES | Q61493 |
| **TQFLYPKV** | TQFLYPKV | 8 | 68.3 | 0.2 | 454969.11 | YES | 6723200 | YES | | YES | P15066 |
| **TQQLYPSL** | TQQLYPSL | 8 | 248.3 | 0.6 | | YES | 12020000 | YES | 9969800 | YES | P69566 |

| | | | | | | | | | | | |
|---|---|---|---|---|---|---|---|---|---|---|---|
| **TQYIFNNM** | TQYIFNNM | 8 | 6.4 | 0.015 | 86678 | YES | 5618300 | NO | 8366.5 | NO | P27046 |
| **TQYIFNNM** | TQYIFNNM(+15.99) | 8 | 6.4 | 0.015 | 774970 | YES | 4094200 | NO | 7762500 | NO | P27046 |
| **TQYSFYQQL** | TQYSFYQQL | 9 | 3.8 | 0.01 | 705046.309 | NO | | YES | 1428100 | NO | Q9Z329 |
| **TSFMFQRV** | TSFMFQRV | 8 | 3.3 | 0.01 | 4558650.25 | YES | 9173400 | YES | 552770 | YES | P97429 |
| **TSFRYSSL** | TSFRYSSL | 8 | 2.3 | 0.01 | 805630 | NO | 13743000 | NO | 21160000 | NO | Q8BX90 |
| **TSFTFRKV** | TSFTFRKV | 8 | 7.3 | 0.02 | | YES | 1292600 | YES | 855270 | YES | Q80Y20 |
| **TSIAFKNI** | TSIAFKNI | 8 | 30.6 | 0.09 | 251210 | YES | 5640100 | YES | 5274600 | YES | Q8VHJ5:Q03141 |
| **TSIQFNLRNL** | TSIQFNLRNL | 10 | 81.8 | 0.25 | 131808.602 | NO | 4166200 | YES | | YES | Q8BJ56 |
| **TSLKYLEM** | TSLKYLEM | 8 | 70.9 | 0.2 | 717472.264 | NO | 1380700 | YES | | YES | Q9CWG9 |
| **TSPEYQKL** | TSPEYQKL | 8 | 40.9 | 0.125 | 26719000 | YES | 1862700 | YES | 7601600 | YES | Q920R0 |
| **TSPLFLHF** | TSPLFLHF | 8 | 110.4 | 0.3 | | YES | 21702000 | YES | 6210600 | YES | Q5SSZ5 |
| **TSVRFTQL** | TSVRFTQL | 8 | 5.3 | 0.015 | 8610400 | YES | 134680000 | YES | 290030000 | YES | Q9D6T0 |
| **TSVVFNKL** | TSVVFNKL | 8 | 16.7 | 0.05 | 282890 | YES | 13397000 | YES | 9304700 | YES | Q791N7 |
| **TSYIFVSV** | TSYIFVSV | 8 | 4.2 | 0.01 | 110960 | NO | 3372200 | NO | | YES | P42337 |
| **TSYRFLAL** | TSYRFLAL | 8 | 2.1 | 0.01 | 4515300 | NO | 48499000 | NO | 730320 | YES | Q8QZX2 |
| **TSYSYIRL** | TSYSYIRL | 8 | 2.3 | 0.01 | | YES | 1072100 | YES | | YES | F8VPZ5 |
| **TTFEHAHNM** | TTFEHAHNM | 9 | 268.3 | 0.7 | 229989.222 | YES | 571140 | YES | | YES | P58252 |
| **TTFEHAHNM** | TTFEHAHNM(+15.99) | 9 | 268.3 | 0.7 | 107010 | YES | 334450 | YES | 2947900 | YES | P58252 |
| **TTLIFQKL** | TTLIFQKL | 8 | 18.8 | 0.06 | 420370 | YES | 11678000 | YES | 77775000 | YES | P15307 |
| **TTLLYKPI** | TTLLYKPI | 8 | 89.8 | 0.25 | 44701 | YES | | YES | 402240 | NO | Q9R078 |
| **TTVAFTQV** | TTVAFTQV | 8 | 95.5 | 0.3 | 5370000 | YES | 13987000 | YES | 1149600 | YES | P12970 |
| **TTYKYEMI** | TTYKYEM(+15.99)I | 8 | 28.6 | 0.09 | 1389700 | YES | 34702000 | YES | 23347000 | NO | Q8QZY1 |
| **TTYKYFAL** | TTYKYFAL | 8 | 2.7 | 0.01 | 3912300 | YES | | YES | 115940000 | YES | Q5BLK4 |
| **TTYVHKGL** | TTYVHKGL | 8 | 26.3 | 0.08 | 136760 | YES | | YES | | YES | Q9CZW5 |
| **TTYVHKGLL** | TTYVHKGLL | 9 | 41.6 | 0.125 | 105277.827 | YES | 126810 | NO | 126130 | NO | Q9CZW5 |
| **TVQSFHHL** | TVQSFHHL | 8 | 41.8 | 0.125 | 68927.0411 | NO | | YES | 84697 | NO | Q9R1J0 |
| **TVRFFNSV** | TVRFFNSV | 8 | 298.8 | 0.7 | | YES | | YES | | YES | Q9ERV1 |
| **TVTEFKQL** | TVTEFKQL | 8 | 368 | 0.9 | 89607 | NO | | YES | | YES | Q8BY87 |
| **VAALFKNL** | VAALFKNL | 8 | 9.7 | 0.03 | 348210 | YES | 3538100 | YES | 1208000 | YES | Q5U464 |

| | | | | | | | | | | | |
|---|---|---|---|---|---|---|---|---|---|---|---|
| **VAASFKGL** | VAASFKGL | 8 | 36.5 | 0.125 | 1284313.92 | YES | 2843300 | YES | 605700 | YES | Q6P5B0 |
| **VADKFSEL** | VADKFSEL | 8 | 100.9 | 0.3 | 1715755.51 | NO | 16091000 | YES | 7016600 | YES | Q7TNP2 |
| **VADKFTEL** | VADKFTEL | 8 | 140.9 | 0.4 | 2136700 | YES | 90223000 | YES | 58585000 | YES | Q76MZ3 |
| **VAFAFKKL** | VAFAFKKL | 8 | 4.6 | 0.01 | 816030 | YES | 4007500 | YES | 5848600 | YES | Q8CFA1 |
| **VAFAYKNV** | VAFAYKNV | 8 | 4.4 | 0.01 | 3911700 | NO | 59748000 | NO | 30798000 | NO | Q9DCF9 |
| **VAFDFTKV** | VAFDFTKV | 8 | 10.6 | 0.03 | 6244100 | YES | 304680000 | YES | 323470000 | YES | Q7M6Y3 |
| **VAFIFNQKF** | VAFIFNQKF | 9 | 53.7 | 0.175 | 85914.3724 | NO | 2216400 | YES | | YES | Q8CJ19:Q8BML1 |
| **VAFNHQNL** | VAFNHQNL | 8 | 5.5 | 0.015 | 1351390.7 | YES | 3213200 | YES | 923520 | YES | Q8BX90 |
| **VAHTFVIGV** | VAHTFVIGV | 9 | 72.7 | 0.2 | 139060 | YES | 8479300 | YES | 12603000 | YES | P50580 |
| **VAIGFKTKL** | VAIGFKTKL | 9 | 40.9 | 0.125 | 1021109.61 | NO | | YES | | YES | Q64310 |
| **VAIRFDSGL** | VAIRFDSGL | 9 | 9.5 | 0.03 | 1269122.31 | NO | 7503700 | YES | 8532100 | YES | Q9ERU3 |
| **VAITYKEL** | VAITYKEL | 8 | 15.3 | 0.05 | 1452200.63 | YES | 14976000 | YES | | YES | Q03963 |
| **VALDFEQEM** | VALDFEQEM | 9 | 87.5 | 0.25 | 352363.448 | NO | 30764000 | YES | 5250600 | YES | P60710:P63260:Q8BFZ3 |
| **VALDFEQEM** | VALDFEQEM(+15.99) | 9 | 87.5 | 0.25 | 505240 | NO | 30454000 | YES | 99404000 | YES | P60710:P63260:Q8BFZ3 |
| **VALLFRQL** | VALLFRQL | 8 | 3.6 | 0.01 | 2950365.61 | YES | 24914000 | YES | | YES | Q6ZPE2 |
| **VAMVFKTL** | VAMVFKTL | 8 | 8.3 | 0.025 | 273117.801 | YES | 3321900 | YES | 809390 | YES | Q65Z40 |
| **VAPDRFPTL** | VAPDRFPTL | 9 | 24.2 | 0.07 | 2309279.86 | NO | 20794000 | YES | 23681000 | YES | Q9DBS8 |
| **VAPFFKSYI** | VAPFFKSYI | 9 | 52 | 0.15 | 152140 | NO | 11649000 | YES | | YES | Q9JHU4 |
| **VAPHHLFL** | VAPHHLFL | 8 | 242.8 | 0.6 | 228415.528 | NO | 4538500 | NO | 2948700 | YES | B2RQC6 |
| **VAPQYQEL** | VAPQYQEL | 8 | 17.2 | 0.05 | 6267900.64 | YES | | YES | 2697600 | YES | Q9D711 |
| **VAPRYVALL** | VAPRYVALL | 9 | 5.9 | 0.015 | 1492700 | YES | 2346500 | NO | 1665700 | NO | Q8CGC7 |
| **VAPSAVNL** | VAPSAVNL | 8 | 108.9 | 0.3 | | YES | | YES | | YES | Q8BTI8 |
| **VAQKFNHL** | VAQKFNHL | 8 | 10.2 | 0.03 | 106259.438 | NO | 112870 | YES | | YES | Q80ZE4 |
| **VAVIHQSL** | VAVIHQSL | 8 | 98.9 | 0.3 | | YES | 215020 | YES | | YES | Q8K158 |
| **VAYGFRNI** | VAYGFRNI | 8 | 3.8 | 0.01 | 1082200 | YES | | YES | 15263000 | YES | Q7TMW6 |
| **VAYKFPEL** | VAYKFPEL | 8 | 2.6 | 0.01 | 3324936.07 | YES | 40786000 | YES | 114680000 | YES | Q6WKZ8 |
| **VAYKFPELL** | VAYKFPELL | 9 | 9.5 | 0.03 | 3719200 | YES | 16675000 | YES | 110820 | YES | Q6WKZ8 |
| **VAYRHLVGV** | VAYRHLVGV | 9 | 12.3 | 0.04 | 175018.982 | YES | 1768900 | NO | 386090 | NO | Q9D0M3 |

| | | | | | | | | | | | |
|---|---|---|---|---|---|---|---|---|---|---|---|
| **VAYRYEVL** | VAYRYEVL | 8 | 3.4 | 0.01 | 2236515.96 | NO | 2496900 | NO | | YES | Q5U4C9 |
| **VAYSHDGAFL** | VAYSHDGAFL | 10 | 89.7 | 0.25 | 504945.597 | NO | 8629000 | YES | 1187400 | YES | O88342 |
| **VAYWRQAGL** | VAYWRQAGL | 9 | 6.3 | 0.015 | 2112900 | YES | 59981000 | YES | | YES | P56382 |
| **VEYDFHLL** | VEYDFHLL | 8 | 67.5 | 0.2 | 2727243.77 | YES | 13793000 | YES | | YES | P46664 |
| **VFIDKQTNL** | VFIDKQTNL | 9 | 267.2 | 0.7 | 433130 | YES | 7778400 | YES | 5870600 | YES | P28659:Q9Z0H4 |
| **VFRLLPQL** | VFRLLPQL | 8 | 299.9 | 0.7 | | YES | 49991000 | YES | 30905000 | YES | Q9EST5 |
| **VFTEVANL** | VFTEVANL | 8 | 195.2 | 0.5 | 532560 | YES | | YES | | YES | Q62141 |
| **VFVKVINL** | VFVKVINL | 8 | 239.3 | 0.6 | 59712 | NO | 20249000 | YES | 12375000 | YES | Q69Z37 |
| **VFYQVQSL** | VFYQVQSL | 8 | 32.5 | 0.1 | 47380 | NO | | YES | | YES | B1AY13 |
| **VGFDYKERL** | VGFDYKERL | 9 | 37.1 | 0.125 | 16326301.6 | YES | 668150000 | YES | 46848000 | YES | Q60598 |
| **VGFTFPNRL** | VGFTFPNRL | 9 | 7.8 | 0.02 | 2852243.42 | YES | 42465000 | YES | 10415000 | YES | Q9Z0E0 |
| **VGIGFSNL** | VGIGFSNL | 8 | 4.6 | 0.01 | | YES | | YES | | YES | Q9JJT0 |
| **VGITYQHI** | VGITYQHI | 8 | 19.6 | 0.06 | 1683100 | YES | 79695000 | YES | 70998000 | YES | Q9DBZ5 |
| **VGLKFPGL** | VGLKFPGL | 8 | 5.6 | 0.015 | 1436775.31 | YES | 2401200 | YES | | YES | Q62086 |
| **VGLRYEKI** | VGLRYEKI | 8 | 124.7 | 0.4 | | YES | 6228900 | YES | 9261500 | YES | Q9WUN2 |
| **VGLYYINKI** | VGLYYINKI | 9 | 200.2 | 0.5 | 194170 | YES | | YES | 865020 | YES | Q5RL79 |
| **VGMKYRNL** | VGMKYRNL | 8 | 4.5 | 0.01 | | YES | 92171 | YES | | YES | Q91YP2 |
| **VGNEFSHL** | VGNEFSHL | 8 | 15 | 0.05 | 460461.46 | NO | | YES | 502140 | YES | Q99ML9 |
| **VGNNFHNL** | VGNNFHNL | 8 | 18.9 | 0.06 | 100591.801 | NO | 557820 | YES | | YES | Q8K2H6 |
| **VGPKFRGV** | VGPKFRGV | 8 | 43.4 | 0.15 | 70001 | YES | 5180300 | YES | 7039400 | YES | Q9D2V5 |
| **VGPRYTNL** | VGPRYTNL | 8 | 4.6 | 0.01 | 194610000 | YES | 645770000 | NO | 1217400000 | YES | P63085 |
| **VGPRYTQL** | VGPRYTQL | 8 | 6.8 | 0.02 | 8149500 | YES | 88811000 | NO | 47054000 | NO | Q63844 |
| **VGPYTYREL** | VGPYTYREL | 9 | 30.7 | 0.09 | 567776.52 | YES | 424210 | YES | 217010 | YES | O35114 |
| **VGQEYLERL** | VGQEYLERL | 9 | 123.6 | 0.4 | 315658.192 | NO | | YES | | YES | Q3URE1 |
| **VGTAFSRL** | VGTAFSRL | 8 | 10.4 | 0.03 | 59138.1938 | NO | | YES | | YES | Q9CZ42 |
| **VGVKYVNKL** | VGVKYVNKL | 9 | 97.9 | 0.3 | | YES | 2269700 | NO | 1390200 | YES | Q9WVL3 |
| **VGVTYRTL** | VGVTYRTL | 8 | 12.2 | 0.04 | | YES | 6915400 | YES | 6355100 | YES | B1AUR6 |
| **VGYLHEGL** | VGYLHEGL | 8 | 9.3 | 0.025 | 7129926.88 | YES | 16219000 | YES | 2568000 | YES | Q6P4T2 |
| **VGYNPYSHL** | VGYNPYSHL | 9 | 4.2 | 0.01 | 164349.8 | NO | 4113000 | YES | 8202800 | YES | O54941 |

| | | | | | | | | | | | |
|---|---|---|---|---|---|---|---|---|---|---|---|
| **VGYRFVTAI** | VGYRFVTAI | 9 | 6.8 | 0.02 | 259655.701 | NO | 189760000 | NO | 168190000 | YES | Q80TM9 |
| **VGYRTQPM** | VGYRTQPM | 8 | 20.8 | 0.06 | 211541.16 | NO | 739700 | NO | | YES | Q8C2Q3 |
| **VGYRYETL** | VGYRYETL | 8 | 2.9 | 0.01 | 1133600 | NO | 61989000 | NO | 113300000 | NO | Q9DBT5 |
| **VHYVFDTTI** | VHYVFDTTI | 9 | 45.1 | 0.15 | 725574.453 | NO | 2164900 | YES | 1138900 | NO | Q9CYQ7 |
| **VIAGFNRL** | VIAGFNRL | 8 | 34.7 | 0.125 | 11570550 | YES | | YES | | YES | O70591 |
| **VIASFKVL** | VIASFKVL | 8 | 173.6 | 0.5 | 2183501.15 | NO | 23296000 | YES | 10493000 | YES | P57780 |
| **VIEEFRHL** | VIEEFRHL | 8 | 34.6 | 0.125 | | YES | 2790500 | YES | 945650 | YES | P52633 |
| **VIFKPALL** | VIFKPALL | 8 | 14.3 | 0.04 | 290202.514 | NO | | YES | | YES | A3KG59 |
| **VIFNYKGKNV** | VIFNYKGKNV | 10 | 122.2 | 0.4 | 3172357.82 | NO | 9450900 | NO | 1464000 | NO | P14211 |
| **VIFQPHIL** | VIFQPHIL | 8 | 271.4 | 0.7 | 1674100 | YES | 8919500 | YES | | YES | Q2EMV9 |
| **VILEYFTRL** | VILEYFTRL | 9 | 4.6 | 0.01 | | YES | 10700000 | YES | | YES | Q9CR08 |
| **VILSFRSL** | VILSFRSL | 8 | 4 | 0.01 | 21148000 | YES | | YES | | YES | P28660 |
| **VIMKLFPQL** | VIM(+15.99)KLFPQL | 9 | 27.8 | 0.08 | 96680.9359 | YES | 5808500 | YES | 6332300 | YES | Q64FW2 |
| **VIMKLFPQL** | VIMKLFPQL | 9 | 27.8 | 0.08 | | YES | 17881000 | YES | 406810 | YES | Q64FW2 |
| **VINELIGNL** | VINELIGNL | 9 | 477.2 | 1 | | YES | 9735200 | YES | | YES | Q99J56 |
| **VINPYKNL** | VINPYKNL | 8 | 34.5 | 0.1 | 1788869.98 | YES | 13419000 | YES | 3004300 | YES | Q8VDD5 |
| **VINSFVHV** | VINSFVHV | 8 | 47.4 | 0.15 | 1576800 | YES | 15507000 | YES | 3914700 | YES | Q9D4H8 |
| **VINVFHHL** | VINVFHHL | 8 | 11.3 | 0.04 | 137840 | YES | 10331000 | YES | 14264000 | YES | Q52KE7 |
| **VIQDFQASVL** | VIQDFQASVL | 10 | 324.3 | 0.8 | 397241.009 | NO | 3140400 | YES | 523550 | YES | Q9Z2N8 |
| **VIQDFVKM** | VIQDFVKM | 8 | 189.8 | 0.5 | 6404625.09 | NO | 23903000 | YES | 764290 | NO | Q8BHG9 |
| **VIQDFVKM** | VIQDFVKM(+15.99) | 8 | 189.8 | 0.5 | 3131963.73 | NO | 21591000 | YES | 14070000 | NO | Q8BHG9 |
| **VIQKFLYL** | VIQKFLYL | 8 | 13.7 | 0.04 | 443945.458 | NO | 21942000 | YES | | YES | Q9R111 |
| **VIQVFQQL** | VIQVFQQL | 8 | 7.5 | 0.02 | 3871989.68 | YES | 143810000 | YES | 30172000 | YES | Q8BHC4 |
| **VISDFITRL** | VISDFITRL | 9 | 68.6 | 0.2 | 322270000 | YES | | YES | 1318000 | YES | Q9DBG1 |
| **VITEFARI** | VITEFARI | 8 | 13.9 | 0.04 | | YES | | YES | 19962000 | YES | Q3TX08 |
| **VITNFSARI** | VITNFSARI | 9 | 31.1 | 0.09 | 128300 | YES | 8624500 | YES | | YES | Q3UVL4 |
| **VIVDTFHGL** | VIVDTFHGL | 9 | 79.1 | 0.25 | 321828.321 | NO | 641430 | YES | 272730 | YES | P35123:Q99K46 |
| **VIVEFRDL** | VIVEFRDL | 8 | 18.4 | 0.06 | 1364100 | YES | | YES | 6583800 | NO | Q922X9 |
| **VIVKFAQL** | VIVKFAQL | 8 | 3 | 0.01 | 89171 | YES | | YES | 9241200 | YES | Q6NS46 |

| | | | | | | | | | | | |
|---|---|---|---|---|---|---|---|---|---|---|---|
| **VIVPHIVNL** | VIVPHIVNL | 9 | 91 | 0.25 | 242384.014 | NO | | YES | 9020400 | YES | Q8BW70 |
| **VIVRFLTV** | VIVRFLTV | 8 | 40 | 0.125 | 13656000 | YES | 112680000 | YES | 7495700 | YES | P62245 |
| **VIVRFLTVM** | VIVRFLTVM | 9 | 96.4 | 0.3 | | YES | 27445000 | YES | | YES | P62245 |
| **VIWGKYAQV** | VIWGKYAQV | 9 | 16.2 | 0.05 | 119290 | YES | 3688300 | YES | | YES | Q9CR47 |
| **VIYDVSHNI** | VIYDVSHNI | 9 | 293.1 | 0.7 | 143829.447 | NO | | YES | 508770 | YES | Q99LF4 |
| **VIYNPRNL** | VIYNPRNL | 8 | 11 | 0.04 | 4286100 | YES | 225420000 | YES | 31169000 | YES | P97481 |
| **VIYPFMQGL** | VIYPFMQGL | 9 | 3.5 | 0.01 | | YES | 10682000 | YES | 3130900 | YES | Q91VV4 |
| **VLIPKLPQL** | VLIPKLPQL | 9 | 430.2 | 1 | 586720 | YES | 6630800 | YES | | YES | Q9CPZ6 |
| **VLLRYQQL** | VLLRYQQL | 8 | 12 | 0.04 | 416240 | YES | 2071900 | YES | | YES | Q9EQ20 |
| **VMYKFLTV** | VM(+15.99)YKFLTV | 8 | 6.1 | 0.015 | 254680 | NO | 62765000 | YES | 206300 | NO | Q9WVC3 |
| **VMYRVIQV** | VM(+15.99)YRVIQV | 8 | 34.4 | 0.1 | 1513785.06 | NO | 7405800 | YES | 15807000 | NO | Q61069 |
| **VMYRVIQV** | VMYRVIQV | 8 | 34.4 | 0.1 | 41132 | NO | | YES | 271150 | NO | Q61069 |
| **VNAQFPRF** | VNAQFPRF | 8 | 322.2 | 0.8 | | YES | | YES | 792860 | YES | P58742 |
| **VNFAFNQI** | VNFAFNQI | 8 | 7.1 | 0.02 | 842260 | NO | 3289700 | YES | 3455800 | YES | Q9JLF7 |
| **VNFEFPEF** | VNFEFPEF | 8 | 43.3 | 0.15 | | YES | 216020000 | YES | 4050400 | YES | P62082 |
| **VNFGRQGLNL** | VNFGRQGLNL | 10 | 270.9 | 0.7 | 309367.309 | NO | 5085300 | YES | | YES | Q99KK1 |
| **VNFIKENLL** | VNFIKENLL | 9 | 55.6 | 0.175 | 44333 | NO | 4352000 | YES | 2934200 | YES | Q61687 |
| **VNFKHEVSV** | VNFKHEVSV | 9 | 65.8 | 0.2 | | YES | | YES | 458810 | YES | Q9CQT2 |
| **VNFLHSNKL** | VNFLHSNKL | 9 | 34.1 | 0.1 | 314622.046 | YES | | YES | 1008400 | YES | P22518 |
| **VNFPFLVKL** | VNFPFLVKL | 9 | 15.1 | 0.05 | 194760 | NO | 48739000 | YES | 281020 | YES | P05132 |
| **VNFTYQFL** | VNFTYQFL | 8 | 2.9 | 0.01 | | YES | 10012000 | YES | | YES | Q3UMB9 |
| **VNFVHTNL** | VNFVHTNL | 8 | 3.6 | 0.01 | 1515537.94 | YES | 18761000 | YES | 45280000 | YES | Q9D8E6 |
| **VNIKLNQL** | VNIKLNQL | 8 | 136 | 0.4 | | YES | | YES | | YES | Q9DBC3 |
| **VNIPFVRL** | VNIPFVRL | 8 | 5 | 0.015 | 1587500 | YES | 75922000 | YES | | YES | Q8R151 |
| **VNIVINNL** | VNIVINNL | 8 | 64.5 | 0.2 | 943476.611 | YES | 1014200 | YES | | YES | Q80TY5 |
| **VNLQYSEV** | VNLQYSEV | 8 | 16.7 | 0.05 | | YES | 3769400 | YES | | YES | P62700 |
| **VNLTFRTV** | VNLTFRTV | 8 | 11 | 0.04 | 728885.444 | YES | | YES | | YES | Q8K1E6 |
| **VNLVFEKI** | VNLVFEKI | 8 | 74.7 | 0.25 | 3389325.11 | YES | 17730000 | YES | 12673000 | YES | Q6AW69 |

| | | | | | | | | | | | |
|---|---|---|---|---|---|---|---|---|---|---|---|
| **VNMVPFPRL** | VNM(+15.99)VPFPRL | 9 | 14.9 | 0.05 | 2087400 | YES | 31147000 | YES | 12370000 | YES | P99024:P68372:Q9CWF2:Q922F4:Q9ERD7:A2AQ07 |
| **VNMVPFPRL** | VNMVPFPRL | 9 | 14.9 | 0.05 | 1052800 | YES | 66374000 | YES | | YES | P99024:P68372:Q9CWF2:Q922F4:Q9ERD7:A2AQ07 |
| **VNNIFQLTV** | VNNIFQLTV | 9 | 316.5 | 0.8 | 76592.959 | NO | | YES | | YES | Q9JKY5 |
| **VNNLFVQL** | VNNLFVQL | 8 | 19.2 | 0.06 | 472225.216 | YES | | YES | 603820 | YES | P97393 |
| **VNRKYEYL** | VNRKYEYL | 8 | 20.6 | 0.06 | | YES | 342350 | YES | 956300 | YES | Q505B7 |
| **VNRVFDKL** | VNRVFDKL | 8 | 49.7 | 0.15 | | YES | 137320000 | YES | 145780000 | YES | P28076 |
| **VNSIFQHL** | VNSIFQHL | 8 | 13.7 | 0.04 | 3323500 | NO | 137350000 | YES | 32646000 | YES | Q80SU7 |
| **VNSNFYLRM** | VNSNFYLRM | 9 | 19.3 | 0.06 | 1023953.17 | NO | 2740300 | YES | 413830 | NO | Q9CQJ2 |
| **VNSNFYLRM** | VNSNFYLRM(+15.99) | 9 | 19.3 | 0.06 | 425793.437 | NO | 1605500 | YES | 3586500 | NO | Q9CQJ2 |
| **VNVAKLRYM** | VNVAKLRYM | 9 | 256.1 | 0.7 | 474874.009 | NO | 1830400 | NO | 377100 | NO | Q9CQS5 |
| **VNVAKLRYM** | VNVAKLRYM(+15.99) | 9 | 256.1 | 0.7 | 281730.273 | NO | 988290 | NO | 7248300 | NO | Q9CQS5 |
| **VNVCYKEL** | VNVC(+119.00)YKEL | 8 | 66.9 | 0.2 | 1670920.8 | NO | 2033700 | NO | 1774300 | NO | Q8BKT7 |
| **VNVDHPINL** | VNVDHPINL | 9 | 178.5 | 0.5 | 4120041.33 | YES | 17227000 | YES | 3182600 | YES | Q3UIW5 |
| **VNVDYSKL** | VNVDYSKL | 8 | 26 | 0.08 | 152950000 | YES | 230980000 | YES | 626740000 | YES | Q62425 |
| **VNVEFVRV** | VNVEFVRV | 8 | 17.9 | 0.05 | 406290 | YES | 10933000 | YES | | YES | Q8BML1 |
| **VNVERVLNV** | VNVERVLNV | 9 | 239.5 | 0.6 | 1890692.69 | YES | 13378000 | YES | 3734400 | YES | Q6DFV5 |
| **VNVPFHLAL** | VNVPFHLAL | 9 | 15.8 | 0.05 | 49018.2473 | NO | 7272000 | YES | 866010 | YES | Q8BMG7 |
| **VNVQKISNL** | VNVQKISNL | 9 | 124.3 | 0.4 | 353839.761 | NO | | YES | 375550 | YES | Q8CFI7 |
| **VNVRFTGV** | VNVRFTGV | 8 | 8.5 | 0.025 | 9266100.34 | YES | 12450000 | YES | 9581000 | YES | Q6PDI6;Q76LS9 |
| **VNVVFIGHV** | VNVVFIGHV | 9 | 28.6 | 0.09 | 320983.044 | YES | 5655200 | YES | 6154700 | YES | Q149F3;Q8R050 |
| **VNWDFVEQV** | VNWDFVEQV | 9 | 101.4 | 0.3 | 1257787.03 | YES | 13764000 | YES | 257840 | YES | Q3UIW5 |
| **VNWEKHVLI** | VNWEKHVLI | 9 | 97.3 | 0.3 | 716576.348 | NO | 7715300 | YES | 2005300 | NO | O09117 |
| **VNYDFGHM** | VNYDFGHM(+15.99) | 8 | 3.9 | 0.01 | 30730 | NO | 630870 | NO | 3974700 | NO | P58283 |
| **VNYDYSTLIL** | VNYDYSTLIL | 10 | 36.8 | 0.125 | 98677 | NO | 5810400 | YES | | YES | D2EAC2 |
| **VNYEPLGL** | VNYEPLGL | 8 | 37 | 0.125 | 142250 | NO | 8613600 | YES | 811920 | YES | Q80YE7 |
| **VNYRHLAL** | VNYRHLAL | 8 | 3.3 | 0.01 | 1033340.8 | YES | 5841000 | YES | 17383000 | YES | P08775 |
| **VNYRHLALL** | VNYRHLALL | 9 | 3.7 | 0.01 | 260890 | YES | 13170000 | YES | 61216000 | NO | P08775 |

| | | | | | | | | | | | |
|---|---|---|---|---|---|---|---|---|---|---|---|
| **VNYRVPNM** | VNYRVPNM(+15.99) | 8 | 9.3 | 0.025 | 156340 | NO | 4680000 | NO | 9607400 | YES | Q62077 |
| **VNYYFERNM** | VNYYFERNM(+15.99) | 9 | 5 | 0.01 | 5242.4 | NO | 374050 | YES | 3394900 | NO | Q9D4H9 |
| **VQEEFLQRL** | VQEEFLQRL | 9 | 367.5 | 0.9 | 19873 | NO | 468860 | YES | 567510 | YES | Q9JK81 |
| **VQFLYREL** | VQFLYREL | 8 | 5.5 | 0.015 | 1332420.67 | YES | 5220100 | YES | 3740900 | NO | B9EJR8 |
| **VQQYYRVL** | VQQYYRVL | 8 | 168.9 | 0.5 | 1297207.3 | YES | 2458300 | YES | | YES | E9PVA8 |
| **VQRSFSQV** | VQRSFSQV | 8 | 70.6 | 0.2 | 359700 | NO | 403260 | YES | 310730 | YES | Q3U1N2 |
| **VQWEYGRL** | VQWEYGRL | 8 | 8.9 | 0.025 | | YES | | YES | 844860 | YES | Q8BML9 |
| **VQYEMRTL** | VQYEMRTL | 8 | 36.7 | 0.125 | 553714.546 | NO | 2878700 | YES | 621270 | NO | Q80ZK0 |
| **VQYEPAHL** | VQYEPAHL | 8 | 15.5 | 0.05 | 261346.254 | NO | | YES | 1839400 | YES | Q9WUK6 |
| **VQYKFSHL** | VQYKFSHL | 8 | 2.3 | 0.01 | 6963987.8 | YES | 20528000 | YES | 45164000 | YES | Q9JHD1;Q9JHD2 |
| **VQYLYRVF** | VQYLYRVF | 8 | 54.4 | 0.175 | 134087.343 | NO | 4096700 | YES | 699490 | YES | Q80Y44 |
| **VQYVLPRL** | VQYVLPRL | 8 | 12.3 | 0.04 | | YES | 9579900 | YES | 13573000 | YES | Q62245 |
| **VRVFFSGL** | VRVFFSGL | 8 | 49 | 0.15 | | YES | | YES | | YES | Q66JV4;Q80YR9 |
| **VSAPYGRI** | VSAPYGRI | 8 | 94.8 | 0.3 | 287908.192 | YES | | YES | | YES | Q5SQX6;Q7TMB8 |
| **VSDAFQKL** | VSDAFQKL | 8 | 65.8 | 0.2 | | YES | | YES | 69105 | YES | Q9CQK3 |
| **VSFPFGKI** | VSFPFGKI | 8 | 7.5 | 0.02 | 222697.252 | YES | 2848300 | YES | | YES | Q6NS46 |
| **VSFTYRYL** | VSFTYRYL | 8 | 1.9 | 0.01 | 6218700 | YES | 186290000 | YES | 623610000 | YES | Q920Q4 |
| **VSIIFCEAV** | VSIIFC(+119.00)EAV | 9 | 31.1 | 0.09 | 61857.1016 | NO | 2759500 | NO | 163030 | NO | Q91V37 |
| **VSIQFYHL** | VSIQFYHL | 8 | 2.5 | 0.01 | | YES | 26313000 | YES | 36751000 | YES | Q5H8C4 |
| **VSISFKSL** | VSISFKSL | 8 | 4.4 | 0.01 | 1055582.87 | YES | 1602200 | YES | 3360500 | YES | Q3UHA3 |
| **VSLDGYFHL** | VSLDGYFHL | 9 | 35.9 | 0.125 | 225182.443 | NO | 3580500 | YES | | YES | Q8BGZ3 |
| **VSLKYAHM** | VSLKYAHM(+15.99) | 8 | 2.5 | 0.01 | 122190 | NO | 410630 | YES | 2438000 | YES | P46664 |
| **VSMDFVQRF** | VSMDFVQRF | 9 | 49.8 | 0.15 | 183179.601 | NO | | YES | | YES | Q921L5 |
| **VSNAFVRL** | VSNAFVRL | 8 | 7 | 0.02 | | YES | | YES | | YES | Q9D483 |
| **VSPEFHTL** | VSPEFHTL | 8 | 7.2 | 0.02 | 423962.654 | YES | 457930 | NO | | YES | Q7TMK6 |
| **VSPLFQKL** | VSPLFQKL | 8 | 4.7 | 0.01 | 14781000 | YES | 50370000 | YES | 81709000 | YES | Q68FL6 |
| **VSPRLTFL** | VSPRLTFL | 8 | 30 | 0.09 | 2920371.93 | YES | 81597000 | YES | 16639000 | YES | P36371 |
| **VSPTLYKQL** | VSPTLYKQL | 9 | 15.8 | 0.05 | 110770 | YES | | YES | | YES | Q3TCH7 |
| **VSQKFTSI** | VSQKFTSI | 8 | 26.9 | 0.08 | 449710.477 | NO | | YES | 1396200 | YES | Q3UCV8 |

| | | | | | | | | | | | |
|---|---|---|---|---|---|---|---|---|---|---|---|
| **VSQYYPKL** | VSQYYPKL | 8 | 10.1 | 0.03 | 901060 | YES | 198170000 | YES | 61723000 | YES | Q3TEA8 |
| **VSRSPSLL** | VSRSPSLL | 8 | 409 | 0.9 | 111432.374 | NO | | YES | | YES | Q7TPV4 |
| **VSTKFEHL** | VSTKFEHL | 8 | 13.1 | 0.04 | 1105754.12 | YES | 4105400 | YES | 8420600 | YES | B2RXC1 |
| **VSVEYTEKM** | VSVEYTEKM | 9 | 225.2 | 0.6 | 215977.888 | YES | 8288800 | YES | 1585500 | YES | O35130 |
| **VSVEYTEKM** | VSVEYTEKM(+15.99) | 9 | 225.2 | 0.6 | 2140200 | YES | 3635600 | YES | 21084000 | YES | O35130 |
| **VSVSFPHF** | VSVSFPHF | 8 | 26.2 | 0.08 | 216270.034 | NO | 6365000 | YES | | YES | Q3U1V6 |
| **VSVSFRVL** | VSVSFRVL | 8 | 7.8 | 0.02 | 292037.193 | NO | | YES | | YES | Q8BG28 |
| **VSYKNPSL** | VSYKNPSL | 8 | 34.3 | 0.1 | 129837.589 | NO | 4433500 | YES | 1540800 | YES | Q920B9 |
| **VSYKVDNL** | VSYKVDNL | 8 | 10.1 | 0.03 | 389185.563 | NO | 1072100 | YES | | YES | Q8K284 |
| **VSYKYSKV** | VSYKYSKV | 8 | 2.7 | 0.01 | 389460 | YES | 392460 | NO | 2141500 | YES | Q07113 |
| **VSYLFSHV** | VSYLFSHV | 8 | 1.8 | 0.01 | 6238600 | YES | 72119000 | YES | 57394000 | YES | Q9D7G0:Q9CS42 |
| **VSYQFPKL** | VSYQFPKL | 8 | 2.1 | 0.01 | | YES | | YES | | YES | Q924W7 |
| **VSYQHAFL** | VSYQHAFL | 8 | 2.3 | 0.01 | 488858.076 | NO | 2562900 | YES | | YES | Q9WV70 |
| **VSYWFDQRF** | VSYWFDQRF | 9 | 12.4 | 0.04 | 4160552.28 | NO | 12934000 | YES | | YES | P54751 |
| **VTFERVEQM** | VTFERVEQM | 9 | 169.8 | 0.5 | 96236.8736 | NO | | YES | | YES | Q923J1 |
| **VTFIYQKL** | VTFIYQKL | 8 | 3.2 | 0.01 | 311520 | YES | | YES | 1571800 | YES | Q8C963 |
| **VTIHYNKL** | VTIHYNKL | 8 | 14.6 | 0.05 | 358973.744 | YES | 911080 | YES | 911290 | NO | Q62083 |
| **VTIKYSKL** | VTIKYSKL | 8 | 5.5 | 0.015 | 95111.1367 | YES | 557520 | NO | | YES | Q8BGF7 |
| **VTNEFVHI** | VTNEFVHI | 8 | 123.1 | 0.4 | 398331.688 | NO | 1464500 | YES | | YES | Q9CQR6 |
| **VTPEGYAHL** | VTPEGYAHL | 9 | 12 | 0.04 | | YES | | YES | | YES | Q9Z2V5 |
| **VTVDFSKL** | VTVDFSKL | 8 | 15.3 | 0.05 | 179060 | YES | 5524900 | YES | 2485700 | YES | Q6NVF4 |
| **VTVNFRKL** | VTVNFRKL | 8 | 9.4 | 0.03 | | YES | 1145400 | YES | 323300 | YES | Q6NZJ6 |
| **VTWRVTNL** | VTWRVTNL | 8 | 11 | 0.03 | 551300 | YES | 17765000 | YES | 5214500 | YES | P10404 |
| **VTYESRKL** | VTYESRKL | 8 | 52.3 | 0.175 | | YES | | YES | | YES | Q9CY00 |
| **VTYHGFPNL** | VTYHGFPNL | 9 | 7.1 | 0.02 | 1450564.61 | YES | 17522000 | YES | 23925000 | YES | O08760 |
| **VTYSFRQSF** | VTYSFRQSF | 9 | 15 | 0.05 | 218920 | YES | 11649000 | YES | | YES | Q8R0S2:Q5DU25 |
| **VTYSKPRL** | VTYSKPRL | 8 | 32.9 | 0.1 | 122440 | YES | 1344900 | YES | | YES | Q9CPQ8 |
| **VVAEFGRI** | VVAEFGRI | 8 | 114.7 | 0.3 | 756050 | YES | | YES | | YES | Q8BXC6 |
| **VVDIFRKL** | VVDIFRKL | 8 | 142.8 | 0.4 | 3052460.99 | YES | | YES | | YES | P51791;Q61418;Q9WVD4 |

| | | | | | | | | | | | |
|---|---|---|---|---|---|---|---|---|---|---|---|
| **VVVDYGTRL** | VVVDYGTRL | 9 | 69.6 | 0.2 | 48278.1434 | YES | | YES | 211140 | YES | P27046 |
| **VVYAVRNL** | VVYAVRNL | 8 | 5.5 | 0.015 | | YES | 3012500 | YES | | YES | P28658 |
| **VVYIYHSL** | VVYIYHSL | 8 | 2.6 | 0.01 | 738520 | YES | 14886000 | YES | 11521000 | YES | Q8CD15 |
| **VVYIYKEHF** | VVYIYKEHF | 9 | 39.8 | 0.125 | 320480 | YES | 11815000 | YES | | YES | Q5XG71 |
| **VVYIYRQI** | VVYIYRQI | 8 | 4.4 | 0.01 | 1625151.22 | YES | 1835600 | YES | | YES | Q62417 |
| **VVYSYHYL** | VVYSYHYL | 8 | 2.3 | 0.01 | 307478.097 | NO | 6055000 | YES | | YES | O08811 |
| **VVYTPWSNL** | VVYTPWSNL | 9 | 10.4 | 0.03 | 1218300 | YES | 11204000 | YES | 399990 | YES | Q78PG9 |
| **VWIRNIQL** | VWIRNIQL | 8 | 441.2 | 1 | 106270 | NO | | YES | | YES | Q8R1T4 |
| **VWIYNSQL** | VWIYNSQL | 8 | 283.7 | 0.7 | 73188.4813 | NO | 2340600 | YES | 1708200 | YES | Q8BRG6 |
| **VWLEAARL** | VWLEAARL | 8 | 61.9 | 0.175 | 6873617.22 | YES | 17261000 | YES | | YES | Q91YR7 |
| **VWYRVIQI** | VWYRVIQI | 8 | 155.6 | 0.4 | 4804832.17 | YES | 23428000 | YES | | YES | P17427 |
| **VWYRVLQI** | VWYRVLQI | 8 | 209.6 | 0.6 | 4804832.17 | YES | 23428000 | YES | | YES | P17426 |
| **VYYRKPLL** | VYYRKPLL | 8 | 95.2 | 0.3 | 38638.0959 | NO | 647450 | NO | 861790 | NO | Q02053 |
| **YAMIYRNL** | YAM(+15.99)IYRNL | 8 | 8.9 | 0.025 | 1935371.6 | YES | 1009700 | YES | 95412 | YES | P23804 |
| **YAMIYRNL** | YAMIYRNL | 8 | 8.9 | 0.025 | 1504007.9 | YES | 2494400 | YES | | YES | P23804 |
| **YAYSFKYL** | YAYSFKYL | 8 | 4.6 | 0.01 | 232902.894 | NO | 4247800 | YES | 6003600 | NO | Q62383 |
| **YGYEHILTL** | YGYEHILTL | 9 | 127.1 | 0.4 | 221563.724 | NO | 1725900 | YES | | YES | Q9D2N9 |
| **YGYHFPEL** | YGYHFPEL | 8 | 10.8 | 0.03 | 2092200 | YES | 44736000 | YES | 3081300 | YES | Q9D6Z1 |
| **YNFQYISL** | YNFQYISL | 8 | 7.8 | 0.02 | 1744400 | YES | 8064300 | YES | 195380 | YES | Q8K2C8 |
| **YNWRYKNL** | YNWRYKNL | 8 | 9.2 | 0.025 | 528473.109 | NO | 4153100 | NO | 897640 | YES | Q8CFQ3 |
| **YQFVYQNL** | YQFVYQNL | 8 | 6.3 | 0.015 | 438764.731 | NO | 2842200 | NO | 1895500 | NO | Q5NCI0 |
| **YSLVYQAL** | YSLVYQAL | 8 | 7.3 | 0.02 | 130382.929 | NO | 5381600 | YES | | YES | Q8BSP2 |
| **YSPAYAHL** | YSPAYAHL | 8 | 5.6 | 0.015 | 3701493.12 | YES | 5109700 | YES | 7457600 | YES | O09012 |
| **YSPEFKGQI** | YSPEFKGQI | 9 | 498.2 | 1.1 | 1354467.97 | YES | 6545500 | YES | 1048500 | YES | Q8CIE6 |
| **YTFVYRVL** | YTFVYRVL | 8 | 14.1 | 0.04 | 428130.607 | YES | 2060000 | YES | | YES | Q62136 |

### Supplementary Table 4

| PEPTIDE NUMBER | Sequence | Length | H-2K$^b$ IC50 (nM) | Product of Spectral Intensity Values | Found in DIA | CD44$^+$ PD-1$^{hi}$ Male B10.BR | CD44$^+$ PD-1$^-$ Male B10.BR | CD44$^+$ PD-1$^{hi}$ Female B10.BR | CD44$^+$ PD-1$^-$ Female B10.BR | CD44$^+$ PD-1$^{hi}$ Male BALB/c | CD44$^+$ PD-1$^-$ Male BALB/c |
|---|---|---|---|---|---|---|---|---|---|---|---|
| **PEPTIDE 1** | SNYLFTKL | 8 | 2.4 | 8.67E+27 | YES | 19.5 (18.1-22.1) | 0.5 (0.2-1.1) | 14.6 (10.3-17) | 0.1 (0.1-0.2) | 20.7 (17.3-26.2) | 1.1 (0.6-1.7) |
| **PEPTIDE 2** | ATLVFHNL | 8 | 6.9 | 6.86E+25 | YES | 7.1 (2.6-11.6) | 0.2 (0.1-0.2) | 4.6 (4-5.6) | 0.1 (0-0.1) | 17.8 (14.4-22.9) | 2.3 (1.9-2.9) |
| **PEPTIDE 3** | VGPRYTNL | 8 | 4.6 | 1.53E+26 | NO | 12.2 (10.8-14.5) | 0.2 (0.1-0.3) | 8.1 (6.7-9.7) | 0.1 (0.1-0.1) | 6.7 (5.8-8.1) | 1.2 (1-1.6) |
| **PEPTIDE 4** | RTYTYEKL | 8 | 9.4 | 1.78E+25 | YES | 8.4 (7.7-9) | 0.2 (0.1-0.3) | 8 (7.1-8.6) | 0.1 (0.1-0.1) | 16.2 (13.2-17.7) | 0.6 (0.4-0.8) |
| **PEPTIDE 5** | INFDFPKL | 8 | 6.2 | 1.41E+26 | YES | 9.1 (5.2-15.1) | 0.1 (0-0.1) | 2.8 (1.5-4.3) | 0.1 (0.1-0.1) | 5.4 (2.9-9) | 0.8 (0.3-1) |
| **PEPTIDE 6** | SVYVYKVL | 8 | 6.7 | 1.25E+25 | YES | 11.9 (9.8-13.3) | 0.3 (0.2-0.4) | 10.1 (8.3-12) | 0.1 (0-0.1) | 13.9 (10.1-21.4) | 0.7 (0.2-0.9) |
| **PEPTIDE 7** | VNVDYSKL | 8 | 26 | 2.21E+25 | YES | 2.9 (1.3-3.7) | 0.2 (0-0.4) | 2.7 (1.7-3.9) | 0.1 (0.1-0.1) | 4.3 (2.1-7.2) | 2.2 (0.5-3.4) |
| **PEPTIDE 8** | ASYEFVQRL | 9 | 4.2 | 2.89E+24 | YES | 0.6 (0.5-0.9) | 0.2 (0.1-0.2) | 1 (0.8-1.3) | 0.1 (0.1-0.1) | 1 (0.5-1.6) | 1.4 (0.4-2.1) |
| **PEPTIDE 9** | HIYEFPQL | 8 | 7.9 | 3.85E+24 | YES | 7.9 (5.6-9.5) | 0.2 (0.1-0.2) | 6.9 (2.7-9.9) | 0.1 (0-0.1) | 5.2 (1.8-8.6) | 0.6 (0.4-0.8) |
| **PEPTIDE 10** | VAFDFTKV | 8 | 10.6 | 6.15E+23 | YES | 6.1 (3.1-10.7) | 0.2 (0.1-0.2) | 2.2 (1.4-2.9) | 0.1 (0-0.1) | 4.1 (1-8.9) | 1.7 (1-2.3) |
| **PEPTIDE 11** | TSVRFTQL | 8 | 5.3 | 3.36E+23 | YES | 2.2 (1.2-3.6) | 0.1 (0.1-0.2) | 1.7 (1.4-1.9) | 0 (0-0.1) | 3.6 (2.7-4.6) | 1.5 (0.9-2) |
| **PEPTIDE 12** | AVVAFVMKM | 9 | 339.2 | 2.74E+23 | YES | 0.4 (0.2-0.5) | 0.3 (0.2-0.5) | 0.6 (0.5-0.6) | 0.4 (0.3-0.5) | 0.6 (0.5-0.7) | 0.2 (0.1-0.3) |
| **PEPTIDE 13** | SAYEFYHAL | 9 | 2.8 | 1.99E+23 | YES | 2.8 (1.4-5.5) | 0.2 (0.2-0.2) | 1 (0.6-1.7) | 0.1 (0-0.1) | 2.2 (0.2-6) | 0.9 (0.5-1.4) |
| **PEPTIDE 14** | SVIKFENL | 8 | 7.1 | 2.26E+23 | YES | 2.6 (1-5.7) | 0.1 (0.1-0.1) | 1 (0.4-1.6) | 0.1 (0-0.2) | 1.3 (0.5-1.9) | 1.4 (1.1-1.7) |
| **PEPTIDE 15** | SGYDFENRL | 9 | 14.2 | 6.01E+22 | YES | 0.6 (0.6-0.6) | 0.1 (0.1-0.2) | 0.5 (0.3-1) | 0 (0-0) | 0.4 (0.1-0.6) | 1 (0.3-1.5) |
| **PEPTIDE 16** | VSFTYRYL | 8 | 1.9 | 7.22E+23 | YES | 5.4 (2.9-9.7) | 0.4 (0.3-0.5) | 10 (7.9-13.5) | 0.1 (0.1-0.2) | 11.2 (7.8-16.6) | 2.2 (1.5-3) |
| **PEPTIDE 17** | ISFKFDHL | 8 | 2.7 | 2.31E+23 | YES | 1.6 (1.3-1.9) | 0.1 (0-0.1) | 1.6 (0.8-2.6) | 0.1 (0-0.1) | 2.3 (2.1-2.5) | 0.9 (0.6-1.1) |
| **PEPTIDE 18** | RNYSYEKL | 8 | 11.6 | 9.41E+22 | YES | 6.7 (6.2-7) | 0.1 (0.1-0.1) | 7.6 (6-8.6) | 0.1 (0.1-0.2) | 17.5 (12.4-23) | 0.6 (0.3-0.8) |
| **PEPTIDE 19** | SGYKFGVL | 8 | 4.6 | 6.19E+22 | NO | 0.5 (0.4-0.7) | 0.1 (0.1-0.2) | 1.2 (0.9-1.8) | 0.1 (0-0.1) | 3.3 (2.3-3.9) | 0.3 (0.1-0.4) |
| **PEPTIDE 20** | VGFDYKERL | 9 | 37.1 | 5.11E+23 | YES | 2.1 (1.1-3.4) | 0.1 (0.1-0.2) | 5.1 (1.9-6.8) | 0.1 (0.1-0.1) | 0.9 (0.2-2) | 0.7 (0.5-1.2) |
| **PEPTIDE 21** | SSPKFSEL | 8 | 9.3 | 8.97E+22 | NO | 2 (0.7-4.3) | 0.1 (0-0.1) | 0.7 (0.4-1.1) | 0.1 (0-0.1) | 1.1 (0.7-1.8) | 0.9 (0.7-1.3) |
| **PEPTIDE 22** | ASPEFTKL | 8 | 20.9 | 1.03E+23 | YES | 1.1 (1-1.4) | 0.1 (0-0.2) | 1 (0.6-1.2) | 0.1 (0.1-0.3) | 2.9 (2.3-4) | 1.4 (1.1-1.9) |

| | | | | | | | | | | | |
|---|---|---|---|---|---|---|---|---|---|---|---|
| **PEPTIDE 23** | ATQVYPKL | 8 | 118.9 | 1.17E+23 | YES | 3.1 (1.7-4.2) | 0.2 (0.2-0.2) | 4.1 (3.2-5.3) | 0.1 (0-0.2) | 7.1 (6.3-7.7) | 0.7 (0.2-1.2) |
| **PEPTIDE 24** | SGLKYVNV | 8 | 14.6 | 7.07E+22 | YES | 3.1 (1.8-5.1) | 0.2 (0.2-0.3) | 2 (1.4-3.1) | 0.1 (0.1-0.2) | 3.7 (3.2-4) | 2.2 (1.7-2.4) |
| **PEPTIDE 25** | QIIPFKTL | 8 | 139.6 | 2.57E+23 | YES | 1.9 (0.8-2.5) | 0.2 (0.1-0.3) | 2.5 (1.5-3.5) | 0.2 (0.1-0.3) | 1.9 (1.8-2) | 0.8 (0.5-1.1) |
| **PEPTIDE 26** | ATRSFPQL | 8 | 37.4 | 2.00E+22 | YES | 1.8 (0.6-3.5) | 0.1 (0-0.1) | 1.1 (0.5-1.7) | 0 (0-0.1) | 11.3 (5.7-18.1) | 0.1 (0-0.2) |
| **PEPTIDE 27** | SGYIYHKL | 8 | 4.8 | 4.96E+21 | YES | 13.5 (9.7-16.8) | 0.4 (0.3-0.5) | 8.1 (5.8-11.1) | 0.1 (0.1-0.2) | 23.5 (14.4-29.5) | 0.5 (0.2-0.9) |
| **PEPTIDE 28** | VSPLFQKL | 8 | 4.7 | 6.08E+22 | YES | 3.1 (1.3-5.3) | 0.2 (0.1-0.2) | 3 (2.3-4) | 0.1 (0-0.1) | 15.4 (6.5-23.7) | 0.5 (0.3-0.8) |
| **PEPTIDE 29** | QSIAFISRL | 9 | 13.5 | 8.84E+22 | YES | 2.6 (1.2-4.8) | 0.5 (0.4-0.5) | 3 (2.4-3.9) | 0.3 (0.1-0.4) | 0.8 (0.4-1.3) | 0.8 (0.5-1.1) |
| **PEPTIDE 30** | SSYTFPKM | 8 | 3.4 | 4.75E+21 | YES | 5.4 (3.3-8.2) | 0.2 (0.2-0.3) | 3.1 (2-5) | 0.1 (0-0.1) | 9.6 (7.1-12.4) | 0.2 (0.1-0.3) |
| **PEPTIDE 31** | VSQYYPKL | 8 | 10.1 | 1.10E+22 | YES | 2.3 (2-2.5) | 0.1 (0.1-0.1) | 6.3 (4.5-7.5) | 0 (0-0) | 12.5 (3.8-21.8) | 0.2 (0.2-0.3) |
| **PEPTIDE 32** | QSIEFSRL | 8 | 5.5 | 2.19E+22 | YES | 1.7 (1.1-2.1) | 0.1 (0.1-0.1) | 2.9 (1.3-5.6) | 0.1 (0-0.1) | 0.7 (0.5-1) | 0.4 (0.3-0.6) |
| **PEPTIDE 33** | SGIDFKQL | 8 | 68.8 | 8.01E+21 | YES | 1 (0.8-1.2) | 0 (0-0.1) | 2.9 (0.4-7.1) | 0 (0-0) | 0.2 (0.2-0.3) | 0.2 (0.1-0.4) |
| **PEPTIDE 34** | AVLSFSTRL | 9 | 20.1 | 3.24E+22 | YES | 4.8 (3.3-5.6) | 0.7 (0.6-0.9) | 6.5 (4.1-10.5) | 0.2 (0.1-0.3) | 0.7 (0.4-1.2) | 0.8 (0.6-1.2) |
| **PEPTIDE 35** | VGPRYTQL | 8 | 6.8 | 3.41E+22 | NO | 3.9 (1.9-5.8) | 0.1 (0-0.1) | 8.5 (4.6-12.5) | 0 (0-0) | 3.1 (1.4-4) | 0.1 (0-0.1) |
| **PEPTIDE 36** | RNYEYLIRL | 9 | 11.9 | 3.28E+22 | YES | 2 (1.6-2.2) | 0.2 (0.1-0.3) | 2.8 (1.2-5.4) | 0 (0-0.1) | 0.3 (0.2-0.4) | 0.3 (0.2-0.6) |
| **PEPTIDE 37** | VAYKFPEL | 8 | 2.6 | 1.56E+22 | YES | 0.9 (0.6-1.3) | 0.1 (0-0.1) | 3.7 (2.5-5.4) | 0 (0-0) | 1.9 (0.6-3.5) | 0.6 (0.2-0.9) |
| **PEPTIDE 38** | LQYEFTKL | 8 | 10.8 | 7.90E+22 | YES | 1 (0.6-1.8) | 0 (0-0.1) | 2.8 (0.9-6) | 0 (0-0) | 6.1 (1.1-8.6) | 0.3 (0.2-0.4) |
| **PEPTIDE 39** | VSYLFSHV | 8 | 1.8 | 2.58E+22 | YES | 4.4 (4-5.1) | 0.1 (0.1-0.1) | 7.3 (4.1-13.5) | 0 (0-0) | 8.5 (3.9-13.6) | 1.8 (1.1-2.6) |
| **PEPTIDE 40** | SAFSFRTL | 8 | 3.3 | 4.94E+22 | YES | 5.2 (4.5-6.4) | 0.4 (0.3-0.5) | 6.1 (4.6-9.1) | 0.1 (0.1-0.2) | 19.4 (13.3-22.9) | 3.1 (2.8-3.2) |
| **PEPTIDE 41** | SNYHFYSSI | 9 | 3.9 | 2.15E+21 | YES | 1.2 (0.9-1.4) | 0.4 (0.3-0.4) | 2.5 (1-5.3) | 0.1 (0.1-0.1) | 0.6 (0.4-0.8) | 0.6 (0.5-0.7) |
| **PEPTIDE 42** | HGYTFANL | 8 | 3 | 1.28E+22 | YES | 1.1 (0.8-1.4) | 0.1 (0-0.1) | 2.8 (1.1-5.8) | 0 (0-0) | 15 (13.2-17) | 1.1 (0.6-2) |
| **PEPTIDE 43** | SGYDFSRL | 8 | 3.6 | 7.90E+21 | YES | 1.9 (1.9-2.1) | 0.1 (0.1-0.1) | 2.4 (2-2.9) | 0.1 (0-0.2) | 4.3 (1-7.4) | 0.7 (0.7-0.8) |
| **PEPTIDE 44** | TAYEFAKL | 8 | 2.4 | 9.71E+21 | YES | 1.6 (1.4-1.8) | 0.1 (0-0.1) | 2.9 (2.2-3.6) | 0.1 (0-0.1) | 7.5 (0.7-18) | 0.7 (0.2-0.9) |
| **PEPTIDE 45** | VADKFTEL | 8 | 140.9 | 1.13E+22 | YES | 0.3 (0.2-0.5) | 0 (0-0) | 0.7 (0.4-1.2) | 0 (0-0.1) | 0.5 (0.5-0.5) | 0.2 (0.1-0.2) |
| **PEPTIDE 46** | TAFRFSEL | 8 | 3.3 | 2.39E+22 | NO | 0.9 (0.6-1.2) | 0 (0-0) | 1.8 (1.6-2.1) | 0.1 (0-0.1) | 5.2 (1.4-9.2) | 0.7 (0.4-1) |
| **PEPTIDE 47** | KILTFDQL | 8 | 66.5 | 7.62E+21 | NO | 2.3 (1.2-4.2) | 0.1 (0-0.1) | 3.2 (2.8-3.8) | 0.1 (0.1-0.1) | 9.3 (6.2-11.1) | 1.8 (1-2.4) |
| **PEPTIDE 48** | VGYRYETL | 8 | 2.9 | 7.96E+21 | NO | 7.8 (2.7-16.1) | 0.1 (0-0.1) | 3.9 (2.6-6.7) | 0.1 (0.1-0.1) | 4.4 (2.7-6.7) | 1.4 (0.9-1.9) |
| **PEPTIDE 49** | RVAEFTTNL | 9 | 132.4 | 2.10E+21 | YES | 1.5 (0.7-2) | 0.1 (0.1-0.1) | 1.2 (1.1-1.3) | 0.1 (0.1-0.1) | 0.2 (0.2-0.4) | 1.2 (1-1.5) |
| **PEPTIDE 50** | RSLKFYSL | 8 | 5.7 | 6.97E+21 | NO | 3 (2.1-4.5) | 0.3 (0.3-0.3) | 2.3 (1.9-2.7) | 0.3 (0.2-0.3) | 2.2 (1.4-3.4) | 2 (1.5-2.5) |
| **PEPTIDE 51** | TNQDFIQRL | 9 | 149.4 | 1.80E+21 | YES | 2.5 (1.7-3.7) | 0.2 (0.2-0.2) | 1.8 (1.4-2.3) | 0 (0-0.1) | 0.6 (0.5-0.9) | 2 (1.8-2.3) |

| | | | | | | | | | | | |
|---|---|---|---|---|---|---|---|---|---|---|---|
| **PEPTIDE 52** | TTYKYFAL | 8 | 2.7 | #N/A | YES | 1.6 (1.2-2.1) | 0.1 (0-0.1) | 1.6 (0.9-2) | 0 (0-0) | 2 (1.7-2.3) | 0.5 (0.3-0.7) |
| **PEPTIDE 53** | VGITYQHI | 8 | 19.6 | 9.52E+21 | YES | 4.3 (3.5-4.7) | 0.3 (0.2-0.4) | 4.8 (2.8-8.5) | 0.2 (0.1-0.3) | 5.3 (3.1-7.5) | 2.3 (2.1-2.5) |
| **PEPTIDE 54** | TAPQYYRL | 8 | 11.9 | 6.99E+21 | YES | 4.3 (3.4-5) | 0.6 (0.5-0.7) | 3.6 (2.3-5.8) | 0.2 (0.2-0.3) | 1.6 (1.2-2.1) | 2.7 (2.6-2.9) |
| **PEPTIDE 55** | KNYDFAQVL | 9 | 29.5 | 3.14E+21 | YES | 0.3 (0.2-0.6) | 0.1 (0-0.1) | 0.4 (0.3-0.5) | 0 (0-0) | 0.5 (0.1-1.2) | 0.5 (0.3-0.8) |
| **PEPTIDE 56** | FAYRFSNL | 8 | 2 | 3.16E+21 | NO | 4.8 (1.7-7.7) | 0.1 (0.1-0.2) | 2 (1.8-2.3) | 0 (0-0.1) | 3.6 (3.1-4.2) | 0.5 (0.4-0.8) |
| **PEPTIDE 57** | ATYTFIQQL | 9 | 7.6 | 8.87E+21 | YES | 1 (0.7-1.7) | 0.2 (0-0.5) | 4.4 (2.8-6.4) | 1.8 (0-4.2) | 1.3 (0.2-2.7) | 2.7 (0.4-5.6) |
| **PEPTIDE 58** | SGYKFFSL | 8 | 2.6 | 1.37E+22 | YES | 1 (0.7-1.3) | 0.1 (0.1-0.2) | 1.7 (0.8-2.7) | 0 (0-0.1) | 1.8 (1.5-2.2) | 0.4 (0.3-0.6) |
| **PEPTIDE 59** | RAYLFAHV | 8 | 2.5 | 1.79E+21 | YES | 4.9 (4.1-6.4) | 0.5 (0.4-0.7) | 6.7 (5.5-7.6) | 0.2 (0.2-0.3) | 12.6 (9.1-16.3) | 1.9 (1.8-2) |
| **PEPTIDE 60** | VIVRFLTV | 8 | 40 | 1.15E+22 | YES | 7.3 (6.3-9.1) | 3.3 (3-3.4) | 9.3 (8.9-10.1) | 0.9 (0.6-1.1) | 7.5 (4.4-10.2) | 1.2 (0.8-1.5) |
| **PEPTIDE 61** | SGYKYVGM | 8 | 4.8 | 6.63E+21 | YES | 2 (1.7-2.4) | 0.2 (0.1-0.3) | 2.6 (2.1-3) | 0.1 (0-0.1) | 7.1 (4-10.2) | 0.1 (0.1-0.2) |
| **PEPTIDE 62** | RNYQFDFL | 8 | 11.1 | 4.42E+21 | YES | 4.1 (2.5-7) | 0.2 (0.2-0.3) | 12.9 (10.9-16.9) | 0.1 (0-0.1) | 5.9 (3.5-7.8) | 0.1 (0-0.3) |
| **PEPTIDE 63** | ISILYHQL | 8 | 3.7 | 1.15E+20 | NO | 1.3 (1-1.6) | 0.3 (0.2-0.3) | 2.4 (2.2-2.7) | 0.1 (0-0.1) | 4.2 (2-8.2) | 0.2 (0.1-0.3) |
| **PEPTIDE 64** | VNSIFQHL | 8 | 13.7 | 1.49E+22 | NO | 1.3 (1.1-1.8) | 0.2 (0.1-0.3) | 3.4 (2.9-4.3) | 0 (0-0) | 7.8 (3.7-12.1) | 0.3 (0.2-0.4) |
| **PEPTIDE 65** | SSPHYTTL | 8 | 12.2 | 7.07E+21 | YES | 3.6 (1.6-6.8) | 0.1 (0.1-0.1) | 5.5 (4.9-6.1) | 0.1 (0.1-0.2) | 7.7 (5.3-11.1) | 0.9 (0.7-1) |
| **PEPTIDE 66** | VGYRFVTAI | 9 | 6.8 | 8.29E+21 | NO | 2.5 (0.4-3.6) | 0.1 (0-0.1) | 5.8 (5.2-6.5) | 0.1 (0.1-0.2) | 3.9 (1-6.7) | 0.2 (0-0.3) |
| **PEPTIDE 67** | TGPKYIHL | 8 | 26.7 | 8.63E+21 | YES | 3.4 (0.5-8.9) | 0 (0-0.1) | 2.1 (1.9-2.4) | 0.1 (0.1-0.2) | 2.8 (1.8-4) | 0.4 (0.1-0.8) |
| **PEPTIDE 68** | VQYKFSHL | 8 | 2.3 | 6.46E+21 | YES | 3.6 (0.9-8.8) | 0 (0-0) | 2.8 (1.7-3.9) | 0.1 (0-0.1) | 4.7 (3.4-5.4) | 0 (0-0.1) |
| **PEPTIDE 69** | RVLLFSQM | 8 | 31.8 | 1.31E+22 | YES | 0.8 (0.5-1.3) | 0.3 (0.2-0.3) | 0.2 (0.1-0.3) | 0.2 (0.1-0.3) | 0.6 (0.4-1) | 0 (0-0) |
| **PEPTIDE 70** | RVLIFSQM | 8 | 42.3 | 1.31E+22 | YES | 1.1 (1-1.3) | 0.3 (0.2-0.4) | 0.8 (0.3-1.3) | 0.4 (0.2-0.6) | 0.2 (0.1-0.3) | 0 (0-0.1) |
| **PEPTIDE 71** | KIFEFKETL | 9 | 63.4 | 4.45E+21 | NO | 0.5 (0.1-0.9) | 0 (0-0) | 0.9 (0.6-1.3) | 0 (0-0) | 0.2 (0.1-0.3) | 0.1 (0.1-0.2) |
| **PEPTIDE 72** | TTYKYEMI | 8 | 28.6 | 6.70E+21 | NO | 3.6 (0.4-9.6) | 0 (0-0.1) | 2.2 (1.7-2.6) | 0.1 (0.1-0.1) | 13.3 (5.4-19.8) | 0.4 (0.2-0.5) |
| **PEPTIDE 73** | ISVSFYHV | 8 | 6 | 1.12E+21 | YES | 2.7 (2-3.8) | 0.1 (0-0.2) | 2.9 (2.5-3.1) | 0.1 (0.1-0.1) | 4.4 (2-8) | 0.9 (0.4-1.7) |
| **PEPTIDE 74** | KNFPFERL | 8 | 11.2 | 2.52E+20 | YES | 4.9 (2.2-6.3) | 0.1 (0-0.1) | 6.3 (5-9) | 0.1 (0.1-0.1) | 16.3 (7.6-22.9) | 0.6 (0.4-0.7) |
| **PEPTIDE 75** | RVYEFLDKL | 9 | 34.9 | 2.66E+21 | YES | 0.6 (0.1-1.2) | 0 (0-0) | 1.3 (1-1.6) | 0 (0-0.1) | 0.3 (0.2-0.5) | 0.2 (0.1-0.3) |
| **PEPTIDE 76** | AFYYIHNL | 8 | 87.6 | 9.76E+20 | YES | 0.7 (0.2-1.6) | 0 (0-0) | 2 (1.9-2) | 0.1 (0.1-0.1) | 0.8 (0.4-1.6) | 0.5 (0.2-0.9) |
| **PEPTIDE 77** | TNYNFQYI | 8 | 7 | 1.28E+21 | YES | 4.1 (2.8-6.6) | 0.1 (0.1-0.1) | 3.9 (3.4-4.5) | 0 (0-0) | 8.4 (4.6-16) | 0.4 (0.2-0.8) |
| **PEPTIDE 78** | RAYLFNSV | 8 | 7 | 8.32E+20 | YES | 5.1 (4.2-6) | 0.1 (0.1-0.2) | 6 (3.5-7.4) | 0.1 (0.1-0.1) | 18 (13.6-21.1) | 1 (0.8-1.2) |
| **PEPTIDE 79** | ISARFVQL | 8 | 6.4 | 5.36E+20 | NO | 0.9 (0.6-1.1) | 0.1 (0-0.1) | 2.3 (2.1-2.8) | 0 (0-0) | 5.5 (5-6) | 0.5 (0.3-0.8) |
| **PEPTIDE 80** | SAYEVIKL | 8 | 56.8 | 4.63E+20 | YES | 1 (0.6-1.2) | 0.1 (0-0.1) | 2.5 (1.7-3.3) | 0.1 (0-0.2) | 3.9 (1.7-5.6) | 0.2 (0.1-0.3) |

| | | | | | | | | | | | |
|---|---|---|---|---|---|---|---|---|---|---|---|
| **PEPTIDE 81** | LAPVYQRL | 8 | 17.7 | #N/A | YES | 1.1 (1-1.3) | 0.1 (0.1-0.1) | 2.1 (1.9-2.2) | 0.2 (0.1-0.3) | 11 (7.7-16.7) | 0.5 (0.2-1) |
| **PEPTIDE 82** | KGFTFSAL | 8 | 4.2 | #N/A | YES | 2.3 (0.8-5) | 0 (0-0) | 2.4 (1.2-4.3) | 0 (0-0.1) | 10.3 (3.9-16) | 0.2 (0.1-0.3) |
| **PEPTIDE 83** | VSPRLTFL | 8 | 30 | 3.96E+21 | YES | 1.4 (1.1-2) | 0.1 (0.1-0.2) | 3.2 (1.3-6) | 0.1 (0.1-0.1) | 5.8 (5.3-6.6) | 0.6 (0.2-0.8) |
| **PEPTIDE 84** | VNMVPFPRL | 9 | 14.9 | 8.64E+20 | YES | 2.8 (1.6-4.6) | 0.6 (0.3-0.9) | 1.6 (1.3-2.1) | 0.1 (0.1-0.2) | 2.2 (1-4.1) | 1.1 (0.9-1.4) |
| **PEPTIDE 85** | TNVLFNHL | 8 | 11.1 | 7.84E+20 | YES | 0.4 (0.3-0.5) | 0.1 (0-0.1) | 0.9 (0.6-1.5) | 0.2 (0.2-0.3) | 0.3 (0.1-0.6) | 0.3 (0.3-0.4) |
| **PEPTIDE 86** | IGPTYYQRL | 9 | 3.9 | 4.04E+18 | YES | 2.1 (1.4-3) | 0.1 (0-0.1) | 2.8 (2.3-3) | 0.1 (0-0.1) | 7.4 (4.1-10.4) | 0.5 (0.4-0.8) |
| **PEPTIDE 87** | SVYTHSYL | 8 | 4.6 | 2.27E+20 | YES | 5.9 (2.8-11.1) | 0 (0-0.1) | 6.9 (4.6-9.6) | 0 (0-0.1) | 15 (8.4-26.4) | 0.2 (0.1-0.3) |
| **PEPTIDE 88** | KTYQFLNDI | 9 | 89.3 | 2.23E+20 | YES | 0.9 (0.5-1.5) | 0 (0-0.1) | 0.9 (0.5-1.4) | 0 (0-0) | 2.2 (0.6-4.3) | 0 (0-0.1) |
| **PEPTIDE 89** | TSFRYSSL | 8 | 2.3 | 2.34E+20 | NO | 1.9 (1-3.4) | 0.1 (0-0.2) | 1.2 (0.6-1.8) | 0.1 (0-0.1) | 6.3 (3.9-7.5) | 0.1 (0-0.3) |
| **PEPTIDE 90** | AMYIFLHTV | 9 | 31.7 | 3.31E+21 | YES | 0.7 (0.7-0.8) | 1.7 (1.2-2.1) | 0.6 (0.4-0.7) | 0.3 (0.1-0.3) | 0.8 (0.1-1.2) | 0.3 (0.1-0.7) |
| **PEPTIDE 91** | SGLKYVAV | 8 | 39.3 | 6.19E+20 | NO | 2.5 (2-2.7) | 0.1 (0.1-0.1) | 1.1 (0.5-1.6) | 0.1 (0.1-0.2) | 3.1 (0.7-5.5) | 0.9 (0.6-1.1) |
| **PEPTIDE 92** | VNFVHTNL | 8 | 3.6 | 1.29E+21 | YES | 3.5 (2.1-5.2) | 0.2 (0.1-0.3) | 3.8 (2.8-4.5) | 0 (0-0) | 9.6 (7.6-11) | 0.3 (0-0.5) |
| **PEPTIDE 93** | IFYYVQKL | 8 | 52.7 | 2.24E+21 | YES | 1.7 (0.9-2.9) | 0.2 (0.1-0.3) | 2.1 (1.9-2.3) | 0.1 (0-0.1) | 5.7 (2.6-7.6) | 0.1 (0-0.4) |
| **PEPTIDE 94** | IRYFPTQAL | 9 | 437.1 | 1.12E+20 | YES | 0.4 (0.3-0.5) | 0.1 (0.1-0.1) | 0.5 (0.3-0.7) | 0.1 (0-0.1) | 0.5 (0.1-1.3) | 0.1 (0-0.2) |
| **PEPTIDE 95** | KNVLFSHL | 8 | 7.9 | 1.63E+21 | YES | 1.2 (0.6-2.1) | 0.1 (0-0.1) | 1.4 (1.1-1.7) | 0 (0-0) | 5.3 (2.6-7.9) | 0.3 (0.1-0.5) |
| **PEPTIDE 96** | SSPKFSEI | 8 | 53.5 | 8.97E+22 | NO | 0.8 (0.3-1.8) | 0.1 (0-0.1) | 0.8 (0.7-0.9) | 0 (0-0.1) | 4.1 (2.3-6.1) | 0.2 (0-0.4) |
| **PEPTIDE 97** | INFDFNTI | 8 | 19.8 | #N/A | YES | 3.7 (1-6.4) | 2.4 (1.6-4) | 3.2 (2.3-4.1) | 0.4 (0.3-0.5) | 3.4 (2.7-4.8) | 0.1 (0-0.2) |
| **PEPTIDE 98** | KIITYRNL | 8 | 7.5 | #N/A | YES | 2.2 (1.7-2.8) | 0.4 (0.2-0.5) | 3.7 (2.9-5.1) | 0.2 (0.1-0.2) | 2.1 (1.7-2.4) | 1.1 (1-1.2) |
| **PEPTIDE 99** | KVLHFYNV | 8 | 52.8 | #N/A | NO | 3.5 (2.9-4.2) | 0.3 (0.2-0.4) | 4.2 (3.1-4.8) | 0.2 (0.1-0.2) | 3.4 (1.9-4.3) | 1 (0.6-1.4) |
| **PEPTIDE 100** | KVITFIDL | 8 | 135.5 | 4.55E+20 | YES | 2.7 (0.4-6) | 1.4 (0-4.1) | 1.5 (0.7-2.9) | 0.1 (0-0.1) | 3.5 (1.6-7) | 0.2 (0-0.3) |

#### Supplementary Table 5

| Antibodies used for IHC | | | | |
|---|---|---|---|---|
| Antibody Target | Clone | Format | Supplier | Catalogue # |
| H-2K$^b$ | AF6-88.5 | FITC | BD Biosciences | 553569 |
| H-2K$^d$ | SF1-1.1 | FITC | BD Biosciences | 553564 |
| H-2D$^b$/H-2K$^b$ | 28-8-6 | FITC | BioLegend | 114605 |
| CD4 | GK1.5 | FITC | BD Biosciences | 553729 |
| CD8a | 53-6.7 | FITC | BioLegend | 100706 |
| F4/80 | BM8 | FITC | BioLegend | 123108 |
| CD19 | 6D5 | FITC | BioLegend | 115506 |
| CD11c | N418 | FITC | BioLegend | 117306 |
| FITC | polyclonal | HRP | Bio-Rad | 4510-7864 |

| Antibodies used for hepatocyte/RMA-S staining (flow) | | | | |
|---|---|---|---|---|
| Antibody Target | Clone | Format | Supplier | Catalogue # |
| H-2K$^b$ | AF6-88.5 | biotin | BD Biosciences | 553568 |
| H-2K$^b$ | Y-3 | purified | WEHI* | N/A |
| H-2K$^d$ | SF1-1.1 | biotin | BioLegend | 116604 |
| H-2K$^b$-SIINFEKL | 25D-1.16 | APC | BioLegend | 141606 |
| Mouse IgG2b | RMG2b-1 | PE | BioLegend | 406708 |
| biotin | streptavidin | PE | BioLegend | 405204 |

| Antibodies used for Des-RAG Adoptive Transfer/Screening experiment | | | | |
|---|---|---|---|---|
| Antibody Target | Clone | Format | Supplier | Catalogue # |
| CD4 | GK1.5 | FITC | BD Biosciences | 553729 |
| CD8a | 53-6.7 | PE | BD Biosciences | 553033 |
| CD90.2 | 53-2.1 | PerCPCy5.5 | BioLegend | 140322 |
| CD44 | IM7 | APC | BD Biosciences | 559250 |
| PD-1 | 29F.1A12 | BV421 | BioLegend | 135218 |
| Vbeta2 | B20.6 | PE | BioLegend | 127908 |
| CD45.1 | A20 | BV711 | BioLegend | 110730 |
| CD45.2 | 104 | PerCPCy5.5 | BioLegend | 109828 |

| Antibodies used for multimer staining | | | | |
|---|---|---|---|---|
| Antibody Target | Clone | Format | Supplier | Catalogue # |
| CD8a | KT-15 | FITC | Invitrogen | MA5-16760 |
| CD90.2 | 53-2.1 | PerCPCy5.5 | BioLegend | 140322 |
| CD44 | IM7 | APC | BioLegend | 103012 |
| CD44 | IM7 | BV605 | BioLegend | 103047 |
| PD-1 | 29F.1A12 | BV421 | BioLegend | 135218 |
| CD19 | 6D5 | PECy7 | BioLegend | 115520 |
| CD14 | Sa14-2 | PECy7 | BioLegend | 740357 |
| PE | PE001 | biotin | BioLegend | 408104 |
| APC | APC003 | biotin | BioLegend | 408004 |

| Antibodies used for transplant immune response monitoring | | | | |
|---|---|---|---|---|
| Antibody Target | Clone | Format | Supplier | Catalogue # |
| CD69 | H1.2F3 | BUV737 | BD Biosciences | 612793 |
| CD4 | GK1.5 | BUV805 | BD Biosciences | 612900 |
| PD-1 | 29F.1A12 | BV421 | BioLegend | 135218 |
| KLRG1 | 2F1/KLRG1 | BV510 | BioLegend | 138421 |
| CD44 | IM7 | BV605 | BioLegend | 103047 |
| Lag-3 | C9B7W | BV650 | BioLegend | 125227 |
| Tim-3 | RMT3-23 | BV711 | BioLegend | 119727 |
| CD62L | MEL-14 | BV785 | BioLegend | 104440 |
| CD8a | KT-15 | FITC | Invitrogen | MA5-16760 |
| CD90.2 | 53-2.1 | PerCPCy5.5 | BioLegend | 140322 |
| TIGIT | 1G9 | PEDazzle594 | BioLegend | 142110 |
| CD14 | Sa14-2 | PECy7 | BioLegend | 123316 |
| CD19 | 6D5 | PECy7 | BioLegend | 115520 |
| CD186 | SA051D1 | APC | BioLegend | 151106 |
| CD127 | A7R34 | APC/Cy7 | BioLegend | 135040 |

| Antibodies used for confocal imaging | | | | |
|---|---|---|---|---|
| Antibody Target | Clone | Format | Supplier | Catalogue # |
| CD31 | PECAM-1 | AF488 | BioLegend | 102414 |
| CD45 | 30-F11 | AF647 | BioLegend | 103124 |
| CK19 | EPNCIR127B | purified | Abcam | Ab133496 |
| H-2K$^{b}$ | Y-3 | purified | WEHI* | N/A |
| Rabbit IgG | Polyclonal | AF750 | Invitrogen | A21039 |
| Mouse IgG2b | RMG2b-1 | PE | BioLegend | 406708 |

| Antibodies used for immunoaffinity purification | | | | |
|---|---|---|---|---|
| Antibody Target | Clone | Format | Supplier | Catalogue # |
| H-2K$^{b}$ | K9-178 | purified | In-house | In-house |
| H-2K$^{b/k}$ | Y-3 | purified | In-house | In-house |
| H-2K$^{d}$ | SF1-1.1.10 | purified | In-house | In-house |
| H-2D$^{b}$ | 28-14-8 | purified | In-house | In-house |

*Antibody core, Walter & Eliza Hall Institute, Melbourne, Australia

Supplementary Table 6: Variable Window widths used for DIA acquisition

| | Start | End | Width |
|---|---|---|---|
| MS1 | 375 | 1000 | |
| DIA | 375 | 386.1 | 11.1 |
| DIA | 385.1 | 398.7 | 13.6 |
| DIA | 397.7 | 412 | 14.3 |
| DIA | 411 | 425.4 | 14.4 |
| DIA | 424.4 | 441 | 16.6 |
| DIA | 440 | 457 | 17 |
| DIA | 456 | 474.7 | 18.7 |
| DIA | 473.7 | 493.7 | 20 |
| DIA | 492.7 | 510.7 | 18 |
| DIA | 509.7 | 528.8 | 19.1 |
| DIA | 527.8 | 545.8 | 18 |
| DIA | 544.8 | 565.7 | 20.9 |
| DIA | 564.7 | 591.3 | 26.6 |
| DIA | 590.3 | 616.3 | 26 |
| DIA | 615.3 | 643.3 | 28 |
| DIA | 642.3 | 668.3 | 26 |
| DIA | 667.3 | 693.3 | 26 |
| DIA | 692.3 | 851.3 | 159 |
| DIA | 850 | 1000 | 150 |
